# Molecular archaeology of the human brain

**DOI:** 10.1101/598094

**Authors:** Joanna Kaczanowska, Florian Ganglberger, Bence Galik, Andreas Hess, Yoshan Moodley, Katja Bühler, Wulf Haubensak

**Author notes:** These authors contributed equally (Joanna Kaczanowska, Florian Ganglberger).

## Abstract

Mapping the origin of human cognitive traits typically relies on comparing behavioral or neuroanatomical features in human phylogeny. However, such studies rely on inferences from comparative relationships and do not incorporate neurogenetic information, as these approaches are restricted to experimentally accessible species. Here, we fused evolutionary genomics with human functional neuroanatomy to reconstruct the neurogenetic evolution of human brain functions more directly and comprehensively. Projecting genome-wide selection pressure (dN/dS ratios) in sets of chronologically ordered mammalian species onto the human brain reference space unmasked spatial patterns of cumulative neurogenetic selection and co-evolving brain networks from task-evoked functional MRI and functional neuroanatomy. Importantly, this evolutionary atlas allowed imputing functional features to archaic brains from extinct hominin genomes. These data suggest accelerated neurogenetic selection for language and verbal communication across all hominin lineages. In addition, the predictions identified strategic thought and decision making as the dominant traits that may have separated anatomically modern humans (AMH) from archaic hominins.

## Introduction

Exploring the origins of the human mind holds the anthropocentric promise of uncovering the traits that may set us apart from our closest relatives, alive or extinct. Various initiatives have provided valuable insight into the evolution of the extraordinary size, shape (*1*–*5*) and function (*6, 7*) of the AMH brain. Comparative neuroanatomy traced cognitive evolution from incremental neuroanatomical changes along the mammalian lineage (*8*–*13*). Typically, descriptive quantitative measures, such as increases in regional volumes, gene expression boundaries or cell numbers, serve as proxies for evolutionary selection (Methods Table 1). However, these changes primarily reflect structural organization of a single parameter at gross resolution and omit a functional network context. Therefore, these approaches rather indirectly trace the functional evolution of the brain, which occurs often with similarly sized brains or brain areas that show different functional organization and/or have undergone different evolutionary pressures. Moreover, comparative neuroanatomy does not trace the underlying neurogenetic evolution per se and is limited to experimentally accessible species, mapped at deep anatomical resolution. The latter is a problem for archaic hominins as only endocasts of skulls remain, which restricts insights to inferences from the brain surface. Another line of research in evolutionary genetics, which can analyze archaic material, infers differences in cognitive traits from genetic mutations (*14, 15*) in human phylogeny. While extremely informative, these studies typically interpret these mutations in isolation, omitting the compound effect of multi-genic co-evolution of functional brain networks.

Here, we propose that the selection pressure on the mammalian brain over evolutionary history left genomic signatures that can be mined for direct and deeper insight into the neurogenetic functional evolution across the human lineage. Genetic information has the key advantage that it not only contains genome-wide signatures of evolutionary events but also correlates with the mesoscale functional organization of brain networks (*16, 17*). Newly developed, highly parallel sequencing techniques have generated whole genomes of both extant (*18*) and extinct species (*19*– *21*) along the mouse-AMH evolutionary tree. Using genetic (*22*), connectomic (*23*) and behavioral or psychiatric (*24*) brain-data initiatives, we can now relate genetic features directly to functional brain networks and behavioral traits (*16, 17, 25, 26*). Importantly, these approaches can trace genetic and functional co-evolution directly within the AMH brain framework and do not rely on interpolating associations between analogous structures as in classical comparative strategies.

## Results

Here, we adapted such computational workflows (*17*) to systematically integrate genome-wide phylogenetic selection pressure into a brain-wide evolutionary atlas of the AMH brain. These resources allow reconstruction of the neurogenetic and functional evolution from the common ancestor with rodents, along the primate lineage, to bipedal hominins and humans.

Initially, this strategy required tracking genome-wide patterns of selection pressure on the mammalian phylogeny linking the mouse with AMH (Fig. 1A). This evolutionary framework covered eight major diversification episodes along the primate lineage leading to AMH - from common ancestry with rodents (mouse) through prosimians (bush baby), New World monkeys (marmoset), Old World monkeys (macaque), lesser apes (gibbon), great apes (chimpanzee) and finally the known extinct hominins Denisovan and Neanderthal. We inferred gene-wise selection pressure between successive pairwise species comparisons (PSCs, Fig. 1A bottom) by calculating the ratio of nonsynonymous to synonymous nucleotide substitutions (dN/dS) from pairwise alignments of protein-coding gene orthologs (data retrieved from Ensembl (*18*) and JBrowse (*27*), see Methods). The dN/dS ratio is a robust, gene-wise measure of selection pressure that is sensitive to evolutionary adaptation (*28, 29*) and does not rely on population data, which are mostly unavailable. dN/dS values are also a suitable proxy for overall functional selection and should also reflect the overall phenotypic contributions from co-evolving non-coding regulatory sequences, indels, and species-specific genes/duplications. Using brain-expressed gene sets built on PSCs homologs provides sufficient details to model multi-genic neurocognitive traits of AMH (*16*). Thus, in our PSC approach, dN/dS values should reliably indicate the most dominant genetic evolution that occurred after the split from each Last Common Ancestor (LCA). Therefore, the combination of phylogenetically ordered, pair-wise comparisons is a straightforward means of tracing selection pressure along evolutionary lines, in this case, along the human lineage.

**Fig. 1.**
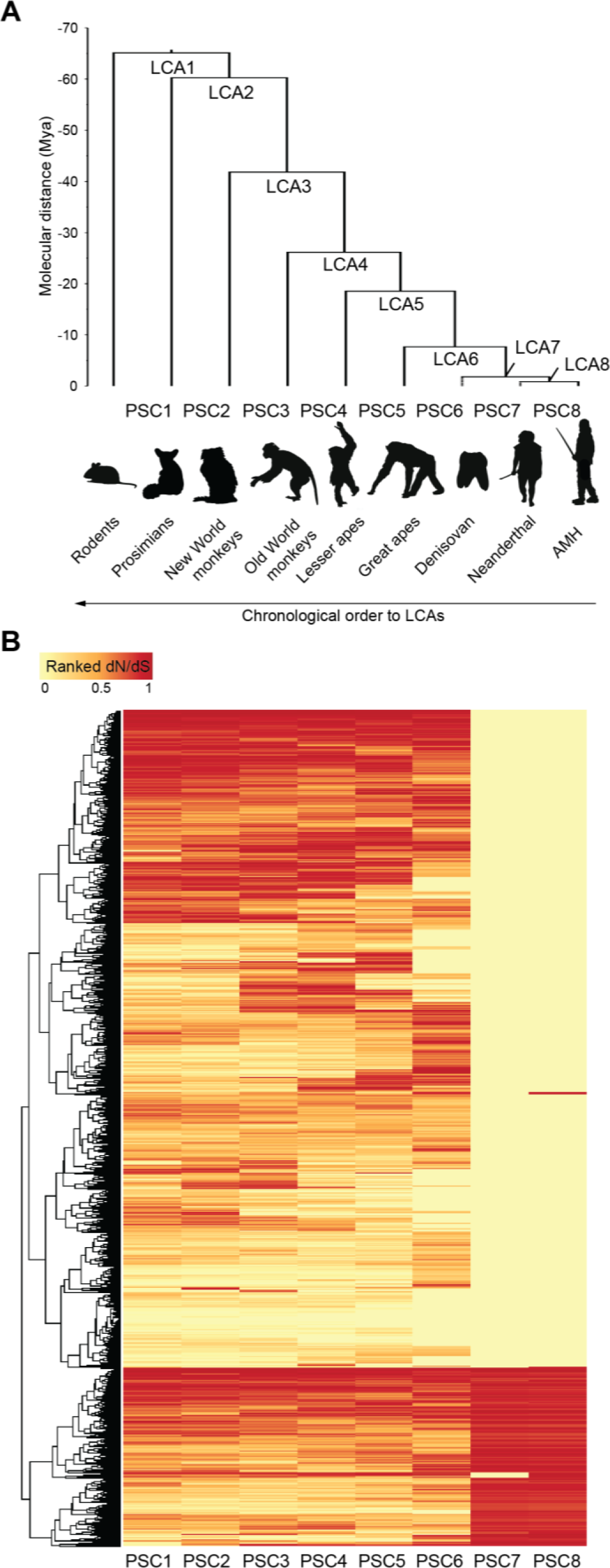
Reconstructing genomic selection pressure in the human lineage. (**A**) Phylogeny of mammalian species in the human branch, including extinct hominins (Denisovan and Neanderthal). Species were ordered by molecular distance based on mitochondrial genome sequences in millions of years ago (Mya). Tree nodes represent Last Common Ancestors (LCAs) of the consecutive species pairs. LCAs were first ordered based on chronological appearance in evolutionary history (LCA1-8), then by molecular distance to AMHs (LCA7, 8). Pairwise species comparisons (PSCs) continuously bridged mouse-to-AMH evolutionary history. (**B**) Hierarchical clustering of dN/dS values assigned to the respective LCA of a pairwise species comparison. dN/dS values were column-wise rank normalized for each PSC to eliminate bias due to different evolutionary times after the split from LCAs. Color code indicates normalized rank. Note that the binary like appearance of dN/dS spread for PSC7-PSC8 results from the evolutionary proximity of these species.

We then focused our analysis on neurocognitive functions by constraining the homolog pool to brain-expressed genes of the Allen Brain Atlas (ABA), a high resolution database of brain gene expression (Methods)(*22*). We thus obtained the continuous neuro-genetic evolution along the mammalian evolutionary sequence leading to AMHs using multiple species as reference. As expected, we observed diverse trends in selection pressure, as represented by several clusters of co-evolving genes (Fig. 1B). To trace selective forces, our further analyses utilized the rank-ordered dN/dS genes found in the ABA within each PSC to capture synergistic, multi-genic effects of coevolving genes in brain networks.

Fusing these genetic data with the AMH brain gene expression atlas approximates synergies in molecular evolution directly onto AMH neuroanatomy. So, we first generated cumulative neuroanatomical dN/dS weighted maps for every PSC (Fig. 2) in the AMH reference brain at subregional mesoscale resolution (*23*) using the ABA gene expression database (*22*). Then, we combined these data (Methods, Computational neuroanatomy, Generating evolutionary maps) into a temporo-spatial map of hot spots of high selection pressure (Fig. 2). Consequently, the resulting evolutionary landscape traced the phylogenetic history of the AMH brain through successive LCAs. We note this map is purely genetically driven and largely independent of establishing phylogenetic homology among brain areas. We found that the load of the most highly selected genes accumulated in different neuroanatomical areas for different PSCs (see the Methods Table 2 for anatomical-neurocognitive annotation). There was a gradual shift in selection pressure, acting more on subcortical regions in PSCs 1-5, then increasing over cortical regions (PSCs 6-8). Basal forebrain, basal ganglia, brain stem and cerebellum were under selection pressure in all comparisons. Different thalamic nuclei, hippocampal, amygdalar and cortical parts selection varied between PSCs (Fig. 2).

**Fig. 2.**
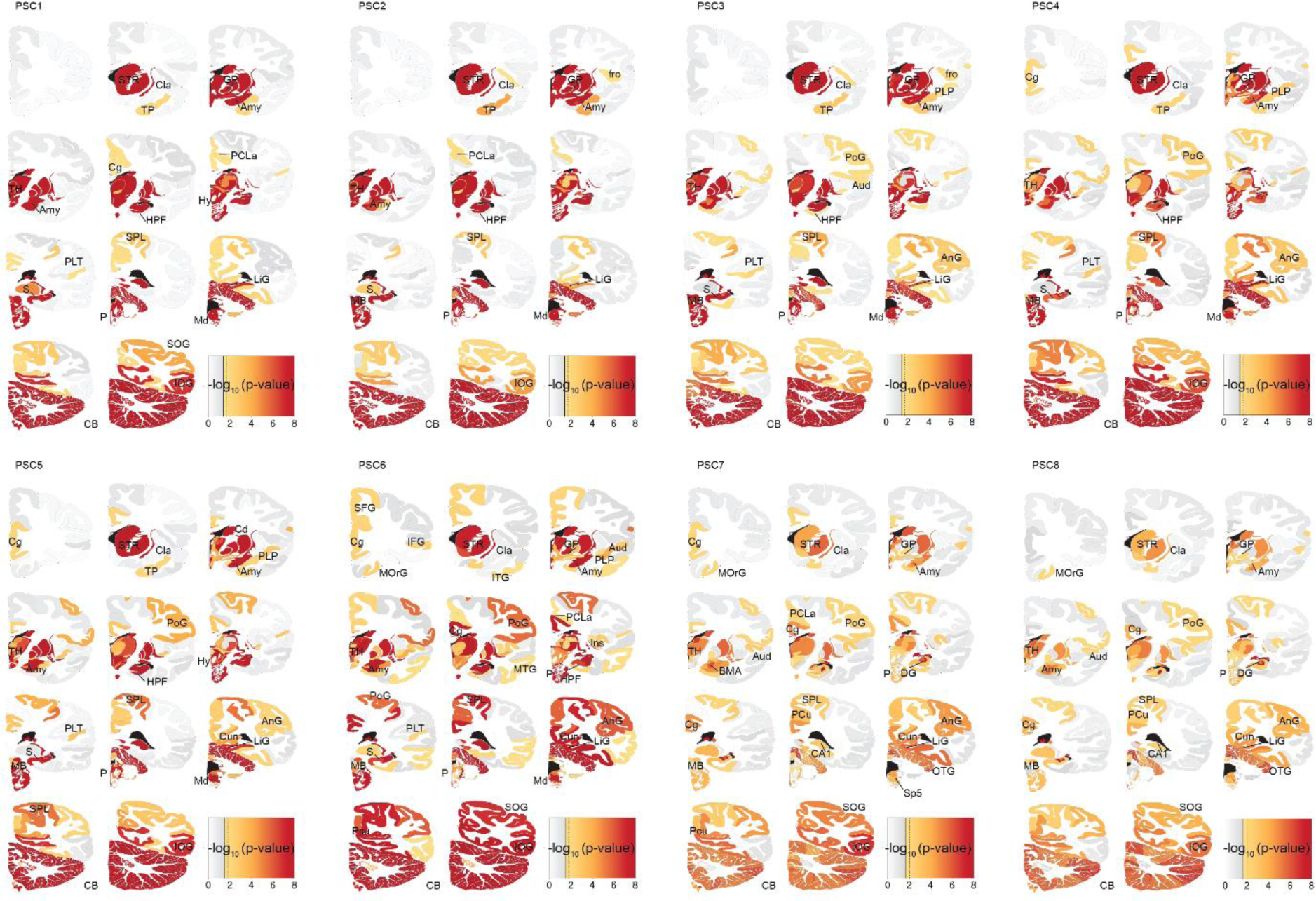
Individual genetic maps for PSC1-8 projected onto the AMH reference brain. Each map represents cumulative load of ranked dN/dS genes. Regions show the most significant p-value of their individual biopsy-sites. Significant regions are indicated with color scale, non-significant with grey scale. Amy – amygdala, Ang – angular gyrus, Aud – auditory cortex, BMA – basomedial amygdala, CA1 – cornus ammoni 1, Cg – cingulate cortex, Cd – caudate, Cun – cuneus, DG – dentate gyrus, FuG – fusiform gyrus, GP – globus pallidus, GRe – gyrus rectus, Hy – hypothalamus, IFG – inferior frontal gyrus, Ins – insula, IOG – inferior occipital gyrus, HPF – hippocampal formation, ITG – inferior temporal gyrus, LiG – lingual gyrus, MB – midbrain, Md – medulla, MFG – medial frontal gyrus, MOrG – medial orbital gyrus, MTG – middle temporal gyrus, OP – occipital pole, OTG – occipito-temporal gyrus, P – pons, PCLa – paracentral lobule anterior part, PLP – planum polare, PLT – planum temporale, PCu – precuneus, PoG – postcentral gyrus, S – subiculum, SFG – superior frontal gyrus, SMG – supramarginal gyrus, SOG – superior occipital gyrus, Sp5 – spinal trigeminal nucleus, SPL – superior parietal lobule, STR - striatum, TH – thalamus, TP – temporal pole.

To highlight such changes lineage-wide, we computed a combined dN/dS atlas, weighted by the temporal profile of selection pressure amongst the PSCs. In general, subcortical regions (e.g. the striatum, basal forebrain and brain stem) showed signatures of selection from rodent to primates (e.g. Fig. 3, PSC1-5, dark blue-yellow). Notably, one structure evolving at the highest genetic rate early in the evolutionary transition to primates was the claustrum (Fig. 3, PSC1, dark blue), which is a central element in a framework for consciousness (*30*).

**Fig. 3.**
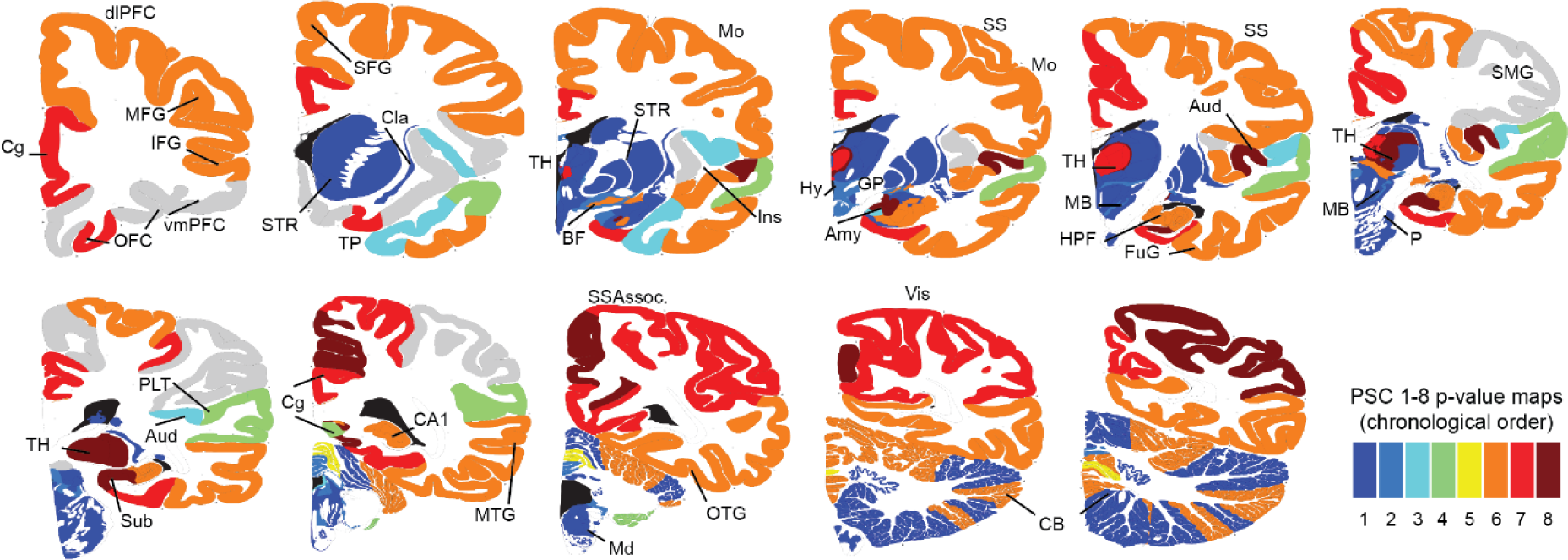
The compound evolutionary landscape of selection pressure in the AMH brain. Color indicates the PSC with the significant absolute peak in cumulative gene expression in a given brain region. Amy – amygdala, Aud – auditory cortex, BF – basal forebrain, CA1 – cornus ammoni 1, CB – cerebellum, Cg – cingulate cortex, CnF – cuneiform nucleus, FuG – fusiform gyrus, HPF – hippocampal formation, Hy – hypothalamus, Ins – insula, MB – midbrain, Mo – motor cortex, MTG – middle temporal gyrus, OFC – orbitofrontal cortex, OTG – occipito-temporal gyrus, P – pons, dlPFC/vmPFC – dorsolateral/ventromedial prefrontal cortex, SS(Assoc.) – somatosensory (associative) cortex, STR – striatum, Sub – subiculum, TH – thalamus, TP – temporal pole, Vis – visual cortex.

In addition, our approach can track neuroanatomical selection across evolutionary transitions from genomes where brain-wide (non-surface) neuroanatomical data is out of reach. In contrast to early PSCs, traces of late selection pressure in hominid evolution (great apes and hominins) predominantly accumulated in the cortex (Fig. 3, PSC6-8, orange-dark red). Accelerated selections within archaic hominins (Fig. 3, PSC7, light red) and AMHs (Fig. 3, PSC8, dark red) localized mainly within sub-regions of prefrontal, temporal or somatosensory associative cortex, but also in the thalamus and amygdala. It is striking that all these structures subserve higher cognitive functions, spiritual beliefs (*31*), language (speech comprehension (Wernicke’s area) and production (Broca’s area), reading (angular gyrus (*32*)) or facial recognition (*33*). Brain structures with the highest selection pressure marking divergence of Neanderthal and AMH involved cortical (auditory, parietal – especially precuneus, occipito-temporal), subiculum, thalamic and amygdalar areas (Fig. 3, PSC8, dark red). Collectively, these evolutionary maps are in line with comparative data from primate cognitive evolution and provide a neuroanatomical framework for many previous inferences about cognitive evolution in hominins (Methods Table 1).

Since these evolutionary brain maps suggested that neurogenetic selection follows specific functional needs, we utilized data from functional brain networks initiatives, in conjunction with evolutionary neuroanatomy data, to directly trace this functional evolution. To link temporal evolutionary patterns with spatially registered brain function, we performed a temporo-spatial analysis by GABi biclustering (*34*). Specifically, we clustered genes based on their ranked dN/dS values with their spatial expression correlated to multiple AMH functional brain networks. These functional networks consisted of task-evoked functional MRI (TNs) (Fig. S1, Methods Table 3, note TNs in left and associated traits in right columns) from the Human Connectome Project (HCP) (*24*) and networks for specific brain functions from literature (FNs) (Fig. S1, Methods Table 4, note FNs in left and associated traits in right columns). We tuned GABi biclustering with custom criteria to identify the largest diverse biclusters of co-evolving genes (i.e. genes that have >=0.90 ranked dN/dS over multiple PSCs) with high specificity for functional networks (i.e. genes with high network correlation for the same networks) from Methods Table 3 & 4 (Fig. 4A). This strategy yielded statistically stable biclusters (Methods), which could be further organized into modules (M1-7). Importantly, each module containing the same PSCs could then be chronologically ordered (Fig. 4A). To visualize these neurogenetic co-evolution networks in the brain, we generated a 3D representation of the most prominent networks within each bicluster module (Fig. 4B). These data illustrate the evolutionary selection of networks for attention, consciousness and emotion during early primate evolution and for language, working memory, strategic thought, and motor control in hominin and AMH lineages. In clusters representing the earliest LCAs in the mammalian lineage, most selection pressure accumulated in networks for attention (visuospatial and fronto-parietal FNs) (Fig. 4B, Module 1). In addition to attention (salience FN), selection pressure then extended to networks for consciousness (default mode FN (DMN)) and emotion (EM-FACES TN) (Fig. 4B, Module 2). Subsequent evolution selected for motor control (sensorimotor FN), higher cognition (central executive (CEN) and RE-AVG networks) and attention (ventral attention FN) (Fig. 4B, Module 3, 4 and 5). In contrast, great ape-hominin diversification selected for theory of mind (SO-TOM TN), cognitive emotional control (prefrontal-amygdala FN), motivation (prefrontal-accumbens FN) and awareness (dorsal attention FN) (Fig. 4B, Module 6). Most strikingly, in hominins language (LA-AVG TN), working memory (WM-FACE, WM-REST TNs), strategic thought (GA-REWARD, GA-AVG TNs) and motor control (MO-RH-LH, MO-RF-LF TNs) emerged as the traits under highest evolutionary pressure (Fig. 4B, Module 7).

**Fig. 4.**
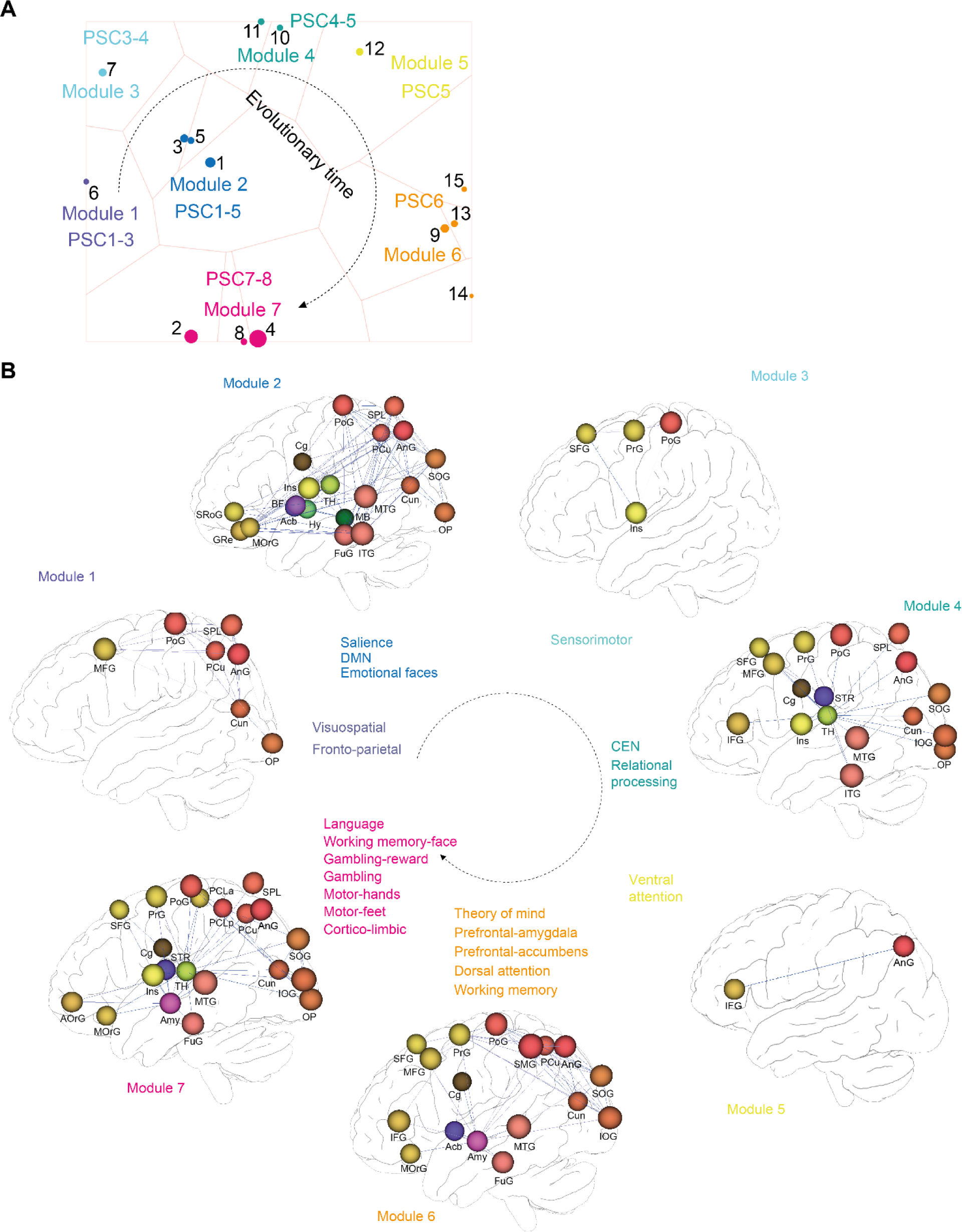
Evolution of neurocognitive tasks in AMH brain networks. (**A**) Biclustering of functional networks (Methods Table 3 & 4) and highest dN/dS values (at 0.90 rank cut-off) across time (PSC1-8). Biclusters embedded in a 2D space using t-SNE with bicluster overlap as distance measure to trace coevolving functional brain networks modules in the brain over time. Individual clusters (1-15) highlight gene sets and associated networks with similar evolutionary history. Circle size corresponds to the cluster size. Closely related clusters are assigned to common modules (M1-M7). (**B**) 3D visualization of top ranked functional brain networks from A, highlighting the highest differences between PSCs corresponding to the associated functional networks. The nodes represent anatomical regions, the edges gene-expression correlations of the respective networks. Note that for visibility, not all networks components are shown in each case. Acb – nucleus accumbens, Amy – amygdala, Ang – angular gyrus, AorG – anterior orbital gyrus, BF – basal forebrain, Cg – cingulate cortex, Cun – cuneus, FuG – fusiform gyrus, GRe – gyrus rectus, Hy – hypothalamus, IFG – inferior frontal gyrus, Ins – insula, IOG – inferior occipital gyrus, ITG – inferior temporal gyrus, MB – midbrain, MFG – medial frontal gyrus, MOrG – medial orbital gyrus, MTG – middle temporal gyrus, OP – occipital pole, PCLa/p – paracentral lobule anterior/posterior part, PCu – precuneus, PrG – precentral gyrus, PoG – postcentral gyrus, SFG – superior frontal gyrus, SMG – supramarginal gyrus, SOG – superior occipital gyrus, SPL – superior parietal lobule, SRoG – superior rostral gyrus, STR - striatum, TH – thalamus.

The biclustering approach directly related neurocognitive evolution to functional co-evolution at the genetic level, identifying gene sets potentially driving the observed functional evolution within a given brain network. Among the genes correlating with language task, we identified a key gene FOXP1, which has been linked to speech impairment and intellectual disability (*35, 36*), and is the closest homolog of FOXP2, which shows similar expression patterns in the brain. While FOXP2 itself differs between chimpanzees and hominins (*14*), it is too similar among hominins for detection here. Several other genes from this cluster are associated with cognitive disabilities and neuroanatomical abnormalities like macrocephaly (DVL1(*37*)), microcephaly (MCPH1 (*38*)), autism (SHANK2 (*39*)), bipolar disorder-and circadian rhythm-linked gene CLOCK (*40*), which was also identified as a hub and enriched gene in AMH-specific transcriptional networks of the frontal pole (*41*). In addition, we performed Ingenuity Pathway Analysis (IPA) (*42*) on all clusters to gain a deeper insight into the genetic composition of the clustered networks. We focused our IPA analysis on Diseases & Functions in Nervous System, to find the functional associations within molecular-to-system-level neurobiology. IPA results show multiple behavioral, psychiatric, neurophysiological, structural and molecular functions which coincide with the diversification periods analyzed here. Many associations point to memory, cognition, brain size, and organization of synapses. The highlights of this study are shown in Table 1. Interestingly, genes in modules M1-4 are linked to neurodegenerative diseases. Module 6 covering chimp-Denisovan selection pressure shows associations with increasing brain size. The genetic component of Module 7 (PSCs 7&8), which correlates with language, working memory and strategic thought (Fig. 4B, Module 7), was associated with memory and learning. Complete analysis results can be found in Supplementary Tables 1-7.

**Table 1.**
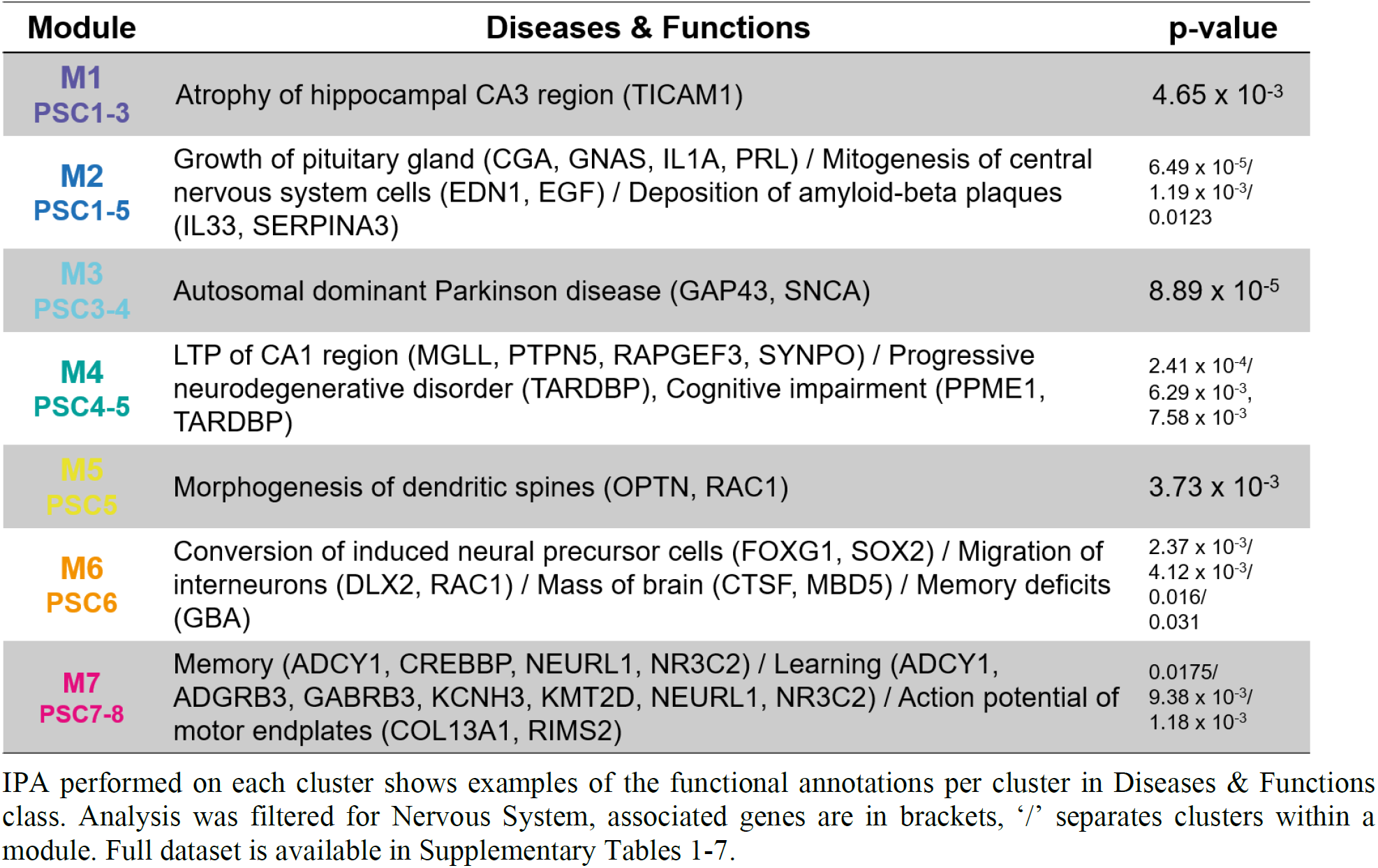
Summary of mammalian lineage Ingenuity Pathway Analysis (IPA).

To explore archaic-to-AMH brain evolution more deeply, we next ran the biclustering solely on the hominid data, which includes PSCs 6-8 (Fig. 5). With this, we achieved a better segregation of functional features under selection pressure among the closest relatives of AMH. The method delineated clusters of functional networks selected between great apes and hominins or among anatomically modern and archaic humans, which pointed to the similar findings as in the mammalian lineage approach (e.g. language, working memory and motor functions in hominins). Importantly, we obtained a PSC8 specific cluster highly correlating with the GA-REWARD TN, indicating strategic thought as the fastest evolving trait between Neanderthals and AMHs. Paralleling these patterns of functional neuroanatomical evolution, Ingenuity analysis (Table 2) functionally associated the genetic clusters with cognitive traits (working memory, learning, cognition) and psychiatric disorders (schizophrenia, impulsivity), among others (Supplementary Tables 8-10).

**Table 2.**
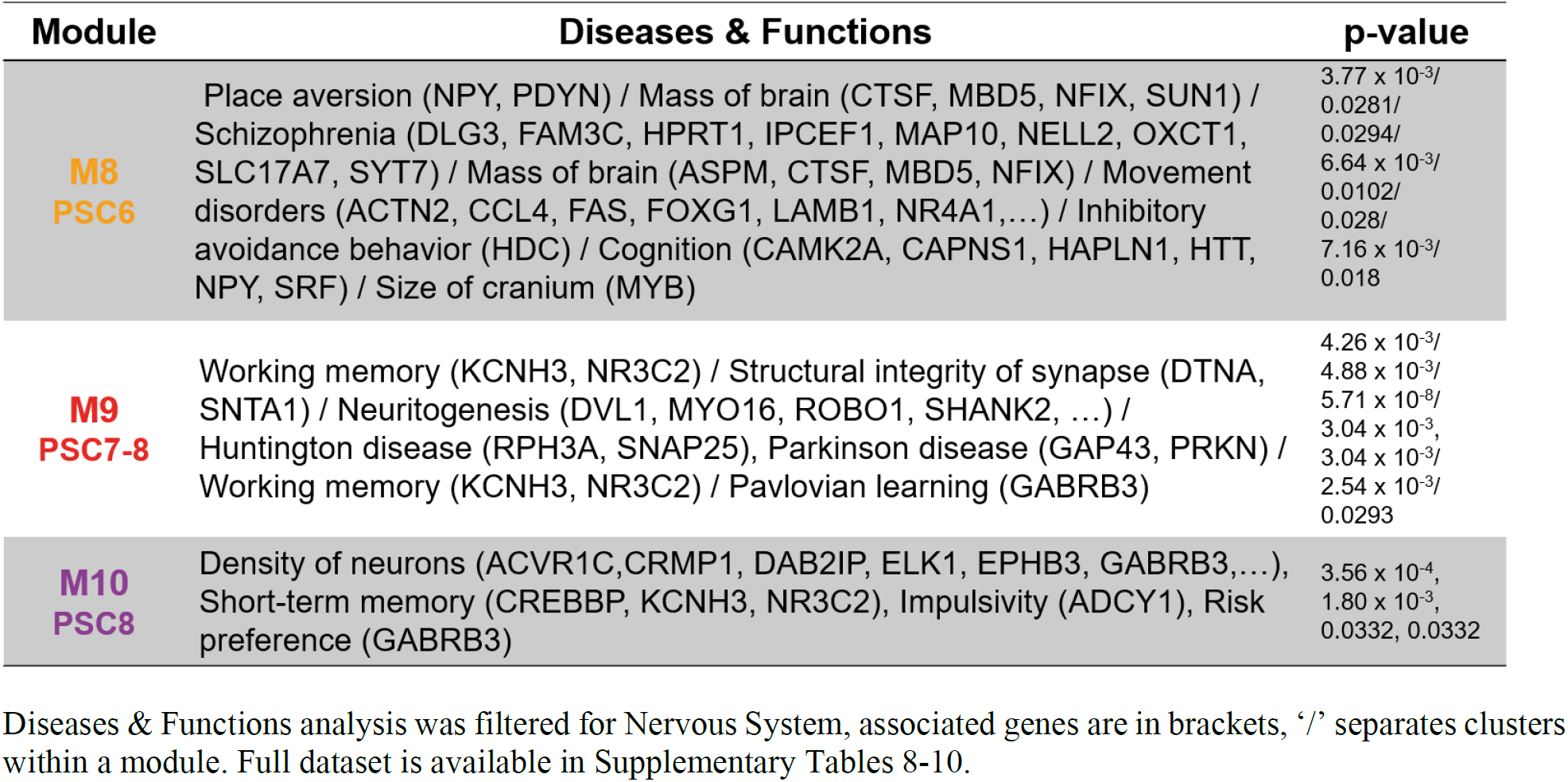
Summary of hominid Ingenuity Pathway Analysis (IPA).

| Module | Diseases & Functions | p-value |
| --- | --- | --- |
| **M8**<br>**PSC6** | Place aversion (NPY, PDYN) / Mass of brain (CTSF, MBD5, NFIX, SUN1) / Schizophrenia (DLG3, FAM3C, HPRT1, IPCEF1, MAP10, NELL2, OXCT1, SLC17A7, SYT7) / Mass of brain (ASPM, CTSF, MBD5, NFIX) / Movement disorders (ACTN2, CCL4, FAS, FOXP1, LAMB1, NR4A1,...) / Inhibitory avoidance behavior (HDC) / Cognition (CAMK2A, CAPNS1, HAPLN1, HTT, NPY, SRF) / Size of cranium (MYB) | 3.77 x 10<sup>-3</sup>/<br>0.0281/<br>0.0294/<br>6.64 x 10<sup>-3</sup>/<br>0.0102/<br>0.028/<br>7.16 x 10<sup>-3</sup>/<br>0.018 |
| **M9**<br>**PSC7-8** | Working memory (KCNH3, NR3C2) / Structural integrity of synapse (DTNA, SNTA1) / Neuritogenesis (DVL1, MYO16, ROBO1, SHANK2, ...) / Huntington disease (RPH3A, SNAP25), Parkinson disease (GAP43, PRKN) / Working memory (KCNH3, NR3C2) / Pavlovian learning (GABRB3) | 4.26 x 10<sup>-3</sup>/<br>4.88 x 10<sup>-3</sup>/<br>5.71 x 10<sup>-8</sup>/<br>3.04 x 10<sup>-3</sup>,<br>3.04 x 10<sup>-3</sup>/<br>2.54 x 10<sup>-3</sup>/<br>0.0293 |
| **M10**<br>**PSC8** | Density of neurons (ACVR1C, CRMP1, DAB2IP, ELK1, EPHB3, GABRB3,...), Short-term memory (CREBBP, KCNH3, NR3C2), Impulsivity (ADCY1), Risk preference (GABRB3) | 3.56 x 10<sup>-4</sup>,<br>1.80 x 10<sup>-3</sup>,<br>0.0332, 0.0332 |

Diseases & Functions analysis was filtered for Nervous System, associated genes are in brackets, ‘/’ separates clusters within a module. Full dataset is available in Supplementary Tables 8-10.

**Fig. 5.**
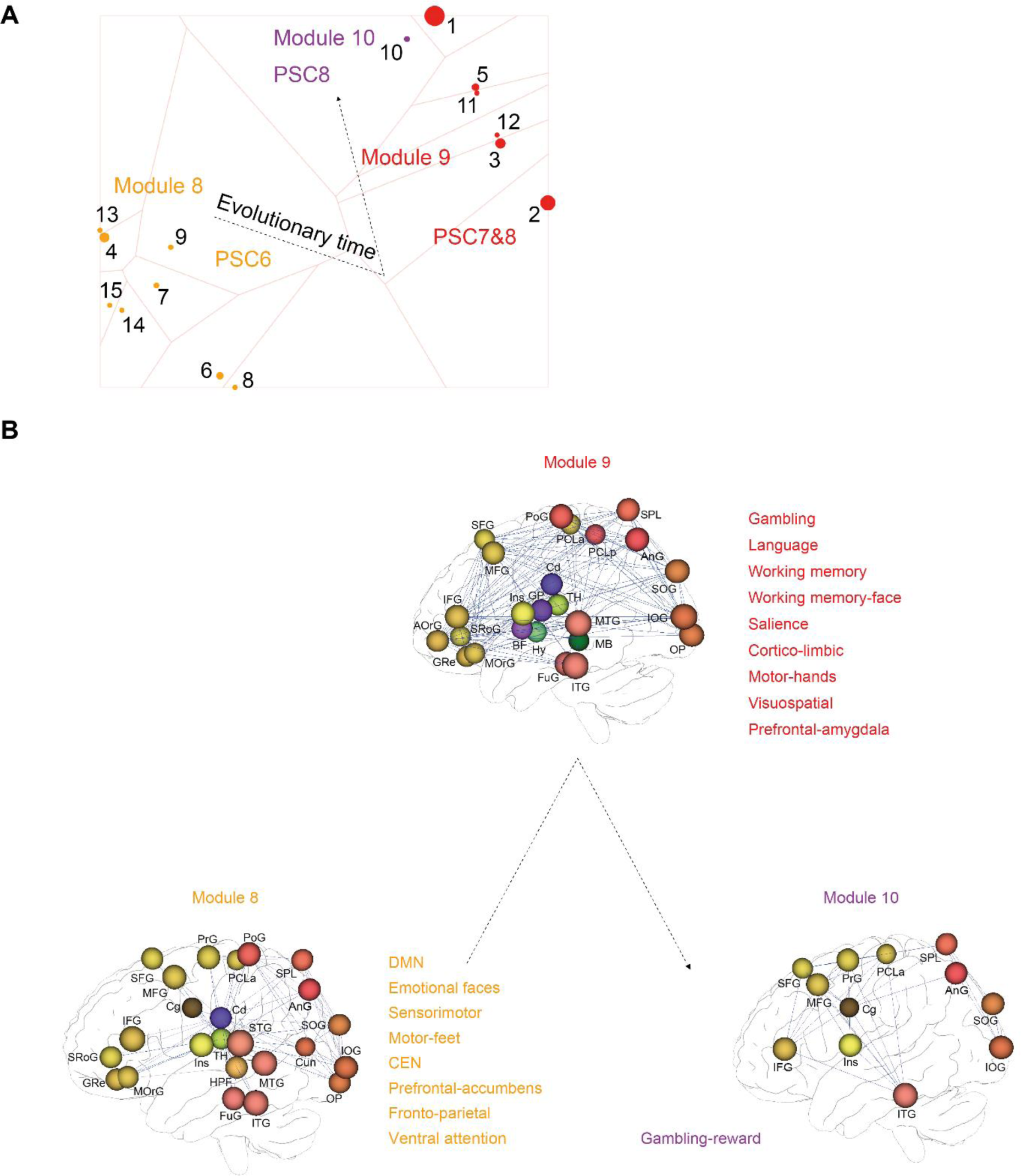
Hominin-specific distribution of selection pressure in AMH brain networks. (**A**) Biclustering of functional networks as in Fig. 4 performed solely on the hominid branch (PSCs 6-8). Similarly, closely related clusters are assigned to common modules (M8-M10). (**B**) 3D visualization of top ranked functional brain networks from A, highlighting the highest differences between PSCs corresponding to the associated functional networks. The nodes represent anatomical regions, the edges gene-expression correlations of the respective networks. Note that for visibility, not all networks components are shown in each case. Acb – nucleus accumbens, Ang – angular gyrus, AorG – anterior orbital gyrus, BF – basal forebrain, Cd – caudate, Cg – cingulate cortex, Cun – cuneus, FuG – fusiform gyrus, GP – globus pallidus, GRe – gyrus rectus, HPF – hippocampal formation, Hy – hypothalamus, IFG – inferior frontal gyrus, Ins – insula, IOG – inferior occipital gyrus, ITG – inferior temporal gyrus, MB – midbrain, MFG – medial frontal gyrus, MOrG – medial orbital gyrus, MTG – middle temporal gyrus, OP – occipital pole, PCLa/p – paracentral lobule anterior/posterior part, PCu – precuneus, PrG – precentral gyrus, PoG – postcentral gyrus, SFG – superior frontal gyrus, SMG – supramarginal gyrus, SOG – superior occipital gyrus, SPL – superior parietal lobule, SRoG – superior rostral gyrus, STR - striatum, TH – thalamus.

Together, these findings provide a critical neurogenetic framework explaining evolutionary ethology, archeological records and genetic data (Methods Table 1). Importantly, they identify neurocognitive traits critical in recent hominin evolution and the underlying gene sets driving their evolution.

## Discussion

This study integrates rates of genetic evolution along the mammalian/primate lineage into an evolutionary atlas of the AMH brain. We exploited recent big data initiatives from genomics and brain functions to explore evolutionary events that ultimately shaped the mind and behavior of AMHs. Our findings utilize an innovative *in silico* approach, fusing genomic information with brain data to associate multi-genic behavioral traits with brain functional networks (*43*). We could map the cumulative load of evolutionary-weighted genes into AMH functional brain networks, genome-and brain-wide, which reconstructed relative selection pressures on brain functions at each evolutionary ‘transition’ from mouse to AMH.

This computational strategy uses and explores compound neurogenetic effects on brain networks, which are untraceable when studied in isolation. From the genetic perspective, the evolution of functional traits is inherently multi-genic. Therefore, single gene studies in mouse or organoid models, while mechanistically insightful (*44*), cannot easily assess such genetic synergies on cognitive traits emerging from brain-wide functional networks. This makes it difficult to unmask the evolution of complex cognitive traits and to identify potential candidate gene sets driving evolution in this way. Our study will therefore complement such functional evolutionary genetic models with an atlas of coevolving genes and functional neuroanatomical networks. Future functional exploration of these gene sets and networks in suitable experimental systems should reveal potential mechanisms driving the neurocognitive divergence between the respective species.

We showcase an approach for reconstructing the evolutionary history of functional selection using the genetic remains of species long extinct by projecting compound phylogenetic evolutionary weights onto a functional reference framework. This strategy may be useful to functionally explore biological systems not available for traditional experimental studies. Of note, the straightforward evolutionary genetic analysis could be refined by including an extended phylogeny into the computation of dN/dS values and/or other evolutionary measures, possibly leading to even deeper understanding of neurogenetic evolution. This notwithstanding, our workflow unraveled neurogenetic selection for complex neurocognitive traits in archaic hominin brains, like social interaction and communication, hand motor control, working memory for faces along with symbolic thought and abstract thinking (*33*). These findings delineate a critical neurogenetic framework for archaic cognitive abilities inferred from ancient art (*45, 46*). Strikingly, our computational neuro-archaeological data provide initial neurogenetic evidence that all ancient hominin and AMH lineages evolved networks for language. The active selection for language networks strongly suggests that all these archaic hominins could speak. Furthermore, this indicates that verbal communication evolved in the LCA to all three hominin species (i.e. *Homo erectus*) several hundred thousand years ago, adding critical insights to a longstanding debate about the timing and origin of human language (*47*). Surprisingly, the transition to AMHs further accelerated evolution of reward-related decision making and strategic thought as those features most prominently separating us from archaic hominins. It is tempting to speculate that these abilities may have contributed to the selective advantage for evolutionary success of anatomically modern humans.

## Acknowledgments

We thank Shiva Alemzadeh for bicluster visualization. We thank Attila Gyenesei for supervising the computation of evolutionary genetic data. We thank Life Science Editors for editing assistance.;

## Author contributions

JK designed and interpreted functional genetic analysis and computational neuroanatomy and wrote the manuscript. FG designed and performed computational neuroanatomy and wrote the manuscript. BG and YM designed and performed evolutionary genetic analyses. AH designed computational neuroanatomy. KB designed and supervised computational neuroanatomy. WH initiated the study, designed and interpreted the computational neuroanatomy and wrote the manuscript. All authors contributed to writing and commented on the manuscript.;

## Data and materials availability

All data were analyzed with either published or custom code. Sources of published code are identified in the Method section.

## Methods

### Genetic analysis

#### Mammalian lineage setup

For our evolutionary analysis of the primate lineage, we selected nine species including the mouse (*Mus musculus*), bush baby (*Otolemur garnetti*), marmoset (*Callithrix jacchus*), macaque (*Macaca mulatta*), gibbon (*Nomascus leucogenys*), chimpanzee (*Pan troglodytes*), extinct hominins Denisovan (*Homo Denisovan*) and Neanderthal (*Homo neanderthalensis*) and anatomically modern human (AMH, *Homo sapiens*) (Methods Table 1).

To determine the general evolutionary relationship among the selected species, we reconstructed a phylogenetic tree from mitochondrial genomes using a Bayesian approach in BEAST 2.5 (*48*). First, the best nucleotide substitution model was determined using JMODELTEST (*49*). We then implemented this model in BEAST. To account for variable rates of evolution among different primate lineages, we used a relaxed lognormal prior on the clock rate. We used three independent normally-distributed and soft-bounded calibration priors (after Perez et al. 2013) (*50*) to place a timeframe onto our phylogeny. We used a AMH-chimpanzee mean divergence of 7.8 Myr (SD 1.2 Myr), and Old World monkey-ape divergence of 28 Myr (SD 3 Myr) and lastly we placed a 60 Myr mean (SD 2.8 Myr) time for the coalescence of all primate lineages. This fully parameterized model was run five times, each time for 200 million simulations, logging parameters every 20,000 steps, and discarding the first 20% as burn-in. MCMC convergence was assessed by viewing MCMC traces directly and by ESS values in TRACER 1.6(*51*). A maximum clade credibility tree was calculated and annotated in Figtree 1.4 (*52*).

The results of the mt-derived phylogeny, LCA and PSC order is given in Fig. 1A. Note that, formally, Denisovans and Neanderthals split after the split from the human ancestor. However, mtDNA derived phylogeny (Fig. 1A) places Neanderthal closer to AMH (*53*) and support a Chimp-to-Denisovan-Neanderthal-AMH species order.

**Methods Table 1.** Main evolutionary characteristics of analyzed taxa.

|  | *Mus musculus* | *Otolemur garnettii* | *Callithrix jacchus* | *Macaca mulatta* | *Nomascus leucogenys* | *Pan troglodytes* | *Homo denisovan* | *Homo neanderthalensis* | *Homo sapiens* |
| --- | --- | --- | --- | --- | --- | --- | --- | --- | --- |
| Brain size (cc) | 0.5 | 10.87 (54) | 7.77 (54) | 86.55 (55)* | 82.30 (56) | 400.90 (54) | 1200 (57) | 1473.46 (57) | 1473.84 (57) |
| Encephalization quotient (EQ=Ea/Ee) | 0.5 (58) | 1.25 (54)* | 1.70 (54)* | 2.10 (58) | 2.30 (58)* | 2.91 (54)* | no data available | no data available | 7.59 (58)* |
| Lifestyle | nocturnal territorial | nocturnal territorial sedentary | diurnal motile | diurnal motile/ sedentary | diurnal territorial | diurnal motile and sedentary | no data available | forager/hunter-gatherers (59) art and culture (45, 46) | territorial colonial nomadic/sedentary forager/hunter-gatherers urban agricultural art and culture |
| Diet | omnivorous | omnivorous | herbivorous | omnivorous | herbivorous | omnivorous | no data available | Omnivorous (59) | omnivorous |
| Social behavior | polygamous hierarchical dominance vocal communication | vocal communication scent markings | groups may include mated polyandrous groups nonmaternal infant care tactile, chemical communication | polygynadrous hierarchical dominance visual, tactile, acoustic and chemical communication | hierarchical dominance female dominance vocal communication | hierarchical dominance polygynadrous cooperative breeder visual, vocal, tactile, chemical communication | no data available | no data available | monogamous cooperative breeder hierarchical dominance language, visual, tactile communication |
| Group size/type | small | solitary | 2-13 individuals | 80-100 members | small families | fusion-fission society | no data available | small, isolated groups(60) | complex societies |
| Neuro-cognitive features |  |  |  |  | Advanced vocal communication (61–63) |  | no data available | Language (64–66) |  |
|  |  |  |  |  | Social behavior (67)<br>Cooperation (68, 69) |  |  | Social cognition (70) | Social learning and culture (71, 72) |
|  | Higher cognitive functions (73, 74)<br>Perceptual decision making (75)<br>Social decision making (76)<br>Sequential learning (77) |  |  |  |  |  |  | Higher intelligence (working memory, abstract thinking (78)) | Higher intelligence (tool use and production, abstract thinking, metacognition, theory of mind, social learning (58)) |
|  | Visual processing and visual attention (79–82) |  |  |  |  |  |  | Visuo-spatial integration (83, 84) | Spatio-temporal visual integration (85, 86) |
|  | Sensorimotor system (87) |  |  |  |  |  |  | Olfaction (88) |  |

Brain size given in cubic cm (cc), Encephalization quotient (EQ) defined as the ratio between actual brain mass (Ea) and brain mass predicted for an animal of given size (Ee), \* mean value from published samples. Descriptive species data based on https://animaldiversity.org. Blue indicates neurocognitive features inferred from endocasts, gross neuroanatomy or archeological remains.

#### Data collection

Pre-calculated non-synonymous (dN) and synonymous (dS) nucleotide substitutions for all homologous genes for the selected species were downloaded from Ensembl (version=78) (*89*) using the Bioconductor BiomaRt package (version 2.18.0) (*90*) except for Neanderthal and Denisovan genes. The pairwise values, based on the assembled lineage, were also obtained from Ensembl. For instance, bush baby vs. marmoset and marmoset vs. macaque. Other available entries like Ensembl gene IDs, Entrez IDs and HGNC symbols were downloaded for all genes for all species.

To obtain the dN/dS values for ancient hominins vs. chimpanzee and AMH, all protein coding genes for these species were downloaded from Ensembl (*18*). High coverage Denisovan genome alignment against the AMH (hg19/GRCh37) reference genome was downloaded from the http://cdna.eva.mpg.de/denisova/alignments/ repository. The data necessary for Neanderthal-AMH dN/dS calculation was gathered form the high coverage variant list from the Vindija Neanderthal genome (*91*).

#### Ancient hominin gene generation

Two different approaches were used to generate the AMH homolog, Denisovan and Neanderthal genes without any missing values.

The approach applied to the Denisovan dataset was based on mapping reads in BAM format to the AMH genome, and the data was segregated by chromosomes. Consensus sequences for Denisovan chromosomes were created using SAMtools mpileup (version 1.3.1) (*92*) and BCFtools (version 1.3) (*93*) utilities. Genes were extracted based on the hg19/GRCh37 Ensembl annotation (ftp://ftp.ensembl.org/pub/, version=78) using an in-house R script. The hominin genes were renamed using the AMH homolog Ensembl gene IDs for easier identification during the later processes.

To retrieve gene homologs from Vindija dataset, high coverage Neanderthal variants were downloaded in VCF format, and only SNPs predicted in protein coding genes were processed. Then, the Neanderthal genes were generated by exchanging AMH nucleotides to the high coverage SNPs in the corresponding AMH reference genes using the SNPs in the coding regions and its coordinates.

#### dN/dS calculation

For Ensembl downloaded dN and dS data, dN/dS values were calculated using a simple division. For ancient hominin vs. chimpanzee and AMH values, we downloaded and pairwise aligned codons from previously generated gene sequences using PRANK (v.140603) (*94*). Next, additional ratios were calculated using codeml tool from PAML package (version 4.9) (*95*) applying basic model.

#### Table organization

The combined dN/dS table was built from all dN/dS values and the corresponding Ensembl genes and Entrez IDs. The results from the mouse to AMH lineage including hominins were organized in the before mentioned order (Fig. 1A, Methods Table 1). The first mouse vs. bush baby step was used as the basis of the table. The values for the second step (bush baby vs. marmoset) were collected based on the gene IDs from the previous step. In this case, the bush baby gene IDs were used to organize the following column containing the results for marmoset genes. The same approach was applied in the following steps until the Neanderthal vs. AMH values.

### Computational neuroanatomy

#### Table preparation

To compute the evolutionary signatures in the mammalian brain, we used the dN/dS lineage table from above (Methods, Genetic data, Table organization) with 37173 genes/rows along 8 PSCs/columns (Pairwise Species Comparisons). We next sought to generate a brain gene set compatible across the Allen Brain Atlas platform (mouse and AMH) to be most versatile for this and future applications. To this end, we restricted genes to those present in both Allen Mouse Brain Atlas (AMBA) and Allen Human Brain Atlas (AHBA). We merged this table with spatial gene expression data from the AHBA via gene Entrez ID. We avoided merging duplicate entries with AMBA/AHBA by conflating the table rows for unique mouse and AMH Entrez ID combinations (=mouse/AMH homologues).

For homologue genes dN/dS were averaged. We omitted infinite dN/dS ratios (dS=0) for this purpose (*96*). We further removed rows with all dN/dS ratios=0, since static absent selection pressure was not relevant for our analysis. After filtering the table for genes with spatial gene expression available in AMBA/AHBA, we ended up with 10445 rows.

Wolf et al. (*97*) states that mean dN/dS between two species decreases with evolutionary distance. The 8 PSCs are not equidistant in evolutionary time, so we normalized them individually (=column-wise normalization). We used a rank-normalization (=rank / amount of rows), since their dN/dS ratios are not equally distributed. This brings the dN/dS of each column to a uniform distribution between 0 and 1. The rank-normalized dN/dS can then be interpreted as their percentile (e.g. value of 0.9 means the dN/dS represents the 90^th^ percentile) and a rank-normalized dN/dS of 0 is still a dN/dS of 0. The resulting table of dN/dS ranks is visualized in Fig. 1B.

#### Generating evolutionary maps

We visualized the evolutionary landscape throughout phylogenetic history of the brain by creating brain-region level evolutionary maps that color-code each region by its evolutionary time point. Time points were encoded by associating each brain region with its most specific PSC (pairwise species comparison), i.e. the LCA (last common ancestor) with the strongest structural association of genes under high selection pressure.

Therefore, we thought to predict the association of highly selective genes with functional neuroanatomical maps according to Ganglberger et al. *(17)* for each of the eight PSCs. To our better knowledge, there is no general threshold for highly selected genes based on (ranked) dN/dS ratio, so it is not possible to choose gene sets to predict these maps. Hence, we modeled selection pressure by weighting spatial gene expression from AHBA/AMBA by their dN/dS rank for each PSC. This creates 8 sets of 10445 genes for which genes with low selection pressure have low expression values, and genes with high selection pressure are similar to their original expression values (because dN/dS ranks are between 0 and 1). The resulting eight functional maps for mouse and AMH were all significantly different (FDR=0.1) from functional maps with randomly associated dN/dS ranks (i.e. we shuffled the dN/dS ranks before weighting). We computed the prediction of functional maps purely on expression site level (biopsy site), without applying higher order network measures. Since the 10445 genes do not represent single multigenic traits as used in Ganglberger et al. *(17)*, the gene expression synergy was not region specific, which did not lead to region specific changes in gene expression synergy weighted networks. This indicated that the genes with the highest selection pressure for each PCS modulate multiple brain functions/networks.

The AMH data, assembled from gene expression of 3702 biopsy-sites from microarray data of the Allen Human Brain Atlas (*22*), was visualized on a region level (most significant p-value of all biopsy sites within a brain region) in Fig. 3.

Finally, we wanted to combine the maps for each PSC to a single evolutionary map. To generate these, the highest specificity for each biopsy site was represented by the PSC with the most significant p-value (Fig. 2). Please note that the individual PSC maps showed already accumulated biopsy site values at (sub)region level, thus it was not possible to derive the most specific PSC directly from Fig. 3. Theses maps showed the most significant p-value of the biopsy-sites within a brain region (and therefore only the most significant biopsy-site), while the evolutionary map was computed first on biopsy-level and then visualized the most frequent PSC of all biopsy sites.

**Methods Table 2.**
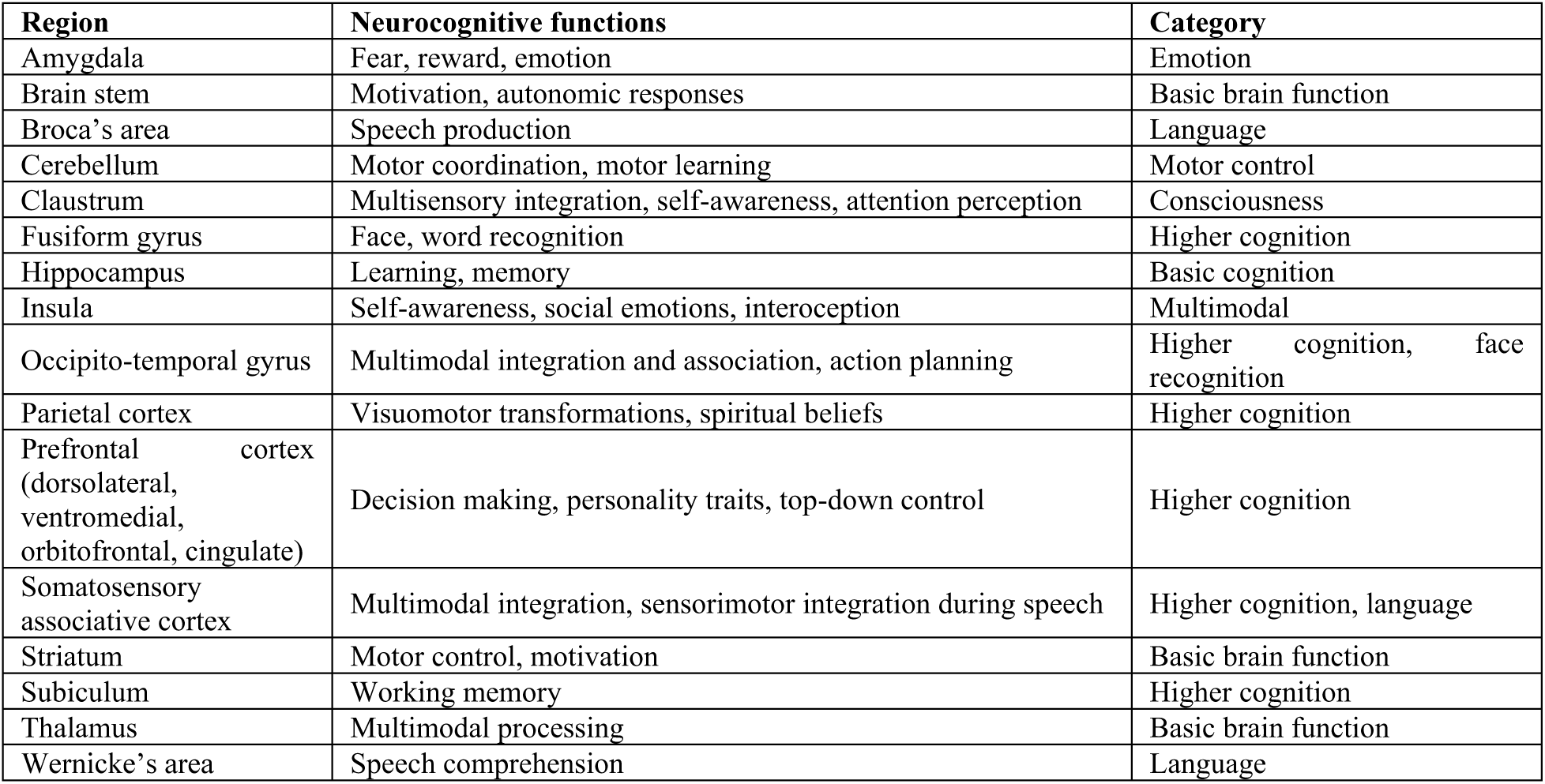
Functional neuroanatomical annotation.

#### Task-evoked functional brain activity

Task-specific brain activity maps were downloaded from Human Connectome Project (HCP) website (www.humanconnectome.org). We used data available for seven major domains, detailed description can be found in Barch et al. (*98*). Contrasts selected for comparison with dN/dS functional maps were collected in Methods Table 3. The contrasts labels and behavioral signatures descriptions correspond to Tavor et al. (*99*).

#### Functional network meta-comparison

Literature-based comparison of regions involved in several functional networks extracted from fMRI scans was correlated to functional maps (Methods, Computational neuroanatomy, Generating evolutionary maps). Networks used in the study are collected in Methods Table 4.

#### Subspace pattern mining for network evolution via biclustering

To identify genes linked to specific tasks or functional networks, we mined co-evolving genes with high spatial correlations to those networks. Therefore, we retrieved AMH spatial gene expression data from the AHBA for each of the 10445 genes in the dN/dS ranked table. Spatial gene expression data was available for ∼3700 biopsy sites in the brain. We mapped both, the task fMRI data of 11 networks (Methods, Computational neuroanatomy, Correlation against task-evoked functional brain activity) and the literature-based data on another 11 networks (Methods, Computational neuroanatomy, Functional network meta-comparison) to the biopsy sites and computed the Spearman rank correlation coefficient between every gene and network.

To make both types of functional data comparable, we normalized each block (104445×11 fMRI-to-gene-expression correlations and 104445×11 literature-to-gene-expression correlations) with z-score standardization. Functional network specificity for each gene was then computed by rank-normalization (rank / amount of networks) for each gene over all networks, so they were mapped to a range between 0 and 1. As a result, 0 was assigned to the network with the lowest correlation. The network with the highest correlation to a gene gets the value of 1. We concatenated this data with the dN/dS ranked table to receive a 10445 × 30 spatio-temporal network table (8 PSCs with dN/dS ranks, 11 fMRI-to-gene-expression and 11 literature-to-gene-expression correlation ranks). From this table, we removed all genes with an overall low correlation to all networks (genes for which all network correlations were smaller than 0.1), which reduced the table to 6239 rows.

We performed data mining by using GABi (*34*), a framework that facilitates a genetic algorithm for biclustering (simultaneous clustering of the rows and columns of a matrix), available as R package. Compared to other biclustering algorithms, the advantage of this framework allows the definition of a customized criteria. This enables the user to specify the properties a bicluster should have, such as coherence, consistency, size etc. We used these custom criteria, in terms of genetic algorithms also called “fitness function” to find biclusters of highly selected genes (genes that have high dN/dS ranks over multiple PSCs) with high specificity for similar functional networks (genes with high network correlation ranks for the same networks). Therefore, GABi creates a set of candidate solutions (a candidate solution is a set of genes) and applies the fitness function to see which PSCs and networks fit the custom criteria. It iteratively optimizes the candidate solutions to find the largest bicluster fitting these criteria by means of evolution inspired operators such as mutation, crossover and selection (*34*).

We defined the custom criteria for fitness function to find the largest bicluster:

-with **at least one PSC and one network**, since biclusters without one of them do not represent genes with high dN/dS ranks and high specificity for similar functional networks
- with **PSCs having a mean dN/dS rank >=0.9** for the genes in the bicluster. This selects only genes that have PSCs with an average dN/dS rank above equal 0.9, which puts them in the top 10 percentile of dN/dS ranks. Therefore, they can be considered as the top 10 percent genes regarding their selection pressure, therefore highly selective.
- with **networks having a mean network correlation specificity rank >=0.75** for the genes in the bicluster. Therefore, only genes with an average network specificity in the top 25% genes (the networks with the 1-8th highest correlation to each gene) were in the bicluster.
- where **the gene’s dN/dS and network correlation ranks are above 0**. The genetic algorithm optimizes the fitness function of the candidate solutions and therefore finds the largest bicluster. If a bicluster has a mean network correlation rank >0.75 for a specific network, the algorithm will add as many genes with a network correlation rank <0.75 until its size is maximized. We limited this by omitting genes that have zero network correlation.

If a bicluster has this property, then we define its fitness as

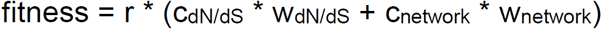

where r is the number of rows, c_dN/dS_ the amount of dN/dS rank columns and c_network_ the amount of gene-to-network correlation ranks columns. w_dN/dS_ and w_network_ are weighting factors to account for the different amount of dN/dS rank columns (8) and network correlation ranks columns (22). Otherwise it would be more likely to find biclusters with network correlation ranks columns than dN/dS rank columns. w_dN/dS_ = 1.875 and w_network_ =0.6818, to sum up to 30 again (similar to not weighted).

Stability tests were performed on random subsets of the data to empirically estimate parameters for the genetic algorithm utilized by GABi. We found that a population size of at least 10-20 times the ‘chromosome’ length (amount of rows of the data set (*34*)) with ∼2000 demes (separate subpopulations (*34*)) led to stable results over multiple runs and therefore reproducible results. This amount of demes was necessary to minimize the chances of locally-optima solutions for GABi(*34*), since it must converge to solutions that are relatively small compared to the search space (identified biclusters had about 1-2% of the total amount of genes). Each deme needed at least 32 individual solutions (*34*) (=chromosomes), therefore the 2000 demes constrained the population size to be at least ∼10 times the chromosome length (2000*32/6239 = 10.26).

We applied GABi with a population size of 124780 (20x the chromosome length of 6239) chromosomes divided into 2000 demes. We ran the biclustering with these parameters several times with similar results. The algorithm found for each bicluster up to 100 variations (i.e. biclusters with the same fitness/size, but differ at several genes), so we grouped biclusters with at least 75% overlapping genes to form one representative. A representative bicluster did not fulfil the criteria of the fitness function (otherwise it would have been found by GABi directly) but summarized all genes of all of its variations. Statistical evaluation was performed by permutation tests to verify that the representative biclusters had significantly higher mean dN/dS ranks, and gene-to-network correlation ranks than random sets of similar size. P-values were highly significant (<0.0001), which was expected since the fitness function was specifically designed to find highly consistent biclusters.

Biclusters, respectively bicluster modules, shown in Fig. 4, 5 and Fig. S1 were visualized with newly developed tool (*100*) (VRVis, Vienna). Nodes were selected from the bicluster’s networks (Methods Table 3 & 4, Fig. S1), the edges represented the strongest spatial gene expression correlation of the bicluster’s genes. The networks were created similarly to Methods, Computational neuroanatomy and Tracing the evolution of spatial gene expression correlation networks. Originally the edges showed a region-bias, i.e. regions that had higher correlation between them than others over all network, resulting from the amount of genes in a bicluster (the correlation of gene sets converges to the genome-wide spatial gene expression correlation with increasing size). We targeted this by generating an empirical distribution for each individual edge by 1000 random drawn gene sets from the genome (of same size as the bicluster). These distributions (i.e. their mean and standard distribution) were then used for z-score normalization of the bicluster edges.

#### Functional genetics

For functional profiling of genes clustered with brain networks we applied the knowledgebase from Ingenuity Pathway Analysis (IPA) (QIAGEN Inc., https://www.qiagenbioinformatics.com/products/ingenuitypathway-analysis). Each cluster from biclustering of full lineage or hominid branch was analyzed separately. We applied Nervous System filter to avoid non-specific functional associations. All results can be found in the Supplementary Materials.

### Methodological remarks

Taken together this workflow maps dN/dS data onto spatial brain gene expression and correlates it with task-evoked fMRI and known functional networks (see Main text and Methods, Methods Tables 3 and 4). Both, cumulative correlation and biclustering, build on dN/dS-driven spatial correlations of highly selected genes per PSC with fMRI networks. Cumulative correlation focuses on functional correlations for each PSC top-selected genes, while biclustering relates ranked dN/dS of all PSCs and fMRI networks simultaneously, building clusters of highly selected genes (bound to any PSC) which highly correlate with fMRI networks. Jointly, these approaches are in support of each other and reconstruct a congruent ancestral history of the brain’s functional evolution.

Of note, overall, results obtained with PSCs along the AMH lineage, shown here, are similar to predictions from PSCs rooted in AMH (AMH vs all species).

**Methods Table 3.** Functional maps from AMH task-evoked fMRI.

| Task network | Contrast | Neurocognitive functions | Category |
| --- | --- | --- | --- |
| Emotional faces | EM-FACES | Valance judgments (faces) | Emotion |
| Gambling<br>Gambling-reward | GA-AVG<br>GA-REWARD | Reward, punishment, decision making | Strategic thought |
| Language | LA-AVG | Sentences, stories, mental arithmetic (auditory) | Language |
| Motor hand control<br>Motor foot control<br>Motor tongue control | MO-RH+LH<br>MO-RF+LF<br>MO-T | Hand, foot, tongue movements | Motor control |
| Relational processing | RE-AVG | Higher-order cognition (attention, working memory) | Higher cognition |
| Theory of mind | SO-TOM | Interpret social vs random interactions | Social cognition |
| Working memory<br>Working memory-faces | WM-REST<br>WM-FACE | N-back working memory, body parts (face), tools, places | Working memory |

**Methods Table 4.**
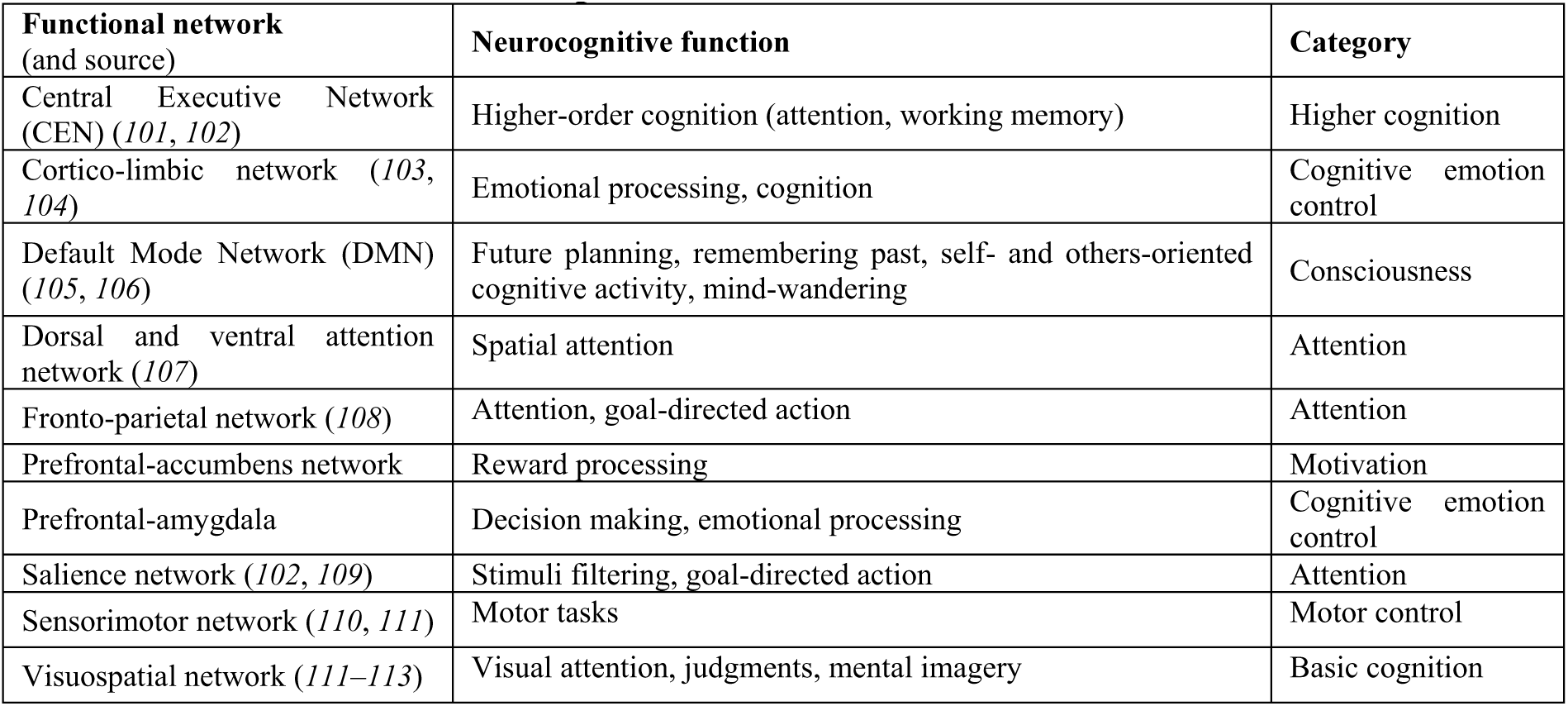
Functional maps from literature.

### Statistics

Statics are described in the corresponding paragraphs of the Methods section.

## References

1. C. P. van S. & K. I. Ana Navarrete, Energetics and the evolution of human brain size. Nature (2011), doi:10.1038/nature10629.

2. M. Hofman, Evolution of the human brain: when bigger is better. Front. Neuroanat. 8 (2014), p. 15.

3. S. Neubauer, J.-J. Hublin, P. Gunz, The evolution of modern human brain shape. Sci. Adv. 4, eaao5961 (2018).

4. A. Du et al., Pattern and process in hominin brain size evolution are scale-dependent. Proc. R. Soc. B Biol. Sci. 285, 20172738 (2018).

5. M. González-Forero, A. Gardner, Inference of ecological and social drivers of human brain-size evolution. Nature. 557, 554–557 (2018).

6. K. Leah, In Search of a Unifying Theory of Complex Brain Evolution. Ann. N. Y. Acad. Sci. 1156, 44–67 (2009).

7. G. Roth, Convergent evolution of complex brains and high intelligence. Philos. Trans. R. Soc. B Biol. Sci. 370 (2015) (available at http://rstb.royalsocietypublishing.org/content/370/1684/20150049.abstract).

8. S. Herculano-Houzel, The remarkable, yet not extraordinary, human brain as a scaled-up primate brain and its associated cost. Proc. Natl. Acad. Sci. 109, 10661–10668 (2012).

9. E. L. MacLean, Unraveling the evolution of uniquely human cognition. Proc. Natl. Acad. Sci. 113, 6348–6354 (2016).

10. Z. He et al., Comprehensive transcriptome analysis of neocortical layers in humans, chimpanzees and macaques. Nat. Neurosci. 20, 886–895 (2017).

11. A. L. Bauernfeind et al., Evolutionary divergence of gene and protein expression in the brains of humans and chimpanzees. Genome Biol. Evol. 7, 2276–2288 (2015).

12. C. C. Sherwood, F. Subiaul, T. W. Zawidzki, A natural history of the human mind: Tracing evolutionary changes in brain and cognition. J. Anat. 212, 426–454 (2008).

13. A. M. M. Sousa, K. A. Meyer, G. Santpere, F. O. Gulden, N. Sestan, Evolution of the Human Nervous System Function, Structure, and Development. Cell. 170, 226–247 (2017).

14. W. Enard et al., Molecular evolution of FOXP2, a gene involved in speech and language. Nature (2002), doi:10.1038/nature01025.

15. X. Nuttle et al., Emergence of a Homo sapiens-specific gene family and chromosome 16p11.2 CNV susceptibility. Nature. 536, 205–209 (2016).

16. J. Richiardi et al., Correlated gene expression supports synchronous activity in brain networks. Science (80-.). 348, 1241–1244 (2015).

17. F. Ganglberger et al., Predicting functional neuroanatomical maps from fusing brain networks with genetic information. Neuroimage (2018), doi:10.1016/j.neuroimage.2017.08.070.

18. D. R. Zerbino et al., Ensembl 2018. Nucleic Acids Res. (2018), doi:10.1093/nar/gkx1098.

19. R. E. Green et al., A Draft Sequence of the Neandertal Genome. Science (80-.). 328, 710– 722 (2010).

20. K. Prüfer et al., The complete genome sequence of a Neanderthal from the Altai Mountains. Nature. 505, 43–49 (2014).

21. M. Meyer et al., A high-coverage genome sequence from an archaic Denisovan individual. Science (80-.). (2012), doi:10.1126/science.1224344.

22. M. J. Hawrylycz et al., An anatomically comprehensive atlas of the adult human brain transcriptome. Nature. 489, 391–399 (2012).

23. S. L. Ding et al., Comprehensive cellular-resolution atlas of the adult human brain. J. Comp. Neurol. (2016), doi:10.1002/cne.24080.

24. D. C. Van Essen et al., The WU-Minn Human Connectome Project: An overview. Neuroimage (2013), doi:10.1016/j.neuroimage.2013.05.041.

25. G. Z. Wang et al., Correspondence between Resting-State Activity and Brain Gene Expression. Neuron. 88, 659–666 (2015).

26. M. Hawrylycz et al., Canonical genetic signatures of the adult human brain. Nat. Neurosci. 18, 1832–1844 (2015).

27. M. E. Skinner, A. V. Uzilov, L. D. Stein, C. J. Mungall, I. H. Holmes, JBrowse: A nextgeneration genome browser. Genome Res. (2009), doi:10.1101/gr.094607.109.

28. A. M. Boddy et al., Evidence of a Conserved Molecular Response to Selection for Increased Brain Size in Primates. Genome Biol. Evol. 9, 700–713 (2017).

29. J. Wang et al., Peregrine and saker falcon genome sequences provide insights into evolution of a predatory lifestyle. Nat. Genet. 45, 563–566 (2013).

30. F. Crick, C. Koch, A framework for consciousness. Nat. Neurosci. (2003), doi:10.1038/nn0203-119.

31. C. Urgesi, S. M. Aglioti, M. Skrap, F. Fabbro, The Spiritual Brain: Selective Cortical Lesions Modulate Human Self-Transcendence. Neuron (2010), doi:10.1016/j.neuron.2010.01.026.

32. C. J. Price, The anatomy of language: Contributions from functional neuroimaging. J. Anat. (2000), doi:10.1017/S0021878299006901.

33. M. P. Mattson, Superior pattern processing is the essence of the evolved human brain. Front. Neurosci. (2014), doi:10.3389/fnins.2014.00265.

34. E. W. J. Curry, A framework for generalized subspace pattern mining in high-dimensional datasets. BMC Bioinformatics (2014), doi:10.1186/s12859-014-0355-5.

35. F. F. Hamdan et al., De novo mutations in FOXP1 in cases with intellectual disability, autism, and language impairment. Am. J. Hum. Genet. (2010), doi:10.1016/j.ajhg.2010.09.017.

36. A. K. Le Fevre et al., FOXP1 mutations cause intellectual disability and a recognizable phenotype. Am. J. Med. Genet. Part A (2013), doi:10.1002/ajmg.a.36174.

37. K. J. Bunn et al., Mutations in DVL1 cause an osteosclerotic form of robinow syndrome. Am. J. Hum. Genet. (2015), doi:10.1016/j.ajhg.2015.02.010.

38. A. P. Jackson et al., Identification of Microcephalin, a Protein Implicated in Determining the Size of the Human Brain. Am. J. Hum. Genet. (2002), doi:10.1086/341283.

39. S. Berkel et al., Mutations in the SHANK2 synaptic scaffolding gene in autism spectrum disorder and mental retardation. Nat. Genet. (2010), doi:10.1038/ng.589.

40. K. Roybal et al., Mania-like behavior induced by disruption of CLOCK. Proc. Natl. Acad. Sci. (2007), doi:10.1073/pnas.0609625104.

41. G. Konopka et al., Human-Specific Transcriptional Networks in the Brain. Neuron. 75, 601–617 (2012).

42. A. Krämer, J. Green, J. Pollard, S. Tugendreich, Causal analysis approaches in ingenuity pathway analysis. Bioinformatics (2014), doi:10.1093/bioinformatics/btt703.

43. L. a. O’Connell, H. a. Hofmann, Evolution of a Vertebrate Social Decision-Making Network. Science (80-.). 336, 1154–1157 (2012).

44. E. Di Lullo, A. R. Kriegstein, The use of brain organoids to investigate neural development and disease. Nat. Rev. Neurosci. (2017), doi:10.1038/nrn.2017.107.

45. J. Jaubert et al., Early Neanderthal constructions deep in Bruniquel Cave in southwestern France. Nature. 534, 111–114 (2016).

46. D. L. Hoffmann et al., U-Th dating of carbonate crusts reveals Neandertal origin of Iberian cave art. Science (80-.). 359, 912–915 (2018).

47. D. Dediu, S. C. Levinson, Neanderthal language revisited: not only us. Curr. Opin. Behav. Sci. (2018), doi:10.1016/j.cobeha.2018.01.001.

48. A. J. Drummond, A. Rambaut, BEAST: Bayesian evolutionary analysis by sampling trees. BMC Evol. Biol. (2007), doi:10.1186/1471-2148-7-214.

49. D. Posada, jModelTest: Phylogenetic model averaging. Mol. Biol. Evol. (2008), doi:10.1093/molbev/msn083.

50. S. I. Perez, M. F. Tejedor, N. M. Novo, L. Aristide, Divergence Times and the Evolutionary Radiation of New World Monkeys (Platyrrhini, Primates): An Analysis of Fossil and Molecular Data. PLoS One (2013), doi:10.1371/journal.pone.0068029.

51. A. Rambaut, A. J. Drummond, Tracer v1.4. Encycl. Atmos. Sci. (2007), doi:10.1111/1469-8986.ep10972559.

52. Rambaut, FigTree v. 1.4.0. http://tree.bio.ed.ac.uk/software/figtree/ (2012).

53. M. Meyer et al., A mitochondrial genome sequence of a hominin from Sima de los Huesos. Nature. 505, 403–406 (2014).

54. M. T. Silcox, C. K. Dalmyn, J. I. Bloch, Virtual endocast of Ignacius graybullianus (Paromomyidae, Primates) and brain evolution in early primates. Proc. Natl. Acad. Sci. (2009), doi:10.1073/pnas.0812140106.

55. K. Isler et al., Endocranial volumes of primate species: scaling analyses using a comprehensive and reliable data set. J. Hum. Evol. 55, 967–978 (2008).

56. K. Semendeferi, H. Damasio, The brain and its main anatomical subdivisions in living hominoids using magnetic resonance imaging. J. Hum. Evol. (2000), doi:10.1006/jhev.1999.0381.

57. E. Pearce, C. Stringer, R. I. M. Dunbar, New insights into differences in brain organization between Neanderthals and anatomically modern humans. Proc. R. Soc. B Biol. Sci. (2013), doi:10.1098/rspb.2013.0168.

58. G. Roth, U. Dicke, Evolution of the brain and intelligence. Trends Cogn. Sci. (2005), doi:10.1016/j.tics.2005.03.005.

59. L. S. Weyrich et al., Neanderthal behaviour, diet, and disease inferred from ancient DNA in dental calculus. Nature. 544, 357–361 (2017).

60. S. Castellano et al., Patterns of coding variation in the complete exomes of three Neandertals. Proc. Natl. Acad. Sci. (2014), doi:10.1073/pnas.1405138111.

61. B. Wilson, C. I. Petkov, Communication and the primate brain: insights from neuroimaging studies in humans, chimpanzees and macaques. Hum. Biol. (2011), doi:10.3378/027.083.0203.Communication.

62. F. Aboitiz, A brain for speech. Evolutionary continuity in primate and human auditory-vocal processing. Front. Neurosci. (2018), doi:10.3389/fnins.2018.00174.

63. J. K. Rilling et al., The evolution of the arcuate fasciculus revealed with comparative DTI. Nat. Neurosci. (2008), doi:10.1038/nn2072.

64. S. Johansson, Institutional repository of Jönköping University The Talking Neanderthals?: What Do Fossils, Genetics, and Archeology Say?, 35–74 (2013).

65. P. Lieberman, The evolution of language and thought. J. Anthropol. Sci. 94, 127–146 (2016).

66. I. Tattersall, The material record and the antiquity of language. Neurosci. Biobehav. Rev. 81, 247–254 (2017).

67. J. Sliwa, W. A. Freiwald, Neuroscience: A dedicated network for social interaction processing in the primate brain. Science (80-.). (2017), doi:10.1126/science.aam6383.

68. S. F. Brosnan, F. B. M. De Waal, Monkeys reject unequal pay. Nature (2003), doi:10.1038/nature01963.

69. S. F. Brosnan, L. Salwiczek, R. Bshary, The interplay of cognition and cooperation. Philos. Trans. R. Soc. B Biol. Sci. (2010), doi:10.1098/rstb.2010.0154.

70. T. Kochiyama et al., Reconstructing the Neanderthal brain using computational anatomy, 1–9 (2018).

71. A. Whiten, Primate culture and social learning. Cogn. Sci. (2000), doi:10.1207/s15516709cog2403_6.

72. E. Herrmann, J. Call, M. V. Hernández-Lloreda, B. Hare, M. Tomasello, Humans have evolved specialized skills of social cognition: The cultural intelligence hypothesis. Science (80-.). (2007), doi:10.1126/science.1146282.

73. S. Herculano-Houzel, C. E. Collins, P. Wong, J. H. Kaas, Cellular scaling rules for primate brains. Proc. Natl. Acad. Sci. (2007), doi:10.1073/pnas.0611396104.

74. M. J. Beran et al., Primate cognition: attention, episodic memory, prospective memory, selfcontrol, and metacognition as examples of cognitive control in nonhuman primates. Wiley Interdiscip. Rev. Cogn. Sci. (2016), doi:10.1002/wcs.1397.

75. T. D. Hanks, C. Summerfield, Perceptual Decision Making in Rodents, Monkeys, and Humans. Neuron (2017), doi:10.1016/j.neuron.2016.12.003.

76. S. Tremblay, K. M. Sharika, M. L. Platt, Social Decision-Making and the Brain: A Comparative Perspective. Trends Cogn. Sci. (2017), doi:10.1016/j.tics.2017.01.007.

77. C. M. Conway, M. H. Christiansen, Sequential learning in non-human primates. Trends Cogn. Sci. (2001), doi:10.1016/S1364-6613(00)01800-3.

78. T. Wynn, F. L. Coolidge, The expert Neandertal mind. J. Hum. Evol. 46, 467–487 (2004).

79. D. L. Robinson, M. E. Goldberg, G. B. Stanton, Parietal association cortex in the primate: sensory mechanisms and behavioral modulations. J. Neurophysiol. (1978), doi:10.1152/jn.1978.41.4.910.

80. D. J. Felleman, D. C. Van Essen, Distributed hierarchical processing in the primate cerebral cortex. Cereb. Cortex (1991), doi:10.1093/cercor/1.1.1.

81. J. C. I. Belmonte et al., Brains, Genes, and Primates. Neuron. 86, 617–631 (2015).

82. M.-E. LaramÃ©e, D. Boire, Visual cortical areas of the mouse: comparison of parcellation and network structure with primates. Front. Neural Circuits (2015), doi:10.3389/fncir.2014.00149.

83. E. Bruner, M. Lozano, Extended mind and Visuo-Spatial integration: Three hands for the Neandertal lineage. J. Anthropol. Sci. 92, 173–280 (2014).

84. E. Pearce, C. Stringer, R. I. M. Dunbar, New insights into differences in brain organization between Neanderthals and anatomically modern humans. Proc. R. Soc. B Biol. Sci. 280, 20130168–20130168 (2013).

85. T. Imura, M. Tomonaga, Differences between chimpanzees and humans in visual temporal integration. Sci. Rep. (2013), doi:10.1038/srep03256.

86. E. E. Hecht et al., Differences in Neural Activation for Object-Directed Grasping in Chimpanzees and Humans. J. Neurosci. (2013), doi:10.1523/JNEUROSCI.2172-13.2013.

87. J. H. Kaas, in Evolution of Nervous Systems (2010).

88. M. Bastir et al., Evolution of the base of the brain in highly encephalized human species. Nat. Commun. 2 (2011), doi:10.1038/ncomms1593.

89. J. Herrero et al., Ensembl comparative genomics resources. Database (2016), doi:10.1093/database/bav096.

90. S. Durinck et al., BioMart and Bioconductor: A powerful link between biological databases and microarray data analysis. Bioinformatics (2005), doi:10.1093/bioinformatics/bti525.

91. K. Prüfer et al., A high-coverage Neandertal genome from Vindija Cave in Croatia (Prufer,2017).pdf. 1887, 1–13 (2017).

92. H. Li et al., The Sequence Alignment/Map format and SAMtools. Bioinformatics (2009), doi:10.1093/bioinformatics/btp352.

93. H. Li, A statistical framework for SNP calling, mutation discovery, association mapping and population genetical parameter estimation from sequencing data. Bioinformatics (2011), doi:10.1093/bioinformatics/btr509.

94. A. Löytynoja, Phylogeny-aware alignment with PRANK. Methods Mol. Biol. (2014), doi:10.1007/978-1-62703-646-7_10.

95. Z. Yang, PAML 4: Phylogenetic analysis by maximum likelihood. Mol. Biol. Evol. (2007), doi:10.1093/molbev/msm088.

96. J. L. Villanueva-Cañas, S. Laurie, M. M. Albà, Improving genome-wide scans of positive selection by using protein isoforms of similar length. Genome Biol. Evol. (2013), doi:10.1093/gbe/evt017.

97. J. B. W. Wolf, A. Kunstner, K. Nam, M. Jakobsson, H. Ellegren, Nonlinear Dynamics of Nonsynonymous (dN) and Synonymous (dS) Substitution Rates Affects Inference of Selection. Genome Biol. Evol. (2010), doi:10.1093/gbe/evp030.

98. D. M. Barch et al., Function in the human connectome: Task-fMRI and individual differences in behavior. Neuroimage (2013), doi:10.1016/j.neuroimage.2013.05.033.

99. I. Tavor et al., Task-free MRI predicts individual differences in brain activity during task performance. Science (80-.). 352, 216–220 (2016).

100. F. Ganglberger et al., in Eurographics Proceedings (The Eurographics Association, 2018; https://doi.org/10.2312/vcbm.20181231).

101. V. Menon, in Brain Mapping: An Encyclopedic Reference (2015).

102. W. W. Seeley et al., Dissociable Intrinsic Connectivity Networks for Salience Processing and Executive Control. J. Neurosci. (2007), doi:10.1523/JNEUROSCI.5587-06.2007.

103. P. Fang et al., Increased Cortical-Limbic Anatomical Network Connectivity in Major Depression Revealed by Diffusion Tensor Imaging. PLoS One (2012), doi:10.1371/journal.pone.0045972.

104. B. Vai et al., Abnormal cortico-limbic connectivity during emotional processing correlates with symptom severity in schizophrenia. Eur. Psychiatry (2015), doi:10.1016/j.eurpsy.2015.01.002.

105. P. Lin et al., Dynamic Default Mode Network across Different Brain States. Sci. Rep. (2017), doi:10.1038/srep46088.

106. J. R. Andrews-Hanna, J. Smallwood, R. N. Spreng, The default network and self-generated thought: Component processes, dynamic control, and clinical relevance. Ann. N. Y. Acad. Sci. 1316, 29–52 (2014).

107. S. Vossel, J. J. Geng, G. R. Fink, Dorsal and Ventral Attention Systems. Neurosci. (2014), doi:10.1177/1073858413494269.

108. T. P. Zanto, A. Gazzaley, Fronto-parietal network: Flexible hub of cognitive control. Trends Cogn. Sci. (2013), doi:10.1016/j.tics.2013.10.001.

109. L. Q. Uddin, L. Q. Uddin, Anatomy of the Salience Network. Salience Netw. Hum. Brain, 5–10 (2017).

110. F. Ferri, F. Frassinetti, M. Ardizzi, M. Costantini, V. Gallese, A sensorimotor network for the bodily self. J. Cogn. Neurosci. (2012), doi:10.1162/jocn_a_00230.

111. S. M. Smith et al., Correspondence of the brain’s functional architecture during activation and rest. Proc. Natl. Acad. Sci. 106, 13040–13045 (2009).

112. T. A. De Graaf, A. Roebroeck, R. Goebel, A. T. Sack, Brain network dynamics underlying visuospatial judgment: An FMRI connectivity study. J. Cogn. Neurosci. (2010), doi:10.1162/jocn.2009.21345.

113. K. Whittingstall, M. Bernier, J. C. Houde, D. Fortin, M. Descoteaux, Structural network underlying visuospatial imagery in humans. Cortex (2014), doi:10.1016/j.cortex.2013.02.004.

