## Supplementary Material for "Molecular archaeology of the human brain"

### Supplementary Materials content:

**Figure S1** ........ **3**
**Supplementary Table 1** ........ **4**
Module 1 Cluster 6 ........ 4
**Supplementary Table 2** ........ **5**
Module 2 Cluster 1 ........ 5
Module 2 Cluster 3 ........ 12
Module 2 Cluster 5 ........ 15
**Supplementary Table 3** ........ **16**
Module 3 Cluster 7 ........ 16
**Supplementary Table 4** ........ **24**
Module 4 Cluster 10 ........ 24
Module 4 Cluster 11 ........ 26
**Supplementary Table 5** ........ **28**
Module 5 Cluster 12 ........ 28
**Supplementary Table 6** ........ **30**
Module 6 Cluster 9 ........ 30
Module 6 Cluster 13 ........ 34
Module 6 Cluster 14 ........ 36
Module 6 Cluster 15 ........ 37
**Supplementary Table 7** ........ **38**
Module 7 Cluster 2 ........ 38
Module 7 Cluster 4 ........ 45
Module 7 Cluster 8 ........ 65
**Supplementary Table 8** ........ **69**
Module 8 Cluster 4 ........ 69
Module 8 Cluster 6 ........ 73
Module 8 Cluster 7 ........ 75
Module 8 Cluster 8 ........ 79
Module 8 Cluster 9 ........ 80
Module 8 Cluster 13 ........ 86
Module 8 Cluster 14 ........ 88
Module 8 Cluster 15 ........ 100
**Supplementary Table 9** ........ **103**
Module 9 Cluster 1 ........ 103
Module 9 Cluster 2 ........ 119
Module 9 Cluster 3 ........ 131
Module 9 Cluster 5 ........ 154
Module 9 Cluster 11 ........ 175

Module 9 Cluster 12 ........ 193
**Supplementary Table 10** ........ **206**
Module 10 Cluster 10 ........ 206

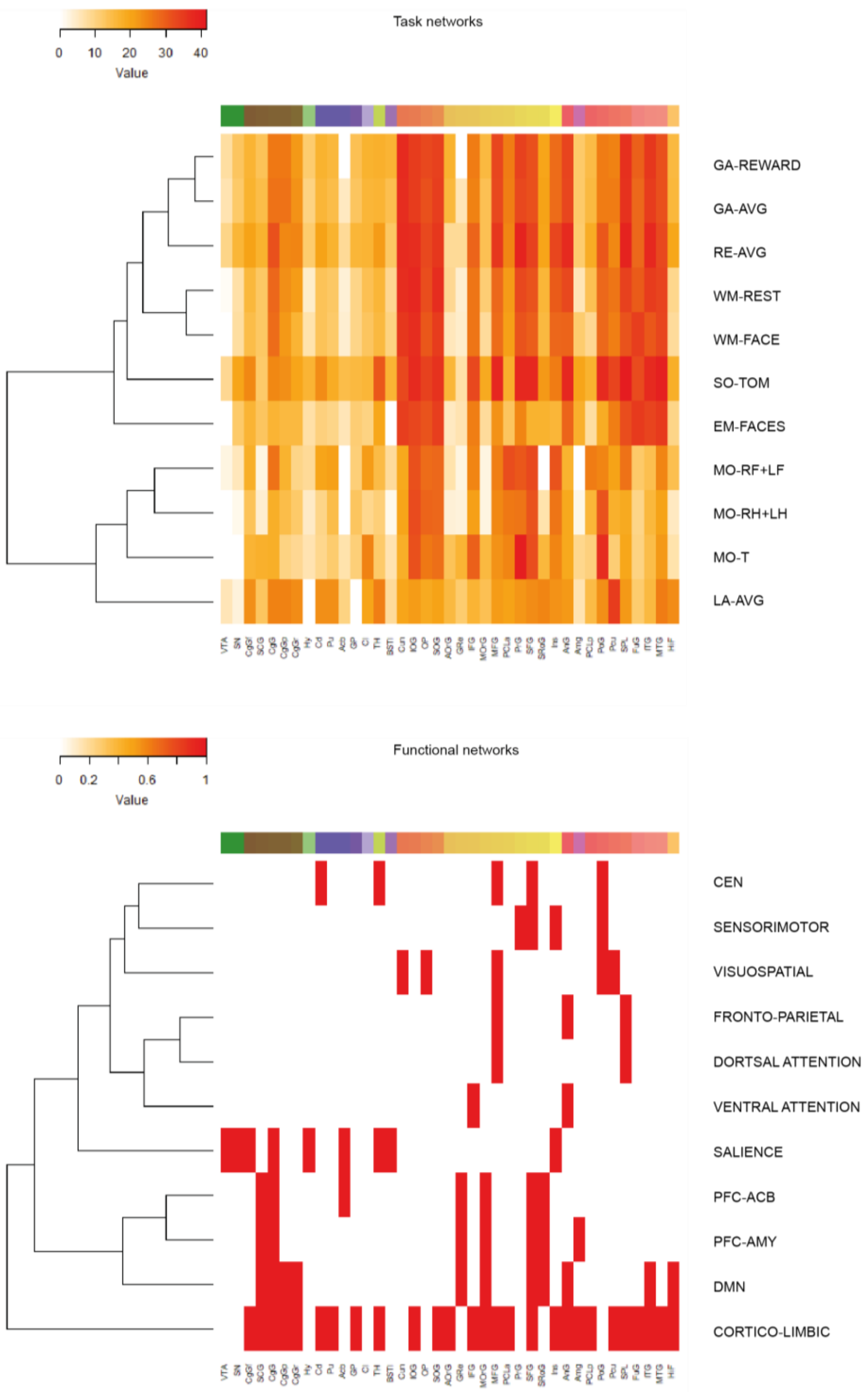


**Fig. S1. AMH task fMRI (TNs) and functional neuroanatomical (FNs) reference networks.** *Top*, Correlation matrix of AMH task-evoked fMRI with anatomical reference framework (ABA). Value corresponds to z-thresholded task contrast correlation to activation of selected regions. *Bottom*, literature-selected functional networks in a AMH anatomical reference framework (ABA). Value is a presence/absence call for selected regions (0 or 1). For network details and abbreviations see Methods Tables 3 & 4.

**Supplementary Table 1: Module 1**
**Cluster 6**

| Categories | Diseases or Functions Annotation | p-Value | Molecules | # Molecules |
|---|---|---|---|---|
| **Neurological Disease,Organ Morphology,Organismal Injury and Abnormalities** | Atrophy of hippocampal CA3 region | 0.00465 | TICAM1 | 1 |
| **Nervous System Development and Function** | Mechanical nociception | 0.00465 | CX3CL1 | 1 |
| **Endocrine System Development and Function,Organ Morphology,Organismal Development** | Enlargement of pituitary gland | 0.00465 | PTTG1 | 1 |
| **Cell Cycle** | Aneuploidy of cells | 0.00465 | PTTG1 | 1 |
| **Inflammatory Response,Organismal Injury and Abnormalities,Renal and Urological Disease** | Inflammation of urinary tract | 0.00465 | PTTG1 | 1 |
| **Cell Cycle,Cell Morphology,Cellular Assembly and Organization,Cellular Function and Maintenance** | Length of telomeres | 0.00465 | PTTG1 | 1 |
| **Cell Death and Survival,Neurological Disease,Organismal Injury and Abnormalities** | Apoptosis of lactotropes | 0.00928 | FAS | 1 |
| **Nervous System Development and Function,Organ Morphology,Organismal Development** | Mass of cerebellum | 0.00928 | IGFBP6 | 1 |
| **Organismal Functions** | Recovery of mice | 0.00928 | FAS | 1 |
| **Infectious Diseases,Neurological Disease** | Infection of ependyma | 0.00928 | TICAM1 | 1 |
| **Cellular Development,Nervous System Development and Function,Tissue Development** | Postsynaptic differentiation of cortical neurons | 0.00928 | IL1RAP | 1 |
| **Neurological Disease,Skeletal and Muscular Disorders** | Hypernociception of hindpaw | 0.00928 | CX3CL1 | 1 |
| **Neurological Disease,Organismal Injury and Abnormalities,Psychological Disorders** | Preclinical stage amyotrophic lateral sclerosis | 0.00928 | FAS | 1 |
| **Cellular Movement,Nervous System Development and Function** | Migration of hippocampal neurons | 0.00928 | CX3CL1 | 1 |
| **Small Molecule Biochemistry** | Quantity of kynurenic acid | 0.00928 | KMO | 1 |
| **Carbohydrate Metabolism,Drug Metabolism,Molecular Transport,Small Molecule Biochemistry** | Accumulation of hyaluronic acid | 0.00928 | HEXB | 1 |
| **Cell Death and Survival,Neurological Disease,Organismal Injury and Abnormalities** | Apoptosis of somatotrophs | 0.00928 | FAS | 1 |
| **Cardiovascular Disease,Organismal Injury and Abnormalities,Tissue Morphology** | Size of infarct | 0.0122 | BID, CX3CL1, TICAM1 | 3 |
| **Lipid Metabolism,Molecular Transport,Small Molecule Biochemistry** | Quantity of steroid | 0.0134 | FAS, PTTG1 | 2 |
| **Cell-To-Cell Signaling and Interaction,Cellular Function and Maintenance,Inflammatory Response,Nervous System Development and Function** | Phagocytosis of neurons | 0.0139 | CX3CL1 | 1 |
| **Infectious Diseases** | Infection by human herpesvirus 1 | 0.0139 | TICAM1 | 1 |
| **Cell-To-Cell Signaling and Interaction,Nervous System Development and Function** | Adhesion of hippocampal neurons | 0.0139 | CX3CL1 | 1 |
| **Cellular Movement,Hematological System Development and Function,Immune Cell Trafficking,Inflammatory Response,Nervous System Development and Function** | Migration of microglia | 0.0139 | CX3CL1 | 1 |
| **Cell-To-Cell Signaling and Interaction,Nervous System Development and Function** | Adhesion of cerebellar granule cell | 0.0139 | CX3CL1 | 1 |
| **Neurological Disease,Organismal Injury and Abnormalities,Psychological Disorders** | Multiple system atrophy | 0.0139 | FAS | 1 |
| **Cell Death and Survival,Cellular Compromise,Neurological Disease,Organismal Injury and Abnormalities,Tissue Morphology** | Neurodegeneration of Purkinje cells | 0.0201 | HEXB, MAP1A | 2 |
| **Cardiovascular Disease,Organismal Injury and Abnormalities,Tissue Morphology** | Volume of infarct | 0.0234 | BID, TICAM1 | 2 |

| | | | | |
|---|---|---|---|---|
| **Endocrine System Disorders,Gastrointestinal Disease,Metabolic Disease,Organismal Injury and Abnormalities** | Diabetes mellitus | 0.0276 | FAS | 1 |
| **Cellular Movement,Hematological System Development and Function,Immune Cell Trafficking** | Cell movement of leukocytes | 0.0307 | CX3CL1, FAS | 2 |
| **Cellular Function and Maintenance** | Endocytosis | 0.0327 | CX3CL1, KIAA1107 | 2 |
| **Cell Morphology,Nervous System Development and Function,Neurological Disease,Organ Morphology,Organismal Development,Organismal Injury and Abnormalities,Tissue Morphology** | Abnormal morphology of Purkinje cells | 0.0357 | HEXB, MAP1A | 2 |
| **Cardiovascular System Development and Function,Cell-To-Cell Signaling and Interaction** | Adhesion of endothelial cells | 0.0366 | FAS | 1 |
| **Cancer,Endocrine System Disorders,Neurological Disease,Organismal Injury and Abnormalities,Reproductive System Disease** | Hyperplasia of pituitary cells | 0.0366 | PTTG1 | 1 |
| **Cell Death and Survival,Embryonic Development** | Cell death of neuroblasts | 0.0366 | FAS | 1 |
| **Drug Metabolism,Endocrine System Development and Function,Lipid Metabolism,Molecular Transport,Small Molecule Biochemistry** | Concentration of testosterone | 0.0366 | PTTG1 | 1 |
| **Cellular Movement,Hematological System Development and Function,Immune Cell Trafficking** | Cell rolling of leukocytes | 0.0366 | FAS | 1 |
| **Dermatological Diseases and Conditions,Neurological Disease,Organismal Injury and Abnormalities** | Allodynia | 0.0411 | CX3CL1 | 1 |
| **Cell Death and Survival,Neurological Disease,Organismal Injury and Abnormalities** | Apoptosis of neural precursor cells | 0.0411 | FAS | 1 |
| **Neurological Disease,Organismal Injury and Abnormalities,Psychological Disorders** | Lesioning of brain | 0.0411 | CX3CL1 | 1 |
| **Cell Death and Survival** | Apoptosis of dopaminergic neurons | 0.0411 | FAS | 1 |
| **Cell Morphology,Nervous System Development and Function,Organ Morphology,Organismal Development** | Morphology of brain cells | 0.0465 | BID, HEXB, MAP1A | 3 |

**Supplementary Table 2: Module 2**
**Cluster 1**

| Categories | Diseases or Functions Annotation | p-Value | Molecules | # Molecules |
|---|---|---|---|---|
| **Cell-To-Cell Signaling and Interaction,Cellular Movement** | Recruitment of myeloid cells | 0.0000267 | CXCL2, IL33, TLR2, TLR4 | 4 |
| **Cell-To-Cell Signaling and Interaction,Cellular Movement,Hematological System Development and Function,Immune Cell Trafficking** | Recruitment of leukocytes | 0.0000427 | CXCL2, IL33, TLR2, TLR4 | 4 |
| **Endocrine System Development and Function,Organ Development** | Growth of pituitary gland | 0.0000649 | CGA, GNAS, IL1A, PRL | 4 |
| **Cellular Movement,Immune Cell Trafficking** | Leukocyte migration | 0.000092 | CD44, CXCL2, IL33, SELPLG, TLR2, TLR4 | 6 |
| **Cell-To-Cell Signaling and Interaction,Cellular Movement,Hematological System Development and Function,Immune Cell Trafficking** | Recruitment of antigen presenting cells | 0.0000958 | IL33, TLR2, TLR4 | 3 |
| **Cell-To-Cell Signaling and Interaction,Cellular Movement,Hematological System Development and Function,Immune Cell Trafficking,Inflammatory Response** | Recruitment of phagocytes | 0.000185 | CXCL2, TLR2, TLR4 | 3 |
| **Cellular Development,Cellular Growth and Proliferation,Endocrine System Development and Function,Nervous System Development and Function,Organ Development** | Proliferation of pituitary cells | 0.000604 | GNAS, IL1A, PRL | 3 |
| **Neurological Disease,Organismal Injury and Abnormalities** | Wallerian degeneration of sciatic nerve | 0.000671 | TLR2, TLR4 | 2 |
| **Cell-To-Cell Signaling and Interaction,Cellular Movement,Hematological System Development** | Recruitment of macrophages | 0.00111 | TLR2, TLR4 | 2 |

| | | | | |
|---|---|---|---|---|
| **and Function,Immune Cell Trafficking,Inflammatory Response** | | | | |
| **Cell-To-Cell Signaling and Interaction,Hematological System Development and Function** | Interaction of leukocytes | 0.00138 | CD44, SELPLG, TLR4 | 3 |
| **Cellular Compromise,Cellular Function and Maintenance** | Endoplasmic reticulum stress response of neurons | 0.00165 | SOD1, XBP1 | 2 |
| **Cardiovascular System Development and Function,Cellular Development,Cellular Function and Maintenance,Cellular Growth and Proliferation,Organismal Development,Tissue Development** | Cell proliferation of vascular endothelial cells | 0.00165 | CD44, PRL | 2 |
| **Nervous System Development and Function** | Remyelination of central nervous system | 0.0023 | PRL, TLR2 | 2 |
| **Cell-To-Cell Signaling and Interaction,Hematological System Development and Function** | Interaction of lymphocytes | 0.0023 | CD44, SELPLG | 2 |
| **Cell Death and Survival** | Apoptosis of microglia | 0.0023 | TLR2, TLR4 | 2 |
| **Cell-To-Cell Signaling and Interaction,Nervous System Development and Function** | Activation of astrocytes | 0.0026 | SOD1, TLR2, TLR4 | 3 |
| **Cell Morphology,Endocrine System Development and Function,Nervous System Development and Function,Organ Morphology,Organismal Development** | Morphology of anterior pituitary cells | 0.00305 | CGA, PRL | 2 |
| **Cell-To-Cell Signaling and Interaction,Cellular Growth and Proliferation,Nervous System Development and Function** | Excitation of sensory neurons | 0.00305 | IL1A, REST | 2 |
| **Cell Morphology,Hematological System Development and Function,Nervous System Development and Function** | Morphology of microglia | 0.00305 | SOD1, TLR4 | 2 |
| **Cellular Movement,Hematological System Development and Function,Immune Cell Trafficking** | Cell rolling of leukocytes | 0.00305 | SELPLG, TLR4 | 2 |
| **Free Radical Scavenging** | Production of reactive oxygen species | 0.00356 | SOD1, TLR2, TLR4 | 3 |
| **Cell Morphology,Cellular Function and Maintenance** | Autophagy of neurons | 0.00389 | SOD1, XBP1 | 2 |
| **Cellular Development,Cellular Growth and Proliferation,Endocrine System Development and Function,Nervous System Development and Function,Organ Development** | Proliferation of anterior pituitary cells | 0.00483 | GNAS, PRL | 2 |
| **Inflammatory Response,Neurological Disease** | Inflammation of central nervous system | 0.00493 | CD4, CD44, CD86, IL18, PTPRC, SOD1 | 6 |
| **Cell Morphology,Nervous System Development and Function,Tissue Morphology** | Size of neurites | 0.00586 | DISC1, SOD1 | 2 |
| **Cellular Development,Cellular Growth and Proliferation** | Proliferation of myeloid cells | 0.00698 | IFNGR1, SOD1 | 2 |
| **Cell Morphology,Nervous System Development and Function,Neurological Disease,Tissue Morphology** | Loss of neurites | 0.00712 | SERPINA3, SOD1, TLR4 | 3 |
| **Inflammatory Disease,Inflammatory Response,Neurological Disease,Organismal Injury and Abnormalities** | Encephalitis | 0.00751 | CD4, CD86, IL18, PTPRC, SOD1 | 5 |
| **Endocrine System Development and Function,Nervous System Development and Function,Organ Morphology,Tissue Morphology** | Quantity of lactotropes | 0.00819 | CGA, PRL | 2 |
| **Cardiovascular Disease,Neurological Disease** | Disruption of blood-brain barrier | 0.00949 | SOD1, TLR2 | 2 |
| **Cell Morphology,Nervous System Development and Function,Tissue Morphology** | Area of neurons | 0.00949 | DISC1, IGFBP1 | 2 |
| **Cancer,Neurological Disease,Organismal Injury and Abnormalities** | Medulloblastoma-like tumor | 0.0107 | GNAS | 1 |
| **Cellular Function and Maintenance** | Uptake by macrophages | 0.0107 | TLR4 | 1 |
| **Neurological Disease,Organismal Injury and Abnormalities** | Neuropathy of striatum | 0.0107 | IL18 | 1 |
| **Cell Morphology,Endocrine System Development and Function,Endocrine System Disorders,Nervous System Development and** | Abnormal morphology of gonadotropes | 0.0107 | PRL | 1 |

| | | | | |
|---|---|---|---|---|
| **Function,Neurological Disease,Organ Morphology,Organismal Development,Organismal Injury and Abnormalities,Reproductive System Disease** | | | | |
| **Cellular Compromise** | Abnormality of mitochondria | 0.0107 | SOD1 | 1 |
| **Cellular Assembly and Organization** | Quantity of rough endoplasmatic reticulum | 0.0107 | SOD1 | 1 |
| **Carbohydrate Metabolism,Molecular Transport** | Accumulation of glycogen | 0.0107 | SOD1 | 1 |
| **DNA Replication, Recombination, and Repair,Energy Production,Nucleic Acid Metabolism,Small Molecule Biochemistry** | Hydrolysis of ATP | 0.0107 | SOD1 | 1 |
| **Behavior** | Climbing behavior | 0.0107 | SOD1 | 1 |
| **Embryonic Development,Nervous System Development and Function,Organ Development,Organismal Development,Tissue Development** | Remyelination of posterior funiculus | 0.0107 | PRL | 1 |
| **Cell Death and Survival** | Cell death of lower motor neuron | 0.0107 | SOD1 | 1 |
| **Metabolic Disease** | Progressive acidosis | 0.0107 | SOD1 | 1 |
| **Dermatological Diseases and Conditions,Neurological Disease,Organismal Injury and Abnormalities** | Hypesthesia | 0.0107 | REST | 1 |
| **Cell Death and Survival** | Cell death of alpha-motor neurons | 0.0107 | SOD1 | 1 |
| **Cell Cycle,Gene Expression** | Binding of Ets element | 0.0107 | SOD1 | 1 |
| **Nervous System Development and Function** | Activation of ventral root | 0.0107 | SOD1 | 1 |
| **Free Radical Scavenging,Molecular Transport** | Accumulation of hydrogen peroxide | 0.0107 | SOD1 | 1 |
| **Cardiovascular Disease,Neurological Disease,Organismal Injury and Abnormalities** | Infarction of cerebral hemisphere | 0.0107 | IL18 | 1 |
| **Cell Death and Survival,Cellular Compromise,Neurological Disease,Organismal Injury and Abnormalities,Tissue Morphology** | Degeneration of subcerebral projection neurons | 0.0107 | SOD1 | 1 |
| **Protein Synthesis** | Oligomerization | 0.0107 | SOD1 | 1 |
| **Nervous System Development and Function,Tissue Morphology** | Quantity of lower motor neuron | 0.0107 | SOD1 | 1 |
| **Cellular Function and Maintenance** | Preservation of motor neurons | 0.0107 | SOD1 | 1 |
| **Cellular Movement,Nervous System Development and Function** | Migration of CA1 neuron | 0.0107 | DISC1 | 1 |
| **Embryonic Development,Hematological System Development and Function,Lymphoid Tissue Structure and Development,Organ Development,Organismal Development,Tissue Development** | Development of lymph follicle | 0.0107 | PDPN | 1 |
| **Neurological Disease,Organismal Injury and Abnormalities** | Atrophy of anterior horn of the spinal cord | 0.0107 | SOD1 | 1 |
| **Cell Morphology,Nervous System Development and Function,Tissue Morphology** | Surface area of dendrites | 0.0107 | DISC1 | 1 |
| **Cellular Assembly and Organization** | Production of cytoplasmic inclusions | 0.0107 | SOD1 | 1 |
| **Cell Signaling,Vitamin and Mineral Metabolism** | Handling of Ca2+ | 0.0107 | SOD1 | 1 |
| **Neurological Disease,Organismal Injury and Abnormalities,Psychological Disorders** | Onset of amyotrophic lateral sclerosis | 0.0107 | SOD1 | 1 |
| **Neurological Disease,Organismal Injury and Abnormalities** | Neuropathy of thalamus | 0.0107 | IL18 | 1 |
| **Tissue Morphology** | Quantity of Streptococcus pneumoniae | 0.0107 | TLR2 | 1 |
| **Neurological Disease,Organismal Injury and Abnormalities** | Neuropathy of hippocampus | 0.0107 | IL18 | 1 |
| **Cellular Development,Cellular Growth and Proliferation,Endocrine System Development and Function,Nervous System Development and Function,Organ Development** | Growth of somatotrophs | 0.0107 | GNAS | 1 |

| | | | | |
|---|---|---|---|---|
| **Cellular Development,Cellular Growth and Proliferation,Hematological System Development and Function,Hematopoiesis,Lymphoid Tissue Structure and Development,Tissue Development** | Proliferation of myeloid progenitor cells | 0.0107 | SOD1 | 1 |
| **Cellular Assembly and Organization** | Quantity of fibrillar inclusions | 0.0107 | SOD1 | 1 |
| **Cell Morphology,Organ Morphology,Organismal Injury and Abnormalities** | Hypertrophy of thyrotropes | 0.0107 | CGA | 1 |
| **Cellular Compromise,Cellular Function and Maintenance** | Endoplasmic reticulum stress response of dopaminergic neurons | 0.0107 | XBP1 | 1 |
| **Cell Death and Survival,Cellular Compromise,Nervous System Development and Function** | Cytotoxicity of motor neurons | 0.0107 | SOD1 | 1 |
| **Hematological Disease,Infectious Diseases,Neurological Disease,Organismal Injury and Abnormalities** | Malaria of brain | 0.0107 | IFNGR1 | 1 |
| **Cellular Assembly and Organization** | Quantity of Golgi vesicles | 0.0107 | SOD1 | 1 |
| **Cancer,Endocrine System Disorders,Organismal Injury and Abnormalities,Reproductive System Disease** | Hyperplasia of pituitary gland | 0.0109 | CGA, PRL | 2 |
| **Cellular Assembly and Organization,Neurological Disease** | Deposition of amyloid-beta plaques | 0.0123 | IL33, SERPINA3 | 2 |
| **Cell-mediated Immune Response,Cellular Movement,Hematological System Development and Function,Immune Cell Trafficking** | Cell movement of T lymphocytes | 0.0123 | CD44, SELPLG | 2 |
| **Cellular Assembly and Organization,Neurological Disease,Organismal Development,Tissue Development** | Formation of amyloid fibrils | 0.0123 | SOD1, TLR4 | 2 |
| **Cell Death and Survival,DNA Replication, Recombination, and Repair** | Fragmentation of DNA | 0.0139 | PRL, SOD1 | 2 |
| **Organismal Injury and Abnormalities,Tissue Morphology** | Size of lesion | 0.014 | IL18, IL1A, SOD1, TLR2, TLR4 | 5 |
| **Neurological Disease** | Neurodegeneration of nerves | 0.0146 | SOD1, TLR2, TLR4 | 3 |
| **Cell Death and Survival,Neurological Disease,Organismal Injury and Abnormalities** | Cell death of pyramidal neurons | 0.0155 | REST, SOD1 | 2 |
| **Cellular Assembly and Organization** | Formation of vacuole | 0.0172 | FUCA1, SOD1 | 2 |
| **Cell-To-Cell Signaling and Interaction,Hematological System Development and Function,Immune Cell Trafficking** | Adhesion of immune cells | 0.0172 | SELPLG, TLR4 | 2 |
| **Endocrine System Development and Function,Endocrine System Disorders,Nervous System Development and Function,Neurological Disease,Organ Morphology,Organismal Development,Organismal Injury and Abnormalities,Reproductive System Disease** | Abnormal morphology of adenohypophysis | 0.019 | CGA, PRL | 2 |
| **Neurological Disease,Organismal Injury and Abnormalities** | Edema of brain | 0.019 | SOD1, TLR4 | 2 |
| **Cell Death and Survival** | Cell death of neuroglia | 0.0202 | CD44, SOD1, TLR2, TLR4 | 4 |
| **Cell Signaling,Small Molecule Biochemistry** | Synthesis of nitric oxide | 0.0209 | TLR2, TLR4 | 2 |
| **Cell-To-Cell Signaling and Interaction,Embryonic Development,Nervous System Development and Function** | Synaptic transmission of pre-Botzinger complex | 0.0213 | CD44 | 1 |
| **Nervous System Development and Function** | Development of phrenic nerve | 0.0213 | CD44 | 1 |
| **Cell-To-Cell Signaling and Interaction,Nervous System Development and Function** | Activation of motor neurons | 0.0213 | SOD1 | 1 |
| **Cell Death and Survival** | Cell death of pre-oligodendrocytes | 0.0213 | SOD1 | 1 |
| **Cellular Assembly and Organization,Cellular Compromise** | Formation of neurofilament inclusion | 0.0213 | SOD1 | 1 |
| **Hematological System Development and Function,Immune Cell Trafficking,Inflammatory Response,Tissue Development** | Accumulation of monocyte-derived macrophages | 0.0213 | SOD1 | 1 |

| | | | | |
|---|---|---|---|---|
| **Cancer,Neurological Disease,Organismal Injury and Abnormalities** | Brain metastasis | 0.0213 | CD44 | 1 |
| **Cell-To-Cell Signaling and Interaction,Nervous System Development and Function** | Compound action potential of optic nerve | 0.0213 | TLR4 | 1 |
| **Cell Signaling,Vitamin and Mineral Metabolism** | Buffering of Ca2+ | 0.0213 | SOD1 | 1 |
| **Organismal Survival** | Delay in organismal death | 0.0213 | SOD1 | 1 |
| **Cell Death and Survival,Cellular Compromise,Neurological Disease,Tissue Morphology** | Degeneration of corticospinal neurons | 0.0213 | SOD1 | 1 |
| **Cell Morphology,Endocrine System Development and Function,Endocrine System Disorders,Nervous System Development and Function,Neurological Disease,Organ Morphology,Organismal Development,Organismal Injury and Abnormalities,Reproductive System Disease** | Lack of lactotropes | 0.0213 | CGA | 1 |
| **Embryonic Development,Tissue Morphology** | Quantity of transit amplifying progenitor cells | 0.0213 | TLR4 | 1 |
| **Cell-To-Cell Signaling and Interaction,Nervous System Development and Function** | Activation of parasympathetic nerve | 0.0213 | REST | 1 |
| **Cardiovascular System Development and Function,Cell-To-Cell Signaling and Interaction** | Activation of endothelial cells | 0.0213 | IL1A | 1 |
| **Cellular Movement,Hematological System Development and Function,Immune Cell Trafficking** | Sequestration of leukocytes | 0.0213 | IFNGR1 | 1 |
| **Neurological Disease,Organismal Injury and Abnormalities,Psychological Disorders** | Delay in amyotrophic lateral sclerosis | 0.0213 | SOD1 | 1 |
| **Hereditary Disorder,Neurological Disease,Organismal Injury and Abnormalities,Psychological Disorders** | Frontotemporal dementia with parkinsonism | 0.0213 | SERPINA3 | 1 |
| **Nervous System Development and Function,Organ Morphology,Organismal Development** | Height of barrel cortex | 0.0213 | IGFBP1 | 1 |
| **Cellular Compromise,Cellular Function and Maintenance** | Endoplasmic reticulum stress response of motor neurons | 0.0213 | SOD1 | 1 |
| **Nervous System Development and Function,Organ Morphology,Organismal Development,Tissue Morphology** | Volume of barrel cortex | 0.0213 | IGFBP1 | 1 |
| **Cell-mediated Immune Response,Cell-To-Cell Signaling and Interaction,Cellular Movement,Hematological System Development and Function,Immune Cell Trafficking** | Adhesion of Th1 cells | 0.0213 | SELPLG | 1 |
| **Cell Death and Survival** | Cell death of cortical astrocytes | 0.0213 | TLR4 | 1 |
| **Cell Morphology,Organ Morphology,Organismal Injury and Abnormalities** | Hypertrophy of gonadotropes | 0.0213 | CGA | 1 |
| **Cell-To-Cell Signaling and Interaction,Nervous System Development and Function** | Binding of neurites | 0.0213 | CD44 | 1 |
| **Nervous System Development and Function,Organ Morphology** | Thickening of brain septum | 0.0213 | CYSLTR1 | 1 |
| **Cardiovascular Disease,Organismal Injury and Abnormalities** | Dysfunction of endothelial tissue | 0.0213 | SOD1 | 1 |
| **Post-Translational Modification** | Tyrosine nitration of protein | 0.0213 | SOD1 | 1 |
| **Cell-mediated Immune Response,Cellular Movement,Hematological System Development and Function,Immune Cell Trafficking** | Cell rolling of Th1 cells | 0.0213 | SELPLG | 1 |
| **Neurological Disease,Organismal Injury and Abnormalities** | Temporal lobe epilepsy with hippocampal sclerosis | 0.0213 | TLR4 | 1 |
| **Neurological Disease,Organismal Injury and Abnormalities,Psychological Disorders** | Corticobasal degeneration | 0.0213 | SERPINA3 | 1 |
| **Neurological Disease,Organismal Injury and Abnormalities** | Gliosis of central nervous system | 0.0219 | AGA, IFNGR1, SOD1 | 3 |
| **Cell Death and Survival,Nervous System Development and Function** | Survival of oligodendrocytes | 0.0228 | IL1A, XAF1 | 2 |

| | | | | |
|---|---|---|---|---|
| **Neurological Disease,Organismal Injury and Abnormalities** | Progressive encephalopathy | 0.023 | GNAS, IL1A, SERPINA3, SOD1 | 4 |
| **Tissue Morphology** | Quantity of cells | 0.025 | AGA, CD86, CGA, DISC1, GNAS, IFNGR1, IGFBP1, IL1A, PRL, PTPRC, SOD1, TLR2, TLR4 | 13 |
| **Cellular Movement,Hematological System Development and Function,Immune Cell Trafficking** | Cell movement of leukocytes | 0.0252 | CD44, SELPLG, TLR4 | 3 |
| **Embryonic Development,Nervous System Development and Function,Organ Development,Organismal Development,Tissue Development** | Neurogenesis of brain | 0.0252 | DISC1, PRL, REST | 3 |
| **Cellular Development,Cellular Growth and Proliferation,Endocrine System Development and Function,Nervous System Development and Function,Organ Development** | Proliferation of lactotropes | 0.0317 | PRL | 1 |
| **Neurological Disease** | Delay in neurodegeneration | 0.0317 | SOD1 | 1 |
| **Cellular Assembly and Organization** | Quantity of vacuole | 0.0317 | SOD1 | 1 |
| **Neurological Disease,Organismal Injury and Abnormalities** | Neurodegeneration of spinal cord | 0.0317 | SOD1 | 1 |
| **Immunological Disease,Neurological Disease** | Autoimmune disease of central nervous system | 0.0317 | CD274 | 1 |
| **Cell Death and Survival,Nervous System Development and Function** | Survival of retinal ganglion cell axon | 0.0317 | XBP1 | 1 |
| **Cardiovascular System Development and Function,Cell-To-Cell Signaling and Interaction** | Interaction of vascular endothelial cells | 0.0317 | CD44 | 1 |
| **Hereditary Disorder,Neurological Disease,Organismal Injury and Abnormalities,Psychological Disorders** | Familial amyotrophic lateral sclerosis | 0.0317 | SOD1 | 1 |
| **Cancer,Endocrine System Disorders,Neurological Disease,Organismal Injury and Abnormalities,Reproductive System Disease** | Hyperplasia of thyrotropes | 0.0317 | CGA | 1 |
| **Cell Death and Survival,Nervous System Development and Function** | Cell survival of oligodendrocytes | 0.0317 | XAF1 | 1 |
| **Cell Morphology,Cellular Function and Maintenance,Nervous System Development and Function,Tissue Morphology** | Length of apical processes | 0.0317 | DISC1 | 1 |
| **Cellular Function and Maintenance,Cellular Growth and Proliferation,Nervous System Development and Function** | Production of neuronal progenitor cells | 0.0317 | PRL | 1 |
| **Nervous System Development and Function,Neurological Disease** | Abnormal morphology of dorsal root | 0.0317 | CD86 | 1 |
| **Hematological System Development and Function,Immune Cell Trafficking,Inflammatory Response,Tissue Development** | Accumulation of regulatory T lymphocytes | 0.0317 | SOD1 | 1 |
| **Antimicrobial Response** | Clearance of measles virus | 0.0317 | IL18 | 1 |
| **Protein Degradation,Protein Synthesis** | Damage of protein | 0.0317 | SOD1 | 1 |
| **Cancer,Hematological Disease,Immunological Disease,Organismal Injury and Abnormalities** | B-cell lymphoma | 0.0317 | IFNGR1 | 1 |
| **Nervous System Development and Function,Organ Morphology,Organismal Development,Tissue Morphology** | Area of barrel cortex | 0.0317 | IGFBP1 | 1 |
| **Cell Morphology,Cellular Function and Maintenance,Nervous System Development and Function,Tissue Morphology** | Length of basal dendrites | 0.0317 | DISC1 | 1 |
| **Cell Signaling,Cellular Function and Maintenance,Molecular Transport,Vitamin and Mineral Metabolism** | Store-operated calcium influx | 0.0317 | SOD1 | 1 |
| **Dermatological Diseases and Conditions,Neurological Disease,Organismal Injury and Abnormalities** | Skin sensation disturbance | 0.0335 | REST, TLR4 | 2 |

| | | | | |
|---|---|---|---|---|
| **Cell Morphology,Nervous System Development and Function,Neurological Disease,Tissue Morphology** | Loss of axons | 0.0358 | SOD1, TLR4 | 2 |
| **Nervous System Development and Function,Tissue Morphology** | Quantity of dopaminergic neurons | 0.0382 | PRL, SOD1 | 2 |
| **Neurological Disease,Organismal Injury and Abnormalities,Psychological Disorders** | Frontotemporal lobar degeneration or amyotrophic lateral sclerosis | 0.0382 | SERPINA3, SOD1 | 2 |
| **Cellular Movement** | Migration of cells | 0.039 | CD44, CXCL2, DISC1, IL33, SELPLG, TLR2, TLR4 | 7 |
| **Nervous System Development and Function,Tissue Morphology** | Quantity of neuroglia | 0.0397 | IFNGR1, IL1A, PTPRC, SOD1 | 4 |
| **Cell Death and Survival** | Apoptosis of neuroglia | 0.0404 | CD44, TLR2, TLR4 | 3 |
| **Cell-To-Cell Signaling and Interaction,Drug Metabolism,Molecular Transport,Small Molecule Biochemistry** | Concentration of dopamine | 0.0404 | PRL, SOD1, TLR4 | 3 |
| **Nervous System Development and Function,Tissue Morphology** | Quantity of astrocytes | 0.0406 | IFNGR1, SOD1 | 2 |
| **Cell Death and Survival,Cellular Compromise,Neurological Disease,Tissue Morphology** | Neurodegeneration of dopaminergic neurons | 0.0406 | SOD1, XBP1 | 2 |
| **Embryonic Development,Nervous System Development and Function,Organ Development,Organismal Development,Tissue Development** | Neurogenesis of olfactory bulb | 0.0421 | PRL | 1 |
| **Cell Death and Survival,Cellular Compromise,Neurological Disease,Tissue Morphology** | Neurodegeneration of interneurons | 0.0421 | SOD1 | 1 |
| **Cell Death and Survival,Nervous System Development and Function** | Cell survival of dopaminergic neurons | 0.0421 | XBP1 | 1 |
| **Energy Production,Molecular Transport,Nucleic Acid Metabolism,Small Molecule Biochemistry** | Release of ATP | 0.0421 | SOD1 | 1 |
| **Cell Morphology,Cellular Compromise** | Swelling of mitochondria | 0.0421 | SOD1 | 1 |
| **Post-Translational Modification** | Carbonylation of protein | 0.0421 | SOD1 | 1 |
| **Cellular Assembly and Organization** | Formation of cytoplasmic aggregates | 0.0421 | SOD1 | 1 |
| **Cellular Assembly and Organization** | Accumulation of PAS-positive granule | 0.0421 | IL33 | 1 |
| **Cellular Assembly and Organization,Cellular Compromise,Cellular Development,Cellular Growth and Proliferation,Nervous System Development and Function,Tissue Development** | Development of peripherin inclusion | 0.0421 | SOD1 | 1 |
| **Cellular Assembly and Organization,Cellular Compromise,Nervous System Development and Function** | Formation of intermediate filament inclusions | 0.0421 | SOD1 | 1 |
| **Nervous System Development and Function,Neurological Disease,Organ Morphology,Organismal Development,Organismal Injury and Abnormalities** | Abnormal morphology of enlarged fourth cerebral ventricle | 0.0421 | COL18A1 | 1 |
| **Cell-To-Cell Signaling and Interaction,Cellular Movement,Hematological System Development and Function,Immune Cell Trafficking,Nervous System Development and Function** | Recruitment of microglia | 0.0421 | IL33 | 1 |
| **Cell Death and Survival,Cell Morphology,Cellular Compromise,Nervous System Development and Function** | Vacuolation of neurons | 0.0421 | AGA | 1 |
| **Small Molecule Biochemistry** | Synthesis of catecholamine | 0.0421 | PRL | 1 |
| **Nervous System Development and Function** | Prepulse inhibition | 0.0421 | GNAS | 1 |
| **Cell Death and Survival,Cellular Compromise,Neurological Disease,Tissue Morphology** | Degeneration of motor axons | 0.0421 | SOD1 | 1 |

| | | | | |
|---|---|---|---|---|
| **Cardiovascular System Development and Function,Cellular Development,Cellular Function and Maintenance,Cellular Growth and Proliferation,Organismal Development,Tissue Development** | Proliferation of microvascular endothelial cells | 0.0421 | CD44 | 1 |
| **Cellular Function and Maintenance,Cellular Growth and Proliferation,Nervous System Development and Function** | Production of granule cells | 0.0421 | REST | 1 |
| **Cell Morphology,Cellular Assembly and Organization,Nervous System Development and Function,Tissue Morphology** | Area of perikaryon | 0.0421 | IGFBP1 | 1 |
| **Cellular Compromise** | Damage of mitochondria | 0.0421 | SOD1 | 1 |
| **Inflammatory Disease,Inflammatory Response,Neurological Disease,Organismal Injury and Abnormalities** | HIV-associated encephalitis | 0.0421 | PTPRC | 1 |
| **Cellular Function and Maintenance,Cellular Growth and Proliferation** | Production of cells | 0.0457 | PRL, REST | 2 |
| **Cell-To-Cell Signaling and Interaction,Hematological System Development and Function,Immune Cell Trafficking,Inflammatory Response,Nervous System Development and Function** | Activation of microglia | 0.0462 | SOD1, TLR2, TLR4 | 3 |
| **Cellular Development,Cellular Growth and Proliferation,Nervous System Development and Function** | Proliferation of neuroglia | 0.0472 | CXCL2, IFNGR1, PRL, SOD1 | 4 |

### Cluster 3

| Categories | Diseases or Functions Annotation | p-Value | Molecules | # Molecules |
|---|---|---|---|---|
| **Cell Cycle** | Mitogenesis of central nervous system cells | 0.00119 | EDN1, EGF | 2 |
| **Cell Morphology** | Modification of neurons | 0.00366 | EDN1, NBN | 2 |
| **Cell-To-Cell Signaling and Interaction** | Binding of brain cells | 0.00499 | CX3CL1, EGF | 2 |
| **Carbohydrate Metabolism,Lipid Metabolism,Small Molecule Biochemistry** | Hydrolysis of phosphatidylinositol polyphosphate | 0.00767 | EDN1 | 1 |
| **Cell Morphology** | Elongation of dopaminergic neurons | 0.00767 | EGF | 1 |
| **Cellular Growth and Proliferation,Nervous System Development and Function,Organ Development** | Proliferation of stellate cells | 0.00767 | EDN1 | 1 |
| **Nervous System Development and Function,Tissue Morphology** | Density of serotonergic neurons | 0.00767 | GAP43 | 1 |
| **Cellular Development,Cellular Growth and Proliferation,Nervous System Development and Function,Organ Development,Tissue Development** | Expansion of mesencephalic neurons | 0.00767 | EGF | 1 |
| **Embryonic Development,Nervous System Development and Function,Organ Development,Organismal Development,Tissue Development** | Development of layer IV | 0.00767 | GAP43 | 1 |
| **Connective Tissue Disorders,Inflammatory Disease,Inflammatory Response,Organismal Injury and Abnormalities,Skeletal and Muscular Disorders** | Degeneration of lumbar intervertebral disc | 0.00767 | EDN1 | 1 |
| **Behavior** | Active avoidance response | 0.00767 | EGF | 1 |
| **Nervous System Development and Function** | Mechanical nociception | 0.00767 | CX3CL1 | 1 |
| **Nervous System Development and Function,Neurological Disease,Organ Morphology,Organismal Development,Organismal Injury and Abnormalities** | Lack of optic tract | 0.00767 | GAP43 | 1 |
| **Nervous System Development and Function,Organ Morphology,Organismal Development** | Size of optic chiasm | 0.00767 | GAP43 | 1 |

| | | | | |
|---|---|---|---|---|
| **Drug Metabolism,Molecular Transport** | Efflux of glutathione | 0.00767 | EDN1 | 1 |
| **Cell Morphology,Cellular Movement,Nervous System Development and Function** | Innervation of serotonergic neurons | 0.00767 | GAP43 | 1 |
| **Cell-To-Cell Signaling and Interaction** | Priming of neural stem cells | 0.00767 | EGF | 1 |
| **Cell Morphology** | Contractility of stellate cells | 0.00767 | EDN1 | 1 |
| **Cardiovascular Disease,Developmental Disorder,Organismal Injury and Abnormalities** | Congenital heart block | 0.00767 | EDN1 | 1 |
| **Cell Morphology,Cellular Assembly and Organization** | Elongation of astrocyte projections | 0.00767 | GAP43 | 1 |
| **Hereditary Disorder,Neurological Disease,Organismal Injury and Abnormalities,Psychological Disorders,Skeletal and Muscular Disorders** | Parkinson-dementia syndrome | 0.00767 | GAP43 | 1 |
| **Neurological Disease** | Ataxia | 0.0101 | FAS, NBN | 2 |
| **Cell Death and Survival,Neurological Disease,Organismal Injury and Abnormalities** | Apoptosis of lactotropes | 0.0153 | FAS | 1 |
| **Organismal Functions** | Recovery of mice | 0.0153 | FAS | 1 |
| **Cell-To-Cell Signaling and Interaction,Endocrine System Development and Function** | Binding of anterior pituitary cells | 0.0153 | EGF | 1 |
| **Cellular Development,Cellular Growth and Proliferation,Nervous System Development and Function,Tissue Development** | Expansion of dopaminergic neurons | 0.0153 | EGF | 1 |
| **Cell Cycle,Nervous System Development and Function** | Mitogenesis of astrocytes | 0.0153 | EDN1 | 1 |
| **Neurological Disease,Skeletal and Muscular Disorders** | Hypernociception of hindpaw | 0.0153 | CX3CL1 | 1 |
| **Cell Morphology,Cellular Assembly and Organization,Cellular Development,Cellular Function and Maintenance,Cellular Growth and Proliferation,Embryonic Development,Nervous System Development and Function,Organismal Development,Tissue Development** | Branching of sympathetic neurites | 0.0153 | GAP43 | 1 |
| **Cell Cycle,Endocrine System Development and Function,Nervous System Development and Function** | Mitogenesis of corticotroph cells | 0.0153 | EGF | 1 |
| **Cell Death and Survival,Cellular Development,Embryonic Development,Organismal Development,Tissue Development,Tissue Morphology** | Regeneration of neural stem cells | 0.0153 | EGF | 1 |
| **Cell Morphology,Cellular Function and Maintenance** | Repair of neurons | 0.0153 | NBN | 1 |
| **Cellular Movement,Nervous System Development and Function** | Migration of hippocampal neurons | 0.0153 | CX3CL1 | 1 |
| **Cell Death and Survival,Neurological Disease,Organismal Injury and Abnormalities** | Apoptosis of somatotrophs | 0.0153 | FAS | 1 |
| **Cell Morphology,Cellular Function and Maintenance,DNA Replication, Recombination, and Repair** | Double-stranded DNA break repair of cells | 0.0153 | NBN | 1 |
| **Cellular Development,Cellular Growth and Proliferation,Embryonic Development,Organismal Development,Tissue Development** | Generation of embryoid body cells | 0.0153 | EGF | 1 |
| **Cell Death and Survival,Cellular Development,Nervous System Development and Function,Organ Morphology,Tissue Development,Tissue Morphology** | Regeneration of pyramidal neurons | 0.0153 | EGF | 1 |
| **Tissue Development** | Formation of white matter | 0.0153 | NBN | 1 |
| **Neurological Disease,Organismal Injury and Abnormalities,Psychological Disorders** | Preclinical stage amyotrophic lateral sclerosis | 0.0153 | FAS | 1 |
| **Hematological System Development and Function,Inflammatory Response,Tissue Morphology** | Quantity of monocytes | 0.0153 | IL12RB2 | 1 |
| **Cell Morphology,Nervous System Development and Function** | Modulation of dopaminergic neurons | 0.0153 | EDN1 | 1 |
| **Cardiovascular System Development and Function,Tissue Morphology** | Contraction of cerebral artery | 0.0153 | EDN1 | 1 |

| | | | | |
|---|---|---|---|---|
| **Embryonic Development,Nervous System Development and Function,Organ Development,Organismal Development,Tissue Development** | Formation of corpus callosum | 0.022 | GAP43, NIN | 2 |
| **Embryonic Development,Nervous System Development and Function,Organ Development,Organismal Development,Tissue Development** | Formation of optic tract | 0.0228 | GAP43 | 1 |
| **Cell-To-Cell Signaling and Interaction,Nervous System Development and Function** | Adhesion of hippocampal neurons | 0.0228 | CX3CL1 | 1 |
| **Cardiovascular System Development and Function** | Vasoconstriction of middle cerebral artery | 0.0228 | EDN1 | 1 |
| **Ophthalmic Disease,Organismal Injury and Abnormalities** | Proliferative vitreoretinopathy | 0.0228 | EDN1 | 1 |
| **Cellular Movement,Hematological System Development and Function,Immune Cell Trafficking,Inflammatory Response,Nervous System Development and Function** | Migration of microglia | 0.0228 | CX3CL1 | 1 |
| **Cellular Development,Cellular Growth and Proliferation,Embryonic Development,Nervous System Development and Function,Organismal Development,Tissue Development** | Generation of neuroblasts | 0.0228 | EGF | 1 |
| **Lipid Metabolism,Small Molecule Biochemistry** | Turnover of inositol phospholipid | 0.0228 | EDN1 | 1 |
| **Cell-To-Cell Signaling and Interaction,Cellular Function and Maintenance,Inflammatory Response,Nervous System Development and Function** | Phagocytosis of neurons | 0.0228 | CX3CL1 | 1 |
| **Cell-To-Cell Signaling and Interaction,Endocrine System Development and Function,Nervous System Development and Function** | Desensitization of pituitary cells | 0.0228 | EDN1 | 1 |
| **Cell Morphology** | Sphere formation of stem cells | 0.0228 | EGF | 1 |
| **Cell Morphology,Nervous System Development and Function** | Contraction of stellate cells | 0.0228 | EDN1 | 1 |
| **Cellular Development,Cellular Growth and Proliferation,Nervous System Development and Function,Tissue Development** | Expansion of cholinergic neurons | 0.0228 | EGF | 1 |
| **Cell-To-Cell Signaling and Interaction,Nervous System Development and Function** | Adhesion of cerebellar granule cell | 0.0228 | CX3CL1 | 1 |
| **Neurological Disease,Organismal Injury and Abnormalities,Psychological Disorders** | Multiple system atrophy | 0.0228 | FAS | 1 |
| **Cell Signaling,Molecular Transport,Vitamin and Mineral Metabolism** | Quantity of Ca2+ | 0.0283 | CCL1, CX3CL1, EDN1 | 3 |
| **Embryonic Development,Nervous System Development and Function,Organ Development,Organismal Development,Tissue Development** | Formation of hippocampal commissure | 0.0303 | GAP43 | 1 |
| **Amino Acid Metabolism,Lipid Metabolism,Small Molecule Biochemistry** | Synthesis of GABA | 0.0303 | EGF | 1 |
| **Nervous System Development and Function** | Prepulse inhibition | 0.0303 | EGF | 1 |
| **Post-Translational Modification** | Activation of protein | 0.0303 | EGF | 1 |
| **Nervous System Development and Function,Neurological Disease,Organ Morphology,Organismal Development,Organismal Injury and Abnormalities** | Abnormal morphology of disorganized barrel cortex | 0.0303 | GAP43 | 1 |
| **Connective Tissue Disorders,Developmental Disorder,Organismal Injury and Abnormalities,Skeletal and Muscular Disorders** | Microcephaly | 0.0378 | NBN | 1 |
| **Embryonic Development,Nervous System Development and Function,Organ Development,Organ Morphology,Organismal Development,Tissue Development** | Morphogenesis of corpus callosum | 0.0378 | NIN | 1 |
| **DNA Replication, Recombination, and Repair** | Mitochondrial DNA damage | 0.0378 | OGG1 | 1 |
| **Cell Morphology,Nervous System Development and Function** | Innervation of optic nerve | 0.0378 | GAP43 | 1 |
| **Cell Death and Survival** | Apoptosis of oligodendrocytes | 0.0394 | NBN, OGG1 | 2 |

| | | | | |
|---|---|---|---|---|
| **Embryonic Development,Organismal Development,Tissue Development** | Growth of embryonic tissue | 0.044 | CX3CL1, EGF, GAP43, NBN | 4 |
| **Endocrine System Development and Function,Nervous System Development and Function,Organ Morphology,Tissue Morphology** | Quantity of corticotroph cells | 0.0452 | EGF | 1 |
| **Reproductive System Development and Function** | Lordosis | 0.0452 | EGF | 1 |
| **Endocrine System Disorders,Gastrointestinal Disease,Metabolic Disease,Organismal Injury and Abnormalities** | Diabetes mellitus | 0.0452 | FAS | 1 |
| **Cellular Function and Maintenance,Nervous System Development and Function,Tissue Development** | Function of GABAergic neurons | 0.0452 | EGF | 1 |

### Cluster 5

| Categories | Diseases or Functions Annotation | p-Value | Molecules | # Molecules |
|---|---|---|---|---|
| **Cell-To-Cell Signaling and Interaction,Nervous System Development and Function** | Sensitization of trigeminovascular neuron | 0.0062 | KNG1 | 1 |
| **Nervous System Development and Function** | Mechanical nociception | 0.0062 | CX3CL1 | 1 |
| **Inflammatory Response** | Inflammatory response of cerebral cortex | 0.0062 | KNG1 | 1 |
| **Cardiovascular System Development and Function,Nervous System Development and Function,Organ Morphology,Organismal Development,Tissue Morphology** | Diameter of basilar artery | 0.0062 | KNG1 | 1 |
| **Cell-To-Cell Signaling and Interaction,Nervous System Development and Function** | Firing of trigeminovascular neuron | 0.0062 | KNG1 | 1 |
| **Cell Death and Survival,Neurological Disease,Organismal Injury and Abnormalities** | Apoptosis of lactotropes | 0.0124 | FAS | 1 |
| **Organismal Functions** | Recovery of mice | 0.0124 | FAS | 1 |
| **Lipid Metabolism,Small Molecule Biochemistry** | Liberation of arachidonic acid | 0.0124 | KNG1 | 1 |
| **Neurological Disease,Skeletal and Muscular Disorders** | Hypernociception of hindpaw | 0.0124 | CX3CL1 | 1 |
| **Neurological Disease,Organismal Injury and Abnormalities,Psychological Disorders** | Preclinical stage amyotrophic lateral sclerosis | 0.0124 | FAS | 1 |
| **Cellular Movement,Nervous System Development and Function** | Migration of hippocampal neurons | 0.0124 | CX3CL1 | 1 |
| **Cell-To-Cell Signaling and Interaction,Nervous System Development and Function** | Action potential of superior cervical ganglion neurons | 0.0124 | KNG1 | 1 |
| **Lipid Metabolism,Small Molecule Biochemistry** | Liberation of stearic acid | 0.0124 | KNG1 | 1 |
| **Cell Death and Survival,Neurological Disease,Organismal Injury and Abnormalities** | Apoptosis of somatotrophs | 0.0124 | FAS | 1 |
| **Cardiovascular Disease,Hematological Disease,Organismal Injury and Abnormalities** | Thrombus | 0.0124 | KNG1 | 1 |
| **Cell Signaling,Molecular Transport,Vitamin and Mineral Metabolism** | Quantity of Ca2+ | 0.0163 | CCL1, CX3CL1, KNG1 | 3 |
| **Cell-To-Cell Signaling and Interaction,Cellular Function and Maintenance,Inflammatory Response,Nervous System Development and Function** | Phagocytosis of neurons | 0.0185 | CX3CL1 | 1 |
| **Cell-To-Cell Signaling and Interaction,Nervous System Development and Function** | Adhesion of hippocampal neurons | 0.0185 | CX3CL1 | 1 |
| **Carbohydrate Metabolism,Molecular Transport,Small Molecule Biochemistry** | Accumulation of inositol phosphate | 0.0185 | KNG1 | 1 |
| **Cellular Movement,Hematological System Development and Function,Immune Cell Trafficking,Inflammatory Response,Nervous System Development and Function** | Migration of microglia | 0.0185 | CX3CL1 | 1 |

| | | | | |
|---|---|---|---|---|
| **Cell-To-Cell Signaling and Interaction,Nervous System Development and Function** | Adhesion of cerebellar granule cell | 0.0185 | CX3CL1 | 1 |
| **Lipid Metabolism,Small Molecule Biochemistry** | Synthesis of 12-hydroxyeicosatetraenoic acid | 0.0185 | KNG1 | 1 |
| **Neurological Disease,Organismal Injury and Abnormalities,Psychological Disorders** | Multiple system atrophy | 0.0185 | FAS | 1 |
| **Cellular Function and Maintenance,Cellular Growth and Proliferation** | Production of neural precursor cells | 0.0246 | MCPH1 | 1 |
| **Cell Morphology,Cell-To-Cell Signaling and Interaction,Cellular Function and Maintenance,Nervous System Development and Function** | Depolarization of superior cervical ganglion neurons | 0.0246 | KNG1 | 1 |
| **Carbohydrate Metabolism,Lipid Metabolism,Small Molecule Biochemistry** | Hydrolysis of phosphatidylinositol 4,5-diphosphate | 0.0246 | KNG1 | 1 |
| **Protein Trafficking** | Interaction of protein | 0.0246 | KNG1 | 1 |
| **Endocrine System Disorders,Gastrointestinal Disease,Metabolic Disease,Organismal Injury and Abnormalities** | Diabetes mellitus | 0.0367 | FAS | 1 |
| **Cell Signaling,Nucleic Acid Metabolism,Small Molecule Biochemistry** | Binding of guanosine 5'-O-(3-thiotriphosphate) | 0.0426 | KNG1 | 1 |
| **Cell-To-Cell Signaling and Interaction,Cellular Growth and Proliferation,Nervous System Development and Function** | Excitation of sensory neurons | 0.0486 | KNG1 | 1 |
| **Cardiovascular System Development and Function,Cell-To-Cell Signaling and Interaction** | Adhesion of endothelial cells | 0.0486 | FAS | 1 |
| **Cell Death and Survival,Embryonic Development** | Cell death of neuroblasts | 0.0486 | FAS | 1 |
| **Cellular Movement,Hematological System Development and Function,Immune Cell Trafficking** | Cell rolling of leukocytes | 0.0486 | FAS | 1 |

### Supplementary Table 3: Module 3
### Cluster 7

| Categories | Diseases or Functions Annotation | p-Value | Molecules | # Molecules |
|---|---|---|---|---|
| **Hereditary Disorder,Neurological Disease,Organismal Injury and Abnormalities** | Autosomal dominant encephalopathy | 0.0000447 | GAP43, MAPT, SNCA | 3 |
| **Neurological Disease,Organismal Injury and Abnormalities** | Syndromic encephalopathy | 0.0000667 | FAS, MAPT, SNCA | 3 |
| **Hereditary Disorder,Organismal Injury and Abnormalities** | MAPT-related disorder | 0.0000889 | GAP43, MAPT | 2 |
| **Hereditary Disorder,Neurological Disease,Organismal Injury and Abnormalities,Psychological Disorders,Skeletal and Muscular Disorders** | Autosomal dominant Parkinson disease | 0.0000889 | GAP43, SNCA | 2 |
| **Neurological Disease,Organismal Injury and Abnormalities,Psychological Disorders** | Multiple system atrophy | 0.000265 | FAS, SNCA | 2 |
| **Metabolic Disease,Neurological Disease,Organismal Injury and Abnormalities,Psychological Disorders** | Braak stage V-VI Alzheimer disease | 0.000265 | GSK3B, MAPT | 2 |
| **Neurological Disease,Organismal Injury and Abnormalities,Psychological Disorders** | Tauopathy of hippocampus | 0.000527 | GSK3B, MAPT | 2 |
| **Metabolic Disease,Neurological Disease,Organismal Injury and Abnormalities,Psychological Disorders** | Braak stage III-IV Alzheimer disease | 0.000527 | GSK3B, MAPT | 2 |
| **Cell-To-Cell Signaling and Interaction,Nervous System Development and Function** | Long-term potentiation of collateral synapses | 0.000838 | GSK3B, LCMT1, MAPT | 3 |
| **Hereditary Disorder,Neurological Disease,Organismal Injury and Abnormalities,Psychological Disorders** | Familial dementia | 0.000872 | GAP43, MAPT | 2 |
| **Cellular Compromise** | Fragmentation of mitochondria | 0.00306 | MAPT, SNCA | 2 |
| **Cell Death and Survival** | Apoptosis of dopaminergic neurons | 0.00306 | FAS, SNCA | 2 |
| **Developmental Disorder,Nervous System Development and Function,Neurological** | Agenesis of anterior commissure | 0.00306 | FBXO45, VTI1B | 2 |

| | | | | |
|---|---|---|---|---|
| **Disease,Organ Morphology,Organismal Development,Organismal Injury and Abnormalities** | | | | |
| **Nervous System Development and Function,Neurological Disease,Organ Morphology,Organismal Development,Organismal Injury and Abnormalities** | Abnormal morphology of optic tract | 0.00306 | GAP43, VTI1B | 2 |
| **Neurological Disease,Organismal Injury and Abnormalities** | Neurodegeneration of hippocampal CA3 region | 0.00381 | MAPT, SNCA | 2 |
| **Cellular Assembly and Organization,Cellular Function and Maintenance,Cellular Movement,Nervous System Development and Function** | Axonal transport of mitochondria | 0.00381 | GSK3B, MAPT | 2 |
| **Neurological Disease,Psychological Disorders** | Schizophrenia | 0.0041 | GAP43, GOT1, GSK3B, MAP10, OXCT1, RCAN2, RTN1 | 7 |
| **Cellular Assembly and Organization,Cellular Compromise,Cellular Development,Cellular Growth and Proliferation,Nervous System Development and Function,Tissue Development** | Formation of neurofibrillary tangles | 0.00462 | GSK3B, MAPT | 2 |
| **Neurological Disease,Organismal Injury and Abnormalities,Psychological Disorders,Skeletal and Muscular Disorders** | Parkinson disease with dementia | 0.00462 | GAP43, SNCA | 2 |
| **Embryonic Development,Nervous System Development and Function,Organ Development,Organismal Development,Tissue Development** | Formation of anterior commissure | 0.00551 | FBXO45, GAP43 | 2 |
| **Cell Morphology,Cellular Assembly and Organization,Cellular Development,Cellular Function and Maintenance,Cellular Growth and Proliferation,Nervous System Development and Function,Organismal Development,Tissue Development** | Polarization of neurons | 0.00551 | GSK3B, MAPT | 2 |
| **Cellular Assembly and Organization,Cellular Compromise** | Formation of cellular inclusion bodies | 0.00633 | GSK3B, MAPT, SNCA | 3 |
| **Neurological Disease,Organismal Injury and Abnormalities** | Neurodegeneration of hippocampus | 0.00679 | GSK3B, MAPT, SNCA | 3 |
| **Cellular Assembly and Organization,Cellular Function and Maintenance** | Stabilization of microtubules | 0.00751 | GSK3B, MAPT | 2 |
| **Cellular Compromise,Lipid Metabolism,Small Molecule Biochemistry** | Peroxidation of lipid | 0.00751 | MAPT, SNCA | 2 |
| **Nervous System Development and Function,Tissue Morphology** | Density of neurites | 0.00751 | MAPT, SNCA | 2 |
| **Hematological System Development and Function,Tissue Morphology** | Quantity of antigen presenting cells | 0.00826 | IL12RB2, MAPT, SNCA | 3 |
| **Nervous System Development and Function** | Development of optic nerve | 0.00861 | THRB, TMEM126A | 2 |
| **Cell Cycle,Gene Expression** | Binding of DNA | 0.00861 | MAPT, YY1 | 2 |
| **Hematological System Development and Function,Tissue Morphology** | Quantity of myeloid cells | 0.00879 | IL12RB2, MAPT, SNCA | 3 |
| **Neurological Disease,Organismal Injury and Abnormalities** | Astrocytosis | 0.00934 | GSK3B, MAPT, SNCA | 3 |
| **Cell Death and Survival,Cellular Compromise,Neurological Disease,Tissue Morphology** | Neurodegeneration of nigrostriatal dopamine system neuron | 0.00947 | SNCA | 1 |
| **Cell Morphology** | Polarity of granule cells | 0.00947 | PAK1 | 1 |
| **Nervous System Development and Function,Tissue Morphology** | Density of serotonergic neurons | 0.00947 | GAP43 | 1 |
| **Post-Translational Modification,Protein Degradation,Protein Synthesis** | Hydrolysis of protein | 0.00947 | SNCA | 1 |
| **Cellular Assembly and Organization,Tissue Development** | Formation of tau filament | 0.00947 | MAPT | 1 |
| **Neurological Disease,Ophthalmic Disease,Organismal Injury and Abnormalities,Psychological Disorders** | Neuroaxonal dystrophy of optic nerve | 0.00947 | MAPT | 1 |

| | | | | |
|---|---|---|---|---|
| **Cellular Compromise,Neurological Disease,Organismal Injury and Abnormalities** | Degeneration of anterior bulbar cells | 0.00947 | MAPT | 1 |
| **Cell Morphology,Cellular Assembly and Organization,Cellular Development,Cellular Function and Maintenance,Cellular Growth and Proliferation,Nervous System Development and Function,Organismal Development,Tissue Development** | Initiation of formation of dendrites | 0.00947 | PAK1 | 1 |
| **Cell-To-Cell Signaling and Interaction,Nervous System Development and Function** | Firing of striatal neurons | 0.00947 | MAPT | 1 |
| **Embryonic Development,Nervous System Development and Function,Organ Development,Organismal Development,Tissue Development** | Development of layer IV | 0.00947 | GAP43 | 1 |
| **Cell Morphology** | Permeabilization of vesicles | 0.00947 | SNCA | 1 |
| **Neurological Disease,Organismal Injury and Abnormalities** | Atrophy of pontine nucleus | 0.00947 | SNCA | 1 |
| **Nervous System Development and Function** | Myelination of vestibulocochlear nerve | 0.00947 | THRB | 1 |
| **Cellular Compromise,Neurological Disease** | Degeneration of glial cell projections | 0.00947 | MAPT | 1 |
| **Nervous System Development and Function,Neurological Disease,Organ Morphology,Organismal Development,Organismal Injury and Abnormalities** | Lack of optic tract | 0.00947 | GAP43 | 1 |
| **Cell Morphology,Cellular Assembly and Organization,Cellular Compromise,Cellular Function and Maintenance** | Loss of microtubules | 0.00947 | MAPT | 1 |
| **Cell-To-Cell Signaling and Interaction,Cellular Assembly and Organization,Nervous System Development and Function** | Binding of synaptic vesicles | 0.00947 | SNCA | 1 |
| **Cellular Assembly and Organization,Cellular Function and Maintenance** | Fluidity of mitochondrial membrane | 0.00947 | SNCA | 1 |
| **Neurological Disease,Organismal Injury and Abnormalities,Psychological Disorders** | Tauopathy of neurons | 0.00947 | MAPT | 1 |
| **Hereditary Disorder,Neurological Disease,Organismal Injury and Abnormalities,Psychological Disorders** | Pick disease | 0.00947 | MAPT | 1 |
| **Cell Morphology,Cellular Assembly and Organization** | Integrity of mitochondrial membrane | 0.00947 | MAPT | 1 |
| **Cellular Development,Cellular Growth and Proliferation,Nervous System Development and Function,Organ Development,Tissue Development** | Proliferation of hippocampal neurons | 0.00947 | MAPT | 1 |
| **Nervous System Development and Function,Organ Morphology,Organismal Development** | Size of optic chiasm | 0.00947 | GAP43 | 1 |
| **Lipid Metabolism,Molecular Transport,Small Molecule Biochemistry** | Quantity of cardiolipin | 0.00947 | SNCA | 1 |
| **Embryonic Development,Nervous System Development and Function,Organ Development,Organismal Development,Tissue Development** | Development of superior temporal gyrus | 0.00947 | GSK3B | 1 |
| **Neurological Disease,Organismal Injury and Abnormalities,Psychological Disorders,Skeletal and Muscular Disorders** | Sporadic Parkinson disease | 0.00947 | SNCA | 1 |
| **Hereditary Disorder,Neurological Disease,Organismal Injury and Abnormalities,Psychological Disorders,Skeletal and Muscular Disorders** | Parkinson-dementia syndrome | 0.00947 | GAP43 | 1 |
| **Cellular Assembly and Organization,Cellular Movement,Nervous System Development and Function** | Translocation of neurofilaments | 0.00947 | MAPT | 1 |
| **Cell Morphology,Cellular Assembly and Organization** | Shape of mitochondria | 0.00947 | SNCA | 1 |
| **Cellular Assembly and Organization,Cellular Compromise** | Formation of fibrillar inclusions | 0.00947 | MAPT | 1 |
| **Neurological Disease,Organismal Injury and Abnormalities** | Mild astrocytosis | 0.00947 | MAPT | 1 |

| | | | | |
|---|---|---|---|---|
| **Developmental Disorder,Hereditary Disorder,Metabolic Disease,Neurological Disease,Organismal Injury and Abnormalities,Psychological Disorders** | Alzheimers disease type 3 | 0.00947 | MAPT | 1 |
| **Cell Cycle,Gene Expression** | Binding of cAMP response element | 0.00947 | MAPT | 1 |
| **Post-Translational Modification** | Ubiquitination of protein | 0.00947 | MAPT | 1 |
| **Cellular Function and Maintenance,Molecular Transport** | Exocytosis of L-glutamic acid | 0.00947 | SNCA | 1 |
| **Cell Morphology,Cellular Assembly and Organization,Cellular Development,Cellular Function and Maintenance,Cellular Growth and Proliferation,Nervous System Development and Function,Organismal Development,Tissue Development** | Formation of Lewy neurites | 0.00947 | SNCA | 1 |
| **Carbohydrate Metabolism,Lipid Metabolism,Molecular Transport,Small Molecule Biochemistry** | Quantity of phosphatidylglycerol | 0.00947 | SNCA | 1 |
| **Cell Morphology,Cellular Movement,Nervous System Development and Function** | Innervation of serotonergic neurons | 0.00947 | GAP43 | 1 |
| **Organismal Injury and Abnormalities** | Dysfunction of mice | 0.00947 | MAPT | 1 |
| **Cell Morphology,Cellular Assembly and Organization** | Elongation of astrocyte projections | 0.00947 | GAP43 | 1 |
| **Hair and Skin Development and Function** | Pallor | 0.00947 | SNCA | 1 |
| **Cell Morphology,Cellular Assembly and Organization,Cellular Function and Maintenance** | Fluidity of plasma membrane | 0.00947 | SNCA | 1 |
| **Cell-To-Cell Signaling and Interaction,Nervous System Development and Function** | Long-term potentiation of cerebral cortex | 0.00969 | CHRD, GSK3B, MAPT, PAK1, SNCA | 5 |
| **Cellular Assembly and Organization** | Quantity of cellular inclusion bodies | 0.00978 | MAPT, SNCA | 2 |
| **Cellular Assembly and Organization,Neurological Disease** | Deposition of amyloid-beta plaques | 0.00978 | MAPT, SNCA | 2 |
| **Cellular Assembly and Organization,Cellular Function and Maintenance,Nervous System Development and Function,Tissue Morphology** | Quantity of neurites | 0.0108 | GSK3B, MAPT, PAK1, SNCA | 4 |
| **Neurological Disease,Organismal Injury and Abnormalities** | Microgliosis | 0.011 | MAPT, SNCA | 2 |
| **Cell Death and Survival,Cellular Compromise,Neurological Disease,Organismal Injury and Abnormalities,Tissue Morphology** | Neurodegeneration of hippocampal neurons | 0.011 | GSK3B, MAPT | 2 |
| **Cell Death and Survival,Neurological Disease,Organismal Injury and Abnormalities** | Cell death of cortical neurons | 0.0113 | FAS, GSK3B, MAPT, MEF2C, PAK1, SNCA | 6 |
| **Cell Death and Survival,Neurological Disease,Organismal Injury and Abnormalities** | Cell death of cerebellar cortex cells | 0.0124 | FAS, GSK3B, MAPT | 3 |
| **Nervous System Development and Function,Tissue Morphology** | Density of neurons | 0.0125 | GAP43, MAPT, MEF2C, PAK1, SNCA | 5 |
| **Cellular Function and Maintenance,Nervous System Development and Function,Tissue Development** | Function of neurons | 0.0131 | COX8A, MAPT, THRB | 3 |
| **Cell Morphology,Cellular Assembly and Organization,Cellular Development,Cellular Function and Maintenance,Cellular Growth and Proliferation,Embryonic Development,Nervous System Development and Function,Neurological Disease,Organismal Development,Tissue Development,Tissue Morphology** | Loss of dendritic spines | 0.0137 | GSK3B, MAPT | 2 |
| **Cell Morphology,Cellular Assembly and Organization,Cellular Development,Cellular Function and Maintenance,Cellular Growth and Proliferation,Embryonic Development,Nervous System Development and Function,Organismal Development,Tissue Development,Tissue Morphology** | Morphology of dendritic spines | 0.0139 | GSK3B, MAGI2, MAPT, PAK1 | 4 |

| | | | | |
|---|---|---|---|---|
| **Neurological Disease,Organismal Injury and Abnormalities,Psychological Disorders** | Parkinsonism | 0.0139 | GAP43, MAPT, SNCA, TUBA1A | 4 |
| **Cellular Development,Cellular Growth and Proliferation,Nervous System Development and Function,Tissue Development** | Growth of axons | 0.0141 | GAP43, GSK3B, MAPT, RNF6, RTN1, VTI1B | 6 |
| **Cell Morphology,Cell-To-Cell Signaling and Interaction,Cellular Assembly and Organization,Nervous System Development and Function,Tissue Morphology** | Morphology of synapse | 0.0144 | FBXO45, GAP43, MAPT, SNCA | 4 |
| **Cell Morphology,Cellular Assembly and Organization,Cellular Development,Cellular Function and Maintenance,Cellular Growth and Proliferation,Embryonic Development,Nervous System Development and Function,Organismal Development,Tissue Development,Tissue Morphology** | Length of dendritic spines | 0.0151 | MAPT, PAK1 | 2 |
| **Neurological Disease,Organismal Injury and Abnormalities** | Progressive encephalopathy | 0.0154 | GAP43, MAPT, SNCA, TUBA1A | 4 |
| **Neurological Disease,Organismal Injury and Abnormalities** | Gliosis of central nervous system | 0.0159 | GSK3B, MAPT, SNCA | 3 |
| **Nervous System Development and Function,Skeletal and Muscular System Development and Function** | Motor function | 0.0166 | MAPT, SNCA | 2 |
| **Neurological Disease** | Cognitive impairment | 0.0166 | LCMT1, MAPT | 2 |
| **Cell Morphology,Cell-To-Cell Signaling and Interaction,Nervous System Development and Function** | Plasticity of neuronal synapse | 0.0166 | GSK3B, SNCA | 2 |
| **Neurological Disease** | Progressive motor neuropathy | 0.0171 | FAS, GAP43, MAPT, SNCA, TUBA1A | 5 |
| **Hematological System Development and Function,Nervous System Development and Function,Tissue Morphology** | Quantity of microglia | 0.0181 | MAPT, SNCA | 2 |
| **Nervous System Development and Function,Organ Morphology,Organismal Development** | Size of forebrain | 0.0183 | CHRD, GAP43, VTI1B | 3 |
| **Cellular Assembly and Organization** | Production of cellular inclusion bodies | 0.0188 | SNCA | 1 |
| **Cell-To-Cell Signaling and Interaction,Nervous System Development and Function** | Long-term potentiation of cortical neurons | 0.0188 | PAK1 | 1 |
| **Inflammatory Disease,Inflammatory Response,Neurological Disease,Organismal Injury and Abnormalities** | Inflammation of substantia nigra | 0.0188 | SNCA | 1 |
| **Cellular Assembly and Organization** | Trafficking of dense-core vesicles | 0.0188 | GSK3B | 1 |
| **Cell Morphology,Cellular Assembly and Organization,Cellular Development,Cellular Function and Maintenance,Cellular Growth and Proliferation,Embryonic Development,Nervous System Development and Function,Organismal Development,Tissue Development** | Branching of sympathetic neurites | 0.0188 | GAP43 | 1 |
| **Hereditary Disorder,Neurological Disease,Organismal Injury and Abnormalities,Psychological Disorders** | Frontotemporal dementia with parkinsonism | 0.0188 | MAPT | 1 |
| **Cell Death and Survival,Neurological Disease,Organismal Injury and Abnormalities** | Apoptosis of somatotrophs | 0.0188 | FAS | 1 |
| **Cell Death and Survival,Cellular Compromise,Nervous System Development and Function** | Cytotoxicity of dopaminergic neurons | 0.0188 | SNCA | 1 |
| **Metabolic Disease,Neurological Disease,Organismal Injury and Abnormalities,Psychological Disorders** | Mild Alzheimer disease | 0.0188 | MAPT | 1 |
| **Cell-To-Cell Signaling and Interaction,Drug Metabolism,Molecular Transport,Small Molecule Biochemistry** | Reuptake of dopamine | 0.0188 | SNCA | 1 |
| **Neurological Disease** | Axonopathy of neurons | 0.0188 | MAPT | 1 |

| | | | | |
|---|---|---|---|---|
| **Neurological Disease,Organismal Injury and Abnormalities** | Atrophy of olivary nucleus | 0.0188 | SNCA | 1 |
| **Cellular Assembly and Organization,Cellular Compromise** | Formation of Lewy bodies | 0.0188 | SNCA | 1 |
| **Neurological Disease,Organismal Injury and Abnormalities,Psychological Disorders** | Corticobasal degeneration | 0.0188 | MAPT | 1 |
| **Organismal Functions** | Recovery of mice | 0.0188 | FAS | 1 |
| **Neurological Disease,Organismal Injury and Abnormalities,Psychological Disorders** | Preclinical stage amyotrophic lateral sclerosis | 0.0188 | FAS | 1 |
| **Hematological System Development and Function,Inflammatory Response,Tissue Morphology** | Quantity of monocytes | 0.0188 | IL12RB2 | 1 |
| **Cellular Assembly and Organization,Cellular Function and Maintenance** | Dynamics of microtubules | 0.0188 | MAPT | 1 |
| **Cell Morphology,Cellular Function and Maintenance** | Autophagy of dopaminergic neurons | 0.0188 | SNCA | 1 |
| **Cell Death and Survival,Neurological Disease,Organismal Injury and Abnormalities** | Apoptosis of lactotropes | 0.0188 | FAS | 1 |
| **Cell-To-Cell Signaling and Interaction,Nervous System Development and Function** | Synaptic transmission of mossy fibers | 0.0188 | MAPT | 1 |
| **Cellular Assembly and Organization,Cellular Compromise,Cellular Function and Maintenance** | Severing of microtubules | 0.0188 | MAPT | 1 |
| **Cell Death and Survival,Cellular Compromise,Neurological Disease,Tissue Morphology** | Degeneration of myelin figure | 0.0188 | SNCA | 1 |
| **Dermatological Diseases and Conditions,Infectious Diseases,Organismal Injury and Abnormalities** | Leprosy | 0.0188 | MAPT | 1 |
| **Embryonic Development,Nervous System Development and Function,Organ Development,Organismal Development,Tissue Development** | Neurogenesis of dentate gyrus | 0.0198 | MAPT, SNCA | 2 |
| **Cellular Development,Cellular Growth and Proliferation,Nervous System Development and Function,Tissue Development** | Growth of neurites | 0.0209 | ARHGEF25, FAS, GAP43, GSK3B, MAPT, PAK1, RNF6, RTN1, TUBA1A, VTI1B | 10 |
| **Cellular Development** | Differentiation of cerebral cortex cells | 0.0214 | GSK3B, MAPT | 2 |
| **Neurological Disease,Organismal Injury and Abnormalities** | Astrocytosis of brain | 0.0214 | GSK3B, MAPT | 2 |
| **Cellular Development,Cellular Growth and Proliferation,Nervous System Development and Function,Tissue Development** | Outgrowth of axons | 0.0223 | GAP43, MAPT, RNF6, VTI1B | 4 |
| **Cellular Development,Cellular Growth and Proliferation,Nervous System Development and Function,Tissue Development** | Proliferation of neuronal cells | 0.0232 | ACSL6, ARHGEF25, FAS, GAP43, GSK3B, MAPT, PAK1, RNF6, RTN1, TUBA1A, VTI1B | 11 |
| **Nervous System Development and Function,Neurological Disease,Organ Morphology,Organismal Development,Organismal Injury and Abnormalities** | Abnormal morphology of striatum | 0.0249 | SNCA, VTI1B | 2 |
| **Gene Expression** | Expression of RNA | 0.0249 | GSK3B, THRB | 2 |
| **Cell Morphology,Cellular Assembly and Organization,Cellular Function and Maintenance,Nervous System Development and Function** | Extension of axons | 0.0264 | GSK3B, MAPT, PAK1, RNF6 | 4 |
| **Hematological System Development and Function,Inflammatory Response,Tissue Morphology** | Quantity of macrophages | 0.0267 | IL12RB2, SNCA | 2 |
| **Cell Morphology,Cellular Assembly and Organization,Cellular Development,Cellular Function and Maintenance,Cellular Growth and Proliferation,Embryonic Development,Nervous** | Dendritic growth/branching | 0.027 | ARHGEF25, GAP43, GSK3B, MAGI2, | 7 |

| | | | | |
|---|---|---|---|---|
| **System Development and Function,Organismal Development,Tissue Development** | | | MAPT, PAK1, SNCA | |
| **Developmental Disorder,Neurological Disease,Organismal Injury and Abnormalities** | Dystrophy of axons | 0.0281 | SNCA | 1 |
| **Cellular Assembly and Organization,Cellular Compromise,Cellular Function and Maintenance** | Disruption of microtubules | 0.0281 | MAPT | 1 |
| **Cellular Assembly and Organization,Cellular Function and Maintenance** | Bundling of microtubules | 0.0281 | MAPT | 1 |
| **Cell Morphology,Cellular Assembly and Organization** | Elongation of mitochondria | 0.0281 | MAPT | 1 |
| **Cancer,Organismal Injury and Abnormalities** | Transformation of Schwann cells | 0.0281 | PAK1 | 1 |
| **Cell-To-Cell Signaling and Interaction,Inflammatory Response,Nervous System Development and Function** | Cytotoxic reaction of neurons | 0.0281 | MAPT | 1 |
| **Nervous System Development and Function,Neurological Disease,Organ Morphology,Organismal Development,Organismal Injury and Abnormalities** | Lack of internal capsule | 0.0281 | FBXO45 | 1 |
| **Auditory and Vestibular System Development and Function** | Electromotility of outer hair cells | 0.0281 | THRB | 1 |
| **Metabolic Disease,Neurological Disease,Organismal Injury and Abnormalities,Psychological Disorders** | Severe Alzheimer disease | 0.0281 | MAPT | 1 |
| **Cell Morphology,Nervous System Development and Function,Organ Morphology,Organismal Development,Tissue Morphology** | Morphology of cortical projection neurons | 0.0281 | TUBA1A | 1 |
| **Cell Morphology,Cellular Function and Maintenance** | Macroautophagy | 0.0281 | SNCA | 1 |
| **Cell Death and Survival,Cellular Compromise,Neurological Disease,Organismal Injury and Abnormalities,Psychological Disorders,Tissue Morphology** | Neurodegeneration of striatonigral neurons | 0.0281 | SNCA | 1 |
| **Nervous System Development and Function,Neurological Disease** | Abnormal morphology of sensory ganglion | 0.0281 | VTI1B | 1 |
| **Nervous System Development and Function** | Remyelination of sciatic nerve | 0.0281 | GSK3B | 1 |
| **Metabolic Disease,Neurological Disease,Organismal Injury and Abnormalities,Psychological Disorders** | Moderate Alzheimer disease | 0.0281 | MAPT | 1 |
| **Neurological Disease,Organismal Injury and Abnormalities** | Neurological disorder of subiculum | 0.0281 | MAPT | 1 |
| **Neurological Disease,Organismal Injury and Abnormalities,Psychological Disorders** | Tauopathy of amygdala | 0.0281 | MAPT | 1 |
| **Nervous System Development and Function,Neurological Disease,Tissue Morphology** | Abnormal morphology of gray matter | 0.0281 | FBXO45 | 1 |
| **Nervous System Development and Function,Organ Morphology,Tissue Morphology** | Quantity of CA1 neuron | 0.0281 | MAPT | 1 |
| **Developmental Disorder,Neurological Disease** | Hypoplasia of vestibular ganglion | 0.0281 | VTI1B | 1 |
| **Cellular Compromise** | Degeneration of mitochondria | 0.0281 | MAPT | 1 |
| **Embryonic Development,Nervous System Development and Function,Organ Development,Organismal Development,Tissue Development** | Formation of optic tract | 0.0281 | GAP43 | 1 |
| **Cellular Assembly and Organization,Cellular Function and Maintenance** | Density of microtubules | 0.0281 | MAPT | 1 |
| **Cellular Compromise,Neurological Disease,Organismal Injury and Abnormalities** | Lesioning of neurons | 0.0281 | MAPT | 1 |
| **Cell Death and Survival,Neurological Disease,Organismal Injury and Abnormalities** | Apoptosis of hippocampal neurons | 0.0286 | FAS, GSK3B | 2 |
| **Cell Death and Survival** | Cell death of motor neurons | 0.0297 | FAS, MAPT, SNCA | 3 |
| **Embryonic Development,Nervous System Development and Function,Organ Development,Organismal Development,Tissue Development** | Development of cerebral cortex | 0.0305 | GAP43, GSK3B, MAPT, PAK1, SNCA | 5 |

| | | | | |
|---|---|---|---|---|
| **Neurological Disease,Organismal Injury and Abnormalities,Psychological Disorders** | Frontotemporal lobar degeneration or amyotrophic lateral sclerosis | 0.0306 | FAS, MAPT | 2 |
| **Free Radical Scavenging** | Production of reactive oxygen species | 0.0306 | COX8A, MAPT | 2 |
| **Cell-To-Cell Signaling and Interaction,Nervous System Development and Function** | Long-term potentiation of hippocampus | 0.0324 | CHRD, GSK3B, MAPT, SNCA | 4 |
| **Nervous System Development and Function** | Myelination of nerves | 0.0325 | GSK3B, THRB | 2 |
| **Nervous System Development and Function** | Development of nerves | 0.0353 | GSK3B, THRB, TMEM126A | 3 |
| **Cell-To-Cell Signaling and Interaction,Molecular Transport,Small Molecule Biochemistry** | Concentration of 5-hydroxytryptamine | 0.0367 | GAP43, MAPT | 2 |
| **Cell Death and Survival,Cellular Compromise,Neurological Disease,Organismal Injury and Abnormalities,Tissue Morphology** | Neurodegeneration of CA3 neurons | 0.0373 | MAPT | 1 |
| **Cellular Assembly and Organization,Nervous System Development and Function,Tissue Morphology** | Quantity of perikaryon | 0.0373 | SNCA | 1 |
| **Neurological Disease,Organismal Injury and Abnormalities** | Neurodegeneration of dentate gyrus | 0.0373 | MAPT | 1 |
| **Cell Morphology,Cellular Assembly and Organization** | Mass of mitochondria | 0.0373 | SNCA | 1 |
| **Embryonic Development,Nervous System Development and Function,Organ Development,Organismal Development,Tissue Development** | Formation of hippocampal commissure | 0.0373 | GAP43 | 1 |
| **Cellular Function and Maintenance** | Movement of centrosome | 0.0373 | GSK3B | 1 |
| **Cellular Assembly and Organization,Nervous System Development and Function** | Replenishment of synaptic vesicles | 0.0373 | SNCA | 1 |
| **Neurological Disease** | Degeneration of dorsal root ganglion | 0.0373 | VTI1B | 1 |
| **Cell Death and Survival,Cellular Compromise,Neurological Disease,Tissue Morphology** | Degeneration of perikaryon | 0.0373 | MAPT | 1 |
| **Cellular Assembly and Organization,Cellular Function and Maintenance,Cellular Movement,Nervous System Development and Function** | Fast axonal transport | 0.0373 | MAPT | 1 |
| **Cellular Compromise,Neurological Disease,Organismal Injury and Abnormalities** | Atrophy of dendrites | 0.0373 | MAPT | 1 |
| **Cell Death and Survival,Cellular Compromise,Neurological Disease,Tissue Morphology** | Degeneration of dendrites | 0.0373 | MAPT | 1 |
| **Neurological Disease,Organismal Injury and Abnormalities** | Degeneration of substantia nigra | 0.0373 | SNCA | 1 |
| **Cell Death and Survival,Nervous System Development and Function** | Cell survival of dopaminergic neurons | 0.0373 | SNCA | 1 |
| **Cell Morphology,Cellular Assembly and Organization,Nervous System Development and Function,Tissue Morphology** | Size of postsynaptic density | 0.0373 | MAPT | 1 |
| **Nervous System Development and Function,Neurological Disease,Organ Morphology,Organismal Development,Organismal Injury and Abnormalities** | Abnormal morphology of disorganized barrel cortex | 0.0373 | GAP43 | 1 |
| **Lipid Metabolism,Molecular Transport,Small Molecule Biochemistry** | Concentration of docosahexaenoic acid | 0.0373 | SNCA | 1 |
| **Cellular Assembly and Organization** | Quantity of granules | 0.0373 | SNCA | 1 |
| **Cellular Assembly and Organization,Neurological Disease,Organismal Development,Tissue Development** | Formation of senile plaques | 0.0373 | MAPT | 1 |
| **Cellular Compromise** | Permeability transition of mitochondria | 0.0373 | COX4I1 | 1 |
| **Neurological Disease,Organismal Injury and Abnormalities** | Brain atrophy | 0.041 | MAPT, SNCA | 2 |

| | | | | |
|---|---|---|---|---|
| **Hematological System Development and Function,Tissue Morphology** | Quantity of mononuclear leukocytes | 0.041 | IL12RB2, SNCA | 2 |
| **Cell-To-Cell Signaling and Interaction,Cellular Assembly and Organization,Cellular Function and Maintenance,Tissue Development** | Organization of synapse | 0.0427 | MAPT, SNCA, TUBA1A | 3 |
| **Cellular Assembly and Organization,Nervous System Development and Function** | Fasciculation of axons | 0.0432 | FBXO45, GAP43 | 2 |
| **Cell Death and Survival,Neurological Disease,Organismal Injury and Abnormalities** | Cell death of cerebellar granule cell | 0.0432 | GSK3B, MAPT | 2 |
| **Embryonic Development,Nervous System Development and Function,Organ Development,Organismal Development,Tissue Development** | Formation of forebrain | 0.0438 | CHRD, FBXO45, GAP43, GSK3B, MRTFB, PAK1 | 6 |
| **Cell Death and Survival,Neurological Disease,Organismal Injury and Abnormalities** | Cell death of granule cells | 0.044 | FAS, GSK3B, MAPT | 3 |
| **Cell Morphology,Cell-To-Cell Signaling and Interaction,Nervous System Development and Function** | Plasticity of synapse | 0.0448 | GSK3B, MAPT, MEF2C, SNCA | 4 |
| **Cellular Assembly and Organization,Nervous System Development and Function,Tissue Morphology** | Quantity of presynaptic terminals | 0.0465 | SNCA | 1 |
| **Cell Morphology,Cell-To-Cell Signaling and Interaction,Cellular Assembly and Organization,Nervous System Development and Function,Tissue Morphology** | Size of excitatory synapses | 0.0465 | SNCA | 1 |
| **Endocrine System Development and Function,Nervous System Development and Function,Organ Morphology,Tissue Morphology** | Quantity of thyrotrophs | 0.0465 | THRB | 1 |
| **Cell Morphology,Cellular Function and Maintenance** | Autophagy of mitochondria | 0.0465 | SNCA | 1 |
| **Cell-To-Cell Signaling and Interaction,Nervous System Development and Function** | Compound action potential of cochlear nerve | 0.0465 | THRB | 1 |
| **Cardiovascular Disease,Neurological Disease,Organismal Injury and Abnormalities** | Infarction of cerebral cortex | 0.0465 | SNCA | 1 |
| **Cell Death and Survival,Cell-To-Cell Signaling and Interaction,Cellular Compromise,Neurological Disease,Tissue Morphology** | Degeneration of synapse | 0.0465 | MAPT | 1 |
| **Nervous System Development and Function,Tissue Development** | Development of fiber tract | 0.0465 | FBXO45 | 1 |
| **Cell Morphology,Nervous System Development and Function** | Innervation of optic nerve | 0.0465 | GAP43 | 1 |
| **Endocrine System Development and Function,Organ Development** | Function of pituitary gland | 0.0465 | THRB | 1 |
| **Cellular Assembly and Organization,Cellular Function and Maintenance** | Quantity of plasma membrane projections | 0.0465 | MAPT | 1 |
| **Cell-To-Cell Signaling and Interaction,Cellular Growth and Proliferation,Nervous System Development and Function** | Excitation of hippocampal neurons | 0.0465 | SNCA | 1 |
| **Embryonic Development,Nervous System Development and Function,Organ Development,Organ Morphology,Organismal Development,Tissue Development** | Morphogenesis of anterior commissure | 0.0465 | FBXO45 | 1 |
| **Cell-To-Cell Signaling and Interaction,Nervous System Development and Function** | Long-term potentiation | 0.0486 | CHRD, GSK3B, LCMT1, MAPT, PAK1, SNCA | 6 |
| **Cell Death and Survival,Neurological Disease,Organismal Injury and Abnormalities** | Cell death of hippocampal neurons | 0.0494 | FAS, GSK3B, SNCA | 3 |

### Supplementary Table 4: Module 4
### Cluster 10

| Categories | Diseases or Functions Annotation | p-Value | Molecules | # Molecules |
|---|---|---|---|---|

| | | | | |
|---|---|---|---|---|
| **Cell-To-Cell Signaling and Interaction,Nervous System Development and Function** | Long-term potentiation of hippocampal CA1 region | 0.000241 | MGLL, PTPN5, RAPGEF3, SYNPO | 4 |
| **Cell-To-Cell Signaling and Interaction,Nervous System Development and Function** | Long-term potentiation of CA1 neuron | 0.00531 | PTPN5, RAPGEF3 | 2 |
| **Cellular Development,Cellular Growth and Proliferation,Embryonic Development,Nervous System Development and Function,Organ Development,Organismal Development,Tissue Development** | Generation of hippocampal neurons | 0.00612 | RAPGEF3 | 1 |
| **Cell-To-Cell Signaling and Interaction,Lipid Metabolism,Molecular Transport,Small Molecule Biochemistry** | Accumulation of 2-arachidonoylglycerol | 0.00612 | MGLL | 1 |
| **Lipid Metabolism,Molecular Transport,Small Molecule Biochemistry** | Quantity of 2-oleoylglycerol | 0.00612 | MGLL | 1 |
| **Cellular Compromise,Neurological Disease,Organismal Injury and Abnormalities,Psychological Disorders** | Injury of corticostriatal neurons | 0.00612 | PTPN5 | 1 |
| **Cell Morphology,Cellular Assembly and Organization,Cellular Development,Cellular Function and Maintenance,Cellular Growth and Proliferation,Nervous System Development and Function,Organismal Development,Tissue Development** | Development of corticothalamic axons | 0.00612 | TTL | 1 |
| **Cell-To-Cell Signaling and Interaction,Cellular Growth and Proliferation,Nervous System Development and Function** | Depolarization-induced suppression of inhibition of pyramidal neurons | 0.00612 | MGLL | 1 |
| **Cell Death and Survival,Neurological Disease,Organismal Injury and Abnormalities** | Apoptosis of lactotropes | 0.0122 | FAS | 1 |
| **Organismal Functions** | Recovery of mice | 0.0122 | FAS | 1 |
| **Neurological Disease,Organismal Injury and Abnormalities,Psychological Disorders** | Preclinical stage amyotrophic lateral sclerosis | 0.0122 | FAS | 1 |
| **Cell-To-Cell Signaling and Interaction,Nervous System Development and Function** | Endocannabinoid-dependent long-term depression of synapse | 0.0122 | MGLL | 1 |
| **Cell-To-Cell Signaling and Interaction,Cellular Assembly and Organization,Nervous System Development and Function** | Induction of axons | 0.0122 | RAPGEF3 | 1 |
| **Cell Death and Survival,Neurological Disease,Organismal Injury and Abnormalities** | Apoptosis of somatotrophs | 0.0122 | FAS | 1 |
| **Nervous System Development and Function,Neurological Disease** | Abnormal morphology of sensory ganglion | 0.0183 | VTI1B | 1 |
| **Cellular Movement,Embryonic Development,Nervous System Development and Function** | Nucleokinesis of neuroblasts | 0.0183 | MGLL | 1 |
| **Developmental Disorder,Neurological Disease** | Hypoplasia of vestibular ganglion | 0.0183 | VTI1B | 1 |
| **Nervous System Development and Function,Neurological Disease,Organ Morphology,Organismal Development,Organismal Injury and Abnormalities** | Lack of internal capsule | 0.0183 | TTL | 1 |
| **Neurological Disease,Organismal Injury and Abnormalities,Psychological Disorders** | Multiple system atrophy | 0.0183 | FAS | 1 |
| **Cell Death and Survival,Neurological Disease,Organismal Injury and Abnormalities** | Apoptosis of cortical neurons | 0.0216 | FAS, RAPGEF3, ZNRF1 | 3 |
| **Neurological Disease** | Degeneration of dorsal root ganglion | 0.0243 | VTI1B | 1 |
| **Cell-To-Cell Signaling and Interaction,Nervous System Development and Function** | Long-term potentiation of dentate granule cells | 0.0243 | RAPGEF3 | 1 |
| **Cell Morphology,Nervous System Development and Function,Tissue Morphology** | Volume of neurons | 0.0243 | FGF13 | 1 |
| **Cellular Development,Nervous System Development and Function,Tissue Development** | Differentiation of striatal neurons | 0.0302 | ARPP21 | 1 |
| **DNA Replication, Recombination, and Repair** | Mitochondrial DNA damage | 0.0302 | OGG1 | 1 |

| | | | | |
|---|---|---|---|---|
| **Nervous System Development and Function,Tissue Morphology** | Quantity of neuroepithelial cells | 0.0302 | BMP6 | 1 |
| **Cell Death and Survival,Neurological Disease,Organismal Injury and Abnormalities** | Cell death of cortical neurons | 0.0331 | FAS, PTPN5, RAPGEF3, ZNRF1 | 4 |
| **Nervous System Development and Function,Organ Morphology,Organismal Development** | Morphology of forebrain | 0.0353 | ABI2, DYNC2LI1, NDST3, TTL, VTI1B | 5 |
| **Cellular Assembly and Organization,Cellular Compromise,Cellular Function and Maintenance** | Disassembly of microtubules | 0.0362 | TTL | 1 |
| **Cellular Development,Cellular Growth and Proliferation,Nervous System Development and Function,Tissue Development** | Growth of dendritic spines | 0.0362 | SYNPO | 1 |
| **Endocrine System Disorders,Gastrointestinal Disease,Metabolic Disease,Organismal Injury and Abnormalities** | Diabetes mellitus | 0.0362 | FAS | 1 |
| **Free Radical Scavenging,Molecular Transport** | Accumulation of reactive oxygen species | 0.0362 | MGLL | 1 |
| **Cell Morphology,Cellular Assembly and Organization,Cellular Development,Cellular Function and Maintenance,Cellular Growth and Proliferation,Nervous System Development and Function,Organismal Development,Tissue Development** | Polarization of hippocampal neurons | 0.0421 | RAPGEF3 | 1 |
| **Neurological Disease** | Degeneration of vestibular ganglion | 0.0421 | VTI1B | 1 |
| **Nutritional Disease,Psychological Disorders** | Anorexia | 0.0421 | CPT1C | 1 |
| **Neurological Disease,Organismal Injury and Abnormalities,Psychological Disorders** | Disorder of basal ganglia | 0.043 | ARPP21, BHLHE40, FAS, FGF13, PTPN5, RPL31 | 6 |
| **Neurological Disease** | Movement Disorders | 0.0464 | ARPP21, BHLHE40, FAS, FGF13, PTPN5, RPL31 | 6 |
| **Neurological Disease,Organismal Injury and Abnormalities** | Abnormality of striatum | 0.0475 | PTPN5, VTI1B | 2 |
| **Cardiovascular System Development and Function,Cell-To-Cell Signaling and Interaction** | Adhesion of endothelial cells | 0.048 | FAS | 1 |
| **Cell Death and Survival,Embryonic Development** | Cell death of neuroblasts | 0.048 | FAS | 1 |
| **Nervous System Development and Function,Neurological Disease,Organ Morphology,Organismal Development,Organismal Injury and Abnormalities** | Lack of hippocampal commissure | 0.048 | VTI1B | 1 |
| **Endocrine System Development and Function,Lipid Metabolism,Molecular Transport,Small Molecule Biochemistry** | Quantity of 2-arachidonoylglycerol | 0.048 | MGLL | 1 |
| **Cellular Movement,Hematological System Development and Function,Immune Cell Trafficking** | Cell rolling of leukocytes | 0.048 | FAS | 1 |

### Cluster 11

| Categories | Diseases or Functions Annotation | p-Value | Molecules | # Molecules |
|---|---|---|---|---|
| **Neurological Disease,Skeletal and Muscular Disorders** | Progressive motor dysfunction | 0.00629 | TARDBP | 1 |
| **Neurological Disease,Organismal Injury and Abnormalities** | Progressive neurodegenerative disorder | 0.00629 | TARDBP | 1 |
| **Cellular Development,Cellular Growth and Proliferation,Nervous System Development and Function,Organ Development,Tissue Development** | Delay in proliferation of granule cells | 0.00629 | NR1D1 | 1 |
| **Cell Morphology,Nervous System Development and Function,Tissue Morphology** | Size of motor axons | 0.00629 | TARDBP | 1 |

| | | | | |
|---|---|---|---|---|
| **Cellular Assembly and Organization** | Formation of mitochondrial cristae | 0.00629 | TARDBP | 1 |
| **Neurological Disease,Skeletal and Muscular Disorders** | Progressive paralysis | 0.00629 | TARDBP | 1 |
| **Cellular Movement,Nervous System Development and Function** | Delay in migration of granule cells | 0.00629 | NR1D1 | 1 |
| **Cellular Assembly and Organization,Cellular Function and Maintenance** | Movement of recycling endosomes | 0.00629 | TARDBP | 1 |
| **Neurological Disease** | Cognitive impairment | 0.00758 | PPME1, TARDBP | 2 |
| **Cardiovascular Disease,Hematological Disease,Metabolic Disease** | Hyperlipidemia | 0.0125 | SH2B1 | 1 |
| **Endocrine System Disorders** | Leptin resistance | 0.0125 | SH2B1 | 1 |
| **Cell Death and Survival,Nervous System Development and Function** | Cell survival of cortical neurons | 0.0125 | TARDBP | 1 |
| **Cellular Assembly and Organization,DNA Replication, Recombination, and Repair** | Formation of nuclear foci | 0.0125 | TARDBP | 1 |
| **Metabolic Disease** | Hyperleptinemia | 0.0125 | SH2B1 | 1 |
| **Cardiovascular Disease,Cardiovascular System Development and Function,Organismal Injury and Abnormalities** | Abnormal morphology of vasculature | 0.0142 | CC2D1A, NDP | 2 |
| **Organismal Injury and Abnormalities,Skeletal and Muscular Disorders** | Atrophy of skeletal muscle | 0.0187 | TARDBP | 1 |
| **Nutritional Disease** | Obesity | 0.0249 | PTPN2, SH2B1 | 2 |
| **Cellular Assembly and Organization** | Aggregation of mitochondria | 0.0249 | TARDBP | 1 |
| **Cellular Assembly and Organization** | Formation of cytoplasmic aggregates | 0.0249 | TARDBP | 1 |
| **Endocrine System Disorders,Hematological Disease,Metabolic Disease** | Hyperglycemia | 0.0249 | SH2B1 | 1 |
| **Endocrine System Disorders,Metabolic Disease** | Impaired glucose tolerance | 0.031 | SH2B1 | 1 |
| **Cellular Compromise,Nervous System Development and Function,Neurological Disease** | Degeneration of myelin sheath | 0.0371 | TARDBP | 1 |
| **Nervous System Development and Function,Neurological Disease,Organ Morphology,Organismal Development,Organismal Injury and Abnormalities** | Abnormal morphology of dorsal column of spinal cord | 0.0371 | NFATC3 | 1 |
| **Cell Death and Survival** | Cell death of striatal neurons | 0.0408 | DNAJB2, SDHA | 2 |
| **Neurological Disease** | Memory deficits | 0.0432 | TARDBP | 1 |
| **Neurological Disease** | Neurological signs | 0.0469 | ANKRD11, DNAJB2, NR1D1, TARDBP, ZNF239 | 5 |
| **Cellular Development,Cellular Growth and Proliferation,Nervous System Development and Function,Tissue Development** | Outgrowth of sensory axons | 0.0492 | NFATC3 | 1 |
| **Endocrine System Disorders,Metabolic Disease** | Insulin resistance | 0.0492 | SH2B1 | 1 |
| **Cellular Development,Cellular Growth and Proliferation,Embryonic Development,Nervous System Development and Function,Organ Development,Organismal Development,Tissue Development** | Development of Purkinje cells | 0.0492 | NR1D1 | 1 |
| **Neurological Disease,Organ Morphology,Organismal Injury and Abnormalities** | Atrophy of hippocampus | 0.0492 | TARDBP | 1 |
| **Neurological Disease,Organismal Injury and Abnormalities** | Reactive gliosis | 0.0492 | TARDBP | 1 |
| **Cellular Assembly and Organization** | Accumulation of mitochondria | 0.0492 | TARDBP | 1 |

**Supplementary Table 5: Module 5**
**Cluster 12**

| Categories | Diseases or Functions Annotation | p-Value | Molecules | # Molecules |
|---|---|---|---|---|
| **Nervous System Development and Function,Neurological Disease,Organ Morphology,Organismal Development,Organismal Injury and Abnormalities** | Abnormal morphology of optic tract | 0.00247 | GAP43, VTI1B | 2 |
| **Cell Morphology,Cellular Assembly and Organization,Cellular Development,Cellular Function and Maintenance,Cellular Growth and Proliferation,Embryonic Development,Nervous System Development and Function,Organismal Development,Tissue Development** | Morphogenesis of dendritic spines | 0.00373 | OPTN, RAC1 | 2 |
| **Embryonic Development,Nervous System Development and Function,Organ Development,Organismal Development,Tissue Development** | Formation of anterior commissure | 0.00446 | GAP43, RAC1 | 2 |
| **Endocrine System Development and Function,Lipid Metabolism,Molecular Transport,Small Molecule Biochemistry** | Concentration of corticosterone | 0.00446 | CRH, FAS | 2 |
| **Nervous System Development and Function** | Neuroprotection of cerebellar granule cell | 0.00849 | CRH | 1 |
| **Neurological Disease** | Axonopathy of central nervous system | 0.00849 | STMN1 | 1 |
| **Cellular Compromise,Neurological Disease,Organismal Injury and Abnormalities,Psychological Disorders** | Injury of corticostriatal neurons | 0.00849 | PTPN5 | 1 |
| **Cellular Assembly and Organization,Cellular Function and Maintenance** | Endocytosis of plasma membrane | 0.00849 | RAC1 | 1 |
| **Nervous System Development and Function,Tissue Morphology** | Density of serotonergic neurons | 0.00849 | GAP43 | 1 |
| **Cellular Movement,Nervous System Development and Function** | Delay in initiation of migration of cortical projection neurons | 0.00849 | RAC1 | 1 |
| **Embryonic Development,Nervous System Development and Function,Organ Development,Organismal Development,Tissue Development** | Development of layer IV | 0.00849 | GAP43 | 1 |
| **Cellular Assembly and Organization,Cellular Function and Maintenance** | Quantity of peripheral lamellae | 0.00849 | RAC1 | 1 |
| **Neurological Disease** | Axonopathy of peripheral nervous system | 0.00849 | STMN1 | 1 |
| **Nervous System Development and Function,Neurological Disease,Organ Morphology,Organismal Development,Organismal Injury and Abnormalities** | Lack of optic tract | 0.00849 | GAP43 | 1 |
| **Nervous System Development and Function,Organ Morphology,Organismal Development** | Size of optic chiasm | 0.00849 | GAP43 | 1 |
| **Cellular Function and Maintenance,Nervous System Development and Function** | Regulation of noradrenergic neurons | 0.00849 | CRH | 1 |
| **Cell Morphology,Cellular Movement,Nervous System Development and Function** | Innervation of serotonergic neurons | 0.00849 | GAP43 | 1 |
| **Cell-To-Cell Signaling and Interaction,Nervous System Development and Function** | Activation of projection neurons | 0.00849 | CRH | 1 |
| **Cell Morphology,Cellular Assembly and Organization** | Elongation of astrocyte projections | 0.00849 | GAP43 | 1 |
| **Neurological Disease** | Formation of myelin ovoid | 0.00849 | RAC1 | 1 |
| **Hereditary Disorder,Neurological Disease,Organismal Injury and Abnormalities,Psychological Disorders,Skeletal and Muscular Disorders** | Parkinson-dementia syndrome | 0.00849 | GAP43 | 1 |

| | | | | |
|---|---|---|---|---|
| **Cell Morphology,Cellular Assembly and Organization,Nervous System Development and Function** | Projection of corticothalamic axons | 0.00849 | RAC1 | 1 |
| **Cell-To-Cell Signaling and Interaction,Drug Metabolism,Molecular Transport,Small Molecule Biochemistry** | Concentration of norepinephrine | 0.0148 | CRH, GAP43 | 2 |
| **Endocrine System Development and Function,Organ Development** | Activation of pituitary gland | 0.0169 | CRH | 1 |
| **Cell Death and Survival,Neurological Disease,Organismal Injury and Abnormalities** | Apoptosis of lactotropes | 0.0169 | FAS | 1 |
| **Organismal Functions** | Recovery of mice | 0.0169 | FAS | 1 |
| **Nervous System Development and Function,Organ Morphology,Tissue Morphology** | Quantity of mossy fiber cells | 0.0169 | RAC1 | 1 |
| **Behavior** | Despair behavior | 0.0169 | CRH | 1 |
| **Cell-To-Cell Signaling and Interaction,Drug Metabolism,Small Molecule Biochemistry** | Depletion of norepinephrine | 0.0169 | CRH | 1 |
| **Cell Morphology,Cellular Assembly and Organization,Cellular Development,Cellular Function and Maintenance,Cellular Growth and Proliferation,Embryonic Development,Nervous System Development and Function,Organismal Development,Tissue Development** | Branching of sympathetic neurites | 0.0169 | GAP43 | 1 |
| **Cell Cycle,Endocrine System Development and Function,Nervous System Development and Function** | Mitogenesis of corticotroph cells | 0.0169 | CRH | 1 |
| **Cellular Movement,Nervous System Development and Function** | Distribution of interneurons | 0.0169 | DCK | 1 |
| **Cell-To-Cell Signaling and Interaction,Cellular Growth and Proliferation,Nervous System Development and Function** | Excitation of type II spiral ganglion neurons | 0.0169 | CRH | 1 |
| **Cell Death and Survival,Neurological Disease,Organismal Injury and Abnormalities** | Apoptosis of somatotrophs | 0.0169 | FAS | 1 |
| **Cellular Assembly and Organization,Nervous System Development and Function** | Delay in sorting of axons | 0.0169 | RAC1 | 1 |
| **Cell Morphology,Cellular Assembly and Organization,Nervous System Development and Function** | Extension of leading processes | 0.0169 | RAC1 | 1 |
| **Neurological Disease,Organismal Injury and Abnormalities,Psychological Disorders** | Preclinical stage amyotrophic lateral sclerosis | 0.0169 | FAS | 1 |
| **Cellular Development,Cellular Growth and Proliferation,Embryonic Development,Nervous System Development and Function,Organ Development,Organismal Development,Tissue Development** | Expansion of neural precursor cells | 0.0169 | AHR | 1 |
| **Cellular Assembly and Organization,Cellular Function and Maintenance** | Depolarization of plasma membrane | 0.0169 | CRH | 1 |
| **Cellular Development,Cellular Growth and Proliferation,Nervous System Development and Function,Organ Development,Tissue Development** | Proliferation of hypothalamic neurons | 0.0169 | NEGR1 | 1 |
| **Cell-To-Cell Signaling and Interaction,Nervous System Development and Function** | Inhibitory postsynaptic potential of GABAergic neurons | 0.0169 | CRH | 1 |
| **Cell Signaling,Molecular Transport,Nucleic Acid Metabolism,Small Molecule Biochemistry** | Concentration of cyclic AMP | 0.0218 | CRH, RAC1 | 2 |
| **Nervous System Development and Function,Neurological Disease** | Abnormal morphology of sensory ganglion | 0.0253 | VTI1B | 1 |
| **Cell-To-Cell Signaling and Interaction,Cellular Growth and Proliferation,Nervous System Development and Function** | Excitation of CA3 neurons | 0.0253 | CRH | 1 |
| **Embryonic Development,Nervous System Development and Function,Organ Development,Organismal Development,Tissue Development** | Formation of optic tract | 0.0253 | GAP43 | 1 |
| **Cell-To-Cell Signaling and Interaction,Nervous System Development and Function** | Activation of myenteric ganglion cells | 0.0253 | CRH | 1 |
| **Developmental Disorder,Neurological Disease** | Hypoplasia of vestibular ganglion | 0.0253 | VTI1B | 1 |

| | | | | |
|---|---|---|---|---|
| **Cell Morphology,Cellular Assembly and Organization,Nervous System Development and Function** | Projection of thalamocortical axons | 0.0253 | RAC1 | 1 |
| **Neurological Disease,Organismal Injury and Abnormalities,Psychological Disorders** | Multiple system atrophy | 0.0253 | FAS | 1 |
| **Cell Morphology,Cellular Assembly and Organization,Cellular Function and Maintenance,Nervous System Development and Function** | Formation of astrocyte projections | 0.0253 | RAC1 | 1 |
| **Cardiovascular Disease,Neurological Disease,Organismal Injury and Abnormalities,Psychological Disorders** | Ischemic injury of brain | 0.03 | NOX4, PTPN5 | 2 |
| **Neurological Disease,Organismal Injury and Abnormalities** | Seizures | 0.03 | GAP43, RBFOX1 | 2 |
| **Cell-To-Cell Signaling and Interaction,Nervous System Development and Function** | Firing of Purkinje cells | 0.0335 | RBFOX1 | 1 |
| **Embryonic Development,Nervous System Development and Function,Organ Development,Organismal Development,Tissue Development** | Formation of hippocampal commissure | 0.0335 | GAP43 | 1 |
| **Nervous System Development and Function** | Prepulse inhibition | 0.0335 | CRH | 1 |
| **Cell Morphology,Cellular Assembly and Organization,Cellular Function and Maintenance** | Formation of lamellipodia | 0.0335 | RAC1 | 1 |
| **Neurological Disease** | Degeneration of dorsal root ganglion | 0.0335 | VTI1B | 1 |
| **Behavior** | Place aversion | 0.0335 | CRH | 1 |
| **Nervous System Development and Function,Neurological Disease,Organ Morphology,Organismal Development,Organismal Injury and Abnormalities** | Abnormal morphology of disorganized barrel cortex | 0.0335 | GAP43 | 1 |
| **Lipid Metabolism,Molecular Transport,Small Molecule Biochemistry** | Concentration of triacylglycerol | 0.0335 | CRH | 1 |
| **Cell-To-Cell Signaling and Interaction,Nervous System Development and Function** | Conduction of motor nerve | 0.0335 | STMN1 | 1 |
| **Cell-To-Cell Signaling and Interaction,Drug Metabolism,Molecular Transport,Small Molecule Biochemistry** | Quantity of epinephrine | 0.0335 | CRH | 1 |
| **Free Radical Scavenging** | Synthesis of reactive oxygen species | 0.0392 | NOX4, RAC1 | 2 |
| **Cellular Assembly and Organization,Nervous System Development and Function,Tissue Morphology** | Quantity of axon terminals | 0.0417 | RAC1 | 1 |
| **Cell Morphology,Nervous System Development and Function** | Innervation of optic nerve | 0.0417 | GAP43 | 1 |
| **Neurological Disease** | Progressive motor neuropathy | 0.0455 | FAS, GAP43, PGK1, STMN1 | 4 |
| **Neurological Disease,Organismal Injury and Abnormalities,Psychological Disorders,Skeletal and Muscular Disorders** | Parkinson's disease | 0.0482 | GAP43, PGK1, STMN1 | 3 |
| **Cellular Function and Maintenance** | Endocytosis by neurons | 0.0499 | RAC1 | 1 |
| **Endocrine System Development and Function,Nervous System Development and Function,Organ Morphology,Tissue Morphology** | Quantity of corticotroph cells | 0.0499 | CRH | 1 |
| **Behavior** | Grooming | 0.0499 | CRH | 1 |
| **Endocrine System Disorders,Gastrointestinal Disease,Metabolic Disease,Organismal Injury and Abnormalities** | Diabetes mellitus | 0.0499 | FAS | 1 |

### Supplementary Table 6: Module 6
### Cluster 9

| Categories | Diseases or Functions Annotation | p-Value | Molecules | # Molecules |
|---|---|---|---|---|
| **Amino Acid Metabolism,Small Molecule Biochemistry** | Accumulation of amino acids | 0.000692 | SLC16A2, SLC17A7 | 2 |
| **Embryonic Development,Nervous System Development and Function,Organ Development,Organ Morphology,Organismal Development,Tissue Development** | Morphogenesis of diencephalon | 0.000692 | OTX1, SOX2 | 2 |
| **Behavior** | Place aversion | 0.000692 | NPY, PDYN | 2 |

| | | | | |
|---|---|---|---|---|
| **Neurological Disease,Organismal Injury and Abnormalities** | Degenerative ataxia | 0.00114 | FAS, IGFBP5 | 2 |
| **Cell-To-Cell Signaling and Interaction** | Adhesion of brain cells | 0.00114 | ADGRG1, NECTIN1 | 2 |
| **Amino Acid Metabolism,Endocrine System Development and Function,Molecular Transport,Small Molecule Biochemistry** | Quantity of L-triiodothyronine | 0.00171 | NPY, SLC16A2 | 2 |
| **Cell Morphology,Cellular Development,Connective Tissue Development and Function** | Conversion of induced neural precursor cells | 0.00237 | FOXG1, SOX2 | 2 |
| **Amino Acid Metabolism,Drug Metabolism,Endocrine System Development and Function,Molecular Transport,Small Molecule Biochemistry** | Concentration of levothyroxine | 0.00401 | NPY, SLC16A2 | 2 |
| **Cell-To-Cell Signaling and Interaction,Nervous System Development and Function** | Adhesion of neuronal cells | 0.00977 | ADGRG1, NECTIN1 | 2 |
| **Cell-To-Cell Signaling and Interaction,Nervous System Development and Function** | Activation of KNDy neurons | 0.0109 | PDYN | 1 |
| **Lipid Metabolism,Molecular Transport,Small Molecule Biochemistry** | Accumulation of glucosylceramide | 0.0109 | GBA | 1 |
| **Connective Tissue Development and Function,Organismal Development,Skeletal and Muscular System Development and Function,Tissue Development** | Size of cranium | 0.0109 | MYB | 1 |
| **Amino Acid Metabolism,Molecular Transport,Small Molecule Biochemistry** | Uptake of L-triiodothyronine | 0.0109 | SLC16A2 | 1 |
| **Embryonic Development,Nervous System Development and Function,Organ Development,Organismal Development,Tissue Development** | Formation of subgranular zone of dentate gyrus | 0.0109 | FOXG1 | 1 |
| **Organismal Functions** | Regulation of pituitary gland | 0.0109 | NPY | 1 |
| **Cell-mediated Immune Response,Cell-To-Cell Signaling and Interaction,Cellular Movement,Hematological System Development and Function,Immune Cell Trafficking** | Sequestration of T lymphocytes | 0.0109 | HDC | 1 |
| **Drug Metabolism,Small Molecule Biochemistry** | Accumulation of epinephrine | 0.0109 | PNMT | 1 |
| **Cellular Development,Cellular Growth and Proliferation,Nervous System Development and Function,Tissue Development** | Development of Cajal-Retzius neurons | 0.0109 | FOXG1 | 1 |
| **Lipid Metabolism,Small Molecule Biochemistry** | Incorporation of sphingomyelin | 0.0109 | GBA | 1 |
| **Carbohydrate Metabolism,Lipid Metabolism,Small Molecule Biochemistry** | Incorporation of phosphatidylcholine | 0.0109 | GBA | 1 |
| **Amino Acid Metabolism,Small Molecule Biochemistry** | Accumulation of levothyroxine | 0.0109 | SLC16A2 | 1 |
| **Organismal Functions** | Flicking of tail | 0.0109 | PDYN | 1 |
| **Cellular Development,Cellular Growth and Proliferation,Embryonic Development,Nervous System Development and Function,Organ Development,Organismal Development,Tissue Development** | Expansion of telencephalic progenitor cells | 0.0109 | FOXG1 | 1 |
| **Auditory and Vestibular System Development and Function,Cellular Development,Cellular Growth and Proliferation,Embryonic Development,Nervous System Development and Function,Organ Development,Organismal Development,Tissue Development** | Development of spiral ganglion cells | 0.0109 | SOX2 | 1 |
| **Lipid Metabolism,Molecular Transport,Small Molecule Biochemistry** | Quantity of fat | 0.0109 | NPY | 1 |
| **Lipid Metabolism,Small Molecule Biochemistry** | Storage of fat | 0.0109 | NPY | 1 |
| **Amino Acid Metabolism,Small Molecule Biochemistry** | Accumulation of L-triiodothyronine | 0.0109 | SLC16A2 | 1 |
| **Infectious Diseases** | Infection by human herpesvirus 2 | 0.0109 | NECTIN1 | 1 |
| **Cancer,Organismal Injury and Abnormalities** | Tumor burden | 0.0109 | MERTK | 1 |
| **Lipid Metabolism,Small Molecule Biochemistry** | Incorporation of choline | 0.0109 | GBA | 1 |
| **Metabolic Disease** | Glucose metabolism disorder | 0.016 | FAS, NPY | 2 |
| **Cell Death and Survival,Cellular Development,Cellular Function and Maintenance,Cellular Growth and** | Self-renewal of neural stem cells | 0.0177 | FOXG1, SOX2 | 2 |

| | | | | |
|---|---|---|---|---|
| **Proliferation,Embryonic Development,Organismal Development,Tissue Development** | | | | |
| **Cell Death and Survival,Neurological Disease,Organismal Injury and Abnormalities** | Apoptosis of lactotropes | 0.0216 | FAS | 1 |
| **Cellular Development,Cellular Growth and Proliferation,Embryonic Development,Nervous System Development and Function,Organ Development,Organismal Development,Tissue Development** | Development of spiny stellate neurons | 0.0216 | SLC17A7 | 1 |
| **Organismal Functions** | Recovery of mice | 0.0216 | FAS | 1 |
| **Nervous System Development and Function,Neurological Disease,Organ Morphology,Organismal Development,Organismal Injury and Abnormalities** | Abnormal morphology of oculomotor nucleus | 0.0216 | OTX1 | 1 |
| **Cellular Growth and Proliferation,Nervous System Development and Function** | Inhibition of presympathetic neurons | 0.0216 | NPY | 1 |
| **Behavior** | Inhibitory avoidance behavior | 0.0216 | HDC | 1 |
| **Cell Morphology,Cell-To-Cell Signaling and Interaction,Cellular Assembly and Organization** | Length of gap junctions | 0.0216 | NRCAM | 1 |
| **Nervous System Development and Function,Organ Morphology,Organismal Development** | Thickness of granule cell layer | 0.0216 | NRCAM | 1 |
| **Cell-To-Cell Signaling and Interaction,Cellular Growth and Proliferation,Nervous System Development and Function** | Excitation of type II spiral ganglion neurons | 0.0216 | NPY | 1 |
| **Cell Death and Survival,Neurological Disease,Organismal Injury and Abnormalities** | Apoptosis of somatotrophs | 0.0216 | FAS | 1 |
| **Cell Morphology,Cell-To-Cell Signaling and Interaction,Cellular Assembly and Organization** | Structural integrity of synapse | 0.0216 | SNTA1 | 1 |
| **Hereditary Disorder,Neurological Disease,Organismal Injury and Abnormalities** | Spinocerebellar ataxia type 7 | 0.0216 | IGFBP5 | 1 |
| **Nervous System Development and Function,Neurological Disease,Organ Morphology,Organismal Development,Organismal Injury and Abnormalities** | Abnormal morphology of enlarged cerebellum | 0.0216 | OTX1 | 1 |
| **Cellular Development,Cellular Growth and Proliferation,Nervous System Development and Function,Tissue Development** | Generation of Cajal-Retzius neurons | 0.0216 | FOXG1 | 1 |
| **Behavior,Nervous System Development and Function** | Contextual fear memory retrieval | 0.0216 | NPY | 1 |
| **Cell-To-Cell Signaling and Interaction,Nervous System Development and Function** | Synaptic transmission of thalamocortical neurons | 0.0216 | SLC17A7 | 1 |
| **Embryonic Development,Organismal Development,Tissue Development** | Formation of preplate | 0.0216 | FOXG1 | 1 |
| **Hereditary Disorder,Neurological Disease,Organismal Injury and Abnormalities** | Spinocerebellar ataxia type 1 | 0.0216 | IGFBP5 | 1 |
| **Cardiovascular System Development and Function,Cell Morphology,Cell-To-Cell Signaling and Interaction,Cellular Assembly and Organization,Tissue Morphology** | Electrical resistance of microvascular endothelial cells | 0.0216 | MERTK | 1 |
| **Cell Morphology,Organ Morphology,Organismal Injury and Abnormalities** | Hypertrophy of gonadotropes | 0.0216 | TG | 1 |
| **Neurological Disease,Organismal Injury and Abnormalities,Psychological Disorders** | Preclinical stage amyotrophic lateral sclerosis | 0.0216 | FAS | 1 |
| **Cardiovascular System Development and Function,Nervous System Development and Function,Organ Development** | Integrity of blood-brain barrier | 0.0216 | HDC | 1 |
| **Nervous System Development and Function** | Development of ventral nerve cord | 0.0216 | OTX1 | 1 |
| **Embryonic Development,Nervous System Development and Function,Organ Development,Organismal Development,Tissue Development** | Formation of pretectum | 0.0216 | OTX1 | 1 |
| **Cell-To-Cell Signaling and Interaction,Cellular Assembly and Organization,Cellular Function and Maintenance** | Formation of spot-like adhesion site | 0.0216 | NECTIN1 | 1 |

| | | | | |
|---|---|---|---|---|
| **Embryonic Development,Nervous System Development and Function,Organ Development,Organismal Development,Tissue Development** | Formation of forebrain | 0.0283 | FOXG1, HPRT1, ID2, NUMBL, OTX1, SOX2, TG | 7 |
| **Cell-To-Cell Signaling and Interaction,Nervous System Development and Function** | Long term synaptic depression of perirhinal cortex | 0.0322 | HAPLN1 | 1 |
| **Embryonic Development,Nervous System Development and Function,Organ Development,Organismal Development,Tissue Development** | Formation of dorsal thalamus | 0.0322 | OTX1 | 1 |
| **Cellular Assembly and Organization** | Density of plasma membrane projections | 0.0322 | NUMBL | 1 |
| **Amino Acid Metabolism,Molecular Transport,Small Molecule Biochemistry** | Accumulation of L-glutamic acid | 0.0322 | SLC17A7 | 1 |
| **Cell-To-Cell Signaling and Interaction,Nervous System Development and Function** | Adhesion of hippocampal neurons | 0.0322 | NECTIN1 | 1 |
| **Nervous System Development and Function,Neurological Disease,Organ Morphology,Organismal Development,Organismal Injury and Abnormalities** | Abnormal morphology of red nucleus | 0.0322 | OTX1 | 1 |
| **Molecular Transport** | Internalization of water | 0.0322 | NPY | 1 |
| **Infectious Diseases** | Infection by human herpesvirus 1 | 0.0322 | NECTIN1 | 1 |
| **Cancer,Endocrine System Disorders,Neurological Disease,Organismal Injury and Abnormalities,Reproductive System Disease** | Hyperplasia of thyrotropes | 0.0322 | TG | 1 |
| **Endocrine System Development and Function,Endocrine System Disorders,Nervous System Development and Function,Neurological Disease,Organ Development,Organ Morphology,Organismal Development,Organismal Injury and Abnormalities** | Abnormal morphology of mammillary body | 0.0322 | OTX1 | 1 |
| **Endocrine System Development and Function,Lipid Metabolism,Molecular Transport,Small Molecule Biochemistry** | Secretion of corticosterone | 0.0322 | NPY | 1 |
| **Cell-To-Cell Signaling and Interaction,Nervous System Development and Function** | Adhesion of cerebellar granule cell | 0.0322 | ADGRG1 | 1 |
| **Embryonic Development,Nervous System Development and Function,Organ Development,Organismal Development,Tissue Development** | Formation of Ammon's horn | 0.0322 | OTX1 | 1 |
| **Neurological Disease,Organismal Injury and Abnormalities,Psychological Disorders** | Multiple system atrophy | 0.0322 | FAS | 1 |
| **Embryonic Development,Nervous System Development and Function,Organ Development,Organismal Development,Tissue Development** | Formation of rhombencephalon | 0.037 | ADGRG1, DYNC2LI1, NRCAM, OTX1, SLC17A7 | 5 |
| **Cellular Development,Cellular Growth and Proliferation,Embryonic Development,Nervous System Development and Function,Organismal Development,Tissue Development** | Proliferation of neuroblasts | 0.0376 | FOXG1, NUMBL, SOX2 | 3 |
| **Cell Death and Survival,Nervous System Development and Function** | Cell viability of granule cells | 0.0417 | FOXG1, NRCAM | 2 |
| **Nervous System Development and Function,Neurological Disease,Organ Morphology,Organismal Development,Organismal Injury and Abnormalities** | Abnormal morphology of mesencephalic nucleus | 0.0427 | OTX1 | 1 |
| **Cellular Function and Maintenance,Nervous System Development and Function,Tissue Development** | Function of sympathetic neuron | 0.0427 | MAGED1 | 1 |
| **Nervous System Development and Function,Neurological Disease,Organ Morphology,Organismal** | Abnormal morphology of pretectum | 0.0427 | OTX1 | 1 |

| | | | | |
|---|---|---|---|---|
| **Development,Organismal Injury and Abnormalities** | | | | |
| **Embryonic Development,Nervous System Development and Function,Organ Development,Organismal Development,Tissue Development** | Formation of barrel cortex | 0.0427 | SLC17A7 | 1 |
| **Lipid Metabolism,Molecular Transport,Small Molecule Biochemistry** | Concentration of triacylglycerol | 0.0427 | NPY | 1 |
| **Cell-To-Cell Signaling and Interaction,Drug Metabolism,Molecular Transport,Small Molecule Biochemistry** | Quantity of epinephrine | 0.0427 | PNMT | 1 |
| **Cellular Function and Maintenance** | Regulation of neuroglia | 0.0427 | SOX2 | 1 |
| **Embryonic Development,Nervous System Development and Function,Organ Development,Organismal Development,Tissue Development** | Formation of brain | 0.0459 | ADGRG1, DYNC2LI1, FOXG1, HPRT1, ID2, NRCAM, NUMBL, OTX1, SLC17A7, SOX2 | 12 |
| **Cellular Development,Cellular Growth and Proliferation,Embryonic Development,Nervous System Development and Function,Organ Development,Organismal Development,Tissue Development** | Proliferation of neural precursor cells | 0.0464 | FOXG1, ID2, SOX2 | 3 |

### Cluster 13

| Categories | Diseases or Functions Annotation | p-Value | Molecules | # Molecules |
|---|---|---|---|---|
| **Cellular Movement,Nervous System Development and Function** | Migration of interneurons | 0.00412 | DLX2, RAC1 | 2 |
| **Cellular Development** | Commitment of cortical precursor cells | 0.00751 | DLX2 | 1 |
| **Carbohydrate Metabolism,Lipid Metabolism,Small Molecule Biochemistry** | Hydrolysis of phosphatidylinositol polyphosphate | 0.00751 | EDN1 | 1 |
| **Lipid Metabolism,Molecular Transport,Small Molecule Biochemistry** | Accumulation of glucosylceramide | 0.00751 | GBA | 1 |
| **Cellular Assembly and Organization,Cellular Function and Maintenance** | Endocytosis of plasma membrane | 0.00751 | RAC1 | 1 |
| **Cellular Growth and Proliferation,Nervous System Development and Function,Organ Development** | Proliferation of stellate cells | 0.00751 | EDN1 | 1 |
| **Cellular Movement,Nervous System Development and Function** | Delay in initiation of migration of cortical projection neurons | 0.00751 | RAC1 | 1 |
| **Cellular Assembly and Organization,Cellular Function and Maintenance** | Quantity of peripheral lamellae | 0.00751 | RAC1 | 1 |
| **Cellular Assembly and Organization,Cellular Development,Embryonic Development,Nervous System Development and Function,Organ Development,Organismal Development,Tissue Development** | Maturation of olfactory glomeruli | 0.00751 | PPP3CA | 1 |
| **Connective Tissue Disorders,Inflammatory Disease,Inflammatory Response,Organismal Injury and Abnormalities,Skeletal and Muscular Disorders** | Degeneration of lumbar intervertebral disc | 0.00751 | EDN1 | 1 |
| **Lipid Metabolism,Small Molecule Biochemistry** | Incorporation of sphingomyelin | 0.00751 | GBA | 1 |
| **Carbohydrate Metabolism,Lipid Metabolism,Small Molecule Biochemistry** | Incorporation of phosphatidylcholine | 0.00751 | GBA | 1 |
| **Drug Metabolism,Molecular Transport** | Efflux of glutathione | 0.00751 | EDN1 | 1 |
| **Cell Morphology** | Contractility of stellate cells | 0.00751 | EDN1 | 1 |
| **Cardiovascular Disease,Developmental Disorder,Organismal Injury and Abnormalities** | Congenital heart block | 0.00751 | EDN1 | 1 |
| **Neurological Disease** | Formation of myelin ovoid | 0.00751 | RAC1 | 1 |

| | | | | |
|---|---|---|---|---|
| **Cell Morphology,Cellular Assembly and Organization,Nervous System Development and Function** | Projection of corticothalamic axons | 0.00751 | RAC1 | 1 |
| **Lipid Metabolism,Small Molecule Biochemistry** | Incorporation of choline | 0.00751 | GBA | 1 |
| **Embryonic Development,Nervous System Development and Function,Organ Development,Organismal Development,Tissue Development** | Formation of telencephalon | 0.008 | DLX2, MKKS, NUMBL, PPP3CA, RAC1 | 5 |
| **Cellular Development,Cellular Growth and Proliferation,Nervous System Development and Function,Tissue Development** | Development of interneurons | 0.00971 | DLX2, LHX6 | 2 |
| **Cancer,Neurological Disease,Organismal Injury and Abnormalities** | Brain astrocytoma | 0.0128 | AK5, ANKRD30B, CACNG3, DBNDD1, DKK3, FAM184A, HSPBP1, NUMBL, ZDHHC4 | 9 |
| **Cellular Development,Cellular Growth and Proliferation,Nervous System Development and Function,Tissue Development** | Outgrowth of sensory neurons | 0.015 | CLU | 1 |
| **Nervous System Development and Function** | Thickness of optic nerve | 0.015 | DLX2 | 1 |
| **Nervous System Development and Function,Organ Morphology,Tissue Morphology** | Quantity of mossy fiber cells | 0.015 | RAC1 | 1 |
| **Cell Cycle,Nervous System Development and Function** | Mitogenesis of astrocytes | 0.015 | EDN1 | 1 |
| **Neurological Disease,Organismal Injury and Abnormalities** | Injury of spinal cord | 0.015 | CLU | 1 |
| **Cellular Development,Cellular Growth and Proliferation,Nervous System Development and Function,Tissue Development** | Development of striatal interneurons | 0.015 | LHX6 | 1 |
| **Cellular Assembly and Organization,Nervous System Development and Function** | Delay in sorting of axons | 0.015 | RAC1 | 1 |
| **Cell Morphology,Cellular Assembly and Organization,Nervous System Development and Function** | Extension of leading processes | 0.015 | RAC1 | 1 |
| **Cell Morphology,Nervous System Development and Function** | Modulation of dopaminergic neurons | 0.015 | EDN1 | 1 |
| **Cardiovascular System Development and Function,Tissue Morphology** | Contraction of cerebral artery | 0.015 | EDN1 | 1 |
| **Embryonic Development,Nervous System Development and Function,Organ Development,Organismal Development,Tissue Development** | Development of striatum | 0.0173 | DLX2, MKKS | 2 |
| **Cell-To-Cell Signaling and Interaction,Nervous System Development and Function** | Long term synaptic depression of perirhinal cortex | 0.0224 | HAPLN1 | 1 |
| **Cellular Assembly and Organization** | Density of plasma membrane projections | 0.0224 | NUMBL | 1 |
| **Cardiovascular System Development and Function** | Vasoconstriction of middle cerebral artery | 0.0224 | EDN1 | 1 |
| **Cellular Development,Cellular Growth and Proliferation,Nervous System Development and Function,Tissue Development** | Development of GABAergic neurons | 0.0224 | DLX2 | 1 |
| **Ophthalmic Disease,Organismal Injury and Abnormalities** | Proliferative vitreoretinopathy | 0.0224 | EDN1 | 1 |
| **Metabolic Disease,Neurological Disease,Organismal Injury and Abnormalities,Psychological Disorders** | Severe Alzheimer disease | 0.0224 | CLU | 1 |
| **Cellular Movement,Nervous System Development and Function** | Migration of GABAergic neurons | 0.0224 | LHX6 | 1 |
| **Infectious Diseases** | Advanced infection | 0.0224 | CLU | 1 |
| **Lipid Metabolism,Small Molecule Biochemistry** | Turnover of inositol phospholipid | 0.0224 | EDN1 | 1 |
| **Cell-To-Cell Signaling and Interaction,Endocrine System Development and Function,Nervous System Development and Function** | Desensitization of pituitary cells | 0.0224 | EDN1 | 1 |

| | | | | |
|---|---|---|---|---|
| **Cell Morphology,Nervous System Development and Function** | Contraction of stellate cells | 0.0224 | EDN1 | 1 |
| **Cellular Development,Cellular Growth and Proliferation,Nervous System Development and Function,Tissue Development** | Generation of cortical interneurons | 0.0224 | DLX2 | 1 |
| **Cell Morphology,Cellular Assembly and Organization,Nervous System Development and Function** | Projection of thalamocortical axons | 0.0224 | RAC1 | 1 |
| **Cell Morphology,Cellular Assembly and Organization,Cellular Development,Cellular Function and Maintenance,Cellular Growth and Proliferation,Nervous System Development and Function,Organismal Development,Tissue Development** | Branching of mandibular nerve | 0.0224 | DLX2 | 1 |
| **Cell Morphology,Cellular Assembly and Organization,Cellular Function and Maintenance,Nervous System Development and Function** | Formation of astrocyte projections | 0.0224 | RAC1 | 1 |
| **Cell Morphology,Cellular Assembly and Organization,Cellular Function and Maintenance** | Formation of lamellipodia | 0.0297 | RAC1 | 1 |
| **Cellular Assembly and Organization,Neurological Disease,Organismal Development,Tissue Development** | Formation of senile plaques | 0.0297 | CLU | 1 |
| **Nervous System Development and Function** | Regeneration of peripheral nerve | 0.0297 | CLU | 1 |
| **Nervous System Development and Function** | Spreading depression of cerebral cortex | 0.0297 | CLU | 1 |
| **Cell Morphology,Nervous System Development and Function,Neurological Disease,Tissue Morphology** | Abnormal morphology of interneurons | 0.0362 | LHX6, VIT | 2 |
| **Developmental Disorder,Embryonic Development,Organismal Development,Tissue Morphology** | Abnormal morphology of central canal | 0.037 | VIT | 1 |
| **Embryonic Development,Nervous System Development and Function,Organ Development,Organismal Development,Tissue Development** | Development of subpallium | 0.037 | DLX2 | 1 |
| **Cellular Assembly and Organization,Nervous System Development and Function,Tissue Morphology** | Quantity of axon terminals | 0.037 | RAC1 | 1 |
| **Cell Morphology,Cellular Development,Embryonic Development,Nervous System Development and Function,Tissue Development** | Patterning of dendrites | 0.037 | CRMP1 | 1 |
| **Nervous System Development and Function,Organ Morphology,Organismal Development** | Size of spinal cord | 0.037 | VIT | 1 |
| **Cell Morphology,Cellular Assembly and Organization,Cellular Development,Cellular Function and Maintenance,Cellular Growth and Proliferation,Embryonic Development,Nervous System Development and Function,Organismal Development,Tissue Development** | Branching morphogenesis of nerves | 0.037 | DLX2 | 1 |
| **Cellular Development,Nervous System Development and Function,Tissue Development** | Differentiation of projection neurons | 0.037 | DLX2 | 1 |
| **Cellular Compromise,Neurological Disease,Organismal Injury and Abnormalities** | Injury of axons | 0.037 | CLU | 1 |
| **Cardiovascular Disease,Neurological Disease,Organismal Injury and Abnormalities** | Cerebral ischemia | 0.0442 | CLU | 1 |
| **Cellular Function and Maintenance** | Endocytosis by neurons | 0.0442 | RAC1 | 1 |
| **Nervous System Development and Function** | Mechanotransduction of hair cells | 0.0442 | ADGRV1 | 1 |
| **Cell-To-Cell Signaling and Interaction,Nervous System Development and Function** | Synaptic transmission of superior cervical ganglion neurons | 0.0442 | DOC2A | 1 |

**Cluster 14**

| Categories | Diseases or Functions Annotation | p-Value | Molecules | # Molecules |
|---|---|---|---|---|

| | | | | |
|---|---|---|---|---|
| **Cancer,Neurological Disease,Organismal Injury and Abnormalities,Tumor Morphology** | Progression of glioma | 0.00294 | MIIP | 1 |
| **Cardiovascular Disease,Organismal Injury and Abnormalities,Tissue Morphology** | Size of ischemic lesion | 0.00587 | LGALS3 | 1 |
| **Embryonic Development,Nervous System Development and Function,Organ Development,Organismal Development,Tissue Development** | Formation of brain septum | 0.00879 | ADAM19 | 1 |
| **Nervous System Development and Function** | Remyelination of sciatic nerve | 0.00879 | ADAM19 | 1 |
| **Cell Death and Survival,Nervous System Development and Function,Tissue Morphology** | Quantity of apoptotic neurons | 0.0117 | LGALS3 | 1 |
| **Nervous System Development and Function,Organ Morphology,Organismal Development** | Mass of brain | 0.016 | CTSF, MBD5 | 2 |
| **Gene Expression** | Expression of mRNA | 0.0175 | GCDH | 1 |
| **Cellular Growth and Proliferation,Nervous System Development and Function,Organ Development** | Proliferation of hippocampal cells | 0.0204 | SLC2A8 | 1 |
| **Cellular Development,Cellular Growth and Proliferation,Hematological System Development and Function,Lymphoid Tissue Structure and Development,Nervous System Development and Function** | Proliferation of microglia | 0.029 | LGALS3 | 1 |
| **Cellular Assembly and Organization,Neurological Disease** | Deposition of amyloid-beta plaques | 0.046 | PDK1 | 1 |

### Cluster 15

| Categories | Diseases or Functions Annotation | p-Value | Molecules | # Molecules |
|---|---|---|---|---|
| **Neurological Disease** | Memory deficits | 0.031 | GBA | 1 |
| **Cell Death and Survival,Cell Morphology,Nervous System Development and Function,Neurological Disease,Tissue Morphology** | Loss of sympathetic neuron | 0.0397 | SHC2 | 1 |
| **Nervous System Development and Function,Neurological Disease,Organ Morphology,Organismal Development,Organismal Injury and Abnormalities** | Abnormal morphology of enlarged third cerebral ventricle | 0.044 | MKKS | 1 |

**Supplementary Table 7: Module 7**
**Cluster 2**

| Categories | Diseases or Functions Annotation | p-Value | Molecules | # Molecules |
|---|---|---|---|---|
| **Cellular Function and Maintenance,Molecular Transport** | Exocytosis | 2.75E-07 | APBA2, CADPS2, PIP5K1C, PPFIA3, RIMS2, RPH3A, UNC13A, UNC13B, UNC13C | 9 |
| **Cellular Assembly and Organization,Cellular Function and Maintenance,Cellular Movement,Molecular Transport,Nervous System Development and Function** | Exocytosis of synaptic vesicles | 0.00000174 | APBA2, PIP5K1C, PPFIA3, RPH3A, UNC13A, UNC13B, UNC13C | 7 |
| **Cellular Assembly and Organization,Cellular Movement,Nervous System Development and Function** | Transport of synaptic vesicles | 0.0000207 | APBA2, KIAA1107, PIP5K1C, PPFIA3, RPH3A, SCRIB, UNC13A, UNC13B, UNC13C | 9 |
| **Cell-To-Cell Signaling and Interaction,Cellular Assembly and Organization,Cellular Function and Maintenance,Molecular Transport** | Priming of synaptic vesicles | 0.0000281 | CADPS2, RIMS2, UNC13A, UNC13B, UNC13C | 5 |
| **Cellular Assembly and Organization,Nervous System Development and Function** | Docking of synaptic vesicles | 0.000106 | PPFIA3, RIMS2, UNC13A, UNC13B | 4 |
| **Neurological Disease,Organismal Injury and Abnormalities** | Hydrocephalus | 0.000471 | CELSR2, DVL1, DVL3, NCOR1 | 4 |
| **Auditory and Vestibular System Development and Function,Cell Morphology,Cellular Assembly and Organization** | Orientation of stereocilia bundles | 0.00128 | ADGRV1, DVL1, DVL3 | 3 |
| **Nervous System Development and Function,Neurological Disease** | Abnormal morphology of nervous system | 0.00302 | ABR, ADCY1, ADGRA2, ASPM, CADPS2, CELSR2, COL13A1, CREBBP, DVL1, HIVEP2, IDUA, KIF1B, MAN2C1, MCPH1, MEGF8, MTHFR, NCOR1, NOS2, NRN1, PER2, RIMS2, ROCK2, SCN1B, SEMA5B, SHANK2, SORL1, SPTBN4, SYNE1, TOP2B, UNC13A, UNC13B, VAX1 | 32 |

| | | | | |
|---|---|---|---|---|
| **Cell Morphology,Cell-To-Cell Signaling and Interaction,Cellular Assembly and Organization,Nervous System Development and Function,Neurological Disease,Tissue Morphology** | Abnormal morphology of synapse | 0.00666 | COL13A1, NRN1, ROCK2, SYNE1, UNC13A, UNC13B | 6 |
| **Behavior,Nervous System Development and Function** | Memory consolidation | 0.00698 | CREBBP, NR3C2 | 2 |
| **Nervous System Development and Function,Neurological Disease** | Abnormal morphology of central nervous system | 0.00767 | ABR, ADCY1, ADGRA2, ASPM, CADPS2, CELSR2, CREBBP, HIVEP2, IDUA, KIF1B, MAN2C1, MCPH1, MTHFR, NCOR1, NOS2, RIMS2, SCN1B, SHANK2, SPTBN4, TOP2B, VAX1 | 21 |
| **Nervous System Development and Function,Neurological Disease,Organ Morphology,Organismal Development,Organismal Injury and Abnormalities** | Abnormal morphology of forebrain | 0.00857 | ADCY1, ADGRA2, ASPM, CELSR2, CREBBP, HIVEP2, MAN2C1, NCOR1, NOS2, TOP2B, VAX1 | 11 |
| **Nervous System Development and Function,Neurological Disease,Ophthalmic Disease** | Abnormal morphology of optic nerve | 0.00887 | SCN1B, SPTBN4, VAX1 | 3 |
| **Nervous System Development and Function** | Morphology of nervous system | 0.00913 | ABR, ADCY1, ADGRA2, ASPM, CADPS2, CELSR2, COL13A1, CREBBP, DVL1, HIVEP2, IDUA, KIF1B, KIF5C, MAN2C1, MCPH1, MEGF8, MTHFR, NCOR1, NOS2, NRN1, PER2, RGS11, RIMS2, ROCK2, SCN1B, SEMA5B, SHANK2, SORL1, SPTBN4, SYNE1, TIAM1, TOP2B, UNC13A, UNC13B, VAX1 | 35 |
| **Cell-To-Cell Signaling and Interaction,Nervous System Development and Function** | Action potential of motor endplates | 0.0092 | COL13A1, RIMS2 | 2 |

| | | | | |
|---|---|---|---|---|
| **Behavior,Nervous System Development and Function** | Short-term memory | 0.0092 | CREBBP, NR3C2 | 2 |
| **Organismal Development,Organismal Injury and Abnormalities** | Abnormal morphology of head | 0.0102 | ABR, ADCY1, ADGRA2, ADGRV1, ASPM, CADPS2, CELSR2, CREBBP, DVL1, HIVEP2, IDUA, KIF1B, MAN2C1, MCPH1, MTHFR, NCOR1, NOS2, SHANK2, SPTBN4, TOP2B, VAX1 | 21 |
| **Lipid Metabolism,Molecular Transport,Small Molecule Biochemistry** | Concentration of arachidonic acid | 0.0117 | ARNTL, PTGS2 | 2 |
| **Nervous System Development and Function,Neurological Disease,Organ Morphology,Organismal Development,Organismal Injury and Abnormalities** | Abnormal morphology of anterior horn of the spinal cord | 0.0117 | KIF1B, RIMS2 | 2 |
| **Nervous System Development and Function,Neurological Disease,Organ Morphology,Organismal Development,Organismal Injury and Abnormalities** | Abnormal morphology of brain | 0.0137 | ABR, ADCY1, ADGRA2, ASPM, CADPS2, CELSR2, CREBBP, HIVEP2, IDUA, KIF1B, MAN2C1, MCPH1, MTHFR, NCOR1, NOS2, SHANK2, SPTBN4, TOP2B, VAX1 | 19 |
| **Cell-To-Cell Signaling and Interaction,Cellular Function and Maintenance,Nervous System Development and Function** | Paired-pulse facilitation of synapse | 0.0144 | APBA2, CADPS2, ROCK2, UNC13C | 4 |
| **Cell Morphology,Cellular Assembly and Organization,Cellular Development,Cellular Function and Maintenance,Cellular Growth and Proliferation,Nervous System Development and Function,Organismal Development,Tissue Development** | Formation of dendrites | 0.0173 | ABI1, ANAPC2, CELSR2, DVL1, FMN1, MYO5B, ROCK2, SYNE1 | 8 |
| **Behavior,Nervous System Development and Function** | Memory | 0.0175 | ADCY1, CREBBP, NEURL1, NR3C2 | 4 |
| **Cellular Assembly and Organization,Nervous System Development and Function,Tissue Morphology** | Abnormal quantity of synaptic vesicles | 0.0185 | KIF1B, UNC13A, UNC13B | 3 |
| **Carbohydrate Metabolism,Drug Metabolism,Molecular Transport,Small Molecule Biochemistry** | Quantity of heparan sulfate | 0.0189 | IDUA | 1 |
| **Cellular Assembly and Organization** | Aggregation of early endosomes | 0.0189 | KIF16B | 1 |
| **Cell Morphology,Cellular Assembly and Organization,Cellular Function and Maintenance** | Formation of axon-like process | 0.0189 | TIAM1 | 1 |
| **Cell Morphology,Cell-To-Cell Signaling and Interaction** | Length of stereocilia | 0.0189 | MYO15A | 1 |
| **Cell-To-Cell Signaling and Interaction,Cellular Assembly and Organization,Nervous System Development and Function,Tissue Morphology** | Quantity of ribbon synapse | 0.0189 | BSN | 1 |

| | | | | |
|---|---|---|---|---|
| **Cellular Compromise** | Acidification of cytosol | 0.0189 | CA4 | 1 |
| **Cell-To-Cell Signaling and Interaction,Nervous System Development and Function** | Inactivation of synapse | 0.0189 | BSN | 1 |
| **Cellular Development,Nervous System Development and Function,Organ Development** | Differentiation of preoptic area | 0.0189 | VAX1 | 1 |
| **Cellular Development,Nervous System Development and Function,Organ Development** | Differentiation of medial ganglionic eminences | 0.0189 | VAX1 | 1 |
| **Cellular Development,Cellular Growth and Proliferation,Embryonic Development,Nervous System Development and Function,Organ Development,Organismal Development,Tissue Development,Visual System Development and Function** | Formation of retinal cells | 0.0189 | ADCY1 | 1 |
| **Nervous System Development and Function,Organ Morphology,Tissue Morphology** | Quantity of rod bipolar cells | 0.0189 | PRDM1 | 1 |
| **Cell Morphology,Cellular Assembly and Organization,Cellular Development,Cellular Function and Maintenance,Cellular Growth and Proliferation,Embryonic Development,Nervous System Development and Function,Organismal Development,Tissue Development** | Delay in initiation of outgrowth of dendrites | 0.0189 | DAB2IP | 1 |
| **Cell-To-Cell Signaling and Interaction,Nervous System Development and Function** | Clustering of hippocampal neurons | 0.0189 | SHANK2 | 1 |
| **Nervous System Development and Function,Tissue Morphology** | Density of parallel fiber | 0.0189 | DAB2IP | 1 |
| **Cell Death and Survival,Cellular Development,Nervous System Development and Function,Tissue Development,Tissue Morphology** | Regeneration of bipolar cells | 0.0189 | PRDM1 | 1 |
| **Cellular Development,Nervous System Development and Function,Tissue Development** | Differentiation of pyramidal neurons | 0.0189 | UNC5D | 1 |
| **Embryonic Development,Nervous System Development and Function,Organ Development,Organismal Development,Tissue Development** | Development of septal area | 0.0189 | VAX1 | 1 |
| **Behavior** | Social exploration | 0.0189 | APBA2 | 1 |
| **Cardiovascular System Development and Function,Organismal Development** | Penetration by blood vessel | 0.0189 | ADGRA2 | 1 |
| **Developmental Disorder,Endocrine System Disorders,Nutritional Disease,Organismal Injury and Abnormalities,Reproductive System Disease** | Delayed puberty | 0.0189 | VAX1 | 1 |
| **Behavior** | Perseverance behavior | 0.0189 | NR3C2 | 1 |
| **Behavior** | Impulsivity | 0.0189 | ADCY1 | 1 |
| **Metabolic Disease,Neurological Disease,Organismal Injury and Abnormalities,Psychological Disorders** | Preclinical stage Alzheimer disease | 0.0189 | NOS2 | 1 |
| **Hematological Disease** | Hyperemia | 0.0189 | PTGS2 | 1 |
| **Nervous System Development and Function,Tissue Morphology** | Density of climbing fiber | 0.0189 | DAB2IP | 1 |
| **Cell-To-Cell Signaling and Interaction,Drug Metabolism,Small Molecule Biochemistry** | Conversion of norepinephrine | 0.0189 | DBH | 1 |
| **Infectious Diseases** | Replication of Murine cytomegalovirus | 0.0189 | NOS2 | 1 |
| **Metabolic Disease,Neurological Disease,Organismal Injury and Abnormalities,Psychological Disorders** | Stage V Alzheimer disease | 0.0189 | PTGS2 | 1 |
| **Cell Morphology,Cellular Assembly and Organization,Nervous System Development and Function** | Projection of mossy fibers | 0.0189 | NR3C2 | 1 |
| **Cell-To-Cell Signaling and Interaction,Nervous System Development and Function** | Activation of mitral cells | 0.0189 | GJD2 | 1 |
| **Cell Morphology,Cellular Assembly and Organization** | Orientation of plasma membrane projections | 0.0189 | SCN1B | 1 |
| **Cell Morphology,Cellular Assembly and Organization,Nervous System Development and Function,Tissue Morphology** | Diameter of nodes of Ranvier | 0.0189 | SPTBN4 | 1 |
| **Nervous System Development and Function** | Discharge of cochlear nerve | 0.0189 | BSN | 1 |
| **Lipid Metabolism,Molecular Transport,Small Molecule Biochemistry** | Concentration of eicosapentenoic acid | 0.0189 | ARNTL | 1 |

| | | | | |
|---|---|---|---|---|
| **Cell-To-Cell Signaling and Interaction,Cellular Function and Maintenance,Nervous System Development and Function** | Paired-pulse facilitation of parallel fiber-Purkinje cell synapses | 0.0189 | UNC13C | 1 |
| **Metabolic Disease,Neurological Disease,Organismal Injury and Abnormalities,Psychological Disorders** | Stage I Alzheimer disease | 0.0189 | PTGS2 | 1 |
| **Cellular Assembly and Organization,Cellular Function and Maintenance** | Microtubule dynamics | 0.0196 | ABI1, ADCY1, ADCY6, ANAPC2, ATCAY, BSN, CELSR2, DAB2IP, DVL1, ELK1, FMN1, GPRIN1, HERC1, MRTFA, MYO5B, NRN1, PIP5K1C, PTGS2, ROCK2, SCN1B, SEMA5B, SHANK2, SPTBN4, SYNE1, TIAM1, TOP2B | 26 |
| **Cellular Assembly and Organization** | Quantity of vesicles | 0.0211 | BSN, KIF1B, UNC13A, UNC13B | 4 |
| **Cell Morphology,Cellular Assembly and Organization,Cellular Development,Cellular Function and Maintenance,Cellular Growth and Proliferation,Nervous System Development and Function,Organismal Development,Tissue Development** | Neuritogenesis | 0.0223 | ABI1, ADCY1, ANAPC2, ATCAY, BSN, CELSR2, DAB2IP, DVL1, ELK1, FMN1, GPRIN1, HERC1, MRTFA, MYO5B, NRN1, PIP5K1C, PTGS2, ROCK2, SHANK2, SPTBN4, SYNE1, TIAM1, TOP2B | 23 |
| **Carbohydrate Metabolism** | Quantity of carbohydrate | 0.0238 | IDUA, PIP5K1C, PTGS2 | 3 |
| **Nervous System Development and Function,Neurological Disease,Organ Morphology,Organismal Development,Organismal Injury and Abnormalities** | Abnormal morphology of barrel cortex | 0.0241 | ADCY1, ASPM | 2 |
| **Drug Metabolism,Lipid Metabolism,Small Molecule Biochemistry** | Synthesis of prostaglandin E2 | 0.0241 | NOS2, PTGS2 | 2 |
| **Carbohydrate Metabolism,Lipid Metabolism,Molecular Transport,Small Molecule Biochemistry** | Concentration of phosphatidic acid | 0.0241 | PIP5K1C, PTGS2 | 2 |
| **Nervous System Development and Function,Neurological Disease,Organ Morphology,Organismal Development,Organismal Injury and Abnormalities** | Third ventricle dilation | 0.0278 | CELSR2, NCOR1 | 2 |
| **Cell Morphology,Cellular Movement,Nervous System Development and Function** | Innervation of motor neurons | 0.0278 | RIMS2, TOP2B | 2 |

| | | | | |
|---|---|---|---|---|
| **Cell-To-Cell Signaling and Interaction** | Activation of central nervous system cells | 0.0278 | GJD2, MGAT5B, PTGS2, SCN1B | 4 |
| **Nervous System Development and Function,Neurological Disease,Organ Morphology,Organismal Development,Organismal Injury and Abnormalities** | Abnormal morphology of telencephalon | 0.0307 | ADGRA2, ASPM, CELSR2, HIVEP2, MAN2C1, NOS2, TOP2B, VAX1 | 8 |
| **Nervous System Development and Function** | Morphology of central nervous system | 0.0313 | ABR, ADCY1, ADGRA2, ASPM, CADPS2, CELSR2, CREBBP, HIVEP2, IDUA, KIF1B, KIF5C, MAN2C1, MCPH1, MTHFR, NCOR1, NOS2, RIMS2, SCN1B, SHANK2, SPTBN4, TOP2B, VAX1 | 22 |
| **Cellular Assembly and Organization,Cellular Function and Maintenance,Cellular Movement,Nervous System Development and Function** | Endocytosis of synaptic vesicles | 0.0321 | KIAA1107, PIP5K1C, SCRIB | 3 |
| **Organismal Development** | Morphology of head | 0.0358 | ABR, ADCY1, ADGRA2, ADGRV1, ASPM, CADPS2, CELSR2, CREBBP, DVL1, HIVEP2, IDUA, KIF1B, KIF5C, MAN2C1, MCPH1, MTHFR, NCOR1, NOS2, SHANK2, SPTBN4, TOP2B, VAX1 | 22 |
| **Cellular Development,Nervous System Development and Function,Tissue Development** | Differentiation of neuron-like cells | 0.0374 | PIP5K1C | 1 |
| **Cell-To-Cell Signaling and Interaction,Nervous System Development and Function** | Short-term depression of synapse | 0.0374 | UNC13A | 1 |
| **Cellular Assembly and Organization,Cellular Function and Maintenance** | Penetration of microtubules | 0.0374 | TIAM1 | 1 |
| **Lipid Metabolism,Molecular Transport,Small Molecule Biochemistry** | Accumulation of ganglioside GM3 | 0.0374 | IDUA | 1 |
| **Cardiovascular System Development and Function,Organismal Development,Tissue Morphology** | Vasodilation of capillary vessel | 0.0374 | UNC13B | 1 |
| **Cell-To-Cell Signaling and Interaction,Nervous System Development and Function** | Neurotransmission of thalamocortical axons | 0.0374 | RIMS2 | 1 |
| **Cardiovascular System Development and Function** | Hypoperfusion of brain | 0.0374 | NOS2 | 1 |
| **Nervous System Development and Function,Neurological Disease** | Abnormal morphology of maxillary nerve | 0.0374 | MEGF8 | 1 |

| | | | | |
|---|---|---|---|---|
| **Reproductive System Development and Function** | Estrous cyclicity | 0.0374 | GJD2 | 1 |
| **Nervous System Development and Function,Organ Morphology** | Dilation of lateral cerebral ventricle | 0.0374 | POLL | 1 |
| **Carbohydrate Metabolism,Lipid Metabolism,Molecular Transport,Small Molecule Biochemistry** | Concentration of phosphatidylserine | 0.0374 | PTGS2 | 1 |
| **Cellular Assembly and Organization,Cellular Function and Maintenance** | Formation of Golgi vesicles | 0.0374 | BSN | 1 |
| **Cellular Growth and Proliferation,Nervous System Development and Function** | Inhibition of dorsal root ganglion cells | 0.0374 | ROCK2 | 1 |
| **Cell-To-Cell Signaling and Interaction,Nervous System Development and Function** | Long term synaptic depression of striatopallidal neurons | 0.0374 | ADCY5 | 1 |
| **Neurological Disease,Organismal Injury and Abnormalities,Psychological Disorders** | Damage of brain tissue | 0.0374 | NOS2 | 1 |
| **Metabolic Disease,Neurological Disease,Organismal Injury and Abnormalities,Psychological Disorders** | Stage II Alzheimer disease | 0.0374 | PTGS2 | 1 |
| **Cellular Assembly and Organization,Cellular Movement** | Segregation of retinogeniculate projection | 0.0374 | GJD2 | 1 |
| **Cell Cycle,Embryonic Development,Nervous System Development and Function** | Cell division of neuroblasts | 0.0374 | ASPM | 1 |
| **Cardiovascular Disease,Organismal Injury and Abnormalities** | Failure of heart | 0.0374 | PTGS2 | 1 |
| **Developmental Disorder,Endocrine System Disorders,Hereditary Disorder,Metabolic Disease,Neurological Disease,Organismal Injury and Abnormalities,Psychological Disorders** | Childhood adrenoleukodystrophy | 0.0374 | PTGS2 | 1 |
| **Nervous System Development and Function,Organ Morphology,Tissue Morphology** | Quantity of spiny stellate neurons | 0.0374 | RIMS2 | 1 |
| **Cellular Assembly and Organization** | Quantity of dense-core vesicles | 0.0374 | BSN | 1 |
| **Cell-To-Cell Signaling and Interaction,Nervous System Development and Function** | Coupling of neurons | 0.0374 | GJD2 | 1 |
| **Cellular Assembly and Organization** | Translocation of Golgi vesicles | 0.0374 | BSN | 1 |
| **Cell-To-Cell Signaling and Interaction,Nervous System Development and Function** | Long term synaptic depression of corticostriatal neurons | 0.0374 | ADCY5 | 1 |
| **Cancer,Neurological Disease,Organismal Injury and Abnormalities,Tumor Morphology** | Production of glioblastoma | 0.0374 | ROS1 | 1 |
| **Cell-To-Cell Signaling and Interaction,Inflammatory Response** | Cytotoxic reaction of macrophages | 0.0374 | NOS2 | 1 |
| **Cell-To-Cell Signaling and Interaction** | Dissociation of endothelial cells | 0.0374 | UNC13B | 1 |
| **Cell Morphology,Cellular Assembly and Organization,Cellular Function and Maintenance,Nervous System Development and Function** | Extension of neurites | 0.04 | ADCY6, ANAPC2, DVL1, MYO5B, SCN1B, SEMA5B, TIAM1 | 7 |
| **Nervous System Development and Function,Organ Morphology,Organismal Development** | Morphology of brain | 0.045 | ABR, ADCY1, ADGRA2, ASPM, CADPS2, CELSR2, CREBBP, HIVEP2, IDUA, KIF1B, KIF5C, MAN2C1, MCPH1, MTHFR, NCOR1, NOS2, SHANK2, SPTBN4, TOP2B, VAX1 | 20 |
| **Psychological Disorders** | Depressive disorder | 0.0468 | ABI1, CCKBR, | 5 |

| | | | | |
|---|---|---|---|---|
| | | | DVL1, LETMD1, MICAL2 | |
| **Nervous System Development and Function,Neurological Disease,Organ Morphology,Organismal Development,Organismal Injury and Abnormalities** | Abnormal morphology of cerebral neocortex | 0.049 | MAN2C1, MCPH1 | 2 |
| **Nervous System Development and Function,Tissue Morphology** | Quantity of oligodendrocytes | 0.0498 | HIVEP2, MGAT5B, NOS2 | 3 |

**Cluster 4**

| Categories | Diseases or Functions Annotation | p-Value | Molecules | # Molecules |
|---|---|---|---|---|
| **Cellular Assembly and Organization,Cellular Function and Maintenance** | Microtubule dynamics | 6.39E-08 | ABI1, ADCY1, ADCY6, ADGRB3, AGRN, APBA1, APC, ARHGEF28, ATCAY, BCL11B, BSN, CAMSAP1, CDKL3, CELSR2, COL25A1, CRMP1, CYFIP1, DBN1, DCC, EIF2AK4, EIF4G3, ELK1, ERC2, FMN1, GABRB3, GAP43, GNAO1, GPRIN1, HERC1, HRAS, HTR1A, KIF3C, KNDC1, LAMB1, LRRK2, MEF2A, MRTFA, MYO16, MYO5B, NRN1, PALLD, PIP5K1C, PTK2B, PTPRZ1, ROBO1, RYR2, SEMA4F, SHANK2, SIPA1L1, SORBS2, TNR, TRPC6, ULK4 | 53 |
| **Cell Morphology,Cellular Assembly and Organization,Cellular Development,Cellular Function and Maintenance,Cellular Growth and Proliferation,Nervous System Development and Function,Organismal Development,Tissue Development** | Neuritogenesis | 7.86E-08 | ABI1, ADCY1, ADGRB3, AGRN, APBA1, APC, ARHGEF28, | 48 |

| | | | | |
|---|---|---|---|---|
| | | | ATCAY, BCL11B, BSN, CAMSAP1, CDKL3, CELSR2, COL25A1, CRMP1, CYFIP1, DBN1, DCC, EIF2AK4, EIF4G3, ELK1, ERC2, FMN1, GABRB3, GAP43, GNAO1, GPRIN1, HERC1, HRAS, KNDC1, LAMB1, LRRK2, MEF2A, MRTFA, MYO16, MYO5B, NRN1, PALLD, PIP5K1C, PTK2B, PTPRZ1, ROBO1, RYR2, SHANK2, SIPA1L1, SORBS2, TNR, TRPC6 | |
| **Cellular Development,Cellular Growth and Proliferation,Nervous System Development and Function,Tissue Development** | Development of neurons | 3.28E-07 | ABI1, ADCY1, ADGRB3, AGRN, APBA1, APC, ARHGEF28, ATCAY, BCL11B, BSN, CAMSAP1, CDKL3, CELSR2, COL25A1, CRMP1, CYFIP1, DBN1, DCC, EIF2AK4, EIF4G3, ELK1, ERC2, FMN1, GABRB3, GAP43, GNAO1, GPRIN1, HERC1, HRAS, KNDC1, LAMB1, LRRK2, LRRTM4, MEF2A, MRTFA, MYO16, MYO5B, NCAN, NRN1, | 53 |

| | | | | |
|---|---|---|---|---|
| | | | PALLD, PCDH8, PIP5K1C, PLXND1, PTK2B, PTPRZ1, ROBO1, RYR2, SEZ6L, SHANK2, SIPA1L1, SORBS2, TNR, TRPC6 | |
| **Cellular Function and Maintenance,Molecular Transport** | Exocytosis | 6.16E-07 | APBA1, APBA2, ERC2, PIP5K1C, PPFIA3, RIMS2, RPH3A, SNAP25, UNC13A, UNC13B | 10 |
| **Cellular Assembly and Organization** | Transport of vesicles | 8.73E-07 | AP2M1, APBA1, APBA2, APC, DENND1A, KIAA1107, PCLO, PIP5K1C, PPFIA3, RPH3A, SNAP25, UNC13A, UNC13B | 13 |
| **Cellular Assembly and Organization,Cellular Function and Maintenance,Cellular Movement,Molecular Transport,Nervous System Development and Function** | Exocytosis of synaptic vesicles | 0.00000163 | APBA1, APBA2, PIP5K1C, PPFIA3, RPH3A, SNAP25, UNC13A, UNC13B | 8 |
| **Cellular Assembly and Organization,Cellular Movement,Nervous System Development and Function** | Transport of synaptic vesicles | 0.00000175 | AP2M1, APBA1, APBA2, DENND1A, KIAA1107, PCLO, PIP5K1C, PPFIA3, RPH3A, SNAP25, UNC13A, UNC13B | 12 |
| **Cellular Assembly and Organization,Nervous System Development and Function** | Docking of synaptic vesicles | 0.0000236 | PPFIA3, RIMS2, SNAP25, UNC13A, UNC13B | 5 |
| **Cell Morphology** | Shape change of neurites | 0.0000913 | ABI1, AGRN, APBA1, BCL11B, BSN, COL25A1, CRMP1, DBN1, DCC, EIF4G3, ELK1, GABRB3, GAP43, GNAO1, HRAS, KNDC1, | 25 |

| | | | | |
|---|---|---|---|---|
| | | | MYO5B, NR3C2, NRN1, ROBO1, RYR2, SHANK2, SIPA1L1, SORBS2, TNR | |
| **Cell Morphology,Cellular Assembly and Organization,Cellular Development,Cellular Function and Maintenance,Cellular Growth and Proliferation,Embryonic Development,Nervous System Development and Function,Organismal Development,Tissue Development** | Dendritic growth/branching | 0.000104 | ABI1, AGRN, APBA1, BSN, CRMP1, DBN1, EIF4G3, ELK1, GABRB3, GAP43, GNAO1, HRAS, KNDC1, MYO5B, NRN1, ROBO1, RYR2, SHANK2, SIPA1L1, SORBS2, TNR | 21 |
| **Cell Morphology,Nervous System Development and Function** | Innervation of optic nerve | 0.000191 | DCC, GAP43, VAX1 | 3 |
| **Cellular Assembly and Organization** | Priming of vesicles | 0.000324 | ERC2, RIMS2, SNAP25, UNC13A, UNC13B | 5 |
| **Nervous System Development and Function,Neurological Disease** | Abnormal morphology of nervous system | 0.000453 | ABR, ADCY1, AGRN, ARNT2, CACNA1C, CELSR2, COL25A1, CRMP1, DBN1, DCC, E2F5, ERC2, FAT3, FES, GABRB3, GAP43, GMFB, GRIA2, HIVEP2, ITGA6, LRRK2, LRRTM4, MYO16, NOS2, NRN1, PRKN, PROKR2, PTPN13, PTPRZ1, RIMS2, ROBO1, SHANK2, SHC2, SIL1, SLC4A3, SNAP25, SNTA1, SORBS2, TCF7L2, TENM2, TNR, ULK4, UNC13A, | 46 |

| | | | | |
|---|---|---|---|---|
| | | | UNC13B, VAX1, VPS13A | |
| **Cell Morphology,Cellular Assembly and Organization,Cellular Development,Cellular Function and Maintenance,Cellular Growth and Proliferation,Nervous System Development and Function,Organismal Development,Tissue Development** | Branching of neurons | 0.000533 | ABI1, AGRN, APBA1, BSN, COL25A1, CRMP1, DBN1, EIF4G3, ELK1, ERC2, GABRB3, GAP43, GNAO1, HRAS, KNDC1, MYO5B, NRN1, ROBO1, RYR2, SHANK2, SIPA1L1, SORBS2, TNR | 23 |
| **Cancer,Organismal Injury and Abnormalities** | Non-hematological solid tumor | 0.000621 | ABCF1, ADAMTS8, AGAP3, ANKRD30B, APC, ASXL3, ATR, COL11A1, DHRS4, DNAH8, DPP10, ERC2, HERC1, HRAS, ITGA6, KMT2D, MYO5B, NPR1, P2RX5, PCLO, PIGQ, PIK3R1, PLB1, PROKR2, PTPRT, PTPRZ1, RIMS2, ROS1, RYR2, SEMA4F, SMARCA4, THEMIS, TSHZ2, UNC13B | 34 |
| **Cellular Assembly and Organization,Cellular Function and Maintenance** | Formation of Golgi vesicles | 0.000737 | BSN, PCLO | 2 |
| **Cellular Assembly and Organization** | Translocation of Golgi vesicles | 0.000737 | BSN, PCLO | 2 |
| **Nervous System Development and Function** | Morphology of nervous system | 0.000883 | ABR, ADCY1, AGRN, AKAP13, APBA1, ARNT2, CACNA1C, CELSR2, COL25A1, CRMP1, DBN1, DCC, E2F5, ERC2, FAT3, FES, GABRB3, GAP43, | 52 |

| | | | | |
|---|---|---|---|---|
| | | | GMFB, GRIA2, HIVEP2, ITGA6, KIF5C, LRRK2, LRRTM4, MYO16, NOS2, NRN1, PLAGL1, PRDM16, PRKN, PROKR2, PTPN13, PTPRZ1, RIMS2, ROBO1, SHANK2, SHC2, SIL1, SIPA1L1, SLC4A3, SNAP25, SNTA1, SORBS2, TCF7L2, TENM2, TNR, ULK4, UNC13A, UNC13B, VAX1, VPS13A | |
| **Nervous System Development and Function,Neurological Disease** | Abnormal morphology of central nervous system | 0.000932 | ABR, ADCY1, ARNT2, CELSR2, CRMP1, DBN1, DCC, E2F5, FES, GABRB3, GAP43, GMFB, GRIA2, HIVEP2, ITGA6, LRRTM4, NOS2, PRKN, PROKR2, PTPRZ1, RIMS2, ROBO1, SHANK2, SIL1, SLC4A3, SNAP25, TCF7L2, TENM2, TNR, ULK4, VAX1 | 31 |
| **Cell Morphology,Cellular Assembly and Organization,Cellular Development,Cellular Function and Maintenance,Cellular Growth and Proliferation,Embryonic Development,Nervous System Development and Function,Organismal Development,Tissue Development** | Branching of neurites | 0.000932 | ABI1, AGRN, APBA1, BSN, COL25A1, CRMP1, DBN1, EIF4G3, ELK1, GABRB3, GAP43, GNAO1, HRAS, KNDC1, MYO5B, NRN1, | 22 |

| | | | | |
|---|---|---|---|---|
| | | | ROBO1, RYR2, SHANK2, SIPA1L1, SORBS2, TNR | |
| **Cancer,Organismal Injury and Abnormalities** | Non-melanoma solid tumor | 0.000989 | ABCF1, ADAMTS8, AGAP3, ANKRD30B, APC, ASXL3, ATR, COL11A1, DNAH8, DPP10, ERC2, HERC1, HRAS, ITGA6, KMT2D, MYO5B, NPR1, P2RX5, PCLO, PIK3R1, PLB1, PROKR2, PTPRT, PTPRZ1, RIMS2, ROS1, RYR2, SEMA4F, SMARCA4, THEMIS, TSHZ2, UNC13B | 32 |
| **Cell-To-Cell Signaling and Interaction,Cellular Assembly and Organization** | Stabilization of synapse | 0.00101 | NCAN, NRN1, TNR | 3 |
| **Cancer,Neurological Disease,Organismal Injury and Abnormalities** | Brain tumor | 0.00101 | ABCF1, ADAMTS8, AGAP3, ANKRD30B, APC, ASXL3, ATR, COL11A1, DHRS4, DNAH8, DPP10, ERC2, HERC1, HRAS, KMT2D, MYO5B, NPR1, P2RX5, PCLO, PIK3R1, PLB1, PROKR2, PTPRT, PTPRZ1, RIMS2, ROS1, RYR2, SEMA4F, SMARCA4, THEMIS, TSHZ2, UNC13B | 32 |
| **Cancer,Neurological Disease,Organismal Injury and Abnormalities** | Nervous system neoplasm | 0.0011 | ABCF1, ADAMTS8, AGAP3, ANKRD30B, APC, ASXL3, | 33 |

| | | | | |
|---|---|---|---|---|
| | | | ATR, COL11A1, DHRS4, DNAH8, DPP10, ERC2, HERC1, HRAS, KMT2D, MYO5B, NPR1, P2RX5, PCLO, PIGQ, PIK3R1, PLB1, PROKR2, PTPRT, PTPRZ1, RIMS2, ROS1, RYR2, SEMA4F, SMARCA4, THEMIS, TSHZ2, UNC13B | |
| **Cellular Assembly and Organization,Nervous System Development and Function,Tissue Morphology** | Abnormal quantity of synaptic vesicles | 0.00117 | ERC2, PCLO, TNR, UNC13A, UNC13B | 5 |
| **Cancer,Neurological Disease,Organismal Injury and Abnormalities** | Brain astrocytoma | 0.00121 | ABCF1, ADAMTS8, AGAP3, ANKRD30B, APC, ASXL3, ATR, COL11A1, DNAH8, DPP10, ERC2, HERC1, HRAS, MYO5B, P2RX5, PCLO, PIK3R1, PLB1, PROKR2, PTPRZ1, RIMS2, ROS1, RYR2, SEMA4F, THEMIS, TSHZ2 | 26 |
| **Organismal Development,Organismal Injury and Abnormalities** | Abnormal morphology of head | 0.00143 | ABR, ADCY1, ADGRV1, ARNT2, CELSR2, CRMP1, DBN1, DCC, E2F5, FAT3, FES, GABRB3, GAP43, GMFB, GRIA2, HIVEP2, ITGA6, LRRTM4, NOS2, PRKN, PROKR2, ROBO1, | 31 |

| | | | | |
|---|---|---|---|---|
| | | | SHANK2, SIL1, SLC4A3, SNAP25, TCF7L2, TENM2, TNR, ULK4, VAX1 | |
| **Nervous System Development and Function** | Morphology of central nervous system | 0.00167 | ABR, ADCY1, AKAP13, ARNT2, CELSR2, CRMP1, DBN1, DCC, E2F5, FES, GABRB3, GAP43, GMFB, GRIA2, HIVEP2, ITGA6, KIF5C, LRRTM4, MYO16, NOS2, PRDM16, PRKN, PROKR2, PTPRZ1, RIMS2, ROBO1, SHANK2, SIL1, SLC4A3, SNAP25, TCF7L2, TENM2, TNR, ULK4, VAX1 | 35 |
| **Cell-To-Cell Signaling and Interaction,Cellular Assembly and Organization** | Cell-cell contact | 0.00178 | AGRN, BSN, DCC, ERC2, GAP43, LRRTM4, NCAN, NEURL1, NRN1, PCDH8, PLXND1, SEZ6L, SHANK2, SNTA1, SORBS2, TNR, UNC13A, UNC13B | 18 |
| **Cancer,Neurological Disease,Organismal Injury and Abnormalities** | Brain glioma | 0.00182 | ABCF1, ADAMTS8, AGAP3, ANKRD30B, APC, ASXL3, ATR, COL11A1, DNAH8, DPP10, ERC2, HERC1, HRAS, KMT2D, MYO5B, NPR1, P2RX5, PCLO, PIK3R1, PLB1, | 30 |

| | | | | |
|---|---|---|---|---|
| | | | PROKR2, PTPRT, PTPRZ1, RIMS2, ROS1, RYR2, SEMA4F, SMARCA4, THEMIS, TSHZ2 | |
| **Cell-To-Cell Signaling and Interaction,Cellular Assembly and Organization,Cellular Development,Cellular Function and Maintenance,Cellular Growth and Proliferation,Nervous System Development and Function,Tissue Development** | Developmental process of synapse | 0.00183 | AGRN, BSN, DCC, LRRTM4, NCAN, NRN1, PCDH8, PLXND1, SEZ6L, SHANK2, SORBS2, TNR | 12 |
| **Cell-To-Cell Signaling and Interaction,Cellular Assembly and Organization,Cellular Function and Maintenance,Molecular Transport** | Priming of synaptic vesicles | 0.00184 | ERC2, RIMS2, UNC13A, UNC13B | 4 |
| **Nervous System Development and Function** | Pathfinding of neurons | 0.00207 | DCC, GAP43, ROBO1 | 3 |
| **Cell-To-Cell Signaling and Interaction,Nervous System Development and Function** | Activation of synapse | 0.00207 | APBA1, APBA2, GRIA2 | 3 |
| **Nervous System Development and Function,Neurological Disease** | Abnormal morphology of ependyma | 0.00207 | E2F5, ULK4, VAX1 | 3 |
| **Embryonic Development,Nervous System Development and Function,Organ Development,Organismal Development,Tissue Development,Visual System Development and Function** | Formation of eye | 0.00207 | ADCY1, FAT3, SLC4A3 | 3 |
| **Organismal Development** | Morphology of head | 0.00212 | ABR, ADCY1, ADGRV1, AKAP13, ARNT2, CELSR2, CRMP1, DBN1, DCC, E2F5, FAT3, FES, GABRB3, GAP43, GMFB, GRIA2, HIVEP2, ITGA6, KIF5C, LRRTM4, MYO16, NOS2, PRDM16, PRKN, PROKR2, ROBO1, SHANK2, SIL1, SLC4A3, SNAP25, TCF7L2, TENM2, TNR, ULK4, VAX1 | 35 |
| **Hereditary Disorder,Neurological Disease,Organismal Injury and Abnormalities,Psychological Disorders,Skeletal and Muscular Disorders** | Huntington disease grade 4 Huntington disease | 0.00217 | RPH3A, SNAP25 | 2 |

| | | | | |
|---|---|---|---|---|
| **Hereditary Disorder,Neurological Disease,Organismal Injury and Abnormalities,Psychological Disorders,Skeletal and Muscular Disorders** | Familial Parkinson disease | 0.00217 | GAP43, PRKN | 2 |
| **Nervous System Development and Function,Organismal Development** | Secretion of cerebrospinal fluid | 0.00217 | CELSR2, E2F5 | 2 |
| **Nervous System Development and Function,Neurological Disease,Organ Morphology,Organismal Development,Organismal Injury and Abnormalities** | Abnormal morphology of brain | 0.00232 | ABR, ADCY1, ARNT2, CELSR2, CRMP1, DBN1, DCC, E2F5, FES, GABRB3, GAP43, GMFB, GRIA2, HIVEP2, ITGA6, LRRTM4, NOS2, PRKN, PROKR2, ROBO1, SHANK2, SIL1, SNAP25, TCF7L2, TENM2, TNR, ULK4, VAX1 | 28 |
| **Cell Morphology,Cellular Assembly and Organization,Cellular Development,Cellular Function and Maintenance,Cellular Growth and Proliferation,Nervous System Development and Function,Organismal Development,Tissue Development** | Formation of dendrites | 0.00285 | ABI1, ADGRB3, CDKL3, CELSR2, FMN1, KNDC1, MEF2A, MYO5B, PTPRZ1, ROBO1, SIPA1L1, TRPC6 | 12 |
| **Nervous System Development and Function,Organ Morphology,Organismal Development** | Morphology of brain | 0.0031 | ABR, ADCY1, AKAP13, ARNT2, CELSR2, CRMP1, DBN1, DCC, E2F5, FES, GABRB3, GAP43, GMFB, GRIA2, HIVEP2, ITGA6, KIF5C, LRRTM4, MYO16, NOS2, PRDM16, PRKN, PROKR2, ROBO1, SHANK2, SIL1, SNAP25, TCF7L2, TENM2, TNR, ULK4, VAX1 | 32 |
| **Cellular Assembly and Organization** | Quantity of vesicles | 0.00399 | BSN, ERC2, PCLO, TNR, | 6 |

| | | | | |
|---|---|---|---|---|
| | | | UNC13A, UNC13B | |
| **Tissue Morphology** | Quantity of cells | 0.00425 | ACVR1C, AGRN, BSN, CACNA1C, COL25A1, DCC, ERC2, GBA, GNAO1, HIVEP2, HRAS, KIF5C, LRRK2, MGAT5B, NEURL1, NOS2, NR3C2, NRN1, PCLO, PRDM16, PRKN, PTPRZ1, RIMS2, SHC2, TCF7L2, TNR, TRPM2, UNC13A, UNC13B, VAX1 | 30 |
| **Behavior,Nervous System Development and Function** | Working memory | 0.00426 | KCNH3, NR3C2 | 2 |
| **Nervous System Development and Function,Neurological Disease,Ophthalmic Disease,Organ Morphology,Organismal Development,Organismal Injury and Abnormalities** | Abnormal morphology of visual cortex | 0.00426 | HIVEP2, TENM2 | 2 |
| **Cell Morphology** | Projection | 0.00499 | BCL11B, DCC, NR3C2, ROBO1 | 4 |
| **Cell Morphology,Cell-To-Cell Signaling and Interaction,Nervous System Development and Function** | Plasticity of synapse | 0.00512 | DBN1, GRIA2, HRAS, KCNH3, NEURL1, PCDH8, PIP5K1C, PPFIA3, PTK2B, SIPA1L1 | 10 |
| **Cell Morphology,Cell-To-Cell Signaling and Interaction,Cellular Assembly and Organization,Nervous System Development and Function,Tissue Morphology** | Morphology of synapse | 0.00549 | AGRN, ERC2, GAP43, NRN1, SNTA1, TNR, UNC13A, UNC13B | 8 |
| **Cellular Assembly and Organization,Cellular Function and Maintenance** | Stabilization of microtubules | 0.0058 | APC, HTR1A, KIF3C | 3 |
| **Nervous System Development and Function,Neurological Disease,Organ Morphology,Organismal Development,Organismal Injury and Abnormalities** | Third ventricle dilation | 0.0058 | CELSR2, E2F5, ULK4 | 3 |
| **Nervous System Development and Function,Tissue Morphology** | Density of neurons | 0.00652 | ACVR1C, CRMP1, ELK1, GABRB3, GAP43, GNAO1, HRAS, LRRTM4, | 10 |

| | | | | |
|---|---|---|---|---|
| | | | MEF2A, RYR2 | |
| **Cell Morphology,Nervous System Development and Function** | Innervation | 0.00695 | DCC, GABRB3, GAP43, GRIA2, RIMS2, SEZ6L, TENM2, VAX1 | 8 |
| **Cell Morphology,Cellular Development,Embryonic Development,Nervous System Development and Function,Tissue Development** | Patterning of dendrites | 0.00698 | CELSR2, CRMP1 | 2 |
| **Cellular Development,Nervous System Development and Function,Tissue Development** | Differentiation of nerve ending | 0.00698 | AGRN, MEF2A | 2 |
| **Hereditary Disorder,Neurological Disease,Organismal Injury and Abnormalities,Psychological Disorders,Skeletal and Muscular Disorders** | Huntington disease grade 3 Huntington disease | 0.00698 | RPH3A, SNAP25 | 2 |
| **Behavior** | Learning | 0.00938 | ADCY1, ADGRB3, GABRB3, KCNH3, KMT2D, NEURL1, NR3C2 | 7 |
| **Amino Acid Metabolism,Molecular Transport,Small Molecule Biochemistry** | Uptake of amino acids | 0.0102 | GAP43, SLC32A1, TNR | 3 |
| **Nervous System Development and Function** | Depotentiation | 0.0103 | ADCY1, GRIA2 | 2 |
| **Endocrine System Development and Function,Nervous System Development and Function,Organ Morphology,Tissue Morphology** | Quantity of lutenizing hormone-releasing hormone neurons | 0.0103 | DCC, VAX1 | 2 |
| **Cell Morphology,Cell-To-Cell Signaling and Interaction,Cellular Assembly and Organization,Nervous System Development and Function,Neurological Disease,Tissue Morphology** | Abnormal morphology of synapse | 0.0106 | AGRN, ERC2, NRN1, SNTA1, TNR, UNC13A, UNC13B | 7 |
| **Nervous System Development and Function,Tissue Morphology** | Quantity of neurons | 0.0115 | ACVR1C, AGRN, BSN, CACNA1C, COL25A1, DCC, ERC2, GNAO1, HRAS, KIF5C, NEURL1, NRN1, PCLO, PRKN, PTPRZ1, RIMS2, SHC2, TNR, TRPM2, UNC13A, UNC13B, VAX1 | 22 |
| **Cell Morphology,Nervous System Development and Function,Neurological Disease,Tissue Morphology** | Abnormal morphology of neurons | 0.0121 | AGRN, CACNA1C, COL25A1, CRMP1, DBN1, ERC2, FAT3, GAP43, HIVEP2, LRRK2, MYO16, NOS2, NRN1, PRKN, PTPN13, | 26 |

| | | | | |
|---|---|---|---|---|
| | | | ROBO1, SHANK2, SHC2, SIL1, SLC4A3, SNTA1, SORBS2, TNR, UNC13A, UNC13B, VPS13A | |
| **Nervous System Development and Function,Neurological Disease,Organ Morphology,Organismal Development,Organismal Injury and Abnormalities** | Abnormal morphology of third cerebral ventricle | 0.0131 | ARNT2, CELSR2, E2F5, ULK4 | 4 |
| **Cell Morphology,Cellular Assembly and Organization,Nervous System Development and Function** | Projection of axons | 0.014 | BCL11B, DCC, NR3C2 | 3 |
| **Cancer,Organismal Injury and Abnormalities** | Malignant solid tumor | 0.014 | APC, DHRS4, HRAS, ITGA6, PIGQ, ROS1, SMARCA4 | 7 |
| **Cell Signaling,Nucleic Acid Metabolism,Small Molecule Biochemistry** | Binding of guanosine 5'-O-(3-thiotriphosphate) | 0.0141 | GNAO1, HTR1A | 2 |
| **Cellular Assembly and Organization** | Clustering of synaptic vesicles | 0.0141 | BSN, PCLO | 2 |
| **Cellular Movement,Nervous System Development and Function** | Pathfinding of axons | 0.0141 | DCC, GAP43 | 2 |
| **Cell-To-Cell Signaling and Interaction,Drug Metabolism,Small Molecule Biochemistry** | Metabolism of dopamine | 0.0141 | PRKN, SORCS2 | 2 |
| **Nervous System Development and Function,Organ Morphology,Organismal Development** | Morphology of forebrain | 0.0159 | ADCY1, AKAP13, ARNT2, CELSR2, DCC, E2F5, FES, GAP43, HIVEP2, NOS2, PROKR2, ROBO1, TENM2, ULK4, VAX1 | 15 |
| **Cell Morphology,Nervous System Development and Function,Neurological Disease,Tissue Morphology** | Abnormal morphology of bipolar cells | 0.0161 | ERC2, FAT3, SLC4A3 | 3 |
| **Embryonic Development,Nervous System Development and Function,Organ Development,Organismal Development,Tissue Development** | Formation of olfactory bulb | 0.0165 | GNAO1, LRRK2, PROKR2, RTP1, VAX1 | 5 |
| **Nervous System Development and Function,Neurological Disease,Organ Morphology,Organismal Development,Organismal Injury and Abnormalities** | Abnormal morphology of telencephalon | 0.017 | CELSR2, DCC, E2F5, FES, HIVEP2, NOS2, PROKR2, ROBO1, TENM2, ULK4, VAX1 | 11 |
| **Cellular Assembly and Organization,Cellular Function and Maintenance,Cellular Movement,Nervous System Development and Function** | Endocytosis of synaptic vesicles | 0.0175 | AP2M1, DENND1A, KIAA1107, PIP5K1C | 4 |
| **Cellular Development,Cellular Growth and Proliferation,Nervous System Development and Function,Tissue Development** | Proliferation of neuronal cells | 0.0176 | ADCY6, AGRN, APBA2, CRMP1, CYFIP1, DCC, GAP43, GNAO1, HRAS, ITGA6, | 25 |

| | | | | |
|---|---|---|---|---|
| | | | ITGA9, KIF3C, KNDC1, LRRK2, MYO10, NCAN, NRN1, PALLD, PLAGL1, PROKR2, PTK2B, PTPRZ1, ROBO1, SNAP25, VAX1 | |
| **Cell-To-Cell Signaling and Interaction,Nervous System Development and Function** | Action potential of motor endplates | 0.0185 | RIMS2, SNAP25 | 2 |
| **Cell Morphology,Embryonic Development,Nervous System Development and Function,Neurological Disease,Ophthalmic Disease,Organ Development,Organ Morphology,Organismal Development,Organismal Injury and Abnormalities,Tissue Development,Tissue Morphology,Visual System Development and Function** | Abnormal morphology of rod bipolar cells | 0.0185 | FAT3, SLC4A3 | 2 |
| **Nervous System Development and Function,Neurological Disease,Organ Morphology,Organismal Development,Organismal Injury and Abnormalities** | Abnormal morphology of cerebral cortex | 0.0195 | ADCY1, CRMP1, DBN1, E2F5, FES, GAP43, GRIA2, HIVEP2, ITGA6, LRRTM4, SNAP25, TNR | 12 |
| **Cancer,Organismal Injury and Abnormalities** | Nonhematologic malignant neoplasm | 0.0204 | APC, DHRS4, ITGA6, PIGQ, ROS1, SMARCA4 | 6 |
| **Nervous System Development and Function,Neurological Disease,Organ Morphology,Organismal Development,Organismal Injury and Abnormalities** | Abnormal morphology of forebrain | 0.021 | ADCY1, ARNT2, CELSR2, DCC, E2F5, FES, HIVEP2, NOS2, PROKR2, ROBO1, TENM2, ULK4, VAX1 | 13 |
| **Tissue Development** | Organization of nervous tissue | 0.0211 | AGRN, ARHGEF28, DCC, LRRTM4, PTPRO, SHANK2, TNR | 7 |
| **Neurological Disease,Organismal Injury and Abnormalities** | Abnormality of cerebral cortex | 0.0219 | ADCY1, CRMP1, DBN1, E2F5, FES, GAP43, GRIA2, HIVEP2, ITGA6, LRRK2, LRRTM4, PRKN, SNAP25, TNR, TP53BP2 | 15 |
| **Cell Morphology** | Shape change of axons | 0.0221 | AGRN, BCL11B, COL25A1, | 7 |

| | | | | |
|---|---|---|---|---|
| | | | DCC, GAP43, NR3C2, NRN1 | |
| **Cell-To-Cell Signaling and Interaction,Nervous System Development and Function** | Long-term potentiation of mossy fibers | 0.0234 | ADCY1, GRIA2 | 2 |
| **Amino Acid Metabolism,Molecular Transport,Small Molecule Biochemistry** | Uptake of L-glutamic acid | 0.0234 | GAP43, TNR | 2 |
| **Nervous System Development and Function,Neurological Disease,Ophthalmic Disease** | Abnormal morphology of optic nerve | 0.0236 | DCC, SLC4A3, VAX1 | 3 |
| **Cell Morphology,Nervous System Development and Function,Tissue Morphology** | Morphology of neurons | 0.0242 | AGRN, APBA1, CACNA1C, COL25A1, CRMP1, DBN1, ERC2, FAT3, GAP43, HIVEP2, LRRK2, MYO16, NOS2, NRN1, PLAGL1, PRKN, PTPN13, RIMS2, ROBO1, SHANK2, SHC2, SIL1, SIPA1L1, SLC4A3, SNTA1, SORBS2, TNR, UNC13A, UNC13B, VPS13A | 30 |
| **Cellular Function and Maintenance** | Endocytosis | 0.0248 | AP2M1, DENND1A, KIAA1107, PIP5K1C, SNAP25 | 5 |
| **Cellular Development,Cellular Growth and Proliferation,Nervous System Development and Function,Tissue Development** | Growth of axons | 0.0269 | CRMP1, CYFIP1, DCC, GAP43, GNAO1, KIF3C, LRRK2, NRN1, PROKR2, ROBO1, SNAP25 | 11 |
| **Free Radical Scavenging,Lipid Metabolism,Molecular Transport,Small Molecule Biochemistry** | Accumulation of lipid peroxide | 0.0272 | KL | 1 |
| **Cell-To-Cell Signaling and Interaction,Cellular Assembly and Organization,Nervous System Development and Function** | Modulation of synapse | 0.0272 | PCDH8 | 1 |
| **Cellular Development,Nervous System Development and Function,Tissue Development** | Differentiation of sympathetic preganglionic neurons | 0.0272 | AGRN | 1 |
| **Cell-To-Cell Signaling and Interaction,Cellular Assembly and Organization,Nervous System Development and Function** | Binding of presynaptic membrane | 0.0272 | AP2M1 | 1 |
| **Cancer,Organismal Injury and Abnormalities,Reproductive System Disease** | Prostatic carcinoma | 0.0272 | ITGA6 | 1 |
| **Cellular Assembly and Organization** | Aggregation of early endosomes | 0.0272 | KIF16B | 1 |
| **Developmental Disorder,Nervous System Development and Function,Neurological Disease,Organ Morphology,Organismal** | Complete agenesis of corpus callosum | 0.0272 | DCC | 1 |

| | | | | |
|---|---|---|---|---|
| **Development,Organismal Injury and Abnormalities** | | | | |
| **Cell-To-Cell Signaling and Interaction,Hematological System Development and Function,Immune Cell Trafficking,Inflammatory Response,Nervous System Development and Function** | Binding of microglia | 0.0272 | TNR | 1 |
| **Cell-To-Cell Signaling and Interaction,Cellular Assembly and Organization,Nervous System Development and Function,Tissue Morphology** | Quantity of ribbon synapse | 0.0272 | BSN | 1 |
| **Carbohydrate Metabolism,Cellular Function and Maintenance,Small Molecule Biochemistry** | Homeostasis of D-glucose | 0.0272 | FOXO1 | 1 |
| **Cell Signaling,Nucleic Acid Metabolism,Small Molecule Biochemistry** | Inhibition of cyclic AMP | 0.0272 | HTR1A | 1 |
| **Nervous System Development and Function,Tissue Morphology** | Density of serotonergic neurons | 0.0272 | GAP43 | 1 |
| **Nervous System Development and Function** | Function of autonomic nerve | 0.0272 | SNAP25 | 1 |
| **Hereditary Disorder,Neurological Disease,Organismal Injury and Abnormalities,Psychological Disorders,Skeletal and Muscular Disorders** | Autosomal recessive juvenile Parkinson disease | 0.0272 | PRKN | 1 |
| **Cell-To-Cell Signaling and Interaction,Nervous System Development and Function** | Inactivation of synapse | 0.0272 | BSN | 1 |
| **Cellular Development,Nervous System Development and Function,Organ Development** | Differentiation of preoptic area | 0.0272 | VAX1 | 1 |
| **Cellular Development,Nervous System Development and Function,Organ Development** | Differentiation of medial ganglionic eminences | 0.0272 | VAX1 | 1 |
| **Cellular Development,Cellular Growth and Proliferation,Embryonic Development,Nervous System Development and Function,Organ Development,Organismal Development,Tissue Development,Visual System Development and Function** | Formation of retinal cells | 0.0272 | ADCY1 | 1 |
| **Cell-To-Cell Signaling and Interaction,Nervous System Development and Function** | Clustering of hippocampal neurons | 0.0272 | SHANK2 | 1 |
| **Molecular Transport,Protein Trafficking** | Translocation of protein fragment | 0.0272 | AGAP3 | 1 |
| **Cell-To-Cell Signaling and Interaction,Nervous System Development and Function** | Activation of medium spiny neurons | 0.0272 | LRRK2 | 1 |
| **Carbohydrate Metabolism,Lipid Metabolism,Small Molecule Biochemistry** | Incorporation of phosphatidylcholine | 0.0272 | GBA | 1 |
| **Nervous System Development and Function,Organ Morphology,Organismal Development** | Size of optic chiasm | 0.0272 | GAP43 | 1 |
| **Cell Morphology,Cellular Assembly and Organization,Cellular Development,Cellular Function and Maintenance,Cellular Growth and Proliferation,Nervous System Development and Function,Organismal Development,Tissue Development** | Sprouting of rod bipolar cells | 0.0272 | ERC2 | 1 |
| **Cellular Development,Nervous System Development and Function,Tissue Development** | Differentiation of pyramidal neurons | 0.0272 | UNC5D | 1 |
| **Cellular Development** | Delay in initiation of transition of radial glial cells | 0.0272 | PLAGL1 | 1 |
| **Cellular Development** | Delay in initiation of transition of neuronal progenitor cells | 0.0272 | PLAGL1 | 1 |
| **Hereditary Disorder,Neurological Disease,Organismal Injury and Abnormalities,Psychological Disorders,Skeletal and Muscular Disorders** | Huntington disease grade 1 Huntington disease | 0.0272 | SNAP25 | 1 |
| **Cell-To-Cell Signaling and Interaction,Nervous System Development and Function** | Conduction of neuroglia | 0.0272 | TNR | 1 |
| **Nervous System Development and Function** | Curvature of nerves | 0.0272 | DCC | 1 |
| **Cellular Function and Maintenance,Skeletal and Muscular System Development and Function** | Depolarization of smooth muscle | 0.0272 | TRPC6 | 1 |
| **Embryonic Development,Nervous System Development and Function,Organ Development,Organismal Development,Tissue Development** | Development of septal area | 0.0272 | VAX1 | 1 |
| **Behavior** | Social exploration | 0.0272 | APBA2 | 1 |

| | | | | |
|---|---|---|---|---|
| **Hereditary Disorder,Neurological Disease,Organismal Injury and Abnormalities,Psychological Disorders,Skeletal and Muscular Disorders** | Parkinson-dementia syndrome | 0.0272 | GAP43 | 1 |
| **Developmental Disorder,Endocrine System Disorders,Nutritional Disease,Organismal Injury and Abnormalities,Reproductive System Disease** | Delayed puberty | 0.0272 | VAX1 | 1 |
| **Lipid Metabolism,Small Molecule Biochemistry** | Incorporation of choline | 0.0272 | GBA | 1 |
| **Neurological Disease** | Mechanical hypersensitivity of neurons | 0.0272 | CACNA2D1 | 1 |
| **Cellular Assembly and Organization,Cellular Compromise,Nervous System Development and Function** | Retraction of glial cell projections | 0.0272 | GRIA2 | 1 |
| **Cellular Development,Cellular Growth and Proliferation,Embryonic Development,Nervous System Development and Function,Organismal Development,Tissue Development** | Outgrowth of neuroblasts | 0.0272 | ITGA6 | 1 |
| **Lipid Metabolism,Molecular Transport,Small Molecule Biochemistry** | Accumulation of glucosylceramide | 0.0272 | GBA | 1 |
| **Behavior** | Perseverance behavior | 0.0272 | NR3C2 | 1 |
| **Cellular Development,Cellular Growth and Proliferation,Nervous System Development and Function,Tissue Development** | Development of corticospinal neurons | 0.0272 | BCL11B | 1 |
| **Behavior** | Impulsivity | 0.0272 | ADCY1 | 1 |
| **Metabolic Disease,Neurological Disease,Organismal Injury and Abnormalities,Psychological Disorders** | Preclinical stage Alzheimer disease | 0.0272 | NOS2 | 1 |
| **Cell Cycle,Nervous System Development and Function** | Arrest in cell cycle progression of oligodendrocyte precursor cells | 0.0272 | HRAS | 1 |
| **Embryonic Development,Nervous System Development and Function,Organ Development,Organismal Development,Tissue Development** | Development of layer IV | 0.0272 | GAP43 | 1 |
| **Endocrine System Development and Function,Endocrine System Disorders,Nervous System Development and Function,Neurological Disease,Organ Morphology,Organismal Development,Organismal Injury and Abnormalities,Reproductive System Disease** | Abnormal morphology of enlarged adenohypophysis | 0.0272 | TCF7L2 | 1 |
| **Cellular Assembly and Organization** | Delay in initiation of priming of vesicles | 0.0272 | SNAP25 | 1 |
| **Infectious Diseases** | Replication of Murine cytomegalovirus | 0.0272 | NOS2 | 1 |
| **Behavior** | Risk preference | 0.0272 | GABRB3 | 1 |
| **Cell Morphology,Cellular Assembly and Organization,Nervous System Development and Function** | Projection of mossy fibers | 0.0272 | NR3C2 | 1 |
| **Lipid Metabolism,Small Molecule Biochemistry** | Incorporation of sphingomyelin | 0.0272 | GBA | 1 |
| **Cell-To-Cell Signaling and Interaction,Nervous System Development and Function** | Excitation of postsynaptic region | 0.0272 | PCDH8 | 1 |
| **Amino Acid Metabolism,Lipid Metabolism,Small Molecule Biochemistry** | Translocation of GABA | 0.0272 | SLC32A1 | 1 |
| **Cellular Assembly and Organization,Cellular Development,Nervous System Development and Function,Tissue Development** | Differentiation of presynaptic terminals | 0.0272 | AGRN | 1 |
| **Endocrine System Development and Function,Nervous System Development and Function,Organ Morphology,Tissue Morphology** | Quantity of oxytocin neurons | 0.0272 | DCC | 1 |
| **Small Molecule Biochemistry** | Generation of hydrogen sulfide | 0.0272 | HMOX2 | 1 |
| **Cell-To-Cell Signaling and Interaction,Drug Metabolism,Small Molecule Biochemistry** | Catabolism of dopamine | 0.0272 | PRKN | 1 |
| **Nervous System Development and Function,Neurological Disease,Organ Morphology,Organismal Development,Organismal Injury and Abnormalities** | Lack of optic tract | 0.0272 | GAP43 | 1 |

| | | | | |
|---|---|---|---|---|
| **Tissue Development** | Organization of axon initial segments | 0.0272 | DCC | 1 |
| **Cellular Development,Nervous System Development and Function,Tissue Development** | Differentiation of postsynaptic neurons | 0.0272 | AGRN | 1 |
| **Nervous System Development and Function** | Discharge of cochlear nerve | 0.0272 | BSN | 1 |
| **Cell Morphology,Cellular Movement,Nervous System Development and Function** | Innervation of serotonergic neurons | 0.0272 | GAP43 | 1 |
| **Cell Morphology,Cellular Assembly and Organization** | Elongation of astrocyte projections | 0.0272 | GAP43 | 1 |
| **Lipid Metabolism,Molecular Transport,Small Molecule Biochemistry** | Concentration of eicosapentenoic acid | 0.0272 | ARNTL | 1 |
| **Embryonic Development,Organismal Development,Tissue Morphology** | Morphology of visceral endoderm | 0.0272 | AMOT | 1 |
| **Cardiovascular Disease,Hematological Disease,Metabolic Disease** | Hypertriglyceridemia | 0.0272 | FOXO1 | 1 |
| **Cell-To-Cell Signaling and Interaction,Nervous System Development and Function** | Synaptic transmission of cells | 0.028 | BSN, ERC2, GRIA2, HTR1A, NR3C2, SNAP25 | 6 |
| **Cellular Assembly and Organization,Cellular Function and Maintenance,Cellular Movement,Nervous System Development and Function** | Axonal transport of mitochondria | 0.0287 | PRKN, RANBP9 | 2 |
| **Cell Morphology,Cellular Movement,Nervous System Development and Function** | Innervation of Purkinje cells | 0.0287 | GRIA2, SEZ6L | 2 |
| **Cell-To-Cell Signaling and Interaction,Cellular Assembly and Organization,Cellular Development,Cellular Function and Maintenance,Cellular Growth and Proliferation,Nervous System Development and Function,Tissue Development** | Formation of excitatory synapses | 0.0287 | LRRTM4, SORBS2 | 2 |
| **Cellular Development,Cellular Growth and Proliferation,Nervous System Development and Function,Tissue Development** | Growth of neurites | 0.0339 | ADCY6, APBA2, CRMP1, CYFIP1, DCC, GAP43, GNAO1, HRAS, ITGA9, KIF3C, KNDC1, LRRK2, MYO10, NCAN, NRN1, PALLD, PROKR2, PTK2B, PTPRZ1, ROBO1, SNAP25 | 21 |
| **Cellular Movement,Nervous System Development and Function** | Guidance of axons | 0.0341 | DCC, GAP43, HRAS, KIF5C, PIK3R1, PTPRO, RANBP9, ROBO1, SEMA4F, UNC5D, VAX1 | 11 |
| **Cellular Development,Cellular Growth and Proliferation,Nervous System Development and Function,Tissue Development** | Outgrowth of axons | 0.0344 | DCC, GAP43, KIF3C, LRRK2, NRN1, PROKR2, ROBO1 | 7 |

| | | | | |
|---|---|---|---|---|
| **Neurological Disease,Organismal Injury and Abnormalities,Psychological Disorders,Skeletal and Muscular Disorders** | Parkinson disease with dementia | 0.0345 | GAP43, NOS2 | 2 |
| **Cancer,Organismal Injury and Abnormalities** | Development of benign tumor | 0.0345 | HRAS, UNC13B | 2 |
| **Cell Morphology,Cellular Assembly and Organization,Cellular Development,Cellular Function and Maintenance,Cellular Growth and Proliferation,Nervous System Development and Function,Organismal Development,Tissue Development** | Axonogenesis | 0.0387 | ADCY1, AGRN, APC, ARHGEF28, BCL11B, COL25A1, CYFIP1, DCC, GAP43, NRN1, PIP5K1C, PTPRZ1, ROBO1 | 13 |
| **Cell Morphology,Cellular Movement,Nervous System Development and Function** | Innervation of neurons | 0.0392 | GABRB3, GAP43, GRIA2, RIMS2, SEZ6L, TENM2 | 6 |
| **Nervous System Development and Function,Neurological Disease,Organ Morphology,Organismal Development,Organismal Injury and Abnormalities** | Abnormal morphology of choroid plexus | 0.0395 | E2F5, ITGA6, ULK4 | 3 |
| **Developmental Disorder,Neurological Disease** | Hypoplasia of cranial nerve | 0.0406 | DCC, GABRB3 | 2 |
| **Inflammatory Disease,Inflammatory Response,Neurological Disease,Organismal Injury and Abnormalities** | Inflammation of cerebral cortex | 0.0406 | LRRK2, TP53BP2 | 2 |
| **Nervous System Development and Function,Neurological Disease,Organ Morphology,Organismal Development,Organismal Injury and Abnormalities** | Abnormal morphology of corpus callosum | 0.0409 | DCC, NOS2, ROBO1, VAX1 | 4 |
| **Nervous System Development and Function,Neurological Disease,Organ Morphology,Organismal Development,Organismal Injury and Abnormalities** | Abnormal morphology of diencephalon | 0.0409 | ADCY1, ARNT2, CELSR2, E2F5, TENM2, ULK4 | 6 |
| **Cellular Development,Cellular Growth and Proliferation,Nervous System Development and Function,Tissue Development** | Outgrowth of neurons | 0.041 | ADCY6, APBA2, DCC, GAP43, GNAO1, HRAS, ITGA6, ITGA9, KIF3C, LRRK2, MYO10, NRN1, PALLD, PROKR2, PTK2B, PTPRZ1, ROBO1 | 17 |
| **Embryonic Development,Organismal Development** | Development of head | 0.0434 | ABCB6, ABR, ADCY1, ARNT2, CDK5RAP1, CELSR2, EIF2AK4, FAT3, GAP43, GNAO1, ITGA6, LRRK2, MRTFA, MYO16, | 26 |

| | | | | |
|---|---|---|---|---|
| | | | NME7, NOS2, PROKR2, ROBO1, ROGDI, RTP1, SLC4A3, SMARCA4, TACC2, TCF7L2, ULK4, VAX1 | |
| **Nervous System Development and Function** | Sensory system development | 0.0437 | ADCY1, FAT3, KIF5C, SHC2, SLC4A3 | 5 |
| **Cell Morphology,Cellular Assembly and Organization,Cellular Development,Cellular Function and Maintenance,Cellular Growth and Proliferation,Embryonic Development,Nervous System Development and Function,Organismal Development,Tissue Development,Tissue Morphology** | Density of dendritic spines | 0.0447 | CRMP1, ELK1, GABRB3, GNAO1, HRAS, RYR2 | 6 |
| **Cell-To-Cell Signaling and Interaction,Nervous System Development and Function** | Neurotransmission | 0.0454 | BSN, ERC2, FOXO1, GNAO1, GRIA2, HTR1A, NR3C2, PRKN, RIMS2, SNAP25 | 10 |
| **Nervous System Development and Function,Neurological Disease,Organ Morphology,Organismal Development,Organismal Injury and Abnormalities** | Abnormal morphology of barrel cortex | 0.0472 | ADCY1, GAP43 | 2 |
| **Energy Production,Molecular Transport,Nucleic Acid Metabolism,Small Molecule Biochemistry** | Concentration of ATP | 0.0472 | HMOX2, SLC25A23 | 2 |
| **Cell-To-Cell Signaling and Interaction** | Synaptic depression | 0.049 | ADCY1, ADCY5, GRIA2, PCLO, PIP5K1C, PTK2B, SHANK2, UNC13A | 8 |
| **Cancer,Neurological Disease,Organismal Injury and Abnormalities** | Cerebellar neoplasm | 0.0493 | APC, DHRS4, HRAS, SMARCA4 | 4 |

### Cluster 8

| Categories | Diseases or Functions Annotation | p-Value | Molecules | # Molecules |
|---|---|---|---|---|
| **Gene Expression** | Transcription of RNA | 0.00034 | CREBBP, KNG1, TNF | 3 |
| **Cell Death and Survival** | Apoptosis of multilineage progenitor cells | 0.000887 | ELP1, TNF | 2 |
| **Cell-To-Cell Signaling and Interaction,Nervous System Development and Function** | Action potential of motor endplates | 0.00118 | COL13A1, RIMS2 | 2 |
| **Nervous System Development and Function,Neurological Disease,Organ Morphology,Organismal Development,Organismal Injury and Abnormalities** | Abnormal morphology of anterior horn of the spinal cord | 0.00151 | KIF1B, RIMS2 | 2 |
| **Lipid Metabolism,Molecular Transport,Small Molecule Biochemistry** | Accumulation of sphingolipid | 0.00273 | IDUA, TNF | 2 |
| **Carbohydrate Metabolism** | Accumulation of carbohydrate | 0.00321 | IDUA, KNG1 | 2 |
| **Nervous System Development and Function,Neurological Disease,Organ Morphology,Organismal** | Abnormal morphology of spinal cord | 0.00366 | CREBBP, KIF1B, RIMS2 | 3 |

| | | | | |
|---|---|---|---|---|
| **Development,Organismal Injury and Abnormalities** | | | | |
| **Cellular Development,Cellular Growth and Proliferation,Embryonic Development,Organismal Development,Tissue Development** | Proliferation of granule neuron progenitor cells | 0.00373 | SCN1B, TSC2 | 2 |
| **Cell Death and Survival,Organismal Injury and Abnormalities** | Apoptosis of endothelial cells | 0.00488 | SMPD1, TNF | 2 |
| **Lipid Metabolism,Small Molecule Biochemistry** | Synthesis of eicosanoid | 0.00617 | KNG1, TNF | 2 |
| **Carbohydrate Metabolism,Drug Metabolism,Molecular Transport,Small Molecule Biochemistry** | Quantity of heparan sulfate | 0.00661 | IDUA | 1 |
| **Cell-To-Cell Signaling and Interaction,Nervous System Development and Function** | Sensitization of trigeminovascular neuron | 0.00661 | KNG1 | 1 |
| **Cell Death and Survival** | Cell death of retinal ganglion cells | 0.00661 | TNF | 1 |
| **Behavior,Nervous System Development and Function** | Spontaneous pain behavior | 0.00661 | TNF | 1 |
| **Behavior,Developmental Disorder,Neurological Disease,Psychological Disorders** | Autistic-like traits | 0.00661 | TSC2 | 1 |
| **Lipid Metabolism,Small Molecule Biochemistry** | Storage of sphingomyelin | 0.00661 | SMPD1 | 1 |
| **Cell Morphology,Cellular Assembly and Organization** | Orientation of plasma membrane projections | 0.00661 | SCN1B | 1 |
| **Hematological System Development and Function,Lymphoid Tissue Structure and Development,Tissue Morphology** | Quantity of CD8+ T lymphocyte | 0.00661 | SMPD1 | 1 |
| **Inflammatory Response** | Inflammatory response of cerebral cortex | 0.00661 | KNG1 | 1 |
| **Cell Death and Survival,Cell Morphology,Nervous System Development and Function,Neurological Disease,Organ Morphology,Organismal Development,Organismal Injury and Abnormalities,Respiratory Disease,Respiratory System Development and Function,Tissue Morphology** | Loss of olfactory receptor neurons | 0.00661 | TNF | 1 |
| **Cardiovascular System Development and Function,Nervous System Development and Function,Organ Morphology,Organismal Development,Tissue Morphology** | Diameter of basilar artery | 0.00661 | KNG1 | 1 |
| **Cell-To-Cell Signaling and Interaction,Nervous System Development and Function** | Firing of trigeminovascular neuron | 0.00661 | KNG1 | 1 |
| **Cell Death and Survival,Nervous System Development and Function** | Survival of retinal ganglion cells | 0.00661 | TNF | 1 |
| **Infectious Diseases** | Infection by measles virus | 0.00661 | SMPD1 | 1 |
| **Nervous System Development and Function,Skeletal and Muscular System Development and Function** | Motor function | 0.00836 | RIMS2, SMPD1 | 2 |
| **Neurological Disease,Psychological Disorders** | Pervasive developmental disorder | 0.00836 | SLC6A7, TSC2 | 2 |
| **Nervous System Development and Function** | Morphology of central nervous system | 0.0115 | CREBBP, EPHB3, IDUA, KIF1B, KNG1, PCM1, RIMS2, SCN1B, SLC2A8, SMPD1 | 11 |
| **Cell Death and Survival,Neurological Disease,Organismal Injury and Abnormalities** | Apoptosis of lactotropes | 0.0132 | TNF | 1 |
| **Lipid Metabolism,Small Molecule Biochemistry** | Liberation of arachidonic acid | 0.0132 | KNG1 | 1 |
| **Cell Morphology,Cellular Assembly and Organization,Nervous System Development and Function** | Extension of axon branches | 0.0132 | EPHB3 | 1 |
| **Lipid Metabolism,Molecular Transport,Small Molecule Biochemistry** | Accumulation of ganglioside GM3 | 0.0132 | IDUA | 1 |

| | | | | |
|---|---|---|---|---|
| **Nervous System Development and Function,Organ Morphology,Tissue Morphology** | Quantity of spiny stellate neurons | 0.0132 | RIMS2 | 1 |
| **Cell-To-Cell Signaling and Interaction,Nervous System Development and Function** | Neurotransmission of thalamocortical axons | 0.0132 | RIMS2 | 1 |
| **Cell-To-Cell Signaling and Interaction,Nervous System Development and Function** | Action potential of superior cervical ganglion neurons | 0.0132 | KNG1 | 1 |
| **Lipid Metabolism,Small Molecule Biochemistry** | Liberation of stearic acid | 0.0132 | KNG1 | 1 |
| **Cell Death and Survival,Neurological Disease,Organismal Injury and Abnormalities** | Apoptosis of somatotrophs | 0.0132 | TNF | 1 |
| **Cell Morphology,Cellular Assembly and Organization,Cellular Development,Cellular Function and Maintenance,Cellular Growth and Proliferation,Nervous System Development and Function,Organismal Development,Tissue Development** | Branching of axon branches | 0.0132 | EPHB3 | 1 |
| **Cellular Assembly and Organization,Cellular Movement,Nervous System Development and Function** | Migration of axons | 0.0132 | EPHB3 | 1 |
| **Cardiovascular Disease,Hematological Disease,Organismal Injury and Abnormalities** | Thrombus | 0.0132 | KNG1 | 1 |
| **Cell Cycle,Gene Expression** | Binding of synthetic promoter | 0.0132 | TNF | 1 |
| **Protein Synthesis** | Synthesis of protein | 0.0167 | TNF, TSC2 | 2 |
| **Nervous System Development and Function,Tissue Morphology** | Quantity of astrocytes | 0.0167 | TNF, TSC2 | 2 |
| **Organismal Injury and Abnormalities** | Edema | 0.0167 | KNG1, TNF | 2 |
| **Cell-To-Cell Signaling and Interaction,Nervous System Development and Function** | Action potential of neurons | 0.0188 | KNG1, TNF | 2 |
| **Cardiovascular System Development and Function,Nervous System Development and Function,Organ Morphology,Tissue Morphology** | Permeability of blood-brain barrier | 0.0188 | KNG1, TNF | 2 |
| **Cardiovascular Disease,Organismal Injury and Abnormalities** | Fibrosis of blood vessel | 0.0197 | TNF | 1 |
| **Lipid Metabolism,Small Molecule Biochemistry** | Synthesis of 12-hydroxyeicosatetraenoic acid | 0.0197 | KNG1 | 1 |
| **Cell-To-Cell Signaling and Interaction,Nervous System Development and Function** | Action potential of GABAergic neurons | 0.0197 | TNF | 1 |
| **Lipid Metabolism,Molecular Transport,Small Molecule Biochemistry** | Accumulation of ceramide | 0.0197 | TNF | 1 |
| **Lipid Metabolism,Small Molecule Biochemistry** | Synthesis of dinoprost | 0.0197 | TNF | 1 |
| **Infectious Diseases,Neurological Disease** | Infection of neurons | 0.0197 | SMPD1 | 1 |
| **Neurological Disease,Organismal Injury and Abnormalities** | Edema of white matter | 0.0197 | TNF | 1 |
| **Cardiovascular Disease,Inflammatory Disease,Neurological Disease,Organismal Injury and Abnormalities** | Vasculitis of central nervous system | 0.0197 | TNF | 1 |
| **Carbohydrate Metabolism,Molecular Transport,Small Molecule Biochemistry** | Accumulation of inositol phosphate | 0.0197 | KNG1 | 1 |
| **Cell Morphology,Cellular Function and Maintenance,Nervous System Development and Function** | Mitochondrial membrane potential of oligodendrocytes | 0.0197 | TNF | 1 |
| **Cellular Function and Maintenance** | Cellular homeostasis | 0.021 | KNG1, TNF, TSC2 | 3 |
| **Nervous System Development and Function,Neurological Disease** | Abnormal morphology of central nervous system | 0.0222 | CREBBP, EPHB3, IDUA, KIF1B, RIMS2, SCN1B, SLC2A8, SMPD1, TNF | 9 |
| **Cell Morphology,Cell-To-Cell Signaling and Interaction,Cellular Assembly and Organization,Nervous System Development and Function,Neurological Disease,Tissue Morphology** | Abnormal morphology of neuromuscular synapse | 0.0223 | COL13A1, SYNE2 | 2 |
| **Cell-To-Cell Signaling and Interaction,Cellular Movement,Hematological System Development and Function,Immune Cell Trafficking,Inflammatory Response** | Recruitment of monocytes | 0.0262 | TNF | 1 |
| **Neurological Disease** | Synaptic dysfunction | 0.0262 | TNF | 1 |
| **Behavior** | Exploratory behavior | 0.0262 | RIMS2 | 1 |

| | | | | |
|---|---|---|---|---|
| **Cell-mediated Immune Response,Cell-To-Cell Signaling and Interaction,Cellular Movement,Hematological System Development and Function,Immune Cell Trafficking** | Recruitment of T lymphocytes | 0.0262 | TNF | 1 |
| **Carbohydrate Metabolism,Lipid Metabolism,Small Molecule Biochemistry** | Hydrolysis of phosphatidylinositol 4,5-diphosphate | 0.0262 | KNG1 | 1 |
| **Protein Trafficking** | Interaction of protein | 0.0262 | KNG1 | 1 |
| **Carbohydrate Metabolism,Molecular Transport,Small Molecule Biochemistry** | Accumulation of glycosaminoglycan | 0.0262 | IDUA | 1 |
| **Cell Morphology,Cell-To-Cell Signaling and Interaction,Cellular Function and Maintenance,Nervous System Development and Function** | Depolarization of superior cervical ganglion neurons | 0.0262 | KNG1 | 1 |
| **Cell Death and Survival** | Apoptosis of neuronal progenitor cells | 0.0262 | ELP1 | 1 |
| **Nervous System Development and Function,Skeletal and Muscular System Development and Function** | Sensorimotor integration | 0.0262 | RIMS2 | 1 |
| **Nervous System Development and Function** | Morphology of nervous system | 0.0306 | COL13A1, CREBBP, EPHB3, IDUA, KIF1B, KNG1, PCM1, RGS11, RIMS2, SCN1B | 14 |
| **Cellular Assembly and Organization** | Modification of cytoskeleton | 0.0326 | TNF | 1 |
| **Cell-To-Cell Signaling and Interaction,Cellular Function and Maintenance,Nervous System Development and Function** | Discharge of neurons | 0.0326 | TNF | 1 |
| **DNA Replication, Recombination, and Repair** | Mitochondrial DNA damage | 0.0326 | TNF | 1 |
| **Cell-To-Cell Signaling and Interaction,Cellular Movement,Hematological System Development and Function,Immune Cell Trafficking,Inflammatory Response** | Recruitment of neutrophils | 0.0326 | TNF | 1 |
| **Lipid Metabolism,Molecular Transport,Small Molecule Biochemistry** | Concentration of sphingomyelin | 0.0326 | SMPD1 | 1 |
| **Cell-To-Cell Signaling and Interaction,Nervous System Development and Function** | Response of sensory neurons | 0.0326 | TNF | 1 |
| **Cell Morphology,Cellular Function and Maintenance** | Autophagy of cortical neurons | 0.0326 | TSC2 | 1 |
| **Cell Morphology,Cellular Assembly and Organization,Cellular Development,Cellular Function and Maintenance,Cellular Growth and Proliferation,Nervous System Development and Function,Neurological Disease,Organismal Development,Tissue Development,Tissue Morphology** | Abnormal pruning of axons | 0.0326 | EPHB3 | 1 |
| **Cell Death and Survival,Cellular Compromise,Neurological Disease,Organismal Injury and Abnormalities,Tissue Morphology** | Neurodegeneration of Purkinje cells | 0.0385 | IDUA, SMPD1 | 2 |
| **Cell-To-Cell Signaling and Interaction,Nervous System Development and Function** | Activation of sensory neurons | 0.039 | TNF | 1 |
| **Neurological Disease,Organismal Injury and Abnormalities** | Reactive astrogliosis | 0.039 | TNF | 1 |
| **Gene Expression** | Expression of mRNA | 0.039 | TNF | 1 |
| **Cellular Development,Cellular Growth and Proliferation,Nervous System Development and Function,Tissue Development** | Myelination of oligodendrocytes | 0.039 | TNF | 1 |
| **Hematological Disease,Neurological Disease,Organismal Injury and Abnormalities** | Disrupted blood brain barrier | 0.039 | TNF | 1 |
| **Inflammatory Disease,Inflammatory Response,Neurological Disease** | Inflammation of meninges | 0.039 | TNF | 1 |
| **Cell Morphology,Nervous System Development and Function,Neurological Disease,Tissue Morphology** | Abnormal morphology of axons | 0.0405 | EPHB3, SCN1B, TNF | 3 |
| **Cell-To-Cell Signaling and Interaction,Nervous System Development and Function** | Neurotransmission | 0.0427 | KIF1B, KNG1, RIMS2, TNF | 4 |

| | | | | |
|---|---|---|---|---|
| **Nervous System Development and Function,Organ Morphology,Organismal Development** | Morphology of brain | 0.0429 | CREBBP, EPHB3, IDUA, KIF1B, KNG1, PCM1, SLC2A8, SMPD1, TNF | 9 |
| **Behavior,Nervous System Development and Function** | Memory consolidation | 0.0454 | CREBBP | 1 |
| **Cellular Assembly and Organization,Nervous System Development and Function** | Accumulation of neurofilaments | 0.0454 | TNF | 1 |
| **Cell Signaling,Nucleic Acid Metabolism,Small Molecule Biochemistry** | Binding of guanosine 5'-O-(3-thiotriphosphate) | 0.0454 | KNG1 | 1 |
| **Molecular Transport,Protein Trafficking** | Transport of protein | 0.0454 | KIF1B | 1 |
| **Cellular Growth and Proliferation,Nervous System Development and Function,Organ Development** | Proliferation of hippocampal cells | 0.0454 | SLC2A8 | 1 |
| **Nervous System Development and Function** | Function of vestibular system | 0.0454 | EPHB3 | 1 |
| **Cell Signaling,Molecular Transport,Small Molecule Biochemistry** | Release of nitric oxide | 0.0454 | TNF | 1 |

### Supplementary Table 8: Module 8
### Cluster 4

| Categories | Diseases or Functions Annotation | p-Value | Molecules | # Molecules |
|---|---|---|---|---|
| **Nervous System Development and Function,Organ Morphology,Organismal Development** | Thickness of granule cell layer | 0.000651 | L1CAM, NRCAM | 2 |
| **Embryonic Development,Nervous System Development and Function,Organ Development,Organismal Development,Tissue Development** | Development of cerebral cortex | 0.000889 | ATAT1, DCLK2, DLX2, EMX1, FOXG1, L1CAM, MKKS, NEUROG2, OTX1, RAC1 | 13 |
| **Embryonic Development,Nervous System Development and Function,Organ Development,Organismal Development,Tissue Development** | Formation of hippocampus | 0.000934 | ATAT1, DCLK2, DLX2, FOXG1, L1CAM, MKKS, OTX1, RAC1, SRF, TMEM108 | 10 |
| **Embryonic Development,Nervous System Development and Function,Organ Development,Organismal Development,Tissue Development** | Formation of telencephalon | 0.00341 | DLX2, EMX1, HPRT1, ID2, L1CAM, MKKS, NEUROG2, NUMBL, PPP3CA, RAC1 | 11 |
| **Amino Acid Metabolism,Small Molecule Biochemistry** | Accumulation of amino acids | 0.00377 | SLC16A2, SLC17A7 | 2 |
| **Behavior** | Place aversion | 0.00377 | NPY, PDYN | 2 |
| **Cellular Movement,Nervous System Development and Function** | Migration of interneurons | 0.00391 | ATAT1, DLX2, RAC1 | 3 |
| **Embryonic Development,Nervous System Development and Function,Organ Development,Organ Morphology,Organismal Development,Tissue Development** | Morphogenesis of brain | 0.0049 | EMX1, L1CAM, MKKS, OTX1, SHANK3, ZNF335 | 6 |
| **Cell-To-Cell Signaling and Interaction,Nervous System Development and Function** | Synaptic transmission of cells | 0.00598 | ANKS1B, DOC2A, SH3GL1, SHANK3, SLC17A7, SYBU, UCHL1 | 7 |

| | | | | |
|---|---|---|---|---|
| **Embryonic Development,Nervous System Development and Function,Organ Development,Organismal Development,Tissue Development** | Formation of brain | 0.00738 | ATAT1, DCLK2, DLX2, DYNC2LI1, EMX1, FOXG1, HPRT1, ID2, L1CAM, LDB1 | 27 |
| **Amino Acid Metabolism,Endocrine System Development and Function,Molecular Transport,Small Molecule Biochemistry** | Quantity of L-triiodothyronine | 0.00912 | NPY, SLC16A2 | 2 |
| **Cell-To-Cell Signaling and Interaction,Nervous System Development and Function** | Synaptic transmission of superior cervical ganglion neurons | 0.00912 | DOC2A, SYBU | 2 |
| **Cellular Assembly and Organization,Cellular Compromise,Nervous System Development and Function** | Retraction of axons | 0.00912 | OTX1, PPP2CA | 2 |
| **Embryonic Development,Nervous System Development and Function,Organ Development,Organismal Development,Tissue Development** | Formation of forebrain | 0.00992 | DLX2, EMX1, FOXG1, HPRT1, ID2, L1CAM, MKKS, NEUROG2, NUMBL, OTX1 | 14 |
| **Cellular Movement** | Cell movement of neurons | 0.0179 | ATAT1, CSPG5, DLX2, FOXG1, L1CAM, MAP2K4, NEUROG2, NRCAM, RAC1, SRF | 10 |
| **Cellular Function and Maintenance** | Endocytosis | 0.0195 | ACTB, MERTK, PPP3CA, RAC1, SCAMP5 | 5 |
| **Nervous System Development and Function,Neurological Disease,Organ Morphology,Organismal Development,Organismal Injury and Abnormalities** | Abnormal morphology of midbrain-hindbrain boundary | 0.0208 | LDB1, OTX1 | 2 |
| **Lipid Metabolism,Molecular Transport,Small Molecule Biochemistry** | Accumulation of glycosphingolipid | 0.0208 | GBA, IDUA | 2 |
| **Amino Acid Metabolism,Drug Metabolism,Endocrine System Development and Function,Molecular Transport,Small Molecule Biochemistry** | Concentration of levothyroxine | 0.0208 | NPY, SLC16A2 | 2 |
| **Cell-To-Cell Signaling and Interaction,Nervous System Development and Function** | Synaptic transmission of hippocampal neurons | 0.0225 | ANKS1B, SHANK3, UCHL1 | 3 |
| **Cellular Movement,Nervous System Development and Function** | Migration of neurons | 0.025 | ATAT1, CSPG5, DLX2, FOXG1, L1CAM, MAP2K4, NEUROG2, RAC1, SRF | 9 |
| **Carbohydrate Metabolism,Drug Metabolism,Molecular Transport,Small Molecule Biochemistry** | Quantity of heparan sulfate | 0.0255 | IDUA | 1 |
| **Cell-To-Cell Signaling and Interaction,Nervous System Development and Function** | Activation of KNDy neurons | 0.0255 | PDYN | 1 |
| **Cellular Development** | Commitment of cortical precursor cells | 0.0255 | DLX2 | 1 |
| **Carbohydrate Metabolism,Lipid Metabolism,Small Molecule Biochemistry** | Hydrolysis of phosphatidylinositol polyphosphate | 0.0255 | EDN1 | 1 |
| **Neurological Disease** | Pathological denervation of | 0.0255 | UCHL1 | 1 |

| | | | | |
|---|---|---|---|---|
| | neuromuscular junctions | | | |
| **Lipid Metabolism,Molecular Transport,Small Molecule Biochemistry** | Accumulation of glucosylceramide | 0.0255 | GBA | 1 |
| **Cardiovascular Disease** | Congestion of vasculature | 0.0255 | PROS1 | 1 |
| **Auditory Disease,Neurological Disease** | Progressive high frequency hearing impairment | 0.0255 | ACTB | 1 |
| **Cell-mediated Immune Response,Cellular Movement,Hematological System Development and Function,Immune Cell Trafficking** | Cellular infiltration by CD4+ T-lymphocytes | 0.0255 | BSG | 1 |
| **Connective Tissue Development and Function,Organismal Development,Skeletal and Muscular System Development and Function,Tissue Development** | Size of cranium | 0.0255 | MYB | 1 |
| **Amino Acid Metabolism,Molecular Transport,Small Molecule Biochemistry** | Uptake of L-triiodothyronine | 0.0255 | SLC16A2 | 1 |
| **Embryonic Development,Nervous System Development and Function,Organ Development,Organismal Development,Tissue Development** | Formation of subgranular zone of dentate gyrus | 0.0255 | FOXG1 | 1 |
| **Organismal Functions** | Regulation of pituitary gland | 0.0255 | NPY | 1 |
| **Cellular Assembly and Organization,Cellular Function and Maintenance** | Endocytosis of plasma membrane | 0.0255 | RAC1 | 1 |
| **Cellular Growth and Proliferation,Nervous System Development and Function,Organ Development** | Proliferation of stellate cells | 0.0255 | EDN1 | 1 |
| **Cellular Movement,Nervous System Development and Function** | Delay in initiation of migration of cortical projection neurons | 0.0255 | RAC1 | 1 |
| **Cellular Compromise** | Acidification of cytosol | 0.0255 | CA4 | 1 |
| **Protein Synthesis** | Initiation of translation of protein | 0.0255 | AMPD2 | 1 |
| **Cellular Assembly and Organization,Cellular Function and Maintenance** | Quantity of peripheral lamellae | 0.0255 | RAC1 | 1 |
| **Cellular Development,Cellular Growth and Proliferation,Embryonic Development,Nervous System Development and Function,Organ Development,Organismal Development,Tissue Development** | Development of mesencephalic neurons | 0.0255 | NEUROG2 | 1 |
| **Cell-mediated Immune Response,Cell-To-Cell Signaling and Interaction,Cellular Movement,Hematological System Development and Function,Immune Cell Trafficking** | Sequestration of T lymphocytes | 0.0255 | HDC | 1 |
| **Cellular Assembly and Organization,Cellular Development,Embryonic Development,Nervous System Development and Function,Organ Development,Organismal Development,Tissue Development** | Maturation of olfactory glomeruli | 0.0255 | PPP3CA | 1 |
| **Cellular Movement,Nervous System Development and Function** | Migration of medial ganglionic eminence-derived interneurons | 0.0255 | ATAT1 | 1 |
| **Connective Tissue Disorders,Inflammatory Disease,Inflammatory Response,Organismal Injury and Abnormalities,Skeletal and Muscular Disorders** | Degeneration of lumbar intervertebral disc | 0.0255 | EDN1 | 1 |
| **Cellular Development,Cellular Growth and Proliferation,Nervous System Development and Function,Tissue Development** | Development of Cajal-Retzius neurons | 0.0255 | FOXG1 | 1 |
| **Lipid Metabolism,Small Molecule Biochemistry** | Incorporation of sphingomyelin | 0.0255 | GBA | 1 |
| **Carbohydrate Metabolism,Lipid Metabolism,Small Molecule Biochemistry** | Incorporation of phosphatidylcholine | 0.0255 | GBA | 1 |
| **Amino Acid Metabolism,Small Molecule Biochemistry** | Accumulation of levothyroxine | 0.0255 | SLC16A2 | 1 |
| **Organismal Functions** | Flicking of tail | 0.0255 | PDYN | 1 |
| **Cell-To-Cell Signaling and Interaction,Nervous System Development and Function** | Activation of superior cervical ganglion neurons | 0.0255 | SYBU | 1 |
| **Organismal Injury and Abnormalities,Reproductive System Disease** | Endometriosis | 0.0255 | UCHL1 | 1 |
| **Drug Metabolism,Molecular Transport** | Efflux of glutathione | 0.0255 | EDN1 | 1 |

| | | | | |
|---|---|---|---|---|
| **Cellular Development,Cellular Growth and Proliferation,Embryonic Development,Nervous System Development and Function,Organ Development,Organismal Development,Tissue Development** | Expansion of telencephalic progenitor cells | 0.0255 | FOXG1 | 1 |
| **Cell Morphology,Skeletal and Muscular System Development and Function** | Hypercontractility of vascular smooth muscle cells | 0.0255 | SRF | 1 |
| **Cell Morphology** | Contractility of stellate cells | 0.0255 | EDN1 | 1 |
| **Cardiovascular Disease,Developmental Disorder,Organismal Injury and Abnormalities** | Congenital heart block | 0.0255 | EDN1 | 1 |
| **Lipid Metabolism,Molecular Transport,Small Molecule Biochemistry** | Quantity of fat | 0.0255 | NPY | 1 |
| **Lipid Metabolism,Small Molecule Biochemistry** | Storage of fat | 0.0255 | NPY | 1 |
| **Cell Morphology** | Projection of thalamic cells | 0.0255 | NEUROG2 | 1 |
| **Amino Acid Metabolism,Small Molecule Biochemistry** | Accumulation of L-triiodothyronine | 0.0255 | SLC16A2 | 1 |
| **Cell Morphology,Cellular Assembly and Organization,Nervous System Development and Function,Tissue Morphology** | Size of synaptic vesicles pool | 0.0255 | SCAMP5 | 1 |
| **Lipid Metabolism,Molecular Transport,Small Molecule Biochemistry** | Concentration of eicosapentenoic acid | 0.0255 | ARNTL | 1 |
| **Neurological Disease** | Formation of myelin ovoid | 0.0255 | RAC1 | 1 |
| **Immunological Disease,Inflammatory Disease,Inflammatory Response,Neurological Disease,Organismal Injury and Abnormalities** | Symptomatic experimental autoimmune encephalomyelitis | 0.0255 | BSG | 1 |
| **Cell Morphology,Cellular Assembly and Organization,Nervous System Development and Function** | Projection of corticothalamic axons | 0.0255 | RAC1 | 1 |
| **Infectious Diseases** | Infection by human herpesvirus 2 | 0.0255 | NECTIN1 | 1 |
| **Cancer,Organismal Injury and Abnormalities** | Tumor burden | 0.0255 | MERTK | 1 |
| **Nervous System Development and Function,Tissue Development** | Formation of nerve fascicle | 0.0255 | L1CAM | 1 |
| **Lipid Metabolism,Small Molecule Biochemistry** | Incorporation of choline | 0.0255 | GBA | 1 |
| **Nervous System Development and Function,Organ Morphology,Organismal Development** | Size of cerebral ventricles | 0.0256 | L1CAM, MYB | 2 |
| **Nervous System Development and Function,Neurological Disease,Organ Morphology,Organismal Development,Organismal Injury and Abnormalities** | Abnormal morphology of cerebral aqueduct | 0.0256 | L1CAM, OTX1 | 2 |
| **Neurological Disease,Organismal Injury and Abnormalities** | Abnormality of substantia nigra | 0.0256 | HDC, SEPT5 | 2 |
| **Cell Morphology** | Projection | 0.0278 | L1CAM, NEUROG2, RAC1 | 3 |
| **Nervous System Development and Function** | Development of central nervous system | 0.0281 | ATAT1, DCLK2, DLX2, DYNC2LI1, EMX1, FOXG1, HPRT1, ID2, L1CAM, LDB1 | 30 |
| **Cell-To-Cell Signaling and Interaction,Nervous System Development and Function** | Synaptic transmission of nervous tissue | 0.0291 | ANKS1B, DOC2A, SH3GL1, SHANK3, SLC17A7, SYBU, UCHL1 | 7 |
| **Embryonic Development,Nervous System Development and Function,Organ Development,Organismal Development,Tissue Development** | Development of striatum | 0.0307 | DLX2, HPRT1, MKKS | 3 |

| | | | | |
|---|---|---|---|---|
| **Embryonic Development,Nervous System Development and Function,Organ Development,Organismal Development,Tissue Development** | Formation of anterior commissure | 0.0363 | RAC1, TG | 2 |
| **Cell Morphology** | Modification of neurons | 0.0363 | EDN1, NEUROG2 | 2 |
| **Cellular Movement** | Movement of growth cone | 0.0363 | ACTB, L1CAM | 2 |
| **Cellular Movement** | Migration of cells | 0.0399 | ATAT1, BSG, CCL4, CSPG5, DLX2, FAS, FOXG1, HDC, L1CAM, MAP2K4 | 13 |
| **Cell Death and Survival,Nervous System Development and Function** | Cell viability of granule cells | 0.0403 | FOXG1, L1CAM, NRCAM | 3 |
| **Cellular Movement** | Cell movement | 0.0451 | ATAT1, BSG, CCL4, CSPG5, DLX2, FAS, FOXG1, HDC, L1CAM, MAP2K4 | 14 |
| **Cellular Movement,Hematological System Development and Function,Immune Cell Trafficking** | Cell movement of mononuclear leukocytes | 0.0475 | BSG, CCL4, HDC | 3 |
| **Cell Morphology,Cellular Assembly and Organization** | Accumulation of vesicles | 0.0484 | SH3GL1, UCHL1 | 2 |

### Cluster 6

| Categories | Diseases or Functions Annotation | p-Value | Molecules | # Molecules |
|---|---|---|---|---|
| **Carbohydrate Metabolism,Molecular Transport,Small Molecule Biochemistry** | Accumulation of glycosaminoglycan | 0.00178 | HEXB, IDUA | 2 |
| **Gene Expression** | Expression of mRNA | 0.00435 | CRHR2, GCDH | 2 |
| **Gene Expression** | Expression of RNA | 0.0102 | CRHR2, GCDH, SRF | 3 |
| **Cancer,Hematological Disease,Immunological Disease,Neurological Disease,Organismal Injury and Abnormalities** | Primary central nervous system lymphoma | 0.0125 | FOXRED1, PRDM1 | 2 |
| **Carbohydrate Metabolism,Drug Metabolism,Molecular Transport,Small Molecule Biochemistry** | Quantity of heparan sulfate | 0.0175 | IDUA | 1 |
| **Organismal Development** | Size of animal | 0.0175 | NR3C1 | 1 |
| **Cellular Assembly and Organization,Nervous System Development and Function** | Fasciculation of parallel fiber | 0.0175 | NFIX | 1 |
| **Cell Death and Survival,Cellular Development,Nervous System Development and Function,Tissue Development,Tissue Morphology** | Regeneration of bipolar cells | 0.0175 | PRDM1 | 1 |
| **Nervous System Development and Function** | Development of dura mater | 0.0175 | NR3C1 | 1 |
| **Endocrine System Development and Function** | Hyperactivation of hypothalamus-pituitary-adrenal axis | 0.0175 | NR3C1 | 1 |
| **Cell Morphology,Skeletal and Muscular System Development and Function** | Hypercontractility of vascular smooth muscle cells | 0.0175 | SRF | 1 |
| **Cell Cycle** | Quiescence of neural stem cells | 0.0175 | NFIX | 1 |
| **Nervous System Development and Function,Neurological Disease,Organ Morphology,Organismal Development,Organismal Injury and Abnormalities** | Abnormal morphology of cerebellar hemisphere | 0.0175 | AHI1 | 1 |
| **Lipid Metabolism,Molecular Transport,Small Molecule Biochemistry** | Concentration of eicosapentenoic acid | 0.0175 | ARNTL | 1 |

| | | | | |
|---|---|---|---|---|
| **Cellular Compromise** | Acidification of cytosol | 0.0175 | CA4 | 1 |
| **Nervous System Development and Function,Organ Morphology,Tissue Morphology** | Quantity of rod bipolar cells | 0.0175 | PRDM1 | 1 |
| **Organismal Development** | Delay in growth of organism | 0.0175 | NR3C1 | 1 |
| **Auditory Disease,Cell Death and Survival,Cellular Compromise,Neurological Disease,Organismal Injury and Abnormalities,Tissue Morphology** | Neurodegeneration of hair cells | 0.0246 | ADGRV1, IDUA, SUN1 | 3 |
| **Cell Morphology,Nervous System Development and Function,Neurological Disease,Organ Morphology,Organismal Development,Organismal Injury and Abnormalities,Tissue Morphology** | Abnormal morphology of Purkinje cells | 0.0255 | HEXB, IDUA, SPARCL1, SUN1 | 4 |
| **Nervous System Development and Function,Organ Morphology,Organismal Development** | Mass of brain | 0.0281 | CTSF, MBD5, NFIX, SUN1 | 4 |
| **Nervous System Development and Function,Neurological Disease,Organ Morphology,Organismal Development,Organismal Injury and Abnormalities** | Abnormal morphology of cerebellar cortex | 0.0331 | AHI1, HEXB, IDUA, SPARCL1, SUN1 | 5 |
| **Cardiovascular Disease,Hematological Disease,Metabolic Disease** | Hyperlipidemia | 0.0346 | SH2B1 | 1 |
| **Cellular Development,Cellular Growth and Proliferation,Embryonic Development,Nervous System Development and Function,Organ Development,Organismal Development,Tissue Development** | Development of spiny stellate neurons | 0.0346 | SLC17A7 | 1 |
| **Infectious Diseases,Neurological Disease,Organismal Injury and Abnormalities** | Infection of cortical neurons | 0.0346 | SEMA7A | 1 |
| **Nervous System Development and Function,Neurological Disease,Organ Morphology,Organismal Development,Organismal Injury and Abnormalities** | Abnormal morphology of oculomotor nucleus | 0.0346 | OTX1 | 1 |
| **Nervous System Development and Function,Neurological Disease,Organ Morphology,Organismal Development,Organismal Injury and Abnormalities** | Abnormal morphology of enlarged cerebellum | 0.0346 | OTX1 | 1 |
| **Cell-To-Cell Signaling and Interaction,Nervous System Development and Function** | Synaptic transmission of thalamocortical neurons | 0.0346 | SLC17A7 | 1 |
| **Endocrine System Disorders** | Leptin resistance | 0.0346 | SH2B1 | 1 |
| **Cell Morphology,Cellular Assembly and Organization,Cellular Function and Maintenance,Nervous System Development and Function** | Extension of parallel fiber | 0.0346 | NFIX | 1 |
| **Endocrine System Development and Function,Lipid Metabolism,Molecular Transport,Small Molecule Biochemistry** | Release of corticosterone | 0.0346 | NR3C1 | 1 |
| **Nervous System Development and Function** | Development of ventral nerve cord | 0.0346 | OTX1 | 1 |
| **Cell-To-Cell Signaling and Interaction** | Gap junctional intercellular communication of neuronal progenitor cells | 0.0346 | NR3C1 | 1 |
| **Cellular Movement,Nervous System Development and Function** | Distribution of interneurons | 0.0346 | DCK | 1 |
| **Lipid Metabolism,Molecular Transport,Small Molecule Biochemistry** | Accumulation of ganglioside GM3 | 0.0346 | IDUA | 1 |
| **Embryonic Development,Nervous System Development and Function,Organ Development,Organismal Development,Tissue Development** | Formation of pretectum | 0.0346 | OTX1 | 1 |
| **Carbohydrate Metabolism,Drug Metabolism,Molecular Transport,Small Molecule Biochemistry** | Accumulation of hyaluronic acid | 0.0346 | HEXB | 1 |
| **Metabolic Disease** | Hyperleptinemia | 0.0346 | SH2B1 | 1 |

**Cluster 7**

| Categories | Diseases or Functions Annotation | p-Value | Molecules | # Molecules |
|---|---|---|---|---|
| **Cellular Movement,Nervous System Development and Function** | Delay in migration of neurons | 0.00238 | CDKL5, RAC1 | 2 |
| **Cellular Development,Cellular Growth and Proliferation,Nervous System Development and Function,Tissue Development** | Development of bipolar cells | 0.00392 | FOXG1, PRDM1 | 2 |
| **Infectious Diseases,Neurological Disease,Organismal Injury and Abnormalities** | Infection of brain | 0.0058 | NECTIN1, SEMA7A | 2 |
| **Cellular Development,Cellular Growth and Proliferation,Nervous System Development and Function,Tissue Development** | Growth of dendritic spines | 0.0058 | CDKL5, ERBB4 | 2 |
| **Cell-To-Cell Signaling and Interaction** | Adhesion of central nervous system cells | 0.00801 | ERBB4, NECTIN1 | 2 |
| **Lipid Metabolism,Molecular Transport,Small Molecule Biochemistry** | Concentration of arachidonic acid | 0.0134 | ARNTL, KNG1 | 2 |
| **Cell-To-Cell Signaling and Interaction,Cellular Assembly and Organization,Cellular Development,Cellular Function and Maintenance,Cellular Growth and Proliferation,Nervous System Development and Function,Tissue Development** | Formation of excitatory synapses | 0.0165 | DLG3, ERBB4 | 2 |
| **Cellular Assembly and Organization,Nervous System Development and Function** | Fasciculation of parallel fiber | 0.0202 | NFIX | 1 |
| **Cellular Compromise,Neurological Disease,Organismal Injury and Abnormalities** | Atrophy of motor axons | 0.0202 | NEFH | 1 |
| **Embryonic Development,Nervous System Development and Function,Organ Development,Organismal Development,Tissue Development** | Formation of subgranular zone of dentate gyrus | 0.0202 | FOXG1 | 1 |
| **Cellular Assembly and Organization,Cellular Function and Maintenance** | Endocytosis of plasma membrane | 0.0202 | RAC1 | 1 |
| **Cell-To-Cell Signaling and Interaction,Nervous System Development and Function** | Conduction of optic nerve | 0.0202 | ERBB4 | 1 |
| **Carbohydrate Metabolism,Drug Metabolism,Small Molecule Biochemistry** | Synthesis of heparan sulfate | 0.0202 | EXTL2 | 1 |
| **Nervous System Development and Function,Organ Morphology,Tissue Morphology** | Quantity of rod bipolar cells | 0.0202 | PRDM1 | 1 |
| **Carbohydrate Metabolism,Lipid Metabolism,Small Molecule Biochemistry** | Incorporation of phosphatidylcholine | 0.0202 | GBA | 1 |
| **Cell Death and Survival,Cellular Development,Nervous System Development and Function,Tissue Development,Tissue Morphology** | Regeneration of bipolar cells | 0.0202 | PRDM1 | 1 |
| **Ophthalmic Disease,Organ Development,Organismal Injury and Abnormalities,Visual System Development and Function** | Formation of cataract | 0.0202 | NF2 | 1 |
| **Cellular Compromise,Neurological Disease,Organismal Injury and Abnormalities** | Dysfunction of motor neurons | 0.0202 | NEFH | 1 |
| **Inflammatory Response** | Inflammatory response of cerebral cortex | 0.0202 | KNG1 | 1 |
| **Neurological Disease** | Misinnervation | 0.0202 | ERBB4 | 1 |
| **Cellular Development,Cellular Growth and Proliferation,Embryonic Development,Nervous System Development and Function,Organ Development,Organismal Development,Tissue Development** | Expansion of telencephalic progenitor cells | 0.0202 | FOXG1 | 1 |
| **Neurological Disease** | Formation of myelin ovoid | 0.0202 | RAC1 | 1 |
| **Cell Morphology,Cellular Assembly and Organization,Nervous System Development and Function** | Projection of corticothalamic axons | 0.0202 | RAC1 | 1 |
| **Cell Death and Survival,Nervous System Development and Function** | Survival of motor axons | 0.0202 | NEFH | 1 |
| **Cellular Movement,Embryonic Development** | Migration of cranial neural crest cells | 0.0202 | ERBB4 | 1 |
| **Cancer,Neurological Disease,Organismal Injury and Abnormalities** | Fibrous meningioma | 0.0202 | NF2 | 1 |

| | | | | |
|---|---|---|---|---|
| **Lipid Metabolism,Small Molecule Biochemistry** | Incorporation of choline | 0.0202 | GBA | 1 |
| **Cell-To-Cell Signaling and Interaction,Nervous System Development and Function** | Sensitization of trigeminovascular neuron | 0.0202 | KNG1 | 1 |
| **Cell-To-Cell Signaling and Interaction,Nervous System Development and Function** | Activation of excitatory synapses | 0.0202 | CDKL5 | 1 |
| **Lipid Metabolism,Molecular Transport,Small Molecule Biochemistry** | Accumulation of glucosylceramide | 0.0202 | GBA | 1 |
| **Cellular Growth and Proliferation,Nervous System Development and Function,Organ Development** | Growth of arachnoidal cells | 0.0202 | NF2 | 1 |
| **Cellular Movement,Nervous System Development and Function** | Delay in initiation of migration of cortical projection neurons | 0.0202 | RAC1 | 1 |
| **Cell Cycle** | Quiescence of neural stem cells | 0.0202 | NFIX | 1 |
| **Cell Cycle,Cell-To-Cell Signaling and Interaction,Cellular Growth and Proliferation** | Contact growth inhibition | 0.0202 | NF2 | 1 |
| **Cellular Assembly and Organization,Cellular Function and Maintenance** | Quantity of peripheral lamellae | 0.0202 | RAC1 | 1 |
| **Cancer,Neurological Disease,Organismal Injury and Abnormalities** | Tumorigenesis of schwannoma | 0.0202 | NF2 | 1 |
| **Cellular Assembly and Organization,Cellular Development,Embryonic Development,Nervous System Development and Function,Organ Development,Organismal Development,Tissue Development** | Maturation of olfactory glomeruli | 0.0202 | PPP3CA | 1 |
| **Carbohydrate Metabolism,Connective Tissue Development and Function,Skeletal and Muscular System Development and Function,Small Molecule Biochemistry,Tissue Development** | Synthesis of chondroitin sulfate | 0.0202 | EXTL2 | 1 |
| **Cellular Development,Cellular Growth and Proliferation,Nervous System Development and Function,Tissue Development** | Development of Cajal-Retzius neurons | 0.0202 | FOXG1 | 1 |
| **Lipid Metabolism,Small Molecule Biochemistry** | Incorporation of sphingomyelin | 0.0202 | GBA | 1 |
| **Cellular Assembly and Organization,Nervous System Development and Function,Tissue Morphology** | Density of neurofilaments | 0.0202 | NEFH | 1 |
| **Cancer,Dermatological Diseases and Conditions,Developmental Disorder,Hereditary Disorder,Neurological Disease,Organismal Injury and Abnormalities** | Neurofibromatosis 2 | 0.0202 | NF2 | 1 |
| **Cell-To-Cell Signaling and Interaction,Nervous System Development and Function** | Activation of superior cervical ganglion neurons | 0.0202 | SYBU | 1 |
| **Cell Morphology,Cellular Assembly and Organization,Cellular Function and Maintenance** | Reorganization of actin filaments | 0.0202 | DLG3 | 1 |
| **Cardiovascular System Development and Function,Nervous System Development and Function,Organ Morphology,Organismal Development,Tissue Morphology** | Diameter of basilar artery | 0.0202 | KNG1 | 1 |
| **Cell-To-Cell Signaling and Interaction,Nervous System Development and Function** | Firing of trigeminovascular neuron | 0.0202 | KNG1 | 1 |
| **Cell Morphology,Cellular Assembly and Organization,Nervous System Development and Function,Tissue Morphology** | Size of synaptic vesicles pool | 0.0202 | SCAMP5 | 1 |
| **Lipid Metabolism,Molecular Transport,Small Molecule Biochemistry** | Concentration of eicosapentenoic acid | 0.0202 | ARNTL | 1 |
| **Infectious Diseases** | Infection by human herpesvirus 2 | 0.0202 | NECTIN1 | 1 |
| **Cellular Growth and Proliferation** | Cytostasis of neuroglia | 0.0202 | NF2 | 1 |
| **Nervous System Development and Function,Neurological Disease,Organ Morphology,Organismal Development,Organismal Injury and Abnormalities** | Abnormal morphology of barrel cortex | 0.0275 | ASPM, DLG3 | 2 |
| **Carbohydrate Metabolism** | Accumulation of carbohydrate | 0.0275 | HEXB, KNG1 | 2 |

| | | | | |
|---|---|---|---|---|
| **Cellular Development,Cellular Growth and Proliferation,Nervous System Development and Function,Tissue Development** | Myelination of Schwann cells | 0.0275 | NF2, RAC1 | 2 |
| **Neurological Disease,Psychological Disorders** | Schizophrenia | 0.0294 | DLG3, FAM3C, HPRT1, IPCEF1, MAP10, NELL2, OXCT1, SLC17A7, SYT7 | 9 |
| **Nervous System Development and Function,Neurological Disease,Organ Morphology,Organismal Development,Organismal Injury and Abnormalities** | Abnormal morphology of third cerebral ventricle | 0.0309 | MKKS, NFIX, OTX1 | 3 |
| **Cell Morphology,Nervous System Development and Function,Neurological Disease,Tissue Morphology** | Abnormal morphology of neurites | 0.0319 | DLG3, ERBB4, NEFH, NRCAM, NRN1, PPP3CA, SEMA5B, SLC17A7 | 8 |
| **Embryonic Development,Nervous System Development and Function,Organ Development,Organismal Development,Tissue Development** | Development of cerebral cortex | 0.0337 | ASPM, ERBB4, FOXG1, MKKS, NF2, OTX1, RAC1, SH2B1 | 8 |
| **Embryonic Development,Nervous System Development and Function,Organ Development,Organismal Development,Tissue Development** | Formation of hippocampus | 0.0339 | FOXG1, MKKS, NF2, OTX1, RAC1, SH2B1 | 6 |
| **Cellular Function and Maintenance** | Endocytosis | 0.0348 | KIAA1107, PPP3CA, RAC1, SCAMP5 | 4 |
| **Cellular Assembly and Organization,Cellular Function and Maintenance,Cellular Movement,Nervous System Development and Function** | Endocytosis of synaptic vesicles | 0.0384 | KIAA1107, PPP3CA, SCAMP5 | 3 |
| **Cellular Development,Cellular Growth and Proliferation,Embryonic Development,Nervous System Development and Function,Organ Development,Organismal Development,Tissue Development** | Development of spiny stellate neurons | 0.0401 | SLC17A7 | 1 |
| **Nervous System Development and Function,Neurological Disease,Organ Morphology,Organismal Development,Organismal Injury and Abnormalities** | Abnormal morphology of oculomotor nucleus | 0.0401 | OTX1 | 1 |
| **Endocrine System Disorders** | Leptin resistance | 0.0401 | SH2B1 | 1 |
| **Cell Morphology,Cell-To-Cell Signaling and Interaction,Cellular Assembly and Organization** | Length of gap junctions | 0.0401 | NRCAM | 1 |
| **Nervous System Development and Function,Organ Morphology,Organismal Development** | Thickness of granule cell layer | 0.0401 | NRCAM | 1 |
| **Cellular Assembly and Organization,Cellular Compromise** | Formation of neurofilament inclusion | 0.0401 | NEFH | 1 |
| **Cell Morphology,Cellular Assembly and Organization,Cellular Function and Maintenance,Nervous System Development and Function** | Extension of parallel fiber | 0.0401 | NFIX | 1 |
| **Carbohydrate Metabolism,Drug Metabolism,Molecular Transport,Small Molecule Biochemistry** | Accumulation of hyaluronic acid | 0.0401 | HEXB | 1 |
| **Cellular Development,Cellular Growth and Proliferation,Nervous System Development and Function,Tissue Development** | Development of striatal interneurons | 0.0401 | LHX6 | 1 |
| **Cardiovascular Disease,Hematological Disease,Organismal Injury and Abnormalities** | Thrombus | 0.0401 | KNG1 | 1 |

| | | | | |
|---|---|---|---|---|
| **Metabolic Disease** | Hyperleptinemia | 0.0401 | SH2B1 | 1 |
| **Infectious Diseases,Neurological Disease,Organismal Injury and Abnormalities** | Infection of cortical neurons | 0.0401 | SEMA7A | 1 |
| **Cell-To-Cell Signaling and Interaction,Nervous System Development and Function** | Synaptic transmission of thalamocortical neurons | 0.0401 | SLC17A7 | 1 |
| **Nervous System Development and Function** | Development of ventral nerve cord | 0.0401 | OTX1 | 1 |
| **Embryonic Development,Nervous System Development and Function,Organ Development,Organismal Development,Tissue Development** | Formation of pretectum | 0.0401 | OTX1 | 1 |
| **Cell-To-Cell Signaling and Interaction,Cellular Assembly and Organization,Cellular Function and Maintenance** | Formation of spot-like adhesion site | 0.0401 | NECTIN1 | 1 |
| **Hereditary Disorder,Neurological Disease,Organismal Injury and Abnormalities** | Small fiber neuropathy | 0.0401 | ERBB4 | 1 |
| **Cancer,Neurological Disease,Organismal Injury and Abnormalities** | Transitional meningioma | 0.0401 | NF2 | 1 |
| **Cell-To-Cell Signaling and Interaction,Nervous System Development and Function** | Synaptic depression of superior cervical ganglion neurons | 0.0401 | SYBU | 1 |
| **Cellular Development,Cellular Growth and Proliferation,Nervous System Development and Function,Tissue Development** | Outgrowth of sensory neurons | 0.0401 | CLU | 1 |
| **Cardiovascular Disease,Hematological Disease,Metabolic Disease** | Hyperlipidemia | 0.0401 | SH2B1 | 1 |
| **Nervous System Development and Function,Organ Morphology,Tissue Morphology** | Quantity of mossy fiber cells | 0.0401 | RAC1 | 1 |
| **Cell-To-Cell Signaling and Interaction,Nervous System Development and Function** | Adhesion of astrocytes | 0.0401 | ERBB4 | 1 |
| **Lipid Metabolism,Small Molecule Biochemistry** | Liberation of arachidonic acid | 0.0401 | KNG1 | 1 |
| **Neurological Disease,Organismal Injury and Abnormalities** | Injury of spinal cord | 0.0401 | CLU | 1 |
| **Cell Cycle,Embryonic Development,Nervous System Development and Function** | Cell division of neuroblasts | 0.0401 | ASPM | 1 |
| **Cell-To-Cell Signaling and Interaction,Nervous System Development and Function** | Action potential of superior cervical ganglion neurons | 0.0401 | KNG1 | 1 |
| **Lipid Metabolism,Small Molecule Biochemistry** | Liberation of stearic acid | 0.0401 | KNG1 | 1 |
| **Nervous System Development and Function,Neurological Disease,Organ Morphology,Organismal Development,Organismal Injury and Abnormalities** | Abnormal morphology of enlarged cerebellum | 0.0401 | OTX1 | 1 |
| **Cellular Development,Cellular Growth and Proliferation,Nervous System Development and Function,Tissue Development** | Generation of Cajal-Retzius neurons | 0.0401 | FOXG1 | 1 |
| **Cellular Assembly and Organization,Nervous System Development and Function** | Delay in sorting of axons | 0.0401 | RAC1 | 1 |
| **Embryonic Development,Organismal Development,Tissue Development** | Formation of preplate | 0.0401 | FOXG1 | 1 |
| **Cell Morphology,Cellular Assembly and Organization,Nervous System Development and Function** | Extension of leading processes | 0.0401 | RAC1 | 1 |
| **Cancer,Neurological Disease,Organismal Injury and Abnormalities** | Malignant meningioma | 0.0401 | NF2 | 1 |
| **Cancer,Neurological Disease,Organismal Injury and Abnormalities** | Astrocytoma in brainstem | 0.0406 | GBF1, THEMIS | 2 |
| **Cellular Assembly and Organization,Cellular Function and Maintenance** | Organization of filaments | 0.0406 | DLG3, NEFH | 2 |
| **Cancer,Neurological Disease,Organismal Injury and Abnormalities** | Brain astrocytoma | 0.0483 | ADAMTS8, AK5, ANKRD30B, ASPM, ATR, BCL11A, CACNG3, CDKL5, COBL, DBNDD1 | 16 |

| | | | | |
|---|---|---|---|---|
| **Neurological Disease,Psychological Disorders** | Bipolar disorder | 0.0497 | COX5A, DLG3, UQCRC2 | 3 |

### Cluster 8

| Categories | Diseases or Functions Annotation | p-Value | Molecules | # Molecules |
|---|---|---|---|---|
| **Cellular Development,Cellular Growth and Proliferation,Nervous System Development and Function,Tissue Development** | Development of bipolar cells | 0.00125 | FOXG1, PRDM1 | 2 |
| **Organismal Development** | Mass of organism | 0.00421 | CRHR1, NR3C1, PTPN2 | 3 |
| **Nervous System Development and Function,Organ Morphology,Organismal Development** | Mass of brain | 0.00664 | ASPM, CTSF, MBD5, NFIX | 4 |
| **Carbohydrate Metabolism,Drug Metabolism,Molecular Transport,Small Molecule Biochemistry** | Quantity of heparan sulfate | 0.0113 | IDUA | 1 |
| **Cellular Development,Cellular Growth and Proliferation,Nervous System Development and Function,Tissue Development** | Development of Cajal-Retzius neurons | 0.0113 | FOXG1 | 1 |
| **Organismal Development** | Size of animal | 0.0113 | NR3C1 | 1 |
| **Cell-To-Cell Signaling and Interaction,Nervous System Development and Function** | Short-term potentiation of neurons | 0.0113 | CRHR1 | 1 |
| **Cellular Assembly and Organization,Nervous System Development and Function** | Fasciculation of parallel fiber | 0.0113 | NFIX | 1 |
| **Visual System Development and Function** | Light sensitivity of rod bipolar cells | 0.0113 | PDC | 1 |
| **Cell Death and Survival,Cellular Development,Nervous System Development and Function,Tissue Development,Tissue Morphology** | Regeneration of bipolar cells | 0.0113 | PRDM1 | 1 |
| **Nervous System Development and Function** | Development of dura mater | 0.0113 | NR3C1 | 1 |
| **Cellular Development,Cellular Growth and Proliferation,Embryonic Development,Nervous System Development and Function,Organ Development,Organismal Development,Tissue Development** | Expansion of telencephalic progenitor cells | 0.0113 | FOXG1 | 1 |
| **Endocrine System Development and Function** | Hyperactivation of hypothalamus-pituitary-adrenal axis | 0.0113 | NR3C1 | 1 |
| **Embryonic Development,Nervous System Development and Function,Organ Development,Organismal Development,Tissue Development** | Formation of subgranular zone of dentate gyrus | 0.0113 | FOXG1 | 1 |
| **Cell Cycle** | Quiescence of neural stem cells | 0.0113 | NFIX | 1 |
| **Cellular Compromise** | Acidification of cytosol | 0.0113 | CA4 | 1 |
| **Nervous System Development and Function,Organ Morphology,Tissue Morphology** | Quantity of rod bipolar cells | 0.0113 | PRDM1 | 1 |
| **Organismal Development** | Delay in growth of organism | 0.0113 | NR3C1 | 1 |
| **Cellular Movement,Hematological System Development and Function,Immune Cell Trafficking,Inflammatory Response** | Cellular infiltration of dendritic cells | 0.0113 | IL11RA | 1 |
| **Cell Morphology,Cellular Assembly and Organization,Cellular Development,Cellular Function and Maintenance,Cellular Growth and Proliferation,Nervous System Development and Function,Organismal Development,Tissue Development** | Delay in initiation of remyelination of axons | 0.0113 | IL11RA | 1 |
| **Cell Death and Survival** | Quantity of apoptotic cells | 0.0155 | LGALS3, MYC | 2 |
| **Cell Death and Survival,Cellular Development,Cellular Function and Maintenance,Cellular Growth and Proliferation,Embryonic Development,Organismal Development,Tissue Development** | Self-renewal of neural stem cells | 0.0193 | FOXG1, MYC | 2 |

| | | | | |
|---|---|---|---|---|
| **Cardiovascular Disease,Hematological Disease,Metabolic Disease** | Hyperlipidemia | 0.0226 | SH2B1 | 1 |
| **Infectious Diseases,Neurological Disease,Organismal Injury and Abnormalities** | Infection of cortical neurons | 0.0226 | SEMA7A | 1 |
| **Cellular Development,Cellular Growth and Proliferation,Nervous System Development and Function,Tissue Development** | Generation of Cajal-Retzius neurons | 0.0226 | FOXG1 | 1 |
| **Endocrine System Disorders** | Leptin resistance | 0.0226 | SH2B1 | 1 |
| **Embryonic Development,Organismal Development,Tissue Development** | Formation of preplate | 0.0226 | FOXG1 | 1 |
| **Cell Cycle,Embryonic Development,Nervous System Development and Function** | Cell division of neuroblasts | 0.0226 | ASPM | 1 |
| **Cell Morphology,Cellular Assembly and Organization,Cellular Function and Maintenance,Nervous System Development and Function** | Extension of parallel fiber | 0.0226 | NFIX | 1 |
| **Endocrine System Development and Function,Lipid Metabolism,Molecular Transport,Small Molecule Biochemistry** | Release of corticosterone | 0.0226 | NR3C1 | 1 |
| **Cell-To-Cell Signaling and Interaction** | Gap junctional intercellular communication of neuronal progenitor cells | 0.0226 | NR3C1 | 1 |
| **Lipid Metabolism,Molecular Transport,Small Molecule Biochemistry** | Accumulation of ganglioside GM3 | 0.0226 | IDUA | 1 |
| **Cardiovascular Disease,Organismal Injury and Abnormalities,Tissue Morphology** | Size of ischemic lesion | 0.0226 | LGALS3 | 1 |
| **Metabolic Disease** | Hyperleptinemia | 0.0226 | SH2B1 | 1 |
| **Nutritional Disease,Psychological Disorders** | Eating Disorders | 0.0233 | IL11RA, SH2B1 | 2 |
| **Nervous System Development and Function** | Remyelination | 0.03 | ADAM19, IL11RA | 2 |
| **Nervous System Development and Function** | Remyelination of sciatic nerve | 0.0337 | ADAM19 | 1 |
| **Cell Morphology,Cellular Assembly and Organization,Nervous System Development and Function** | Orientation of dendrites | 0.0337 | SEMA7A | 1 |
| **Cardiovascular System Development and Function,Cell Morphology,Cellular Development,Organismal Development,Tissue Development** | Tubulation of microvascular endothelial cells | 0.0337 | PLXDC1 | 1 |
| **Embryonic Development,Nervous System Development and Function,Organ Development,Organismal Development,Tissue Development** | Formation of brain septum | 0.0337 | ADAM19 | 1 |
| **Cell Death and Survival,Nervous System Development and Function,Tissue Morphology** | Quantity of apoptotic neurons | 0.0446 | LGALS3 | 1 |
| **Cellular Assembly and Organization,Cellular Function and Maintenance,Cellular Movement,Nervous System Development and Function** | Axonal transport of vesicles | 0.0446 | RAB33A | 1 |
| **Organismal Injury and Abnormalities,Tissue Morphology** | Area of lesion | 0.0446 | IL11RA | 1 |
| **Carbohydrate Metabolism,Molecular Transport,Small Molecule Biochemistry** | Accumulation of glycosaminoglycan | 0.0446 | IDUA | 1 |
| **Inflammatory Disease,Inflammatory Response,Neurological Disease,Organismal Injury and Abnormalities** | HIV-associated encephalitis | 0.0446 | MTDH | 1 |
| **Cancer,Organismal Injury and Abnormalities,Tissue Morphology,Tumor Morphology** | Morphology of malignant tumor | 0.0446 | MYC | 1 |
| **Endocrine System Disorders,Hematological Disease,Metabolic Disease** | Hyperglycemia | 0.0446 | SH2B1 | 1 |

### Cluster 9

| Categories | Diseases or Functions Annotation | p-Value | Molecules | # Molecules |
|---|---|---|---|---|
| **Embryonic Development,Nervous System Development and Function,Organ Development,Organ Morphology,Organismal Development,Tissue Development** | Morphogenesis of diencephalon | 0.000583 | OTX1, SOX2 | 2 |

| | | | | |
|---|---|---|---|---|
| **Cellular Movement,Hematological System Development and Function,Immune Cell Trafficking,Inflammatory Response** | Cell movement of microglia | 0.00144 | BSG, CX3CL1 | 2 |
| **Cell Morphology,Cellular Development,Connective Tissue Development and Function** | Conversion of induced neural precursor cells | 0.002 | FOXG1, SOX2 | 2 |
| **Cellular Movement,Hematological System Development and Function,Immune Cell Trafficking** | Cell movement of leukocytes | 0.00279 | BSG, CCL4, CX3CL1, FAS | 4 |
| **Embryonic Development,Nervous System Development and Function,Organ Development,Organismal Development,Tissue Development** | Development of diencephalon | 0.00353 | MYH10, OTX1, SOX2 | 3 |
| **Nervous System Development and Function,Neurological Disease,Organ Morphology,Organismal Development,Organismal Injury and Abnormalities** | Abnormal morphology of cerebral aqueduct | 0.0042 | MYH10, OTX1 | 2 |
| **Nervous System Development and Function,Neurological Disease,Organ Morphology,Organismal Development,Organismal Injury and Abnormalities** | Abnormal morphology of third cerebral ventricle | 0.0046 | MKKS, MYH10, OTX1 | 3 |
| **Cellular Movement,Nervous System Development and Function** | Migration of granule cells | 0.00631 | CX3CL1, FOXG1, MYH10 | 3 |
| **Cellular Movement** | Cell movement of myeloid cells | 0.00678 | BSG, CCL4, CX3CL1 | 3 |
| **Cellular Movement,Hematological System Development and Function,Immune Cell Trafficking,Inflammatory Response** | Cell movement of phagocytes | 0.00678 | BSG, CCL4, CX3CL1 | 3 |
| **Neurological Disease,Skeletal and Muscular Disorders** | Neuromuscular disease | 0.00778 | ACTN2, BSG, CCL4, FOXG1, LAMB1, NR4A1, PPP3CA, RPL31, SOX2, SPARCL1 | 10 |
| **Cellular Growth and Proliferation,Nervous System Development and Function,Organ Development** | Proliferation of brain cells | 0.00792 | EDN1, FOXG1, ID2, LGI1, SOX2 | 5 |
| **Neurological Disease,Organismal Injury and Abnormalities,Psychological Disorders** | Disorder of basal ganglia | 0.009 | ACTN2, CCL4, FAS, FOXG1, LAMB1, NR4A1, PPP3CA, RPL31, SOX2, SPARCL1 | 10 |
| **Cell Morphology,Nervous System Development and Function,Neurological Disease,Tissue Morphology** | Abnormal morphology of mossy fibers | 0.00949 | LGI1, PPP3CA | 2 |
| **Cell-mediated Immune Response,Cellular Movement,Hematological System Development and Function,Immune Cell Trafficking** | Cellular infiltration by CD4+ T-lymphocytes | 0.00996 | BSG | 1 |
| **Embryonic Development,Nervous System Development and Function,Organ Development,Organismal Development,Tissue Development** | Formation of subgranular zone of dentate gyrus | 0.00996 | FOXG1 | 1 |
| **Cellular Growth and Proliferation,Nervous System Development and Function,Organ Development** | Proliferation of stellate cells | 0.00996 | EDN1 | 1 |
| **Nervous System Development and Function** | Mechanical nociception | 0.00996 | CX3CL1 | 1 |
| **Cellular Development,Cellular Growth and Proliferation,Embryonic Development,Nervous System Development and Function,Organ Development,Organismal Development,Tissue Development** | Expansion of telencephalic progenitor cells | 0.00996 | FOXG1 | 1 |
| **Cell Morphology** | Contractility of stellate cells | 0.00996 | EDN1 | 1 |

| | | | | |
|---|---|---|---|---|
| **Auditory and Vestibular System Development and Function,Cellular Development,Cellular Growth and Proliferation,Embryonic Development,Nervous System Development and Function,Organ Development,Organismal Development,Tissue Development** | Development of spiral ganglion cells | 0.00996 | SOX2 | 1 |
| **Cellular Assembly and Organization,Cellular Function and Maintenance** | Stabilization of lamellipodia | 0.00996 | MYH10 | 1 |
| **Carbohydrate Metabolism,Lipid Metabolism,Small Molecule Biochemistry** | Hydrolysis of phosphatidylinositol polyphosphate | 0.00996 | EDN1 | 1 |
| **Cellular Assembly and Organization,Cellular Development,Embryonic Development,Nervous System Development and Function,Organ Development,Organismal Development,Tissue Development** | Maturation of olfactory glomeruli | 0.00996 | PPP3CA | 1 |
| **Connective Tissue Disorders,Inflammatory Disease,Inflammatory Response,Organismal Injury and Abnormalities,Skeletal and Muscular Disorders** | Degeneration of lumbar intervertebral disc | 0.00996 | EDN1 | 1 |
| **Cellular Development,Cellular Growth and Proliferation,Nervous System Development and Function,Tissue Development** | Development of Cajal-Retzius neurons | 0.00996 | FOXG1 | 1 |
| **Developmental Disorder,Hereditary Disorder,Metabolic Disease,Neurological Disease,Organismal Injury and Abnormalities** | Mitochondrial complex I deficiency of brain | 0.00996 | NDUFS6 | 1 |
| **Drug Metabolism,Molecular Transport** | Efflux of glutathione | 0.00996 | EDN1 | 1 |
| **Cardiovascular Disease,Developmental Disorder,Organismal Injury and Abnormalities** | Congenital heart block | 0.00996 | EDN1 | 1 |
| **Lipid Metabolism,Molecular Transport,Small Molecule Biochemistry** | Concentration of eicosapentenoic acid | 0.00996 | ARNTL | 1 |
| **Immunological Disease,Inflammatory Disease,Inflammatory Response,Neurological Disease,Organismal Injury and Abnormalities** | Symptomatic experimental autoimmune encephalomyelitis | 0.00996 | BSG | 1 |
| **Neurological Disease** | Movement Disorders | 0.0102 | ACTN2, CCL4, FAS, FOXG1, LAMB1, NR4A1, PPP3CA, RPL31, SOX2, SPARCL1 | 10 |
| **Cell Signaling,Molecular Transport,Vitamin and Mineral Metabolism** | Quantity of Ca2+ | 0.0106 | CCL4, CX3CL1, EDN1, VGF | 4 |
| **Cell Death and Survival,Cellular Development,Cellular Function and Maintenance,Cellular Growth and Proliferation,Embryonic Development,Organismal Development,Tissue Development** | Self-renewal of neural stem cells | 0.0151 | FOXG1, SOX2 | 2 |
| **Cell Morphology,Nervous System Development and Function,Neurological Disease,Tissue Morphology** | Abnormal morphology of bipolar cells | 0.0166 | MYH10, SEMA5B | 2 |
| **Cellular Movement,Hematological System Development and Function,Immune Cell Trafficking** | Cellular infiltration by mononuclear leukocytes | 0.0166 | BSG, CCL4 | 2 |
| **Embryonic Development,Nervous System Development and Function,Organ Development,Organismal Development,Tissue Development** | Formation of forebrain | 0.0186 | FOXG1, ID2, MKKS, MYH10, OTX1, PPP3CA, SOX2 | 7 |
| **Nervous System Development and Function,Neurological Disease,Organ Morphology,Organismal Development,Organismal Injury and Abnormalities** | Abnormal morphology of cerebellar cortex | 0.0188 | HEXB, IL1RAPL1, MYH10, SPARCL1 | 4 |
| **Organismal Functions** | Recovery of mice | 0.0198 | FAS | 1 |
| **Nervous System Development and Function,Neurological Disease,Organ Morphology,Organismal** | Abnormal morphology of oculomotor nucleus | 0.0198 | OTX1 | 1 |

| | | | | |
|---|---|---|---|---|
| **Development,Organismal Injury and Abnormalities** | | | | |
| **Behavior** | Hyperlocomotion | 0.0198 | VGF | 1 |
| **Cellular Assembly and Organization,Cellular Function and Maintenance** | Penetration of microtubules | 0.0198 | MYH10 | 1 |
| **Cellular Movement,Nervous System Development and Function** | Migration of hippocampal neurons | 0.0198 | CX3CL1 | 1 |
| **Carbohydrate Metabolism,Drug Metabolism,Molecular Transport,Small Molecule Biochemistry** | Accumulation of hyaluronic acid | 0.0198 | HEXB | 1 |
| **Cell Morphology,Cell-To-Cell Signaling and Interaction,Cellular Assembly and Organization** | Structural integrity of synapse | 0.0198 | SNTA1 | 1 |
| **Neurological Disease,Organismal Injury and Abnormalities,Psychological Disorders** | Preclinical stage amyotrophic lateral sclerosis | 0.0198 | FAS | 1 |
| **Nervous System Development and Function** | Development of ventral nerve cord | 0.0198 | OTX1 | 1 |
| **Embryonic Development,Nervous System Development and Function,Organ Development,Organismal Development,Tissue Development** | Formation of pretectum | 0.0198 | OTX1 | 1 |
| **Cellular Development,Nervous System Development and Function,Tissue Development** | Presynaptic differentiation of cortical neurons | 0.0198 | IL1RAPL1 | 1 |
| **Cell Death and Survival,Neurological Disease,Organismal Injury and Abnormalities** | Apoptosis of lactotropes | 0.0198 | FAS | 1 |
| **Cell Cycle,Nervous System Development and Function** | Mitogenesis of astrocytes | 0.0198 | EDN1 | 1 |
| **Neurological Disease,Skeletal and Muscular Disorders** | Hypernociception of hindpaw | 0.0198 | CX3CL1 | 1 |
| **Cell Death and Survival,Neurological Disease,Organismal Injury and Abnormalities** | Apoptosis of somatotrophs | 0.0198 | FAS | 1 |
| **Neurological Disease,Psychological Disorders** | First-onset paranoid schizophrenia | 0.0198 | VGF | 1 |
| **Nervous System Development and Function,Neurological Disease,Organ Morphology,Organismal Development,Organismal Injury and Abnormalities** | Abnormal morphology of enlarged cerebellum | 0.0198 | OTX1 | 1 |
| **Cellular Development,Cellular Growth and Proliferation,Nervous System Development and Function,Tissue Development** | Generation of Cajal-Retzius neurons | 0.0198 | FOXG1 | 1 |
| **Cell-To-Cell Signaling and Interaction,Cellular Assembly and Organization,Cellular Development,Cellular Function and Maintenance,Cellular Growth and Proliferation,Embryonic Development,Nervous System Development and Function,Organ Development,Organismal Development,Tissue Development** | Synaptogenesis of pyramidal neurons | 0.0198 | IL1RAPL1 | 1 |
| **Embryonic Development,Organismal Development,Tissue Development** | Formation of preplate | 0.0198 | FOXG1 | 1 |
| **Behavior** | Fear memory acquisition | 0.0198 | VGF | 1 |
| **Cell Morphology,Nervous System Development and Function** | Modulation of dopaminergic neurons | 0.0198 | EDN1 | 1 |
| **Cardiovascular System Development and Function,Tissue Morphology** | Contraction of cerebral artery | 0.0198 | EDN1 | 1 |
| **Cellular Function and Maintenance** | Endocytosis | 0.0228 | CX3CL1, KIAA1107, PPP3CA | 3 |
| **Embryonic Development,Nervous System Development and Function,Organ Development,Organ Morphology,Organismal Development,Tissue Development** | Morphogenesis of brain | 0.0238 | MKKS, OTX1, SOX2 | 3 |
| **Nervous System Development and Function,Neurological Disease,Organ Morphology,Organismal Development,Organismal Injury and Abnormalities** | Abnormal morphology of cerebellum | 0.0251 | HEXB, IL1RAPL1, MYH10, OTX1, SPARCL1 | 5 |
| **Cellular Movement** | Migration of cells | 0.028 | BSG, CCL4, CSPG5, CX3CL1, | 7 |

| | | | | |
|---|---|---|---|---|
| | | | FAS, FOXG1, MYH10 | |
| **Hereditary Disorder,Neurological Disease,Organismal Injury and Abnormalities,Psychological Disorders,Skeletal and Muscular Disorders** | Huntington's Disease | 0.0285 | ACTN2, CCL4, FOXG1, LAMB1, NR4A1, PPP3CA, SOX2 | 7 |
| **Embryonic Development,Nervous System Development and Function,Organ Development,Organismal Development,Tissue Development** | Formation of dorsal thalamus | 0.0296 | OTX1 | 1 |
| **Cell-To-Cell Signaling and Interaction,Nervous System Development and Function** | Adhesion of hippocampal neurons | 0.0296 | CX3CL1 | 1 |
| **Ophthalmic Disease,Organismal Injury and Abnormalities** | Proliferative vitreoretinopathy | 0.0296 | EDN1 | 1 |
| **Cellular Movement,Hematological System Development and Function,Immune Cell Trafficking,Inflammatory Response,Nervous System Development and Function** | Migration of microglia | 0.0296 | CX3CL1 | 1 |
| **Nervous System Development and Function,Neurological Disease,Organ Morphology,Organismal Development,Organismal Injury and Abnormalities** | Abnormal morphology of red nucleus | 0.0296 | OTX1 | 1 |
| **Cell-To-Cell Signaling and Interaction,Cellular Function and Maintenance,Inflammatory Response,Nervous System Development and Function** | Phagocytosis of neurons | 0.0296 | CX3CL1 | 1 |
| **Cell-To-Cell Signaling and Interaction,Endocrine System Development and Function,Nervous System Development and Function** | Desensitization of pituitary cells | 0.0296 | EDN1 | 1 |
| **Cell Morphology,Nervous System Development and Function** | Contraction of stellate cells | 0.0296 | EDN1 | 1 |
| **Embryonic Development,Nervous System Development and Function,Organ Development,Organismal Development,Tissue Development** | Development of third cerebral ventricle | 0.0296 | MYH10 | 1 |
| **Cardiovascular System Development and Function,Cell Morphology,Cellular Development,Organismal Development,Tissue Development** | Tubulation of microvascular endothelial cells | 0.0296 | PLXDC1 | 1 |
| **Cell-To-Cell Signaling and Interaction,Nervous System Development and Function** | Adhesion of cerebellar granule cell | 0.0296 | CX3CL1 | 1 |
| **Cell Death and Survival,Cellular Compromise,Neurological Disease,Tissue Morphology** | Degeneration of nerve ending | 0.0296 | ARNTL | 1 |
| **Cell-To-Cell Signaling and Interaction,Nervous System Development and Function** | Long term synaptic depression of perirhinal cortex | 0.0296 | HAPLN1 | 1 |
| **Cardiovascular System Development and Function** | Vasoconstriction of middle cerebral artery | 0.0296 | EDN1 | 1 |
| **Lipid Metabolism,Small Molecule Biochemistry** | Turnover of inositol phospholipid | 0.0296 | EDN1 | 1 |
| **Cellular Movement,Hematological System Development and Function,Immune Cell Trafficking,Inflammatory Response** | Cellular infiltration by microglia | 0.0296 | BSG | 1 |
| **Endocrine System Development and Function,Endocrine System Disorders,Nervous System Development and Function,Neurological Disease,Organ Development,Organ Morphology,Organismal Development,Organismal Injury and Abnormalities** | Abnormal morphology of mammillary body | 0.0296 | OTX1 | 1 |
| **Cellular Movement,Hematological System Development and Function,Immune Cell Trafficking,Inflammatory Response** | Influx of neutrophils | 0.0296 | CCL4 | 1 |
| **Embryonic Development,Nervous System Development and Function,Organ Development,Organismal Development,Tissue Development** | Formation of Ammon's horn | 0.0296 | OTX1 | 1 |
| **Neurological Disease,Organismal Injury and Abnormalities,Psychological Disorders** | Multiple system atrophy | 0.0296 | FAS | 1 |

| | | | | |
|---|---|---|---|---|
| **Cellular Development,Cellular Growth and Proliferation,Embryonic Development,Nervous System Development and Function,Organismal Development,Tissue Development** | Proliferation of neuroblasts | 0.0302 | CX3CL1, FOXG1, SOX2 | 3 |
| **Cell Signaling,Molecular Transport,Vitamin and Mineral Metabolism** | Concentration of Ca2+ | 0.0314 | ARNTL, EDN1 | 2 |
| **Embryonic Development,Nervous System Development and Function,Organ Development,Organismal Development,Tissue Development** | Development of cerebral cortex | 0.0368 | FOXG1, IL1RAPL1, MKKS, OTX1, SOX2 | 5 |
| **Cellular Development,Cellular Growth and Proliferation,Embryonic Development,Nervous System Development and Function,Organ Development,Organismal Development,Tissue Development** | Proliferation of neural precursor cells | 0.0375 | FOXG1, ID2, SOX2 | 3 |
| **Cellular Movement** | Cellular infiltration by myeloid cells | 0.0379 | BSG, CCL4 | 2 |
| **Cellular Movement,Hematological System Development and Function,Immune Cell Trafficking,Inflammatory Response** | Cellular infiltration by phagocytes | 0.0379 | BSG, CCL4 | 2 |
| **Embryonic Development,Nervous System Development and Function,Organ Development,Organismal Development,Tissue Development** | Development of fourth cerebral ventricle | 0.0392 | MYH10 | 1 |
| **Cellular Movement,Hematological System Development and Function,Immune Cell Trafficking,Inflammatory Response** | Cellular infiltration by monocytes | 0.0392 | CCL4 | 1 |
| **Nervous System Development and Function,Neurological Disease,Organ Morphology,Organismal Development,Organismal Injury and Abnormalities** | Abnormal morphology of pretectum | 0.0392 | OTX1 | 1 |
| **Lipid Metabolism,Molecular Transport,Small Molecule Biochemistry** | Concentration of docosahexaenoic acid | 0.0392 | ARNTL | 1 |
| **Nervous System Development and Function,Neurological Disease,Organ Morphology,Organismal Development,Organismal Injury and Abnormalities** | Abnormal morphology of mesencephalic nucleus | 0.0392 | OTX1 | 1 |
| **Cellular Function and Maintenance** | Regulation of neuroglia | 0.0392 | SOX2 | 1 |
| **Cardiovascular Disease,Neurological Disease,Organismal Injury and Abnormalities,Psychological Disorders** | Ischemic injury of brain | 0.0402 | CX3CL1, SNTA1 | 2 |
| **Nervous System Development and Function,Neurological Disease,Organ Morphology,Organismal Development,Organismal Injury and Abnormalities** | Abnormal morphology of Purkinje's layer of cerebellum | 0.0441 | HEXB, MYH10, SPARCL1 | 3 |
| **Cell Morphology,Cell-To-Cell Signaling and Interaction,Cellular Assembly and Organization,Nervous System Development and Function,Neurological Disease,Tissue Morphology** | Abnormal morphology of neuromuscular synapse | 0.0473 | SNTA1, SNTB2 | 2 |
| **Nervous System Development and Function,Neurological Disease,Organ Morphology,Organismal Development,Organismal Injury and Abnormalities** | Abnormal morphology of inferior colliculus | 0.0488 | OTX1 | 1 |
| **Nervous System Development and Function,Neurological Disease** | Abnormal morphology of trochlear nerve | 0.0488 | OTX1 | 1 |
| **Hematological System Development and Function,Lymphoid Tissue Structure and Development,Tissue Morphology** | Quantity of CD4+ T-lymphocytes | 0.0488 | ID2 | 1 |
| **Cellular Development,Cellular Growth and Proliferation,Embryonic Development,Nervous System Development and Function,Organ Development,Organismal Development,Tissue Development** | Proliferation of cerebellar granule neuron precursor cells | 0.0488 | SOX2 | 1 |
| **Cellular Development,Cellular Growth and Proliferation,Nervous System Development and Function,Organ Development,Tissue Development** | Proliferation of cerebellar granule cell | 0.0488 | LGI1 | 1 |

| | | | | |
|---|---|---|---|---|
| **Nervous System Development and Function,Neurological Disease,Organ Morphology,Organismal Development,Organismal Injury and Abnormalities** | Lack of diencephalon | 0.0488 | OTX1 | 1 |

### Cluster 13

| Categories | Diseases or Functions Annotation | p-Value | Molecules | # Molecules |
|---|---|---|---|---|
| **Carbohydrate Metabolism,Lipid Metabolism,Small Molecule Biochemistry** | Hydrolysis of phosphatidylinositol polyphosphate | 0.0141 | EDN1 | 1 |
| **Lipid Metabolism,Molecular Transport,Small Molecule Biochemistry** | Accumulation of glucosylceramide | 0.0141 | GBA | 1 |
| **Connective Tissue Development and Function,Organismal Development,Skeletal and Muscular System Development and Function,Tissue Development** | Size of cranium | 0.0141 | MYB | 1 |
| **Embryonic Development,Nervous System Development and Function,Organ Development,Organismal Development,Tissue Development** | Formation of subgranular zone of dentate gyrus | 0.0141 | FOXG1 | 1 |
| **Organismal Functions** | Regulation of pituitary gland | 0.0141 | NPY | 1 |
| **Cellular Assembly and Organization,Cellular Function and Maintenance** | Endocytosis of plasma membrane | 0.0141 | RAC1 | 1 |
| **Cellular Growth and Proliferation,Nervous System Development and Function,Organ Development** | Proliferation of stellate cells | 0.0141 | EDN1 | 1 |
| **Cellular Movement,Nervous System Development and Function** | Delay in initiation of migration of cortical projection neurons | 0.0141 | RAC1 | 1 |
| **Cellular Assembly and Organization,Cellular Function and Maintenance** | Quantity of peripheral lamellae | 0.0141 | RAC1 | 1 |
| **Cell-mediated Immune Response,Cell-To-Cell Signaling and Interaction,Cellular Movement,Hematological System Development and Function,Immune Cell Trafficking** | Sequestration of T lymphocytes | 0.0141 | HDC | 1 |
| **Connective Tissue Disorders,Inflammatory Disease,Inflammatory Response,Organismal Injury and Abnormalities,Skeletal and Muscular Disorders** | Degeneration of lumbar intervertebral disc | 0.0141 | EDN1 | 1 |
| **Drug Metabolism,Small Molecule Biochemistry** | Accumulation of epinephrine | 0.0141 | PNMT | 1 |
| **Cellular Development,Cellular Growth and Proliferation,Nervous System Development and Function,Tissue Development** | Development of Cajal-Retzius neurons | 0.0141 | FOXG1 | 1 |
| **Lipid Metabolism,Small Molecule Biochemistry** | Incorporation of sphingomyelin | 0.0141 | GBA | 1 |
| **Carbohydrate Metabolism,Lipid Metabolism,Small Molecule Biochemistry** | Incorporation of phosphatidylcholine | 0.0141 | GBA | 1 |
| **Cellular Movement,Nervous System Development and Function** | Positioning of pyramidal neurons | 0.0141 | PTPRA | 1 |
| **Drug Metabolism,Molecular Transport** | Efflux of glutathione | 0.0141 | EDN1 | 1 |
| **Cellular Development,Cellular Growth and Proliferation,Embryonic Development,Nervous System Development and Function,Organ Development,Organismal Development,Tissue Development** | Expansion of telencephalic progenitor cells | 0.0141 | FOXG1 | 1 |
| **Cellular Assembly and Organization,Cellular Function and Maintenance,Nervous System Development and Function,Tissue Development** | Organization of mossy fibers | 0.0141 | CHL1 | 1 |
| **Cell Morphology** | Contractility of stellate cells | 0.0141 | EDN1 | 1 |
| **Cardiovascular Disease,Developmental Disorder,Organismal Injury and Abnormalities** | Congenital heart block | 0.0141 | EDN1 | 1 |
| **Lipid Metabolism,Molecular Transport,Small Molecule Biochemistry** | Quantity of fat | 0.0141 | NPY | 1 |
| **Lipid Metabolism,Small Molecule Biochemistry** | Storage of fat | 0.0141 | NPY | 1 |
| **Lipid Metabolism,Molecular Transport,Small Molecule Biochemistry** | Concentration of eicosapentenoic acid | 0.0141 | ARNTL | 1 |

| | | | | |
|---|---|---|---|---|
| **Neurological Disease** | Formation of myelin ovoid | 0.0141 | RAC1 | 1 |
| **Cell-To-Cell Signaling and Interaction,Nervous System Development and Function** | Long-term potentiation of Schaffer collateral neurons | 0.0141 | PTPRA | 1 |
| **Cell Morphology,Cellular Assembly and Organization,Nervous System Development and Function** | Projection of corticothalamic axons | 0.0141 | RAC1 | 1 |
| **Infectious Diseases** | Infection by human herpesvirus 2 | 0.0141 | NECTIN1 | 1 |
| **Lipid Metabolism,Small Molecule Biochemistry** | Incorporation of choline | 0.0141 | GBA | 1 |
| **Cell Morphology,Nervous System Development and Function,Neurological Disease,Organ Morphology,Organismal Development,Organismal Injury and Abnormalities,Tissue Morphology** | Abnormal morphology of pyramidal neurons | 0.024 | CHL1, CRMP1, PTPRA | 3 |
| **Metabolic Disease** | Glucose metabolism disorder | 0.0262 | FAS, NPY | 2 |
| **Cell Death and Survival,Neurological Disease,Organismal Injury and Abnormalities** | Apoptosis of lactotropes | 0.028 | FAS | 1 |
| **Cellular Development,Cellular Growth and Proliferation,Embryonic Development,Nervous System Development and Function,Organ Development,Organismal Development,Tissue Development** | Development of spiny stellate neurons | 0.028 | SLC17A7 | 1 |
| **Organismal Functions** | Recovery of mice | 0.028 | FAS | 1 |
| **Nervous System Development and Function,Organ Morphology,Tissue Morphology** | Quantity of mossy fiber cells | 0.028 | RAC1 | 1 |
| **Cellular Growth and Proliferation,Nervous System Development and Function** | Inhibition of presympathetic neurons | 0.028 | NPY | 1 |
| **Behavior** | Inhibitory avoidance behavior | 0.028 | HDC | 1 |
| **Cell Cycle,Nervous System Development and Function** | Mitogenesis of astrocytes | 0.028 | EDN1 | 1 |
| **Cell Morphology,Cell-To-Cell Signaling and Interaction,Cellular Assembly and Organization** | Length of gap junctions | 0.028 | NRCAM | 1 |
| **Nervous System Development and Function,Organ Morphology,Organismal Development** | Thickness of granule cell layer | 0.028 | NRCAM | 1 |
| **Cell-To-Cell Signaling and Interaction,Cellular Growth and Proliferation,Nervous System Development and Function** | Excitation of type II spiral ganglion neurons | 0.028 | NPY | 1 |
| **Carbohydrate Metabolism,Drug Metabolism,Molecular Transport,Small Molecule Biochemistry** | Accumulation of hyaluronic acid | 0.028 | HEXB | 1 |
| **Cell Death and Survival,Neurological Disease,Organismal Injury and Abnormalities** | Apoptosis of somatotrophs | 0.028 | FAS | 1 |
| **Cellular Development,Cellular Growth and Proliferation,Nervous System Development and Function,Tissue Development** | Generation of Cajal-Retzius neurons | 0.028 | FOXG1 | 1 |
| **Behavior,Nervous System Development and Function** | Contextual fear memory retrieval | 0.028 | NPY | 1 |
| **Cell-To-Cell Signaling and Interaction,Nervous System Development and Function** | Synaptic transmission of thalamocortical neurons | 0.028 | SLC17A7 | 1 |
| **Cellular Assembly and Organization,Nervous System Development and Function** | Delay in sorting of axons | 0.028 | RAC1 | 1 |
| **Embryonic Development,Organismal Development,Tissue Development** | Formation of preplate | 0.028 | FOXG1 | 1 |
| **Cell Morphology,Cellular Assembly and Organization,Nervous System Development and Function** | Extension of leading processes | 0.028 | RAC1 | 1 |
| **Neurological Disease,Organismal Injury and Abnormalities,Psychological Disorders** | Preclinical stage amyotrophic lateral sclerosis | 0.028 | FAS | 1 |
| **Cardiovascular System Development and Function,Nervous System Development and Function,Organ Development** | Integrity of blood-brain barrier | 0.028 | HDC | 1 |
| **Hematological System Development and Function,Inflammatory Response,Tissue Morphology** | Quantity of monocytes | 0.028 | IL12RB2 | 1 |

| | | | | |
|---|---|---|---|---|
| **Cell Morphology,Nervous System Development and Function** | Modulation of dopaminergic neurons | 0.028 | EDN1 | 1 |
| **Cardiovascular System Development and Function,Tissue Morphology** | Contraction of cerebral artery | 0.028 | EDN1 | 1 |
| **Cell-To-Cell Signaling and Interaction,Cellular Assembly and Organization,Cellular Function and Maintenance** | Formation of spot-like adhesion site | 0.028 | NECTIN1 | 1 |
| **Cell Morphology,Cellular Assembly and Organization,Nervous System Development and Function** | Projection of axons | 0.029 | CHL1, RAC1 | 2 |
| **Cell Death and Survival** | Apoptosis of motor neurons | 0.0413 | FAS, SMPD2 | 2 |
| **Cell-To-Cell Signaling and Interaction,Nervous System Development and Function** | Long term synaptic depression of perirhinal cortex | 0.0418 | HAPLN1 | 1 |
| **Cellular Assembly and Organization** | Density of plasma membrane projections | 0.0418 | NUMBL | 1 |
| **Amino Acid Metabolism,Molecular Transport,Small Molecule Biochemistry** | Accumulation of L-glutamic acid | 0.0418 | SLC17A7 | 1 |
| **Cell-To-Cell Signaling and Interaction,Nervous System Development and Function** | Adhesion of hippocampal neurons | 0.0418 | NECTIN1 | 1 |
| **Cardiovascular System Development and Function** | Vasoconstriction of middle cerebral artery | 0.0418 | EDN1 | 1 |
| **Ophthalmic Disease,Organismal Injury and Abnormalities** | Proliferative vitreoretinopathy | 0.0418 | EDN1 | 1 |
| **Lipid Metabolism,Small Molecule Biochemistry** | Turnover of inositol phospholipid | 0.0418 | EDN1 | 1 |
| **Cell-To-Cell Signaling and Interaction,Endocrine System Development and Function,Nervous System Development and Function** | Desensitization of pituitary cells | 0.0418 | EDN1 | 1 |
| **Molecular Transport** | Internalization of water | 0.0418 | NPY | 1 |
| **Cell Morphology,Nervous System Development and Function** | Contraction of stellate cells | 0.0418 | EDN1 | 1 |
| **Infectious Diseases** | Infection by human herpesvirus 1 | 0.0418 | NECTIN1 | 1 |
| **Cell Morphology,Cellular Assembly and Organization,Nervous System Development and Function** | Projection of thalamocortical axons | 0.0418 | RAC1 | 1 |
| **Endocrine System Development and Function,Lipid Metabolism,Molecular Transport,Small Molecule Biochemistry** | Secretion of corticosterone | 0.0418 | NPY | 1 |
| **Cellular Movement,Hematological System Development and Function,Immune Cell Trafficking,Inflammatory Response** | Influx of neutrophils | 0.0418 | CCL4 | 1 |
| **Neurological Disease,Organismal Injury and Abnormalities,Psychological Disorders** | Multiple system atrophy | 0.0418 | FAS | 1 |
| **Cell Death and Survival,Cellular Compromise,Neurological Disease,Tissue Morphology** | Degeneration of nerve ending | 0.0418 | ARNTL | 1 |
| **Cell Morphology,Cellular Assembly and Organization,Cellular Function and Maintenance,Nervous System Development and Function** | Formation of astrocyte projections | 0.0418 | RAC1 | 1 |

### Cluster 14

| Categories | Diseases or Functions Annotation | p-Value | Molecules | # Molecules |
|---|---|---|---|---|
| **Cell Morphology,Cellular Assembly and Organization** | Permeability of mitochondria | 0.000307 | ADCYAP1, HTT | 2 |
| **Small Molecule Biochemistry** | Quantity of kynurenic acid | 0.000307 | HTT, KMO | 2 |
| **Infectious Diseases,Neurological Disease** | Infection of central nervous system | 0.000584 | NECTIN1, SEMA7A, TICAM1 | 3 |
| **Neurological Disease** | Neurological deficiency | 0.000909 | ADCYAP1, HTT | 2 |
| **Infectious Diseases** | Infection by human herpesvirus 1 | 0.000909 | NECTIN1, TICAM1 | 2 |
| **Cell-To-Cell Signaling and Interaction,Nervous System Development and Function** | Long term synaptic depression of perirhinal cortex | 0.000909 | HAPLN1, HTT | 2 |

| | | | | |
|---|---|---|---|---|
| **Embryonic Development,Nervous System Development and Function,Organ Development,Organismal Development,Tissue Development** | Development of striatum | 0.00118 | FOXP2, HPRT1, HTT, MKKS | 4 |
| **Carbohydrate Metabolism,Small Molecule Biochemistry** | Synthesis of inositol phosphate | 0.0018 | ADCYAP1, HTT | 2 |
| **Nervous System Development and Function,Neurological Disease,Organ Morphology,Organismal Development,Organismal Injury and Abnormalities** | Abnormal morphology of third cerebral ventricle | 0.00283 | CELSR2, MKKS, NFIX, OTX1 | 4 |
| **Nervous System Development and Function,Neurological Disease,Organ Morphology,Organismal Development,Organismal Injury and Abnormalities** | Abnormal morphology of olfactory tract | 0.00296 | MAP3K12, SEMA7A | 2 |
| **Lipid Metabolism,Small Molecule Biochemistry** | Synthesis of glycosphingolipid | 0.00296 | ADCYAP1, HTT | 2 |
| **Cellular Development,Cellular Growth and Proliferation,Nervous System Development and Function,Tissue Development** | Development of bipolar cells | 0.00296 | FOXG1, PRDM1 | 2 |
| **Cellular Assembly and Organization,Cellular Movement,Nervous System Development and Function** | Transport of synaptic vesicles | 0.00296 | HTT, KIAA1107, PNKD, PPP3CA, RPH3A, SCAMP5 | 6 |
| **Nervous System Development and Function,Organ Morphology,Organismal Development** | Size of metencephalon | 0.00302 | ADCYAP1, IGFBP6, OTX1 | 3 |
| **Neurological Disease** | Neurological signs | 0.00355 | ACTN2, ATP6V1A, CAMKK2, CCL4, FOXG1, GBA, HTT, OXR1, PPP3CA, RPH3A, SLC17A7, THBS3, ZMYND8 | 13 |
| **Metabolic Disease** | Glucose metabolism disorder | 0.00358 | FAS, NPY, SH2B1 | 3 |
| **Cell Signaling,Molecular Transport,Vitamin and Mineral Metabolism** | Quantity of Ca2+ | 0.00399 | ADCYAP1, CAMK2A, CCL4, HTT, NPY, NRN1 | 6 |
| **Infectious Diseases,Neurological Disease,Organismal Injury and Abnormalities** | Infection of brain | 0.00439 | NECTIN1, SEMA7A | 2 |
| **Neurological Disease,Organismal Injury and Abnormalities** | Hydrocephalus | 0.00487 | CELSR2, MYB, NFIX | 3 |
| **Cell Death and Survival** | Cell death of central nervous system cells | 0.00595 | ADCYAP1, CAMK2A, CAPNS1, CCL4, FAS, FOXG1, GBA, HTT, MAP3K12, MEF2C, SLC25A12, STK4, TICAM1 | 13 |
| **Hereditary Disorder,Neurological Disease,Organismal Injury and Abnormalities,Psychological Disorders,Skeletal and Muscular Disorders** | Huntington's Disease | 0.00606 | ACTN2, ATP6V1A, CAMKK2, CCL4, FOXG1, HTT, OXR1, PPP3CA, RPH3A, SLC17A7, THBS3, ZMYND8 | 12 |

| | | | | |
|---|---|---|---|---|
| **Cell-To-Cell Signaling and Interaction,Nervous System Development and Function** | Long term synaptic depression of hippocampal neurons | 0.00607 | CAMK2A, HTT | 2 |
| **Cell Death and Survival** | Apoptosis of multilineage progenitor cells | 0.00607 | HTT, ID2 | 2 |
| **Behavior** | Cognition | 0.00716 | CAMK2A, CAPNS1, HAPLN1, HTT, NPY, SRF | 6 |
| **Neurological Disease,Organ Morphology,Organismal Injury and Abnormalities** | Atrophy of hippocampus | 0.00801 | HTT, TICAM1 | 2 |
| **Cellular Movement** | Migration of granule neuron progenitor cells | 0.00801 | CAMKK2, NFIX | 2 |
| **Cell Morphology,Cell-To-Cell Signaling and Interaction,Cellular Function and Maintenance,Nervous System Development and Function** | Depolarization of pyramidal neurons | 0.00801 | HTT, PANX1 | 2 |
| **Cell Death and Survival** | Cell death of brain | 0.0081 | ADCYAP1, CAMK2A, CAPNS1, CCL4, FAS, FOXG1, GBA, HTT, MEF2C, PPP3CA, SLC25A12, STK4, TICAM1 | 13 |
| **Cell Death and Survival,Neurological Disease,Organismal Injury and Abnormalities** | Cell death of cerebral cortex cells | 0.00816 | ADCYAP1, CAMK2A, CAPNS1, CCL4, FAS, GBA, HTT, MEF2C, SLC25A12, STK4, TICAM1 | 11 |
| **Amino Acid Metabolism,Molecular Transport,Small Molecule Biochemistry** | Uptake of L-glutamic acid | 0.0102 | ADCYAP1, HTT | 2 |
| **Cell Death and Survival,Neurological Disease,Organismal Injury and Abnormalities** | Cell death of brain cells | 0.0114 | ADCYAP1, CAMK2A, CAPNS1, CCL4, FAS, FOXG1, GBA, HTT, MEF2C, SLC25A12, STK4, TICAM1 | 12 |
| **Cell Death and Survival,Embryonic Development** | Cell death of embryonic cells | 0.0126 | CAPNS1, FAS | 2 |
| **Behavior,Nervous System Development and Function** | Memory | 0.0138 | CAMK2A, CAPNS1, HAPLN1, NPY | 4 |
| **Nervous System Development and Function,Tissue Morphology** | Quantity of astrocytes | 0.0153 | HTT, IGFBP6, RB1CC1 | 3 |
| **Cell Death and Survival,Nervous System Development and Function** | Cell viability of granule cells | 0.0153 | ADCYAP1, FOXG1, NRCAM | 3 |
| **Cell Death and Survival,Neurological Disease,Organismal Injury and Abnormalities** | Cell death of granule cells | 0.0153 | ADCYAP1, CAPNS1, FAS, FOXG1, STK4 | 5 |
| **Cell Morphology,Cell-To-Cell Signaling and Interaction,Cellular Function and Maintenance,Nervous System Development and Function** | Depolarization of neurons | 0.0167 | ADCYAP1, HTT, PANX1 | 3 |

| | | | | |
|---|---|---|---|---|
| **Cell-To-Cell Signaling and Interaction,Nervous System Development and Function** | Long-term potentiation of brain | 0.0171 | CAMK2A, CAPNS1, CHRD, GBA, HTT, PPP3CA, SYNPO | 7 |
| **Cell Cycle,Gene Expression** | Binding of Sp1 binding site | 0.0175 | HTT | 1 |
| **Cellular Assembly and Organization,Nervous System Development and Function** | Fasciculation of parallel fiber | 0.0175 | NFIX | 1 |
| **Molecular Transport,Nucleic Acid Metabolism,Small Molecule Biochemistry** | Uptake of ADP | 0.0175 | HTT | 1 |
| **Cellular Compromise,Cellular Function and Maintenance,Organismal Injury and Abnormalities** | Endoplasmic reticulum stress response of striatal neurons | 0.0175 | HTT | 1 |
| **Cell Death and Survival,Neurological Disease,Organismal Injury and Abnormalities** | Cell death of mesencephalic neurons | 0.0175 | CAPNS1 | 1 |
| **Embryonic Development,Nervous System Development and Function,Organ Development,Organismal Development,Tissue Development** | Development of putamen | 0.0175 | FOXP2 | 1 |
| **Neurological Disease,Organismal Injury and Abnormalities** | Atrophy of caudate nucleus | 0.0175 | HTT | 1 |
| **Nervous System Development and Function,Organ Morphology,Tissue Morphology** | Quantity of rod bipolar cells | 0.0175 | PRDM1 | 1 |
| **Cellular Assembly and Organization,Cellular Compromise** | Formation of polyQ inclusions | 0.0175 | HTT | 1 |
| **Cell Morphology,Cellular Assembly and Organization,Cellular Development,Cellular Function and Maintenance,Cellular Growth and Proliferation,Embryonic Development,Nervous System Development and Function,Organ Development,Organismal Development,Tissue Development** | Neuritogenesis of cerebellar granule cell | 0.0175 | ADCYAP1 | 1 |
| **Lipid Metabolism,Small Molecule Biochemistry** | Synthesis of ganglioside GM1 | 0.0175 | HTT | 1 |
| **Cellular Assembly and Organization** | Formation of neuritic aggregates | 0.0175 | HTT | 1 |
| **Nervous System Development and Function,Organ Morphology,Organismal Development** | Volume of globus pallidus | 0.0175 | HTT | 1 |
| **Carbohydrate Metabolism,Lipid Metabolism,Small Molecule Biochemistry** | Incorporation of phosphatidylcholine | 0.0175 | GBA | 1 |
| **Free Radical Scavenging** | Clearance of reactive oxygen species | 0.0175 | HTT | 1 |
| **Lipid Metabolism,Small Molecule Biochemistry** | Damage of phospholipid | 0.0175 | HTT | 1 |
| **Neurological Disease,Organismal Injury and Abnormalities,Psychological Disorders** | Atrophy of globus pallidus | 0.0175 | HTT | 1 |
| **Cell Morphology,Cell-To-Cell Signaling and Interaction,Cellular Assembly and Organization** | Electrical resistance of medium spiny neurons | 0.0175 | HTT | 1 |
| **Cell Death and Survival,Cellular Development,Nervous System Development and Function,Tissue Development,Tissue Morphology** | Regeneration of bipolar cells | 0.0175 | PRDM1 | 1 |
| **Neurological Disease,Organismal Injury and Abnormalities** | Neurodegeneration of thalamic nucleus | 0.0175 | HTT | 1 |
| **Neurological Disease** | Mechanosensory defect | 0.0175 | HTT | 1 |
| **Cell Death and Survival** | Cell death of GABAergic neurons | 0.0175 | HTT | 1 |
| **Cellular Development,Cellular Growth and Proliferation,Embryonic Development,Nervous System Development and Function,Organ Development,Organismal Development,Tissue Development** | Expansion of telencephalic progenitor cells | 0.0175 | FOXG1 | 1 |
| **Cell Death and Survival,Embryonic Development,Tissue Morphology** | Quantity of apoptotic neural crest cells | 0.0175 | CAPNS1 | 1 |
| **Cellular Assembly and Organization** | Quantity of polyglutamine inclusion | 0.0175 | HTT | 1 |

| | | | | |
|---|---|---|---|---|
| **Cell Morphology,Nervous System Development and Function,Organ Morphology,Organismal Development,Tissue Morphology** | Area of striatal neurons | 0.0175 | HTT | 1 |
| **Carbohydrate Metabolism,Small Molecule Biochemistry** | Consumption of D-glucose | 0.0175 | HTT | 1 |
| **Cellular Assembly and Organization** | Biogenesis of mitochondria | 0.0175 | HTT | 1 |
| **Cellular Assembly and Organization,Cellular Function and Maintenance** | Formation of membrane blebs | 0.0175 | HTT | 1 |
| **Cellular Assembly and Organization** | Uncoupling of mitochondria | 0.0175 | HTT | 1 |
| **Cellular Compromise,Neurological Disease,Organismal Injury and Abnormalities** | Dysfunction of striatal neurons | 0.0175 | HTT | 1 |
| **Cell Cycle** | Quiescence of neural stem cells | 0.0175 | NFIX | 1 |
| **Cellular Compromise** | Depletion of striatal neurons | 0.0175 | HTT | 1 |
| **Cellular Compromise,Neurological Disease,Organismal Injury and Abnormalities** | Damage of projection neurons | 0.0175 | HTT | 1 |
| **Nervous System Development and Function** | Regulation of superior cervical ganglion | 0.0175 | ADCYAP1 | 1 |
| **Cellular Development,Cellular Growth and Proliferation,Nervous System Development and Function,Tissue Development** | Development of Cajal-Retzius neurons | 0.0175 | FOXG1 | 1 |
| **Drug Metabolism** | Binding of flumazenil | 0.0175 | HTT | 1 |
| **Neurological Disease,Organismal Injury and Abnormalities** | Neurodegeneration of amygdala | 0.0175 | HTT | 1 |
| **Nervous System Development and Function** | Input resistance of medium spiny neurons | 0.0175 | HTT | 1 |
| **Cell Morphology,Inflammatory Disease,Inflammatory Response,Neurological Disease,Organ Morphology,Organismal Injury and Abnormalities** | Swelling of medium spiny neurons | 0.0175 | HTT | 1 |
| **Lipid Metabolism,Molecular Transport,Small Molecule Biochemistry** | Quantity of fat | 0.0175 | NPY | 1 |
| **Lipid Metabolism,Small Molecule Biochemistry** | Storage of fat | 0.0175 | NPY | 1 |
| **Nervous System Development and Function,Organ Morphology,Organismal Development** | Volume of cerebellar cortex | 0.0175 | ADCYAP1 | 1 |
| **Cell Death and Survival,Cellular Assembly and Organization,Cellular Development,Nervous System Development and Function,Tissue Development,Tissue Morphology** | Regeneration of motor axons | 0.0175 | MAP3K12 | 1 |
| **Lipid Metabolism,Molecular Transport,Small Molecule Biochemistry** | Concentration of eicosapentenoic acid | 0.0175 | ARNTL | 1 |
| **Neurological Disease,Organismal Injury and Abnormalities,Psychological Disorders** | Atrophy of putamen | 0.0175 | HTT | 1 |
| **Cell Death and Survival,Cellular Compromise,Neurological Disease,Tissue Morphology** | Neurodegeneration of ASH sensory neurons | 0.0175 | HTT | 1 |
| **Lipid Metabolism,Small Molecule Biochemistry** | Synthesis of sulfatides | 0.0175 | ADCYAP1 | 1 |
| **Cell Morphology,Cellular Assembly and Organization,Cellular Development,Cellular Function and Maintenance,Cellular Growth and Proliferation,Nervous System Development and Function,Organismal Development,Tissue Development** | Repolarization of medium spiny neurons | 0.0175 | HTT | 1 |
| **Cell Signaling,Molecular Transport,Vitamin and Mineral Metabolism** | Localization of Ca2+ | 0.0175 | HTT | 1 |
| **Cellular Function and Maintenance** | State 3 respiration of mitochondria | 0.0175 | HTT | 1 |
| **Cell Death and Survival,Cellular Compromise,Neurological Disease,Organismal Injury and Abnormalities,Psychological Disorders,Tissue Morphology** | Fragmentation of striatal neurons | 0.0175 | HTT | 1 |
| **Cell Morphology,Nervous System Development and Function,Tissue Morphology** | Morphology of somatodendritic region | 0.0175 | HTT | 1 |
| **Nervous System Development and Function,Organ Morphology,Organismal Development** | Volume of substantia nigra pars compacta | 0.0175 | HTT | 1 |
| **Connective Tissue Development and Function,Organismal Development,Skeletal and Muscular System Development and Function,Tissue Development** | Size of cranium | 0.0175 | MYB | 1 |

| | | | | |
|---|---|---|---|---|
| **Embryonic Development,Nervous System Development and Function,Organ Development,Organismal Development,Tissue Development** | Formation of subgranular zone of dentate gyrus | 0.0175 | FOXG1 | 1 |
| **Organismal Functions** | Regulation of pituitary gland | 0.0175 | NPY | 1 |
| **Cellular Compromise** | Acidification of cytosol | 0.0175 | CA4 | 1 |
| **Neurological Disease** | Dystonia | 0.0175 | HTT | 1 |
| **Carbohydrate Metabolism,Lipid Metabolism,Molecular Transport,Small Molecule Biochemistry** | Quantity of sn-glycero-3-phosphocholine | 0.0175 | HTT | 1 |
| **Neurological Disease,Organismal Injury and Abnormalities** | Degeneration of median forebrain bundle | 0.0175 | HTT | 1 |
| **Cellular Development,Cellular Growth and Proliferation,Embryonic Development,Nervous System Development and Function,Organ Development,Organismal Development,Tissue Development** | Neurogenesis of cortical precursor cells | 0.0175 | FOXP2 | 1 |
| **Cardiovascular System Development and Function,Cell Morphology,Nervous System Development and Function** | Electrical capacitance of medium spiny neurons | 0.0175 | HTT | 1 |
| **Neurological Disease** | Onset of motor incoordination | 0.0175 | HTT | 1 |
| **Lipid Metabolism,Molecular Transport,Small Molecule Biochemistry** | Quantity of succinic acid | 0.0175 | HTT | 1 |
| **Cell-To-Cell Signaling and Interaction,Cellular Function and Maintenance,Nervous System Development and Function** | Paired-pulse facilitation of medium spiny neurons | 0.0175 | HTT | 1 |
| **Nervous System Development and Function** | Plasticity of barrel cortex | 0.0175 | HTT | 1 |
| **Cell-To-Cell Signaling and Interaction,Nervous System Development and Function** | Activation of trigeminovascular neuron | 0.0175 | ADCYAP1 | 1 |
| **Neurological Disease** | Aggregation of htt body | 0.0175 | HTT | 1 |
| **Lipid Metabolism,Small Molecule Biochemistry** | Incorporation of choline | 0.0175 | GBA | 1 |
| **Cardiovascular System Development and Function,Cell Morphology,Nervous System Development and Function** | Electrical capacitance of pyramidal neurons | 0.0175 | HTT | 1 |
| **Cell Death and Survival** | Cell death of ASH sensory neurons | 0.0175 | HTT | 1 |
| **Nervous System Development and Function,Neurological Disease,Organ Morphology,Organismal Development,Organismal Injury and Abnormalities** | Abnormal morphology of subthalamic nucleus | 0.0175 | HTT | 1 |
| **Neurological Disease,Organ Morphology,Organismal Injury and Abnormalities** | Atrophy of hippocampal CA3 region | 0.0175 | TICAM1 | 1 |
| **Lipid Metabolism,Molecular Transport,Small Molecule Biochemistry** | Accumulation of glucosylceramide | 0.0175 | GBA | 1 |
| **Cellular Assembly and Organization** | Transport of cytoplasm | 0.0175 | HTT | 1 |
| **Neurological Disease,Organismal Injury and Abnormalities** | Interictal epileptiform discharge of hippocampal neurons | 0.0175 | CAMK2A | 1 |
| **Lipid Metabolism,Small Molecule Biochemistry** | Loss of cholesterol | 0.0175 | HTT | 1 |
| **Behavior** | Self-administration of amphetamine | 0.0175 | CAMK2A | 1 |
| **Amino Acid Metabolism,Molecular Transport,Small Molecule Biochemistry** | Concentration of phosphocreatine | 0.0175 | HTT | 1 |
| **Neurological Disease** | Degeneration of ventral nerve cord | 0.0175 | HTT | 1 |
| **Cellular Assembly and Organization,Cellular Development,Embryonic Development,Nervous System Development and Function,Organ Development,Organismal Development,Tissue Development** | Maturation of olfactory glomeruli | 0.0175 | PPP3CA | 1 |
| **Cell-To-Cell Signaling and Interaction,Nervous System Development and Function** | Activation of mitral cells | 0.0175 | GJD2 | 1 |
| **Lipid Metabolism,Small Molecule Biochemistry** | Incorporation of sphingomyelin | 0.0175 | GBA | 1 |

| | | | | |
|---|---|---|---|---|
| **Cell Morphology** | Modification of pyramidal neurons | 0.0175 | HTT | 1 |
| **Cell-To-Cell Signaling and Interaction,Nervous System Development and Function** | Long-term potentiation of cholinergic neurons | 0.0175 | HTT | 1 |
| **Cell-To-Cell Signaling and Interaction,Nervous System Development and Function** | Excitatory postsynaptic potential of medium spiny neurons | 0.0175 | HTT | 1 |
| **Cellular Movement** | Migration of neuroepithelial cells | 0.0175 | HTT | 1 |
| **Cell Morphology,Skeletal and Muscular System Development and Function** | Hypercontractility of vascular smooth muscle cells | 0.0175 | SRF | 1 |
| **Cell Morphology,Cellular Assembly and Organization,Nervous System Development and Function,Tissue Morphology** | Size of synaptic vesicles pool | 0.0175 | SCAMP5 | 1 |
| **Embryonic Development,Nervous System Development and Function,Organ Development,Organismal Development,Tissue Development** | Development of caudate nucleus | 0.0175 | FOXP2 | 1 |
| **Cell Death and Survival,Cellular Assembly and Organization,Cellular Function and Maintenance** | Formation of apoptotic bodies | 0.0175 | HTT | 1 |
| **Cell Morphology,Cell-To-Cell Signaling and Interaction,Cellular Function and Maintenance,Nervous System Development and Function** | Resting membrane potential of medium spiny neurons | 0.0175 | HTT | 1 |
| **Infectious Diseases** | Infection by human herpesvirus 2 | 0.0175 | NECTIN1 | 1 |
| **DNA Replication, Recombination, and Repair** | Breakage of double-stranded DNA | 0.0175 | HTT | 1 |
| **Neurological Disease,Organismal Injury and Abnormalities** | Gliosis of caudate nucleus | 0.0175 | HTT | 1 |
| **Cellular Development,Cellular Growth and Proliferation,Nervous System Development and Function,Organ Development,Tissue Development** | Proliferation of granule cells | 0.018 | ADCYAP1, CAMKK2 | 2 |
| **Embryonic Development,Nervous System Development and Function,Organ Development,Organismal Development,Tissue Development** | Formation of hippocampus | 0.0184 | FOXG1, HTT, MKKS, OTX1, SH2B1, SRF | 6 |
| **Cell Death and Survival,Neurological Disease,Organismal Injury and Abnormalities** | Cell death of hippocampal neurons | 0.0184 | ADCYAP1, CAMK2A, FAS, HTT, STK4 | 5 |
| **Behavior** | Learning | 0.0184 | CAMK2A, CAPNS1, HAPLN1, NPY, SRF | 5 |
| **Cell Death and Survival** | Neuronal cell death | 0.0192 | ADCYAP1, CAMK2A, CAPNS1, CCL4, FAS, FOXG1, GBA, HTT, ID2, MAP3K12, MEF2C, NRCAM, NRN1, PANX1, PPP3CA, SLC25A12, STK4, TICAM1 | 18 |
| **Cellular Development,Cellular Growth and Proliferation,Embryonic Development,Nervous System Development and Function,Organismal Development,Tissue Morphology** | Quantity of neuroblasts | 0.0211 | NAP1L2, RB1CC1 | 2 |
| **Nervous System Development and Function,Neurological Disease,Organ Morphology,Organismal Development,Organismal Injury and Abnormalities** | Abnormal morphology of telencephalon | 0.0212 | CELSR2, HTT, ID2, MAP3K12, MKKS, NFIX, OTX1, SEMA7A | 8 |

| | | | | |
|---|---|---|---|---|
| **Embryonic Development,Nervous System Development and Function,Organ Development,Organismal Development,Tissue Development** | Formation of forebrain | 0.0214 | CHRD, FOXG1, FOXP2, HPRT1, HTT, ID2, MKKS, OTX1, PPP3CA, SRF | 10 |
| **Cell Death and Survival,Neurological Disease,Organismal Injury and Abnormalities** | Cell death of cortical neurons | 0.0236 | CAPNS1, CCL4, FAS, GBA, HTT, MEF2C, SLC25A12, TICAM1 | 8 |
| **Nervous System Development and Function,Neurological Disease,Organ Morphology,Organismal Development,Organismal Injury and Abnormalities** | Abnormal morphology of diencephalon | 0.0238 | CELSR2, HTT, MKKS, NFIX, OTX1 | 5 |
| **Nucleic Acid Metabolism,Small Molecule Biochemistry** | Synthesis of purine nucleotide | 0.0243 | ADCYAP1, HTT | 2 |
| **Nervous System Development and Function,Neurological Disease,Organ Morphology,Organismal Development,Organismal Injury and Abnormalities** | Third ventricle dilation | 0.0243 | CELSR2, NFIX | 2 |
| **Cell-To-Cell Signaling and Interaction,Nervous System Development and Function** | Long-term potentiation of hippocampus | 0.0244 | CAMK2A, CAPNS1, CHRD, HTT, PPP3CA, SYNPO | 6 |
| **Cell-To-Cell Signaling and Interaction,Nervous System Development and Function** | Long-term potentiation | 0.025 | B3GAT1, CAMK2A, CAMKK2, CAPNS1, CHRD, GBA, HTT, PPP3CA, SLC17A7, SYNPO | 10 |
| **Cellular Assembly and Organization,Cellular Function and Maintenance,Cellular Movement,Nervous System Development and Function** | Endocytosis of synaptic vesicles | 0.0267 | KIAA1107, PPP3CA, SCAMP5 | 3 |
| **Cellular Development,Cellular Growth and Proliferation,Embryonic Development,Nervous System Development and Function,Organ Development,Organismal Development,Tissue Development** | Formation of brain cells | 0.0272 | ADCYAP1, FOXP2, SH2B1, SLC17A7 | 4 |
| **Nervous System Development and Function,Organ Morphology,Organismal Development** | Size of cerebellum | 0.0277 | ADCYAP1, IGFBP6 | 2 |
| **Nutritional Disease,Psychological Disorders** | Hyperphagia | 0.0277 | NPY, SH2B1 | 2 |
| **Cellular Movement,Nervous System Development and Function** | Migration of granule cells | 0.0287 | ADCYAP1, CELSR2, FOXG1 | 3 |
| **Behavior** | Conditioning | 0.0287 | CAMK2A, NPY, SRF | 3 |
| **Lipid Metabolism,Molecular Transport,Small Molecule Biochemistry** | Quantity of steroid | 0.0287 | ADCYAP1, FAS, HTT | 3 |
| **Neurological Disease,Skeletal and Muscular Disorders** | Neuromuscular disease | 0.0289 | ACTN2, ATP6V1A, CAMKK2, CCL4, FOXG1, HTT, NRCAM, OXR1, PPP3CA, RPH3A, SLC17A7, THBS3, ZMYND8 | 13 |
| **Neurological Disease,Organismal Injury and Abnormalities** | Brain lesion | 0.0311 | ADAMTS8, ADCYAP1, | 20 |

| | | | | |
|---|---|---|---|---|
| | | | AK5, ANKRD30B, ARHGEF18, ATP2A2, ATP6V1A, BCL11A, CDH9, DBNDD1, FOXP2, HSPBP1, NAP1L2, PANX1, PRDM1, RB1CC1, ROS1, SV2B, THEMIS, TICAM1 | |
| **Cellular Movement** | Cell movement | 0.0319 | ADCYAP1, CAMKK2, CCL4, CELSR2, FAS, FOXG1, HTT, NFIX, NRCAM, SRF, TIAM2 | 11 |
| **Cellular Assembly and Organization** | Formation of plasma membrane | 0.0329 | CDH9, GJD2, HTT, NECTIN1, NRCAM, NRN1, SEZ6L2 | 7 |
| **Neurological Disease,Organismal Injury and Abnormalities,Psychological Disorders** | Disorder of basal ganglia | 0.0336 | ACTN2, ATP6V1A, CAMKK2, CCL4, FAS, FOXG1, HTT, OXR1, PPP3CA, RPH3A, SLC17A7, THBS3, ZMYND8 | 13 |
| **Cell-To-Cell Signaling and Interaction,Cellular Assembly and Organization** | Formation of intercellular junctions | 0.0338 | CDH9, GJD2, HTT, NECTIN1, NRCAM, NRN1, SEZ6L2 | 7 |
| **Nervous System Development and Function,Neurological Disease,Organ Morphology,Organismal Development,Organismal Injury and Abnormalities** | Abnormal morphology of oculomotor nucleus | 0.0348 | OTX1 | 1 |
| **Cellular Growth and Proliferation,Nervous System Development and Function** | Inhibition of presympathetic neurons | 0.0348 | NPY | 1 |
| **Molecular Transport** | Quantity of acetate | 0.0348 | HTT | 1 |
| **Cellular Assembly and Organization,Cellular Function and Maintenance,Cellular Movement,Nervous System Development and Function** | Retrograde axonal transport | 0.0348 | HTT | 1 |
| **Nervous System Development and Function,Organ Morphology,Organismal Development** | Thickness of granule cell layer | 0.0348 | NRCAM | 1 |
| **Cell Morphology,Cellular Assembly and Organization,Cellular Development,Cellular Function and Maintenance,Cellular Growth and Proliferation,Nervous System Development and Function,Organismal Development,Tissue Development** | Sprouting of motor neurons | 0.0348 | HTT | 1 |
| **Cell-To-Cell Signaling and Interaction,Cellular Growth and Proliferation,Nervous System Development and Function** | Excitation of type II spiral ganglion neurons | 0.0348 | NPY | 1 |
| **Lipid Metabolism,Small Molecule Biochemistry** | Internalization of lactosylceramide | 0.0348 | HTT | 1 |

| | | | | |
|---|---|---|---|---|
| **Nervous System Development and Function,Organ Morphology,Organismal Development** | Mass of cerebellum | 0.0348 | IGFBP6 | 1 |
| **Infectious Diseases,Neurological Disease,Organismal Injury and Abnormalities** | Infection of cortical neurons | 0.0348 | SEMA7A | 1 |
| **Cellular Function and Maintenance** | Respiration of neurons | 0.0348 | SLC25A12 | 1 |
| **Nervous System Development and Function** | Development of ventral nerve cord | 0.0348 | OTX1 | 1 |
| **Cell-To-Cell Signaling and Interaction,Cellular Assembly and Organization,Cellular Function and Maintenance** | Formation of spot-like adhesion site | 0.0348 | NECTIN1 | 1 |
| **Cell Morphology,Cellular Assembly and Organization,Cellular Development,Cellular Function and Maintenance,Cellular Growth and Proliferation,Embryonic Development,Nervous System Development and Function,Organismal Development,Tissue Development** | Branching of basal dendrites | 0.0348 | CAPNS1 | 1 |
| **Embryonic Development,Nervous System Development and Function,Organ Development,Organismal Development,Tissue Development** | Formation of pretectum | 0.0348 | OTX1 | 1 |
| **Cardiovascular Disease,Hematological Disease,Metabolic Disease** | Hyperlipidemia | 0.0348 | SH2B1 | 1 |
| **Cellular Assembly and Organization,Cellular Function and Maintenance,Cellular Movement,Nervous System Development and Function** | Anterograde axonal transport | 0.0348 | HTT | 1 |
| **Cell-To-Cell Signaling and Interaction,Nervous System Development and Function** | Activation of parasympathetic nerve | 0.0348 | ADCYAP1 | 1 |
| **Neurological Disease,Organismal Injury and Abnormalities** | Gliosis of thalamus | 0.0348 | HTT | 1 |
| **Carbohydrate Metabolism,Molecular Transport,Small Molecule Biochemistry,Vitamin and Mineral Metabolism** | Concentration of ascorbic acid | 0.0348 | HTT | 1 |
| **Cell Death and Survival,Cellular Compromise,Neurological Disease,Tissue Morphology** | Degeneration of GABAergic neurons | 0.0348 | HTT | 1 |
| **Amino Acid Metabolism** | Quantity of glutamine | 0.0348 | HTT | 1 |
| **Cellular Assembly and Organization,Cellular Function and Maintenance,Cellular Movement,Nervous System Development and Function** | Axonal transport of autophagosomes | 0.0348 | HTT | 1 |
| **Cell Death and Survival,Neurological Disease,Organismal Injury and Abnormalities** | Apoptosis of somatotrophs | 0.0348 | FAS | 1 |
| **Cellular Assembly and Organization,DNA Replication, Recombination, and Repair** | Formation of nuclear foci | 0.0348 | HTT | 1 |
| **Cell-To-Cell Signaling and Interaction** | Signaling of astrocytes | 0.0348 | HTT | 1 |
| **Cellular Development,Cellular Growth and Proliferation,Nervous System Development and Function,Tissue Development** | Generation of Cajal-Retzius neurons | 0.0348 | FOXG1 | 1 |
| **Cell-To-Cell Signaling and Interaction,Nervous System Development and Function** | Coupling of neurons | 0.0348 | GJD2 | 1 |
| **Cancer,Neurological Disease,Organismal Injury and Abnormalities,Tumor Morphology** | Production of glioblastoma | 0.0348 | ROS1 | 1 |
| **Cellular Assembly and Organization** | Quantity of nuclear inclusions | 0.0348 | HTT | 1 |
| **Cell Morphology,Nervous System Development and Function,Organ Morphology,Organismal Development,Tissue Morphology** | Volume of striatal neurons | 0.0348 | HTT | 1 |
| **Cell-To-Cell Signaling and Interaction,Nervous System Development and Function** | Synaptic transmission of medium spiny neurons | 0.0348 | HTT | 1 |
| **Cellular Development,Cellular Growth and Proliferation,Embryonic Development,Nervous System Development and Function,Organ Development,Organismal Development,Tissue Development** | Development of spiny stellate neurons | 0.0348 | SLC17A7 | 1 |
| **Organismal Functions** | Recovery of mice | 0.0348 | FAS | 1 |
| **Endocrine System Disorders** | Leptin resistance | 0.0348 | SH2B1 | 1 |
| **Cell Morphology,Cell-To-Cell Signaling and Interaction,Cellular Assembly and Organization** | Length of gap junctions | 0.0348 | NRCAM | 1 |
| **Cell-To-Cell Signaling and Interaction,Drug Metabolism,Small Molecule Biochemistry** | Binding of dopamine | 0.0348 | HTT | 1 |

| | | | | |
|---|---|---|---|---|
| **DNA Replication, Recombination, and Repair,Molecular Transport,Nucleic Acid Metabolism,Small Molecule Biochemistry** | Quantity of adenosine | 0.0348 | HTT | 1 |
| **Nervous System Development and Function** | Modification of striatum | 0.0348 | HTT | 1 |
| **Cell Morphology,Cellular Assembly and Organization,Cellular Function and Maintenance,Nervous System Development and Function** | Extension of parallel fiber | 0.0348 | NFIX | 1 |
| **Neurological Disease,Organismal Injury and Abnormalities** | Astrocytosis of forebrain | 0.0348 | HTT | 1 |
| **Carbohydrate Metabolism,Drug Metabolism,Molecular Transport,Small Molecule Biochemistry** | Accumulation of hyaluronic acid | 0.0348 | HEXB | 1 |
| **Metabolic Disease** | Hyperleptinemia | 0.0348 | SH2B1 | 1 |
| **Cell Morphology,Cell-To-Cell Signaling and Interaction,Cellular Function and Maintenance,Nervous System Development and Function** | Depolarization of medium spiny neurons | 0.0348 | HTT | 1 |
| **Behavior,Nervous System Development and Function** | Contextual fear memory retrieval | 0.0348 | NPY | 1 |
| **Cell-To-Cell Signaling and Interaction,Nervous System Development and Function** | Synaptic transmission of thalamocortical neurons | 0.0348 | SLC17A7 | 1 |
| **Infectious Diseases,Neurological Disease** | Infection of ependyma | 0.0348 | TICAM1 | 1 |
| **Cellular Assembly and Organization** | Binding of vesicles | 0.0348 | HTT | 1 |
| **Cell-To-Cell Signaling and Interaction,Nervous System Development and Function** | Neurotransmission of corticostriatal neurons | 0.0348 | HTT | 1 |
| **Neurological Disease,Organismal Injury and Abnormalities** | Degeneration of entorhinal area | 0.0348 | HTT | 1 |
| **Neurological Disease,Organismal Injury and Abnormalities,Psychological Disorders** | Preclinical stage amyotrophic lateral sclerosis | 0.0348 | FAS | 1 |
| **Cell Death and Survival,Cellular Compromise,Nervous System Development and Function** | Cytotoxicity of medium spiny neurons | 0.0348 | HTT | 1 |
| **Reproductive System Development and Function** | Estrous cyclicity | 0.0348 | GJD2 | 1 |
| **Cell Death and Survival,Cellular Compromise,Neurological Disease,Organismal Injury and Abnormalities,Psychological Disorders,Tissue Morphology** | Damage of medium spiny neurons | 0.0348 | HTT | 1 |
| **Cell Death and Survival,Neurological Disease,Organismal Injury and Abnormalities** | Apoptosis of lactotropes | 0.0348 | FAS | 1 |
| **Nervous System Development and Function,Neurological Disease,Organ Morphology,Organismal Development,Organismal Injury and Abnormalities,Tissue Morphology** | Abnormal morphology of cingulum | 0.0348 | MAP3K12 | 1 |
| **Cellular Assembly and Organization,Cellular Movement** | Segregation of retinogeniculate projection | 0.0348 | GJD2 | 1 |
| **Endocrine System Disorders,Nervous System Development and Function,Neurological Disease,Organ Development,Organismal Injury and Abnormalities** | Neurodegeneration of hypothalamus | 0.0348 | HTT | 1 |
| **Cellular Growth and Proliferation,Tissue Development** | Proliferation of neuroepithelial cells | 0.0348 | HTT | 1 |
| **Cell Morphology,Cellular Compromise,Nervous System Development and Function** | Shrinkage of striatal neurons | 0.0348 | HTT | 1 |
| **Nervous System Development and Function,Neurological Disease,Organ Morphology,Organismal Development,Organismal Injury and Abnormalities** | Abnormal morphology of enlarged cerebellum | 0.0348 | OTX1 | 1 |
| **Cellular Assembly and Organization,Nervous System Development and Function,Tissue Morphology** | Quantity of neuropil inclusions | 0.0348 | HTT | 1 |
| **Embryonic Development,Organismal Development,Tissue Development** | Formation of preplate | 0.0348 | FOXG1 | 1 |
| **Neurological Disease,Organ Morphology,Organismal Injury and Abnormalities** | Atrophy of cerebral neocortex | 0.0348 | HTT | 1 |
| **Cell Death and Survival,DNA Replication, Recombination, and Repair** | Fragmentation of DNA | 0.0351 | ADCYAP1, HTT | 2 |

| | | | | |
|---|---|---|---|---|
| **Cell Death and Survival** | Quantity of apoptotic cells | 0.0351 | CAPNS1, HTT | 2 |
| **Cell Death and Survival,Cell Morphology,Nervous System Development and Function,Neurological Disease,Organ Morphology,Organismal Development,Organismal Injury and Abnormalities,Tissue Morphology** | Loss of Purkinje cells | 0.0351 | HTT, MAP1A | 2 |
| **Cellular Movement** | Migration of cells | 0.0354 | ADCYAP1, CAMKK2, CCL4, CELSR2, FAS, FOXG1, HTT, NFIX, SRF, TIAM2 | 10 |
| **Neurological Disease** | Movement Disorders | 0.0382 | ACTN2, ATP6V1A, CAMKK2, CCL4, FAS, FOXG1, HTT, OXR1, PPP3CA, RPH3A, SLC17A7, THBS3, ZMYND8 | 13 |
| **Cell Death and Survival** | Cell viability of brain cells | 0.0401 | ADCYAP1, FOXG1, HTT, NRCAM, NRN1 | 5 |
| **Cell Morphology,Nervous System Development and Function,Neurological Disease,Tissue Morphology** | Abnormal morphology of neurites | 0.0415 | CAMKK2, HTT, MAP1A, NRCAM, NRN1, PPP3CA, SLC17A7 | 7 |
| **Nervous System Development and Function,Organ Morphology,Organismal Development** | Morphology of cerebellum | 0.0425 | ADCYAP1, HEXB, HTT, IGFBP6, MAP1A, NRCAM, OTX1 | 7 |
| **Cell Morphology,Cellular Assembly and Organization,Cellular Development,Cellular Function and Maintenance,Cellular Growth and Proliferation,Embryonic Development,Nervous System Development and Function,Neurological Disease,Organismal Development,Tissue Development,Tissue Morphology** | Loss of dendritic spines | 0.0431 | CAMKK2, HTT | 2 |
| **Cell Signaling,Nucleic Acid Metabolism,Small Molecule Biochemistry** | Synthesis of cyclic AMP | 0.0431 | ADCYAP1, HTT | 2 |
| **Embryonic Development,Nervous System Development and Function,Organ Development,Organismal Development,Tissue Development** | Development of cerebral cortex | 0.0436 | FOXG1, FOXP2, HTT, MKKS, OTX1, SH2B1, SRF | 7 |
| **Cell-To-Cell Signaling and Interaction,Cellular Assembly and Organization** | Cell-cell contact | 0.0437 | CAMK2A, CDH9, GABBR2, GJD2, HTT, NECTIN1, NRCAM, NRN1, SEZ6L2, SNTB2 | 10 |
| **Nervous System Development and Function,Tissue Morphology** | Quantity of nervous tissue | 0.0455 | ADCYAP1, CAMK2A, GABBR2, GSDME, HTT, MAP3K12, NAP1L2, NRCAM, NRN1, OTX1, PRDM1, RB1CC1, | 14 |

| | | | | |
|---|---|---|---|---|
| | | | SHC2, SLC17A7 | |
| **Cellular Movement** | Cell movement of brain cells | 0.0472 | ADCYAP1, CELSR2, FOXG1, NRCAM, TIAM2 | 5 |
| **Lipid Metabolism,Molecular Transport,Small Molecule Biochemistry** | Concentration of lipid | 0.0472 | ADCYAP1, ARNTL, FAS, HTT, NPY | 5 |
| **Nervous System Development and Function,Organ Morphology,Organismal Development** | Volume of brain | 0.0473 | ADCYAP1, HTT | 2 |
| **Cell Death and Survival,Cellular Compromise,Neurological Disease,Organismal Injury and Abnormalities,Tissue Morphology** | Neurodegeneration of Purkinje cells | 0.0492 | HEXB, HTT, MAP1A | 3 |

### Cluster 15

| Categories | Diseases or Functions Annotation | p-Value | Molecules | # Molecules |
|---|---|---|---|---|
| **Cellular Development,Cellular Growth and Proliferation,Embryonic Development,Nervous System Development and Function,Organ Development,Organismal Development,Tissue Development** | Expansion of neural precursor cells | 0.000324 | AHR, FOXG1 | 2 |
| **Cell Morphology,Cellular Development,Embryonic Development,Nervous System Development and Function,Tissue Development** | Patterning of dendrites | 0.00313 | CELSR2, CRMP1 | 2 |
| **Cellular Function and Maintenance** | Endocytosis | 0.00478 | BECN1, KIAA1107, PPP3CA, RAC1, SCAMP5 | 5 |
| **Cellular Assembly and Organization,Cellular Movement,Nervous System Development and Function** | Transport of synaptic vesicles | 0.0154 | KIAA1107, PFN2, PPP3CA, RPH3A, SCAMP5 | 5 |
| **Cell Morphology,Cellular Assembly and Organization,Cellular Development,Cellular Function and Maintenance,Cellular Growth and Proliferation,Embryonic Development,Nervous System Development and Function,Organismal Development,Tissue Development** | Arrest in sprouting of neurites | 0.018 | PFN2 | 1 |
| **Cellular Assembly and Organization** | Aggregation of early endosomes | 0.018 | KIF16B | 1 |
| **Endocrine System Development and Function,Organ Morphology,Organismal Development** | Enlargement of pituitary gland | 0.018 | PTTG1 | 1 |
| **Connective Tissue Development and Function,Organismal Development,Skeletal and Muscular System Development and Function,Tissue Development** | Size of cranium | 0.018 | MYB | 1 |
| **Cell Cycle** | Aneuploidy of cells | 0.018 | PTTG1 | 1 |
| **Embryonic Development,Nervous System Development and Function,Organ Development,Organismal Development,Tissue Development** | Formation of subgranular zone of dentate gyrus | 0.018 | FOXG1 | 1 |
| **Cellular Assembly and Organization,Cellular Function and Maintenance** | Endocytosis of plasma membrane | 0.018 | RAC1 | 1 |
| **Cellular Movement,Nervous System Development and Function** | Delay in initiation of migration of cortical projection neurons | 0.018 | RAC1 | 1 |
| **Cellular Compromise** | Acidification of cytosol | 0.018 | CA4 | 1 |
| **Cellular Assembly and Organization,Cellular Function and Maintenance** | Quantity of peripheral lamellae | 0.018 | RAC1 | 1 |
| **Cell Cycle,Cell Morphology,Cellular Assembly and Organization,Cellular Function and Maintenance** | Length of telomeres | 0.018 | PTTG1 | 1 |
| **Cell-To-Cell Signaling and Interaction,Cellular Assembly and Organization,Cellular** | Differentiation of inhibitory synapse | 0.018 | CLSTN3 | 1 |

| | | | | |
|---|---|---|---|---|
| **Development,Cellular Function and Maintenance,Cellular Growth and Proliferation,Nervous System Development and Function,Tissue Development** | | | | |
| **Cell Morphology,Cellular Assembly and Organization,Nervous System Development and Function** | Arrest in elongation of neurites | 0.018 | PFN2 | 1 |
| **Cellular Assembly and Organization,Cellular Development,Embryonic Development,Nervous System Development and Function,Organ Development,Organismal Development,Tissue Development** | Maturation of olfactory glomeruli | 0.018 | PPP3CA | 1 |
| **Nervous System Development and Function,Neurological Disease,Organ Morphology,Organismal Development,Organismal Injury and Abnormalities** | Abnormal morphology of stratum radiatum | 0.018 | CLSTN3 | 1 |
| **Cell-To-Cell Signaling and Interaction,Cellular Assembly and Organization,Cellular Development,Cellular Function and Maintenance,Cellular Growth and Proliferation,Nervous System Development and Function,Tissue Development** | Differentiation of excitatory synapses | 0.018 | CLSTN3 | 1 |
| **Cell-To-Cell Signaling and Interaction,Nervous System Development and Function** | Synaptic transmission of inhibitory synapse | 0.018 | CLSTN3 | 1 |
| **Cellular Development,Cellular Growth and Proliferation,Nervous System Development and Function,Tissue Development** | Development of Cajal-Retzius neurons | 0.018 | FOXG1 | 1 |
| **Cell-To-Cell Signaling and Interaction,Nervous System Development and Function** | Activation of superior cervical ganglion neurons | 0.018 | SYBU | 1 |
| **Cellular Development,Cellular Growth and Proliferation,Embryonic Development,Nervous System Development and Function,Organ Development,Organismal Development,Tissue Development** | Expansion of telencephalic progenitor cells | 0.018 | FOXG1 | 1 |
| **Cell Morphology,Skeletal and Muscular System Development and Function** | Hypercontractility of vascular smooth muscle cells | 0.018 | SRF | 1 |
| **Endocrine System Development and Function,Small Molecule Biochemistry** | Hydroxylation of melatonin | 0.018 | CYP1B1 | 1 |
| **Cell Morphology,Cellular Assembly and Organization,Nervous System Development and Function,Tissue Morphology** | Size of synaptic vesicles pool | 0.018 | SCAMP5 | 1 |
| **Lipid Metabolism,Molecular Transport,Small Molecule Biochemistry** | Concentration of eicosapentenoic acid | 0.018 | ARNTL | 1 |
| **Neurological Disease** | Formation of myelin ovoid | 0.018 | RAC1 | 1 |
| **Inflammatory Response,Organismal Injury and Abnormalities,Renal and Urological Disease** | Inflammation of urinary tract | 0.018 | PTTG1 | 1 |
| **Cell-To-Cell Signaling and Interaction,Nervous System Development and Function** | Synaptic transmission of excitatory synapses | 0.018 | CLSTN3 | 1 |
| **Cell Morphology,Cellular Assembly and Organization,Nervous System Development and Function** | Projection of corticothalamic axons | 0.018 | RAC1 | 1 |
| **Infectious Diseases** | Infection by human herpesvirus 2 | 0.018 | NECTIN1 | 1 |
| **Cell Morphology,Nervous System Development and Function,Neurological Disease,Organ Morphology,Organismal Development,Organismal Injury and Abnormalities,Tissue Morphology** | Abnormal morphology of Purkinje cells | 0.0282 | ALS2, HEXB, MAP1A, SPARCL1 | 4 |
| **Cellular Assembly and Organization,Cellular Function and Maintenance,Cellular Movement,Nervous System Development and Function** | Endocytosis of synaptic vesicles | 0.0287 | KIAA1107, PPP3CA, SCAMP5 | 3 |
| **Cellular Movement,Nervous System Development and Function** | Migration of granule cells | 0.0307 | CELSR2, FOXG1, RAC1 | 3 |
| **Cell Death and Survival,Neurological Disease,Organismal Injury and Abnormalities** | Apoptosis of lactotropes | 0.0358 | FAS | 1 |
| **Cellular Development,Cellular Growth and Proliferation,Embryonic Development,Nervous System Development and Function,Organ** | Development of spiny stellate neurons | 0.0358 | SLC17A7 | 1 |

| | | | | |
|---|---|---|---|---|
| **Development,Organismal Development,Tissue Development** | | | | |
| **Organismal Functions** | Recovery of mice | 0.0358 | FAS | 1 |
| **Nervous System Development and Function,Organ Morphology,Tissue Morphology** | Quantity of mossy fiber cells | 0.0358 | RAC1 | 1 |
| **Cell Morphology,Cell-To-Cell Signaling and Interaction,Cellular Assembly and Organization** | Length of gap junctions | 0.0358 | NRCAM | 1 |
| **Nervous System Development and Function,Organ Morphology,Organismal Development** | Thickness of granule cell layer | 0.0358 | NRCAM | 1 |
| **Cellular Movement,Nervous System Development and Function** | Distribution of interneurons | 0.0358 | DCK | 1 |
| **Carbohydrate Metabolism,Drug Metabolism,Molecular Transport,Small Molecule Biochemistry** | Accumulation of hyaluronic acid | 0.0358 | HEXB | 1 |
| **Cell Death and Survival,Neurological Disease,Organismal Injury and Abnormalities** | Apoptosis of somatotrophs | 0.0358 | FAS | 1 |
| **Cell-To-Cell Signaling and Interaction,Cellular Assembly and Organization,Nervous System Development and Function,Tissue Morphology** | Density of inhibitory synapse | 0.0358 | CLSTN3 | 1 |
| **Cellular Development,Cellular Growth and Proliferation,Nervous System Development and Function,Tissue Development** | Generation of Cajal-Retzius neurons | 0.0358 | FOXG1 | 1 |
| **Cell-To-Cell Signaling and Interaction,Nervous System Development and Function** | Synaptic transmission of thalamocortical neurons | 0.0358 | SLC17A7 | 1 |
| **Cellular Assembly and Organization,Nervous System Development and Function** | Delay in sorting of axons | 0.0358 | RAC1 | 1 |
| **Embryonic Development,Organismal Development,Tissue Development** | Formation of preplate | 0.0358 | FOXG1 | 1 |
| **Cell Morphology,Cellular Assembly and Organization,Nervous System Development and Function** | Extension of leading processes | 0.0358 | RAC1 | 1 |
| **Neurological Disease,Organismal Injury and Abnormalities,Psychological Disorders** | Preclinical stage amyotrophic lateral sclerosis | 0.0358 | FAS | 1 |
| **Cell-To-Cell Signaling and Interaction,Cellular Assembly and Organization,Cellular Function and Maintenance** | Formation of spot-like adhesion site | 0.0358 | NECTIN1 | 1 |
| **Cell-To-Cell Signaling and Interaction,Nervous System Development and Function** | Synaptic depression of superior cervical ganglion neurons | 0.0358 | SYBU | 1 |
| **Cell Death and Survival,Cell Morphology,Nervous System Development and Function,Neurological Disease,Organ Morphology,Organismal Development,Organismal Injury and Abnormalities,Tissue Morphology** | Loss of Purkinje cells | 0.0369 | ALS2, MAP1A | 2 |
| **Nervous System Development and Function,Neurological Disease,Organ Morphology,Organismal Development,Organismal Injury and Abnormalities** | Abnormal morphology of cerebellar cortex | 0.0372 | ALS2, HEXB, MAP1A, NRCAM, SPARCL1 | 5 |
| **Cell Morphology,Cellular Compromise** | Collapse of growth cone | 0.0375 | NRCAM, RAC1, SEMA4F, SRF | 4 |
| **Cellular Development,Cellular Growth and Proliferation,Embryonic Development,Nervous System Development and Function,Organ Development,Organismal Development,Tissue Development** | Proliferation of neural precursor cells | 0.0446 | AHR, CUX2, FOXG1, ID2 | 4 |
| **Neurological Disease** | Movement Disorders | 0.0459 | ACTN2, ALS2, CACNB3, CCL4, DDX42, FAS, FOXG1, PFN2, PPP3CA, RPH3A | 13 |
| **Cell Morphology,Nervous System Development and Function,Neurological Disease,Tissue Morphology** | Abnormal morphology of neurites | 0.0469 | MAP1A, NRCAM, NRN1, PFN2, | 7 |

| | | | | |
|---|---|---|---|---|
| | | | PPP3CA, SCYL1, SLC17A7 | |
| **Neurological Disease,Organismal Injury and Abnormalities** | Hydrocephalus | 0.0497 | CELSR2, MYB | 2 |

### Supplementary Table 9: Module 9
### Cluster 1

| Categories | Diseases or Functions Annotation | p-Value | Molecules | # Molecules |
|---|---|---|---|---|
| **Cellular Function and Maintenance,Molecular Transport** | Exocytosis | 6.16E-07 | APBA2, CADPS2, ERC2, PIP5K1C, PPFIA3, RIMS2, RPH3A, UNC13A, UNC13B, UNC13C | 10 |
| **Cell-To-Cell Signaling and Interaction,Cellular Assembly and Organization,Cellular Function and Maintenance,Molecular Transport** | Priming of synaptic vesicles | 0.0000108 | CADPS2, ERC2, RIMS2, UNC13A, UNC13B, UNC13C | 6 |
| **Cellular Assembly and Organization,Cellular Function and Maintenance,Cellular Movement,Molecular Transport,Nervous System Development and Function** | Exocytosis of synaptic vesicles | 0.0000193 | APBA2, PIP5K1C, PPFIA3, RPH3A, UNC13A, UNC13B, UNC13C | 7 |
| **Cellular Assembly and Organization,Cellular Movement,Nervous System Development and Function** | Transport of synaptic vesicles | 0.0000652 | APBA2, KIAA1107, PCLO, PIP5K1C, PPFIA3, RPH3A, SCRIB, UNC13A, UNC13B, UNC13C | 10 |
| **Cancer,Neurological Disease,Organismal Injury and Abnormalities** | Brain astrocytoma | 0.000244 | ABCF1, ADAMTS13, ADAMTS8, AGAP3, ANKRD30B, ASPM, ASXL3, ATF7IP2, CREBBP, DNAH8, ERC2, GATAD2B, HERC1, MYO5B, P2RX5, PCLO, PIK3R1, PRDM2, PTGS2, RAB11FIP5, RALY, RIMS2, ROS1, SEMA4F, SPTBN4, SYNE1, THEMIS, TSC2 | 28 |
| **Cell Morphology,Cellular Assembly and Organization,Cellular Development,Cellular** | Neuritogenesis | 0.000337 | ABI1, ADCY1, | 38 |

| | | | | |
|---|---|---|---|---|
| **Function and Maintenance,Cellular Growth and Proliferation,Nervous System Development and Function,Organismal Development,Tissue Development** | | | ADGRB3, AGRN, ANAPC2, ARHGEF28, ATCAY, BCL11B, BSN, CAMSAP1, CELSR2, CRMP1, DAB2IP, DBN1, DVL1, ELK1, EPHB3, ERC2, FMN1, GABRB3, GAP43, GPRIN1, HERC1, MRTFA, MYO16, MYO5B, NIN, NRN1, PIP5K1C, PTGS2, ROCK2, SHANK2, SORBS2, SPTBN4, SYNE1, TIAM1, TOP2B, TSC2 | |
| **Cellular Assembly and Organization,Cellular Function and Maintenance** | Microtubule dynamics | 0.0004 | ABI1, ADCY1, ADCY6, ADGRB3, AGRN, ANAPC2, ARHGEF28, ATCAY, BCL11B, BSN, CAMSAP1, CELSR2, CRMP1, DAB2IP, DBN1, DVL1, ELK1, EPHB3, ERC2, FMN1, GABRB3, GAP43, GPRIN1, HERC1, MRTFA, MYO16, MYO5B, NIN, NRN1, PIP5K1C, PTGS2, ROCK2, SCN1B, SEMA4F, SEMA5B, SHANK2, SORBS2, SPTBN4, SYNE1, TIAM1, TOP2B, TSC2 | 42 |
| **Cellular Assembly and Organization,Nervous System Development and Function** | Docking of synaptic vesicles | 0.000432 | PPFIA3, RIMS2, | 4 |

| | | | | |
|---|---|---|---|---|
| | | | UNC13A, UNC13B | |
| **Nervous System Development and Function,Neurological Disease** | Abnormal morphology of nervous system | 0.000453 | ABR, ADCY1, ADGRA2, AGRN, ARNT2, ASPM, BRCA2, CADPS2, CELSR2, COL13A1, CREBBP, CRMP1, DBN1, DVL1, EPHB3, ERC2, FAT3, GABRB3, GAP43, HIVEP2, IDUA, KIF1B, LRRTM4, MAN2C1, MCPH1, MEGF8, MTHFR, MYO16, NCOR1, NOS2, NRN1, PER2, PRKN, RIMS2, ROCK2, SCN1B, SEMA5B, SHANK2, SORBS2, SORL1, SPTBN4, SYNE1, TOP2B, UNC13A, UNC13B, VAX1 | 46 |
| **Cellular Development,Cellular Growth and Proliferation,Nervous System Development and Function,Tissue Development** | Development of neurons | 0.000601 | ABI1, ADCY1, ADGRB3, AGRN, ANAPC2, ARHGEF28, ATCAY, BCL11B, BSN, CADPS2, CAMSAP1, CELSR2, CRMP1, DAB2IP, DBN1, DVL1, ELK1, EPHB3, ERC2, FMN1, GABRB3, GAP43, GPRIN1, HERC1, LRRTM4, MRTFA, MYO16, MYO5B, NIN, NRN1, PCDH8, PIP5K1C, PLXND1, PRDM1, PTGS2, ROCK2, | 43 |

| | | | | |
|---|---|---|---|---|
| | | | SHANK2, SORBS2, SPTBN4, SYNE1, TIAM1, TOP2B, TSC2 | |
| **Nervous System Development and Function,Tissue Morphology** | Density of neurons | 0.000631 | ACVR1C, CRMP1, DAB2IP, ELK1, EPHB3, GABRB3, GAP43, LRRTM4, PTGS2, SCN1B, TIAM1, TSC2 | 12 |
| **Cellular Assembly and Organization,Cellular Function and Maintenance** | Formation of Golgi vesicles | 0.000737 | BSN, PCLO | 2 |
| **Cellular Assembly and Organization** | Translocation of Golgi vesicles | 0.000737 | BSN, PCLO | 2 |
| **Nervous System Development and Function** | Morphology of nervous system | 0.000883 | ABR, ADCY1, ADGRA2, AGRN, ARNT2, ASPM, BRCA2, CADPS2, CELSR2, COL13A1, CREBBP, CRMP1, DBN1, DVL1, EPHB3, ERC2, FAT3, GABRB3, GAP43, HIVEP2, IDUA, KIF1B, KIF5C, KNG1, LRRTM4, MAN2C1, MCPH1, MEGF8, MTHFR, MYO16, NCOR1, NOS2, NRN1, PER2, PLAGL1, PRDM16, PRKN, RGS11, RIMS2, ROCK2, SCN1B, SEMA5B, SHANK2, SORBS2, SORL1, SPTBN4, SYNE1, TIAM1, TOP2B, UNC13A, UNC13B, VAX1 | 52 |
| **Behavior,Nervous System Development and Function** | Short-term memory | 0.00101 | CREBBP, KCNH3, NR3C2 | 3 |
| **Cellular Assembly and Organization,Nervous System Development and Function,Tissue Morphology** | Abnormal quantity of synaptic vesicles | 0.00117 | ERC2, KIF1B, PCLO, | 5 |

| | | | | |
|---|---|---|---|---|
| | | | UNC13A, UNC13B | |
| **Cancer,Organismal Injury and Abnormalities** | Non-hematological solid tumor | 0.00124 | ABCF1, ADAMTS13, ADAMTS8, AGAP3, ANKRD30B, ASPM, ASXL3, ATF7IP2, CREBBP, CUL9, DNAH8, ERC2, GATAD2B, HERC1, MYO15A, MYO5B, P2RX5, PCLO, PIGQ, PIK3R1, PRDM1, PRDM2, PTGS2, RAB11FIP5, RALY, RIMS2, ROS1, SEMA4F, SPTBN4, SYNE1, THEMIS, TSC2, UNC13B | 33 |
| **Cancer,Organismal Injury and Abnormalities** | Head and neck tumor | 0.00124 | ABCF1, ADAMTS13, ADAMTS8, AGAP3, ANKRD30B, ASPM, ASXL3, ATF7IP2, CREBBP, CUL9, DNAH8, ERC2, GATAD2B, HERC1, MYO15A, MYO5B, P2RX5, PCLO, PIK3R1, PRDM1, PRDM2, PTGS2, RAB11FIP5, RALY, RIMS2, ROS1, SEMA4F, SPTBN4, SYNE1, THEMIS, TSC2, UNC13B | 32 |
| **Cell Morphology,Cell-To-Cell Signaling and Interaction,Cellular Assembly and Organization,Nervous System Development and Function,Tissue Morphology** | Morphology of synapse | 0.00147 | AGRN, COL13A1, ERC2, GAP43, NRN1, ROCK2, SYNE1, UNC13A, UNC13B | 9 |

| | | | | |
|---|---|---|---|---|
| **Lipid Metabolism,Molecular Transport,Small Molecule Biochemistry** | Concentration of arachidonic acid | 0.00148 | ARNTL, KNG1, PTGS2 | 3 |
| **Cell-To-Cell Signaling and Interaction,Cellular Assembly and Organization** | Cell-cell contact | 0.00178 | AGRN, BSN, COL13A1, EPHB3, ERC2, GAP43, GJD2, LRRTM4, NEURL1, NRN1, PCDH8, PLXND1, ROCK2, SHANK2, SORBS2, SYNE1, UNC13A, UNC13B | 18 |
| **Cancer,Neurological Disease,Organismal Injury and Abnormalities** | Brain glioma | 0.00182 | ABCF1, ADAMTS13, ADAMTS8, AGAP3, ANKRD30B, ASPM, ASXL3, ATF7IP2, CREBBP, DNAH8, ERC2, GATAD2B, HERC1, MYO15A, MYO5B, P2RX5, PCLO, PIK3R1, PRDM1, PRDM2, PTGS2, RAB11FIP5, RALY, RIMS2, ROS1, SEMA4F, SPTBN4, SYNE1, THEMIS, TSC2 | 30 |
| **Neurological Disease,Organismal Injury and Abnormalities** | Hydrocephalus | 0.00184 | CELSR2, DVL1, DVL3, NCOR1 | 4 |
| **Cell Morphology,Cellular Assembly and Organization,Cellular Development,Cellular Function and Maintenance,Cellular Growth and Proliferation,Embryonic Development,Nervous System Development and Function,Organismal Development,Tissue Development** | Dendritic growth/branching | 0.00192 | ABI1, AGRN, BSN, CRMP1, DAB2IP, DBN1, ELK1, EPHB3, GABRB3, GAP43, MYO5B, NRN1, PTGS2, ROCK2, SHANK2, SORBS2, TIAM1, TSC2 | 18 |
| **Cancer,Organismal Injury and Abnormalities** | Non-melanoma solid tumor | 0.00195 | ABCF1, ADAMTS13, ADAMTS8, AGAP3, ANKRD30B, ASPM, ASXL3, | 31 |

| | | | | |
|---|---|---|---|---|
| | | | ATF7IP2, CREBBP, DNAH8, ERC2, GATAD2B, HERC1, MYO15A, MYO5B, P2RX5, PCLO, PIK3R1, PRDM1, PRDM2, PTGS2, RAB11FIP5, RALY, RIMS2, ROS1, SEMA4F, SPTBN4, SYNE1, THEMIS, TSC2, UNC13B | |
| **Cancer,Neurological Disease,Organismal Injury and Abnormalities** | Brain tumor | 0.002 | ABCF1, ADAMTS13, ADAMTS8, AGAP3, ANKRD30B, ASPM, ASXL3, ATF7IP2, CREBBP, DNAH8, ERC2, GATAD2B, HERC1, MYO15A, MYO5B, P2RX5, PCLO, PIK3R1, PRDM1, PRDM2, PTGS2, RAB11FIP5, RALY, RIMS2, ROS1, SEMA4F, SPTBN4, SYNE1, THEMIS, TSC2, UNC13B | 31 |
| **Cancer,Neurological Disease,Organismal Injury and Abnormalities** | Nervous system neoplasm | 0.00213 | ABCF1, ADAMTS13, ADAMTS8, AGAP3, ANKRD30B, ASPM, ASXL3, ATF7IP2, CREBBP, DNAH8, ERC2, GATAD2B, HERC1, MYO15A, MYO5B, P2RX5, PCLO, PIGQ, PIK3R1, PRDM1, PRDM2, | 32 |

| | | | | |
|---|---|---|---|---|
| | | | PTGS2, RAB11FIP5, RALY, RIMS2, ROS1, SEMA4F, SPTBN4, SYNE1, THEMIS, TSC2, UNC13B | |
| **Hereditary Disorder,Neurological Disease,Organismal Injury and Abnormalities,Psychological Disorders,Skeletal and Muscular Disorders** | Familial Parkinson disease | 0.00217 | GAP43, PRKN | 2 |
| **Organismal Development,Organismal Injury and Abnormalities** | Abnormal morphology of head | 0.00279 | ABR, ADCY1, ADGRA2, ADGRV1, ARNT2, ASPM, CADPS2, CELSR2, CREBBP, CRMP1, DBN1, DVL1, EPHB3, FAT3, GABRB3, GAP43, HIVEP2, IDUA, KIF1B, LRRTM4, MAN2C1, MCPH1, MTHFR, NCOR1, NOS2, PRKN, SHANK2, SPTBN4, TOP2B, VAX1 | 30 |
| **Cell Morphology,Cell-To-Cell Signaling and Interaction,Cellular Assembly and Organization,Nervous System Development and Function,Neurological Disease,Tissue Morphology** | Abnormal morphology of synapse | 0.00286 | AGRN, COL13A1, ERC2, NRN1, ROCK2, SYNE1, UNC13A, UNC13B | 8 |
| **Nervous System Development and Function,Neurological Disease** | Abnormal morphology of central nervous system | 0.00362 | ABR, ADCY1, ADGRA2, ARNT2, ASPM, CADPS2, CELSR2, CREBBP, CRMP1, DBN1, EPHB3, GABRB3, GAP43, HIVEP2, IDUA, KIF1B, LRRTM4, MAN2C1, MCPH1, MTHFR, NCOR1, NOS2, PRKN, RIMS2, SCN1B, SHANK2, SPTBN4, | 29 |

| | | | | |
|---|---|---|---|---|
| | | | TOP2B, VAX1 | |
| **Auditory and Vestibular System Development and Function,Cell Morphology,Cellular Assembly and Organization** | Orientation of stereocilia bundles | 0.00365 | ADGRV1, DVL1, DVL3 | 3 |
| **Cell Morphology,Nervous System Development and Function,Tissue Morphology** | Morphology of bipolar cells | 0.0037 | ERC2, FAT3, RGS11, SEMA5B | 4 |
| **Organismal Development** | Morphology of head | 0.00388 | ABR, ADCY1, ADGRA2, ADGRV1, ARNT2, ASPM, CADPS2, CELSR2, CREBBP, CRMP1, DBN1, DVL1, EPHB3, FAT3, GABRB3, GAP43, HIVEP2, IDUA, KIF1B, KIF5C, KNG1, LRRTM4, MAN2C1, MCPH1, MTHFR, MYO16, NCOR1, NOS2, PRDM16, PRKN, SHANK2, SPTBN4, TOP2B, VAX1 | 34 |
| **Cellular Assembly and Organization** | Quantity of vesicles | 0.00399 | BSN, ERC2, KIF1B, PCLO, UNC13A, UNC13B | 6 |
| **Behavior,Nervous System Development and Function** | Working memory | 0.00426 | KCNH3, NR3C2 | 2 |
| **Nervous System Development and Function,Neurological Disease,Organ Morphology,Organismal Development,Organismal Injury and Abnormalities** | Abnormal morphology of brain | 0.00453 | ABR, ADCY1, ADGRA2, ARNT2, ASPM, CADPS2, CELSR2, CREBBP, CRMP1, DBN1, EPHB3, GABRB3, GAP43, HIVEP2, IDUA, KIF1B, LRRTM4, MAN2C1, MCPH1, MTHFR, NCOR1, NOS2, PRKN, SHANK2, SPTBN4, TOP2B, VAX1 | 27 |
| **Nervous System Development and Function,Neurological Disease,Organ Morphology,Organismal** | Abnormal morphology of barrel cortex | 0.00465 | ADCY1, ASPM, GAP43 | 3 |

| | | | | |
|---|---|---|---|---|
| **Development,Organismal Injury and Abnormalities** | | | | |
| **Neurological Disease,Organismal Injury and Abnormalities** | Brain lesion | 0.00491 | ABCF1, ADAMTS13, ADAMTS8, AGAP3, ANKRD30B, ASPM, ASXL3, ATF7IP2, CREBBP, DNAH8, ERC2, GATAD2B, HERC1, KNG1, MYO15A, MYO5B, P2RX5, PCLO, PIK3R1, PRDM1, PRDM2, PTGS2, RAB11FIP5, RALY, RIMS2, ROS1, SEMA4F, SPTBN4, SYNE1, THEMIS, TSC2, UNC13B | 32 |
| **Nervous System Development and Function** | Morphology of central nervous system | 0.00559 | ABR, ADCY1, ADGRA2, ARNT2, ASPM, CADPS2, CELSR2, CREBBP, CRMP1, DBN1, EPHB3, GABRB3, GAP43, HIVEP2, IDUA, KIF1B, KIF5C, KNG1, LRRTM4, MAN2C1, MCPH1, MTHFR, MYO16, NCOR1, NOS2, PRDM16, PRKN, RIMS2, SCN1B, SHANK2, SPTBN4, TOP2B, VAX1 | 33 |
| **Nervous System Development and Function,Organ Morphology,Organismal Development** | Morphology of brain | 0.00568 | ABR, ADCY1, ADGRA2, ARNT2, ASPM, CADPS2, CELSR2, CREBBP, CRMP1, DBN1, EPHB3, | 31 |

| | | | | |
|---|---|---|---|---|
| | | | GABRB3, GAP43, HIVEP2, IDUA, KIF1B, KIF5C, KNG1, LRRTM4, MAN2C1, MCPH1, MTHFR, MYO16, NCOR1, NOS2, PRDM16, PRKN, SHANK2, SPTBN4, TOP2B, VAX1 | |
| **Cell Morphology** | Shape change of neurites | 0.00635 | ABI1, AGRN, BCL11B, BSN, CRMP1, DAB2IP, DBN1, ELK1, EPHB3, GABRB3, GAP43, MYO5B, NR3C2, NRN1, PTGS2, ROCK2, SHANK2, SORBS2, TIAM1, TSC2 | 20 |
| **Cell Morphology,Nervous System Development and Function,Neurological Disease,Tissue Morphology** | Abnormal morphology of neurons | 0.0066 | AGRN, BRCA2, CADPS2, COL13A1, CRMP1, DBN1, DVL1, EPHB3, ERC2, FAT3, GAP43, HIVEP2, IDUA, MYO16, NOS2, NRN1, PRKN, ROCK2, SCN1B, SEMA5B, SHANK2, SORBS2, SORL1, SPTBN4, SYNE1, UNC13A, UNC13B | 27 |
| **Cell Morphology,Cellular Development,Embryonic Development,Nervous System Development and Function,Tissue Development** | Patterning of dendrites | 0.00698 | CELSR2, CRMP1 | 2 |
| **Cell Morphology,Cellular Function and Maintenance** | Autophagy of cortical neurons | 0.00698 | PRKN, TSC2 | 2 |
| **Cell Morphology,Nervous System Development and Function** | Innervation of optic nerve | 0.00698 | GAP43, VAX1 | 2 |
| **Behavior** | Learning | 0.00938 | ADCY1, ADGRB3, CREBBP, GABRB3, KCNH3, NEURL1, NR3C2 | 7 |

| | | | | |
|---|---|---|---|---|
| **Amino Acid Metabolism,Endocrine System Development and Function,Molecular Transport,Small Molecule Biochemistry** | Quantity of L-triiodothyronine | 0.0103 | NCOR1, SLCO1C1 | 2 |
| **Nervous System Development and Function,Tissue Morphology** | Morphology of nervous tissue | 0.0118 | AGRN, BRCA2, CADPS2, COL13A1, CRMP1, DBN1, DVL1, EPHB3, ERC2, FAT3, GAP43, HIVEP2, IDUA, MAN2C1, MYO16, NOS2, NRN1, PLAGL1, PRKN, RGS11, RIMS2, ROCK2, SCN1B, SEMA5B, SHANK2, SORBS2, SORL1, SPTBN4, SYNE1, TIAM1, UNC13A, UNC13B | 32 |
| **Lipid Metabolism,Small Molecule Biochemistry** | Synthesis of eicosanoid | 0.012 | KNG1, NOS2, PTGS2 | 3 |
| **Cell Morphology,Cellular Assembly and Organization,Cellular Development,Cellular Function and Maintenance,Cellular Growth and Proliferation,Nervous System Development and Function,Organismal Development,Tissue Development** | Branching of neurons | 0.0123 | ABI1, AGRN, BSN, CRMP1, DAB2IP, DBN1, ELK1, EPHB3, ERC2, GABRB3, GAP43, MYO5B, NRN1, PTGS2, ROCK2, SHANK2, SORBS2, TIAM1, TSC2 | 19 |
| **Cellular Assembly and Organization** | Clustering of synaptic vesicles | 0.0141 | BSN, PCLO | 2 |
| **Behavior,Nervous System Development and Function** | Memory consolidation | 0.0141 | CREBBP, NR3C2 | 2 |
| **Cell-To-Cell Signaling and Interaction,Drug Metabolism,Small Molecule Biochemistry** | Metabolism of dopamine | 0.0141 | PRKN, SORCS2 | 2 |
| **Behavior,Nervous System Development and Function** | Memory | 0.0142 | ADCY1, CREBBP, KCNH3, NEURL1, NR3C2 | 5 |
| **Cell Morphology,Nervous System Development and Function,Tissue Morphology** | Morphology of neurons | 0.0146 | AGRN, BRCA2, CADPS2, COL13A1, CRMP1, DBN1, DVL1, EPHB3, ERC2, FAT3, GAP43, HIVEP2, IDUA, MYO16, NOS2, NRN1, PLAGL1, | 31 |

| | | | | |
|---|---|---|---|---|
| | | | PRKN, RGS11, RIMS2, ROCK2, SCN1B, SEMA5B, SHANK2, SORBS2, SORL1, SPTBN4, SYNE1, TIAM1, UNC13A, UNC13B | |
| **Cellular Movement,Nervous System Development and Function** | Guidance of axons | 0.0151 | DVL1, EPHB3, GAP43, KIF5C, LMTK2, PIK3R1, PTPRO, SCN1B, SEMA4F, SPTBN4, UNC5D, VAX1 | 12 |
| **Cell Morphology,Nervous System Development and Function,Neurological Disease,Tissue Morphology** | Abnormal morphology of bipolar cells | 0.0161 | ERC2, FAT3, SEMA5B | 3 |
| **Neurological Disease,Psychological Disorders** | Pervasive developmental disorder | 0.0184 | MATK, SLC6A7, TSC2 | 3 |
| **Cell-To-Cell Signaling and Interaction,Nervous System Development and Function** | Action potential of motor endplates | 0.0185 | COL13A1, RIMS2 | 2 |
| **Cell Morphology,Cellular Assembly and Organization,Cellular Development,Cellular Function and Maintenance,Cellular Growth and Proliferation,Nervous System Development and Function,Organismal Development,Tissue Development** | Axonogenesis | 0.0188 | ADCY1, AGRN, ARHGEF28, BCL11B, DVL1, EPHB3, GAP43, NIN, NRN1, PIP5K1C, SPTBN4, TIAM1, TOP2B, TSC2 | 14 |
| **Cell-To-Cell Signaling and Interaction,Drug Metabolism,Molecular Transport,Small Molecule Biochemistry** | Concentration of norepinephrine | 0.0209 | DBH, GAP43, PRKN | 3 |
| **Cell-To-Cell Signaling and Interaction,Nervous System Development and Function** | Synaptic depression of collateral synapses | 0.0234 | ADCY1, RAB11FIP5 | 2 |
| **Amino Acid Metabolism,Drug Metabolism,Endocrine System Development and Function,Molecular Transport,Small Molecule Biochemistry** | Concentration of levothyroxine | 0.0234 | NCOR1, SLCO1C1 | 2 |
| **Nervous System Development and Function,Neurological Disease,Organ Morphology,Organismal Development,Organismal Injury and Abnormalities** | Abnormal morphology of anterior horn of the spinal cord | 0.0234 | KIF1B, RIMS2 | 2 |
| **Nervous System Development and Function,Neurological Disease,Organ Morphology,Organismal Development,Organismal Injury and Abnormalities** | Abnormal morphology of optic tract | 0.0234 | EPHB3, GAP43 | 2 |
| **Nervous System Development and Function,Neurological Disease,Ophthalmic Disease** | Abnormal morphology of optic nerve | 0.0236 | SCN1B, SPTBN4, VAX1 | 3 |
| **Cellular Assembly and Organization,Nervous System Development and Function** | Recycling of synaptic vesicles | 0.0264 | DVL1, GAP43, PIP5K1C | 3 |
| **Carbohydrate Metabolism,Drug Metabolism,Molecular Transport,Small Molecule Biochemistry** | Quantity of heparan sulfate | 0.0272 | IDUA | 1 |

| | | | | |
|---|---|---|---|---|
| **Cell-To-Cell Signaling and Interaction,Cellular Assembly and Organization,Nervous System Development and Function** | Modulation of synapse | 0.0272 | PCDH8 | 1 |
| **Cellular Development,Nervous System Development and Function,Tissue Development** | Differentiation of sympathetic preganglionic neurons | 0.0272 | AGRN | 1 |
| **Cellular Assembly and Organization** | Aggregation of early endosomes | 0.0272 | KIF16B | 1 |
| **Cell Morphology,Cellular Assembly and Organization,Cellular Function and Maintenance** | Formation of axon-like process | 0.0272 | TIAM1 | 1 |
| **Cell Morphology,Cell-To-Cell Signaling and Interaction** | Length of stereocilia | 0.0272 | MYO15A | 1 |
| **Cell-To-Cell Signaling and Interaction,Cellular Assembly and Organization,Nervous System Development and Function,Tissue Morphology** | Quantity of ribbon synapse | 0.0272 | BSN | 1 |
| **Nervous System Development and Function,Tissue Morphology** | Density of serotonergic neurons | 0.0272 | GAP43 | 1 |
| **Behavior,Developmental Disorder,Neurological Disease,Psychological Disorders** | Autistic-like traits | 0.0272 | TSC2 | 1 |
| **Hereditary Disorder,Neurological Disease,Organismal Injury and Abnormalities,Psychological Disorders,Skeletal and Muscular Disorders** | Autosomal recessive juvenile Parkinson disease | 0.0272 | PRKN | 1 |
| **Cellular Compromise** | Acidification of cytosol | 0.0272 | CA4 | 1 |
| **Cell-To-Cell Signaling and Interaction,Nervous System Development and Function** | Inactivation of synapse | 0.0272 | BSN | 1 |
| **Cellular Development,Nervous System Development and Function,Organ Development** | Differentiation of preoptic area | 0.0272 | VAX1 | 1 |
| **Cellular Development,Nervous System Development and Function,Organ Development** | Differentiation of medial ganglionic eminences | 0.0272 | VAX1 | 1 |
| **Cellular Development,Cellular Growth and Proliferation,Embryonic Development,Nervous System Development and Function,Organ Development,Organismal Development,Tissue Development,Visual System Development and Function** | Formation of retinal cells | 0.0272 | ADCY1 | 1 |
| **Nervous System Development and Function,Organ Morphology,Tissue Morphology** | Quantity of rod bipolar cells | 0.0272 | PRDM1 | 1 |
| **Cell Morphology,Cellular Assembly and Organization,Cellular Development,Cellular Function and Maintenance,Cellular Growth and Proliferation,Embryonic Development,Nervous System Development and Function,Organismal Development,Tissue Development** | Delay in initiation of outgrowth of dendrites | 0.0272 | DAB2IP | 1 |
| **Cell-To-Cell Signaling and Interaction,Nervous System Development and Function** | Clustering of hippocampal neurons | 0.0272 | SHANK2 | 1 |
| **Molecular Transport,Protein Trafficking** | Translocation of protein fragment | 0.0272 | AGAP3 | 1 |
| **Nervous System Development and Function,Tissue Morphology** | Density of parallel fiber | 0.0272 | DAB2IP | 1 |
| **Cell Death and Survival,Cellular Development,Nervous System Development and Function,Tissue Development,Tissue Morphology** | Regeneration of bipolar cells | 0.0272 | PRDM1 | 1 |
| **Nervous System Development and Function,Organ Morphology,Organismal Development** | Size of optic chiasm | 0.0272 | GAP43 | 1 |
| **Cell Morphology,Cellular Assembly and Organization,Cellular Development,Cellular Function and Maintenance,Cellular Growth and Proliferation,Nervous System Development and Function,Organismal Development,Tissue Development** | Sprouting of rod bipolar cells | 0.0272 | ERC2 | 1 |
| **Cellular Development,Nervous System Development and Function,Tissue Development** | Differentiation of pyramidal neurons | 0.0272 | UNC5D | 1 |
| **Inflammatory Response** | Inflammatory response of cerebral cortex | 0.0272 | KNG1 | 1 |
| **Cellular Development** | Delay in initiation of transition of radial glial cells | 0.0272 | PLAGL1 | 1 |

| | | | | |
|---|---|---|---|---|
| **Cellular Development** | Delay in initiation of transition of neuronal progenitor cells | 0.0272 | PLAGL1 | 1 |
| **Embryonic Development,Nervous System Development and Function,Organ Development,Organismal Development,Tissue Development** | Development of septal area | 0.0272 | VAX1 | 1 |
| **Behavior** | Social exploration | 0.0272 | APBA2 | 1 |
| **Hereditary Disorder,Neurological Disease,Organismal Injury and Abnormalities,Psychological Disorders,Skeletal and Muscular Disorders** | Parkinson-dementia syndrome | 0.0272 | GAP43 | 1 |
| **Cardiovascular System Development and Function,Organismal Development** | Penetration by blood vessel | 0.0272 | ADGRA2 | 1 |
| **Developmental Disorder,Endocrine System Disorders,Nutritional Disease,Organismal Injury and Abnormalities,Reproductive System Disease** | Delayed puberty | 0.0272 | VAX1 | 1 |
| **Cell-To-Cell Signaling and Interaction,Nervous System Development and Function** | Sensitization of trigeminovascular neuron | 0.0272 | KNG1 | 1 |
| **Behavior** | Perseverance behavior | 0.0272 | NR3C2 | 1 |
| **Cellular Development,Cellular Growth and Proliferation,Nervous System Development and Function,Tissue Development** | Development of corticospinal neurons | 0.0272 | BCL11B | 1 |
| **Behavior** | Impulsivity | 0.0272 | ADCY1 | 1 |
| **Metabolic Disease,Neurological Disease,Organismal Injury and Abnormalities,Psychological Disorders** | Preclinical stage Alzheimer disease | 0.0272 | NOS2 | 1 |
| **Hematological Disease** | Hyperemia | 0.0272 | PTGS2 | 1 |
| **Nervous System Development and Function,Tissue Morphology** | Density of climbing fiber | 0.0272 | DAB2IP | 1 |
| **Embryonic Development,Nervous System Development and Function,Organ Development,Organismal Development,Tissue Development** | Development of layer IV | 0.0272 | GAP43 | 1 |
| **Cell-To-Cell Signaling and Interaction,Drug Metabolism,Small Molecule Biochemistry** | Conversion of norepinephrine | 0.0272 | DBH | 1 |
| **Infectious Diseases** | Replication of Murine cytomegalovirus | 0.0272 | NOS2 | 1 |
| **Behavior** | Risk preference | 0.0272 | GABRB3 | 1 |
| **Metabolic Disease,Neurological Disease,Organismal Injury and Abnormalities,Psychological Disorders** | Stage V Alzheimer disease | 0.0272 | PTGS2 | 1 |
| **Cell Morphology,Cellular Assembly and Organization,Nervous System Development and Function** | Projection of mossy fibers | 0.0272 | NR3C2 | 1 |
| **Cell-To-Cell Signaling and Interaction,Nervous System Development and Function** | Activation of mitral cells | 0.0272 | GJD2 | 1 |
| **Cell Morphology,Cellular Assembly and Organization** | Orientation of plasma membrane projections | 0.0272 | SCN1B | 1 |
| **Cell-To-Cell Signaling and Interaction,Nervous System Development and Function** | Excitation of postsynaptic region | 0.0272 | PCDH8 | 1 |
| **Cellular Assembly and Organization,Cellular Development,Nervous System Development and Function,Tissue Development** | Differentiation of presynaptic terminals | 0.0272 | AGRN | 1 |
| **Cell-To-Cell Signaling and Interaction,Drug Metabolism,Small Molecule Biochemistry** | Catabolism of dopamine | 0.0272 | PRKN | 1 |
| **Nervous System Development and Function,Neurological Disease,Organ Morphology,Organismal Development,Organismal Injury and Abnormalities** | Lack of optic tract | 0.0272 | GAP43 | 1 |
| **Cell Morphology,Cellular Assembly and Organization,Nervous System Development and Function,Tissue Morphology** | Diameter of nodes of Ranvier | 0.0272 | SPTBN4 | 1 |
| **Cellular Development,Nervous System Development and Function,Tissue Development** | Differentiation of postsynaptic neurons | 0.0272 | AGRN | 1 |
| **Nervous System Development and Function** | Discharge of cochlear nerve | 0.0272 | BSN | 1 |
| **Cardiovascular System Development and Function,Nervous System Development and Function,Organ Morphology,Organismal Development,Tissue Morphology** | Diameter of basilar artery | 0.0272 | KNG1 | 1 |

| | | | | |
|---|---|---|---|---|
| **Cell Morphology,Cellular Movement,Nervous System Development and Function** | Innervation of serotonergic neurons | 0.0272 | GAP43 | 1 |
| **Cell-To-Cell Signaling and Interaction,Nervous System Development and Function** | Firing of trigeminovascular neuron | 0.0272 | KNG1 | 1 |
| **Cell Morphology,Cellular Assembly and Organization** | Elongation of astrocyte projections | 0.0272 | GAP43 | 1 |
| **Lipid Metabolism,Molecular Transport,Small Molecule Biochemistry** | Concentration of eicosapentenoic acid | 0.0272 | ARNTL | 1 |
| **Cell-To-Cell Signaling and Interaction,Cellular Function and Maintenance,Nervous System Development and Function** | Paired-pulse facilitation of parallel fiber-Purkinje cell synapses | 0.0272 | UNC13C | 1 |
| **Metabolic Disease,Neurological Disease,Organismal Injury and Abnormalities,Psychological Disorders** | Stage I Alzheimer disease | 0.0272 | PTGS2 | 1 |
| **Nervous System Development and Function,Tissue Morphology** | Density of axons | 0.0287 | ACVR1C, DAB2IP | 2 |
| **Embryonic Development,Nervous System Development and Function,Organ Development,Organismal Development,Tissue Development,Visual System Development and Function** | Formation of eye | 0.0287 | ADCY1, FAT3 | 2 |
| **Cell-To-Cell Signaling and Interaction,Cellular Assembly and Organization,Cellular Development,Cellular Function and Maintenance,Cellular Growth and Proliferation,Nervous System Development and Function,Tissue Development** | Formation of excitatory synapses | 0.0287 | LRRTM4, SORBS2 | 2 |
| **Neurological Disease,Organismal Injury and Abnormalities,Psychological Disorders,Skeletal and Muscular Disorders** | Parkinson disease with dementia | 0.0345 | GAP43, NOS2 | 2 |
| **Cell-To-Cell Signaling and Interaction,Cellular Assembly and Organization,Cellular Function and Maintenance** | Development of gap junctions | 0.0375 | AGRN, BSN, GJD2, LRRTM4, NRN1, PCDH8, PLXND1, SHANK2, SORBS2 | 9 |
| **Tissue Morphology** | Quantity of cells | 0.0384 | ACVR1C, AGRN, BSN, CADPS2, DVL1, EPHB3, ERC2, HIVEP2, KIF1B, KIF5C, MGAT5B, NEURL1, NOS2, NR3C2, NRN1, PCLO, PRDM1, PRDM16, PRKN, PTGS2, RIMS2, TRPM2, TSC2, UNC13A, UNC13B, VAX1 | 26 |
| **Cell Morphology,Cellular Assembly and Organization,Cellular Function and Maintenance,Nervous System Development and Function** | Extension of neurites | 0.04 | ADCY6, ANAPC2, DBN1, DVL1, MYO5B, SCN1B, SEMA4F, SEMA5B, TIAM1 | 9 |
| **Cell Morphology,Cellular Compromise** | Collapse of growth cone | 0.0415 | ARHGEF11, EPHB3, ROCK2, | 5 |

| | | | | |
|---|---|---|---|---|
| | | | SEMA4F, SORCS2 | |
| **Nervous System Development and Function,Neurological Disease,Organ Morphology,Organismal Development,Organismal Injury and Abnormalities** | Abnormal morphology of forebrain | 0.0439 | ADCY1, ADGRA2, ARNT2, ASPM, CELSR2, CREBBP, HIVEP2, MAN2C1, NCOR1, NOS2, TOP2B, VAX1 | 12 |
| **Cell Morphology,Cellular Assembly and Organization,Cellular Development,Cellular Function and Maintenance,Cellular Growth and Proliferation,Embryonic Development,Nervous System Development and Function,Organismal Development,Tissue Development,Tissue Morphology** | Density of dendritic spines | 0.0447 | CRMP1, ELK1, GABRB3, PTGS2, TIAM1, TSC2 | 6 |
| **Cell-To-Cell Signaling and Interaction,Cellular Function and Maintenance,Nervous System Development and Function** | Paired-pulse facilitation of synapse | 0.0464 | APBA2, CADPS2, ROCK2, UNC13C | 4 |
| **Embryonic Development,Nervous System Development and Function,Organ Development,Organismal Development,Tissue Development** | Formation of corpus callosum | 0.047 | EPHB3, GAP43, NIN | 3 |
| **Carbohydrate Metabolism** | Accumulation of carbohydrate | 0.0472 | IDUA, KNG1 | 2 |
| **Drug Metabolism,Lipid Metabolism,Small Molecule Biochemistry** | Synthesis of prostaglandin E2 | 0.0472 | NOS2, PTGS2 | 2 |
| **Carbohydrate Metabolism,Lipid Metabolism,Molecular Transport,Small Molecule Biochemistry** | Concentration of phosphatidic acid | 0.0472 | PIP5K1C, PTGS2 | 2 |
| **Cell Morphology,Cellular Assembly and Organization,Cellular Development,Cellular Function and Maintenance,Cellular Growth and Proliferation,Nervous System Development and Function,Organismal Development,Tissue Development** | Formation of dendrites | 0.0481 | ABI1, ADGRB3, ANAPC2, CELSR2, DVL1, FMN1, MYO5B, ROCK2, SYNE1 | 9 |

### Cluster 2

| Categories | Diseases or Functions Annotation | p-Value | Molecules | # Molecules |
|---|---|---|---|---|
| **Cellular Growth and Proliferation** | Expansion of cells | 0.000331 | FTO, IFT88, IRX6, ITGB1, RET | 5 |
| **Neurological Disease** | Chronic fatigue syndrome | 0.000444 | C3, CP, GSN, SERPINA3 | 4 |
| **Cell Morphology,Cell-To-Cell Signaling and Interaction,Cellular Assembly and Organization** | Structural integrity of synapse | 0.000488 | DTNA, SNTA1 | 2 |
| **Cell Morphology,Nervous System Development and Function,Neurological Disease,Tissue Morphology** | Abnormal morphology of neurons | 0.000755 | B2M, C3, CD81, CLCN3, CLCN6, CNR2, COL25A1, CP, CPEB2, CYP19A1, DTNA, ESR1, GLI1, GLI2, ITGB1, NTRK1, PTPRC, RET, ROBO1, SERPINA3, SHC2, SIL1, SLITRK6, SNTA1, | 26 |

| | | | | |
|---|---|---|---|---|
| | | | UTRN, ZNF423 | |
| **Cell Morphology,Cell-To-Cell Signaling and Interaction,Cellular Assembly and Organization,Nervous System Development and Function,Neurological Disease,Tissue Morphology** | Abnormal morphology of synapse | 0.000776 | B2M, C3, CYP19A1, DTNA, ITGB1, RET, SNTA1, UTRN | 8 |
| **Nervous System Development and Function,Neurological Disease** | Abnormal morphology of nervous system | 0.00104 | AXL, B2M, C3, CD81, CLCN3, CLCN6, CNR2, COL18A1, COL25A1, CP, CPEB2, CYP19A1, DTNA, ESR1, GLI1, GLI2, ITGB1, LEF1, NTRK1, PEG3, PHGDH, PRKACA, PTPRC, PTPRZ1, REST, RET, ROBO1, SERPINA3, SHC2, SIL1, SLC44A2, SLC6A3, SLITRK6, SNTA1, TCF7L2, UTRN, YES1, ZNF423 | 38 |
| **Cell Morphology,Cellular Assembly and Organization,Cellular Development,Cellular Function and Maintenance,Cellular Growth and Proliferation,Embryonic Development,Nervous System Development and Function,Organismal Development,Tissue Development** | Branching of motor axons | 0.00144 | COL25A1, NTRK1 | 2 |
| **Cell Morphology** | Surface area of cells | 0.00144 | DISC1, NOTCH4 | 2 |
| **Reproductive System Development and Function** | Onset of puberty | 0.00144 | ESR1, ITGB1 | 2 |
| **Tissue Morphology** | Quantity of cells | 0.00162 | B2M, C3, CDNF, CLCN3, CNR2, COL25A1, CP, DISC1, ESR1, FTO, GLI1, GLI2, ITGB1, LEF1, NTRK1, PGR, PTPRC, PTPRZ1, RET, SHC2, SLC44A2, SLC6A3, TCF7L2, TNC, UTRN, XIAP, ZNF423 | 27 |
| **Tissue Morphology** | Quantity of stem cells | 0.00239 | GLI1, GLI2, ITGB1, TNC | 4 |
| **Nervous System Development and Function,Organ Morphology,Organismal Development** | Morphology of forebrain | 0.00254 | AKAP13, AXL, B2M, CD81, CLCN3, CNR2, COL18A1, ESR1, GLI1, GLI2, PEG3, | 15 |

| | | | | |
|---|---|---|---|---|
| | | | PHGDH, REST, ROBO1, ZNF423 | |
| **Endocrine System Development and Function,Nervous System Development and Function,Organ Morphology,Tissue Morphology** | Quantity of lactotropes | 0.0026 | ESR1, PGR, SLC6A3 | 3 |
| **Embryonic Development,Organismal Development,Tissue Development** | Morphogenesis of ophthalmic vesicle | 0.00284 | FTO, IRX6 | 2 |
| **Nervous System Development and Function,Organ Morphology,Organismal Development** | Size of pallium | 0.00284 | FTO, IRX6 | 2 |
| **Cellular Development,Cellular Growth and Proliferation,Nervous System Development and Function,Tissue Development** | Expansion of motor neurons | 0.00284 | FTO, IRX6 | 2 |
| **Cellular Development,Cellular Growth and Proliferation,Nervous System Development and Function,Tissue Development** | Generation of v3 neuron | 0.00284 | FTO, IRX6 | 2 |
| **Inflammatory Disease,Inflammatory Response,Neurological Disease,Organismal Injury and Abnormalities** | HIV-associated encephalitis | 0.00284 | CNR2, PTPRC | 2 |
| **Cell Death and Survival,Cellular Compromise,Neurological Disease,Tissue Morphology** | Degeneration of motor axons | 0.00284 | COL25A1, NFIL3 | 2 |
| **Endocrine System Development and Function,Endocrine System Disorders,Organ Morphology,Organismal Development,Organismal Injury and Abnormalities,Reproductive System Disease** | Lack of pituitary gland | 0.00284 | GLI1, GLI2 | 2 |
| **Nervous System Development and Function** | Morphology of nervous system | 0.00304 | AKAP13, AXL, B2M, C3, CD81, CLCN3, CLCN6, CNR2, COL18A1, COL25A1, CP, CPEB2, CYP19A1, DISC1, DTNA, ESR1, FTO, GLI1, GLI2, IRX6, ITGB1, LEF1, NTRK1, PEG3, PHGDH, PRKACA, PTPRC, PTPRZ1, REST, RET, ROBO1, SERPINA3, SHC2, SIL1, SLC44A2, SLC6A3, SLITRK6, SNTA1, TCF7L2, UTRN, YES1, ZNF423 | 42 |
| **Cellular Development,Cellular Growth and Proliferation,Nervous System Development and Function,Tissue Development** | Expansion of neurons | 0.00325 | FTO, IRX6, RET | 3 |
| **Nervous System Development and Function,Neurological Disease,Organ Morphology,Organismal Development,Organismal Injury and Abnormalities** | Abnormal morphology of forebrain | 0.00418 | AXL, B2M, CD81, CLCN3, COL18A1, ESR1, GLI1, GLI2, PEG3, PHGDH, REST, ROBO1, ZNF423 | 13 |

| | | | | |
|---|---|---|---|---|
| **Embryonic Development,Nervous System Development and Function,Organ Development,Organismal Development,Tissue Development** | Formation of cerebellar cortex | 0.0046 | CYP19A1, GLI1, GLI2, ZNF423 | 4 |
| **Nervous System Development and Function** | Antinociception of spinal cord | 0.00466 | ESR1, PGR | 2 |
| **Embryonic Development,Nervous System Development and Function,Organ Development,Organ Morphology,Organismal Development,Tissue Development** | Morphogenesis of cerebellar cortex | 0.00466 | GLI1, GLI2 | 2 |
| **Endocrine System Development and Function,Organ Development** | Function of pituitary gland | 0.00466 | ESR1, SLC6A3 | 2 |
| **Cell Morphology,Nervous System Development and Function** | Projection of neurons | 0.00466 | GLI2, ROBO1 | 2 |
| **Cellular Development,Nervous System Development and Function,Tissue Development** | Differentiation of sensory neurons | 0.00517 | ERCC2, MYO7A, NTRK1, RET | 4 |
| **Cell Morphology,Cell-To-Cell Signaling and Interaction,Cellular Assembly and Organization,Nervous System Development and Function,Neurological Disease,Tissue Morphology** | Loss of synapse | 0.00578 | C3, CYP19A1, ITGB1 | 3 |
| **Cellular Development,Cellular Growth and Proliferation,Embryonic Development,Nervous System Development and Function,Organ Development,Organismal Development,Tissue Development** | Proliferation of neural precursor cells | 0.00584 | CNR2, GLI1, GLI2, HUWE1, IFT88, ORC3 | 6 |
| **Nervous System Development and Function** | Analgesia | 0.00683 | ESR1, PGR, REST | 3 |
| **Neurological Disease** | Disorder of autonomic nervous system | 0.00715 | CPEB2, CYP19A1, NTRK1, RET | 4 |
| **Cell Morphology,Cell-To-Cell Signaling and Interaction,Cellular Assembly and Organization,Nervous System Development and Function,Neurological Disease,Tissue Morphology** | Abnormal morphology of neuromuscular synapse | 0.00715 | DTNA, RET, SNTA1, UTRN | 4 |
| **Embryonic Development,Endocrine System Development and Function,Organ Development,Organismal Development,Tissue Development** | Development of pituitary gland | 0.00791 | GLI1, GLI2, SLC6A3, TCF7L2 | 4 |
| **Cancer,Neurological Disease,Organismal Injury and Abnormalities** | Development of neuroectodermal tumor | 0.00791 | FAT1, GLI1, GLI2, RET | 4 |
| **Nervous System Development and Function,Neurological Disease,Organ Morphology,Organismal Development,Organismal Injury and Abnormalities** | Lack of cerebellum | 0.00951 | PHGDH, ZNF423 | 2 |
| **Endocrine System Development and Function,Endocrine System Disorders,Organ Morphology,Organismal Development,Organismal Injury and Abnormalities,Reproductive System Disease** | Abnormal morphology of pituitary gland | 0.00957 | GLI1, GLI2, SLC6A3, TCF7L2 | 4 |
| **Cell Morphology,Nervous System Development and Function,Tissue Morphology** | Morphology of neurons | 0.00958 | B2M, C3, CD81, CLCN3, CLCN6, CNR2, COL25A1, CP, CPEB2, CYP19A1, DISC1, DTNA, ESR1, GLI1, GLI2, ITGB1, NTRK1, PTPRC, RET, ROBO1, SERPINA3, SHC2, SIL1, SLITRK6, SNTA1, | 27 |

| | | | | |
|---|---|---|---|---|
| | | | UTRN, ZNF423 | |
| **Nervous System Development and Function,Tissue Morphology** | Quantity of neurons | 0.0101 | B2M, C3, CDNF, CLCN3, COL25A1, CP, DISC1, ESR1, GLI2, LEF1, NTRK1, PTPRZ1, RET, SHC2, SLC44A2, SLC6A3, UTRN, XIAP, ZNF423 | 19 |
| **Developmental Disorder,Neurological Disease,Organismal Injury and Abnormalities** | Hypoplasia of cerebellum | 0.0106 | IFT88, PHGDH, ZNF423 | 3 |
| **Nervous System Development and Function** | Morphology of central nervous system | 0.0107 | AKAP13, AXL, B2M, C3, CD81, CLCN3, CNR2, COL18A1, CP, ESR1, FTO, GLI1, GLI2, IRX6, LEF1, NTRK1, PEG3, PHGDH, PTPRC, PTPRZ1, REST, ROBO1, SIL1, SLC6A3, TCF7L2, YES1, ZNF423 | 27 |
| **Embryonic Development,Nervous System Development and Function,Organ Development,Organ Morphology,Organismal Development,Tissue Development** | Morphogenesis of brain | 0.0118 | FTO, GLI1, GLI2, IFT88, IRX6 | 5 |
| **Cell Morphology** | Area of cells | 0.0121 | DISC1, NOTCH4, NTRK1 | 3 |
| **Nervous System Development and Function,Neurological Disease,Organ Morphology,Organismal Development,Organismal Injury and Abnormalities** | Abnormal morphology of fourth cerebral ventricle | 0.0125 | COL18A1, CP | 2 |
| **Embryonic Development,Nervous System Development and Function,Organ Development,Organismal Development,Tissue Development** | Formation of external granular layer | 0.0125 | GLI1, GLI2 | 2 |
| **Cell-To-Cell Signaling and Interaction,Nervous System Development and Function** | Long-term potentiation of striatum | 0.0125 | PRKACA, SLC6A3 | 2 |
| **Cell-To-Cell Signaling and Interaction,Cellular Growth and Proliferation,Nervous System Development and Function** | Excitation of sensory neurons | 0.0125 | REST, SCN9A | 2 |
| **Cellular Development,Cellular Growth and Proliferation,Embryonic Development,Nervous System Development and Function,Organ Development,Organismal Development,Tissue Development** | Development of Purkinje cells | 0.0125 | CYP19A1, ZNF423 | 2 |
| **Nervous System Development and Function,Neurological Disease,Organ Morphology,Organismal Development,Organismal Injury and Abnormalities** | Abnormal morphology of lateral cerebral ventricle | 0.0126 | B2M, CLCN3, COL18A1, PHGDH, ZNF423 | 5 |
| **Organismal Development** | Morphology of head | 0.0128 | AKAP13, AXL, B2M, C3, CD81, | 27 |

| | | | | |
|---|---|---|---|---|
| | | | CLCN3, CNR2, COL18A1, CP, ESR1, FTO, GLI1, GLI2, IRX6, LEF1, PEG3, PHGDH, PTPRC, REST, ROBO1, SIL1, SLC44A2, SLC6A3, SLITRK6, TCF7L2, YES1, ZNF423 | |
| **Nervous System Development and Function,Organ Morphology,Organismal Development** | Morphology of brain | 0.0136 | AKAP13, AXL, B2M, C3, CD81, CLCN3, CNR2, COL18A1, CP, ESR1, FTO, GLI1, GLI2, IRX6, LEF1, PEG3, PHGDH, PTPRC, REST, ROBO1, SIL1, SLC6A3, TCF7L2, YES1, ZNF423 | 25 |
| **Cell Death and Survival,Cell Morphology,Nervous System Development and Function,Neurological Disease,Tissue Morphology** | Loss of sympathetic neuron | 0.0158 | NTRK1, SHC2 | 2 |
| **Endocrine System Development and Function,Small Molecule Biochemistry** | Synthesis of hormone | 0.0158 | ESR1, GPR39 | 2 |
| **Cell-To-Cell Signaling and Interaction,Cellular Assembly and Organization** | Cell-cell contact | 0.0187 | B2M, C3, CYP19A1, DISC1, DTNA, ITGB1, RET, SEZ6L, SLITRK6, SNTA1, TNC, UTRN, XIAP | 13 |
| **Cellular Development,Nervous System Development and Function,Tissue Development** | Differentiation of mechanosensory neurons | 0.0191 | ERCC2, MYO7A, NTRK1 | 3 |
| **Cell Morphology** | Projection | 0.0191 | GLI2, NTRK1, ROBO1 | 3 |
| **Cell Morphology** | Length of plasma membrane projections | 0.0195 | CNR2, PTPRZ1 | 2 |
| **Cellular Growth and Proliferation,Nervous System Development and Function,Organ Development** | Proliferation of brain cells | 0.0202 | CNR2, GLI1, GLI2, HUWE1, IFT88, ORC3, ZNF423 | 7 |
| **Embryonic Development,Nervous System Development and Function,Organ Development,Organ Morphology,Organismal Development,Tissue Development** | Morphogenesis of forebrain | 0.0212 | FTO, IFT88, IRX6 | 3 |
| **Nervous System Development and Function,Neurological Disease** | Abnormal morphology of central nervous system | 0.0219 | AXL, B2M, C3, CD81, CLCN3, COL18A1, CP, ESR1, GLI1, GLI2, LEF1, PEG3, | 22 |

| | | | | |
|---|---|---|---|---|
| | | | PHGDH, PTPRC, PTPRZ1, REST, ROBO1, SIL1, SLC6A3, TCF7L2, YES1, ZNF423 | |
| **Nervous System Development and Function,Neurological Disease,Organ Morphology,Organismal Development,Organismal Injury and Abnormalities** | Abnormal morphology of nucleus accumbens | 0.0221 | CD81 | 1 |
| **Cardiovascular System Development and Function,Tissue Morphology** | Enlargement of microvessel | 0.0221 | NOTCH4 | 1 |
| **Cell-To-Cell Signaling and Interaction,Cellular Assembly and Organization,Cellular Development,Cellular Function and Maintenance,Cellular Growth and Proliferation,Embryonic Development,Nervous System Development and Function,Organ Development,Organismal Development,Tissue Development** | Synaptogenesis of Purkinje cells | 0.0221 | CYP19A1 | 1 |
| **Developmental Disorder,Endocrine System Disorders,Organismal Injury and Abnormalities,Reproductive System Disease** | Hypoplasia of intermediate lobe of the pituitary gland | 0.0221 | SLC6A3 | 1 |
| **Cellular Compromise,Nervous System Development and Function** | Acidification of synaptic vesicles | 0.0221 | CLCN3 | 1 |
| **Endocrine System Development and Function,Organ Morphology,Organismal Development,Reproductive System Development and Function,Tissue Morphology** | Size of pituitary tissue | 0.0221 | SLC6A3 | 1 |
| **Cell-To-Cell Signaling and Interaction,Cellular Growth and Proliferation,Nervous System Development and Function** | Hypoexcitation of sympathetic neuron | 0.0221 | SCN9A | 1 |
| **Nervous System Development and Function** | Survival of lumbar 3 dorsal root ganglion | 0.0221 | NTRK1 | 1 |
| **Cardiovascular System Development and Function,Cell Morphology,Organismal Development,Tissue Morphology** | Surface area of endothelial cells | 0.0221 | NOTCH4 | 1 |
| **Cell Death and Survival,Cellular Development,Nervous System Development and Function,Organ Morphology,Tissue Development,Tissue Morphology** | Regeneration of olfactory receptor neurons | 0.0221 | IFT88 | 1 |
| **Cellular Assembly and Organization,Cellular Development,Nervous System Development and Function,Tissue Development** | Maturation of postsynaptic density | 0.0221 | DTNA | 1 |
| **Cell-To-Cell Signaling and Interaction,Cellular Function and Maintenance,Inflammatory Response** | Phagocytosis of photoreceptors | 0.0221 | MYO7A | 1 |
| **Cell Morphology,Cellular Development,Embryonic Development,Tissue Development** | Stratification of Bergmann glia | 0.0221 | HUWE1 | 1 |
| **Dermatological Diseases and Conditions,Neurological Disease,Organismal Injury and Abnormalities** | Hypesthesia | 0.0221 | REST | 1 |
| **Cell Morphology** | Projection of cholinergic fibers | 0.0221 | NTRK1 | 1 |
| **Cell Cycle** | Senescence of oligodendrocyte precursor cells | 0.0221 | C2orf40 | 1 |
| **Nervous System Development and Function,Organ Morphology** | Density of hippocampal CA1 region | 0.0221 | NTRK1 | 1 |
| **Nervous System Development and Function** | Guidance of sympathetic neuron | 0.0221 | RET | 1 |
| **Developmental Disorder,Neurological Disease,Organismal Injury and Abnormalities** | Hypoplasia of ganglionic eminences | 0.0221 | PHGDH | 1 |
| **Cell-To-Cell Signaling and Interaction,Cellular Growth and Proliferation,Nervous System Development and Function** | Hyperexcitation of sensory neurons | 0.0221 | SCN9A | 1 |

| | | | | |
|---|---|---|---|---|
| **Cellular Development,Cellular Growth and Proliferation,Embryonic Development,Organismal Development,Tissue Development** | Expansion of granule neuron progenitor cells | 0.0221 | IFT88 | 1 |
| **Cellular Movement,Nervous System Development and Function** | Migration of CA1 neuron | 0.0221 | DISC1 | 1 |
| **Cellular Assembly and Organization,Nervous System Development and Function** | Folding of postsynaptic membrane | 0.0221 | UTRN | 1 |
| **Cellular Development,Nervous System Development and Function,Tissue Development** | Spinogenesis of Purkinje cells | 0.0221 | CYP19A1 | 1 |
| **Developmental Disorder,Hereditary Disorder,Metabolic Disease,Neurological Disease,Ophthalmic Disease,Organismal Injury and Abnormalities** | Finnish type amyloidosis | 0.0221 | GSN | 1 |
| **Cellular Compromise** | Disorganization of Bergmann glia | 0.0221 | HUWE1 | 1 |
| **Neurological Disease** | Dysfunction of enteric nervous system | 0.0221 | RET | 1 |
| **Embryonic Development,Organismal Development,Tissue Development** | Formation of floor plate | 0.0221 | GLI2 | 1 |
| **Embryonic Development,Organismal Development,Tissue Morphology** | Morphology of visceral endoderm | 0.0221 | AMOT | 1 |
| **Endocrine System Development and Function,Lipid Metabolism,Molecular Transport,Small Molecule Biochemistry** | Uptake of estrogen | 0.0221 | ESR1 | 1 |
| **Embryonic Development,Tissue Morphology** | Quantity of granule neuron progenitor cells | 0.0221 | GLI2 | 1 |
| **Cell Morphology,Nervous System Development and Function,Neurological Disease,Tissue Morphology** | Loss of cholinergic fibers | 0.0221 | ESR1 | 1 |
| **Cell Death and Survival** | Activation-induced cell death of T lymphocytes | 0.0221 | XIAP | 1 |
| **Developmental Disorder,Neurological Disease,Organismal Injury and Abnormalities** | Hypoplasia of diencephalon | 0.0221 | PHGDH | 1 |
| **Inflammatory Response,Neurological Disease,Organismal Injury and Abnormalities** | Inflammation of dorsal column of spinal cord | 0.0221 | ESR1 | 1 |
| **Cell Morphology,Cellular Assembly and Organization,Cellular Function and Maintenance** | Extension of peripheral lamellae | 0.0221 | ITGB1 | 1 |
| **Cellular Movement,Nervous System Development and Function** | Delay in migration of Schwann cells | 0.0221 | ITGB1 | 1 |
| **Cellular Development,Cellular Growth and Proliferation,Nervous System Development and Function,Tissue Development** | Development of parasympathetic neurons | 0.0221 | RET | 1 |
| **Carbohydrate Metabolism,Cellular Function and Maintenance,Small Molecule Biochemistry** | Homeostasis of D-glucose | 0.0221 | FOXO1 | 1 |
| **Cancer,Neurological Disease,Organismal Injury and Abnormalities** | Tumorigenesis of ganglioneuroma | 0.0221 | RET | 1 |
| **Cellular Assembly and Organization,Cellular Development,Cellular Growth and Proliferation,Embryonic Development,Nervous System Development and Function,Organ Development,Organismal Development,Tissue Development** | Delay in initiation of development of olfactory glomeruli | 0.0221 | TNC | 1 |
| **Cellular Function and Maintenance,Skeletal and Muscular System Development and Function** | Depolarization of smooth muscle | 0.0221 | TRPC6 | 1 |
| **Cell-To-Cell Signaling and Interaction,Nervous System Development and Function** | Firing of C fiber | 0.0221 | NTRK1 | 1 |
| **Cellular Assembly and Organization,Cellular Movement,Nervous System Development and Function** | Segregation of sensory axons | 0.0221 | ITGB1 | 1 |
| **Cell Morphology,Cell-To-Cell Signaling and Interaction,Cellular Function and Maintenance,Nervous System Development and Function** | Depolarization of hair cells | 0.0221 | MYO7A | 1 |
| **Cellular Development,Embryonic Development,Organismal Development,Tissue Development** | Differentiation of intermediate progenitor cells | 0.0221 | LEF1 | 1 |
| **Cell-To-Cell Signaling and Interaction,Nervous System Development and Function** | Firing of Adelta fiber | 0.0221 | NTRK1 | 1 |

| | | | | |
|---|---|---|---|---|
| **Endocrine System Development and Function,Lipid Metabolism,Small Molecule Biochemistry** | Binding of estrogen | 0.0221 | ESR1 | 1 |
| **Embryonic Development,Tissue Morphology** | Deformation of floor plate | 0.0221 | GLI1 | 1 |
| **Cell Morphology,Nervous System Development and Function,Tissue Morphology** | Surface area of dendrites | 0.0221 | DISC1 | 1 |
| **Endocrine System Development and Function,Endocrine System Disorders,Nervous System Development and Function,Neurological Disease,Organ Morphology,Organismal Development,Organismal Injury and Abnormalities,Reproductive System Disease** | Abnormal morphology of enlarged adenohypophysis | 0.0221 | TCF7L2 | 1 |
| **Cell-To-Cell Signaling and Interaction,Nervous System Development and Function** | Long-term potentiation of medial vestibular nucleus | 0.0221 | CYP19A1 | 1 |
| **Cellular Assembly and Organization,Nervous System Development and Function,Tissue Morphology** | Quantity of postsynaptic membrane | 0.0221 | UTRN | 1 |
| **Cell-To-Cell Signaling and Interaction,Nervous System Development and Function** | Clustering of commissural neurons | 0.0221 | GLI2 | 1 |
| **Nervous System Development and Function,Neurological Disease,Organ Morphology,Organismal Development,Organismal Injury and Abnormalities** | Lack of mesencephalic tract of the trigeminal nerve | 0.0221 | LEF1 | 1 |
| **Cellular Function and Maintenance,Cellular Movement,Nervous System Development and Function** | Mislocalization of Purkinje cells | 0.0221 | ZNF423 | 1 |
| **Cell Morphology,Cellular Assembly and Organization,Cellular Development,Cellular Function and Maintenance,Cellular Growth and Proliferation,Nervous System Development and Function,Organismal Development,Tissue Development** | Arborization of spinal nerve | 0.0221 | ITGB1 | 1 |
| **Cell Morphology,Cellular Assembly and Organization,Cellular Development,Cellular Function and Maintenance,Cellular Growth and Proliferation,Embryonic Development,Nervous System Development and Function,Organismal Development,Tissue Development,Tissue Morphology** | Density of synaptic spine | 0.0221 | CYP19A1 | 1 |
| **Cardiovascular Disease,Hematological Disease,Metabolic Disease** | Hypertriglyceridemia | 0.0221 | FOXO1 | 1 |
| **Cell Morphology,Nervous System Development and Function** | Projection of commissural neurons | 0.0221 | GLI2 | 1 |
| **Amino Acid Metabolism** | Quantity of L-serine | 0.0221 | PHGDH | 1 |
| **Hematological System Development and Function,Inflammatory Response,Tissue Morphology** | Quantity of bone marrow-derived neutrophils | 0.0221 | CNR2 | 1 |
| **Embryonic Development,Tissue Morphology** | Quantity of neuronal progenitor cells | 0.0235 | GLI1, GLI2 | 2 |
| **Reproductive System Development and Function** | Fertility | 0.0235 | ESR1, ITGB1 | 2 |
| **Developmental Disorder,Neurological Disease,Organismal Injury and Abnormalities** | Hypoplasia of brain | 0.0243 | IFT88, PHGDH, SLC6A3, ZNF423 | 4 |
| **Nervous System Development and Function,Organ Morphology,Organismal Development** | Morphology of telencephalon | 0.0253 | B2M, CD81, CLCN3, CNR2, COL18A1, GLI1, GLI2, PHGDH, ROBO1, ZNF423 | 10 |
| **Nervous System Development and Function,Tissue Morphology** | Quantity of dopaminergic neurons | 0.0256 | CDNF, GLI2, SLC6A3 | 3 |
| **Embryonic Development,Nervous System Development and Function,Organ Development,Organismal Development,Tissue Development** | Development of spinal cord | 0.0277 | ERCC2, GLI1, GLI2, PHGDH | 4 |
| **Cell Death and Survival,Cellular Function and Maintenance** | Self-renewal of cells | 0.028 | GLI1, GLI2, ITGB1 | 3 |

| | | | | |
|---|---|---|---|---|
| **Organismal Development,Organismal Injury and Abnormalities** | Abnormal morphology of head | 0.0285 | AXL, B2M, C3, CD81, CLCN3, COL18A1, CP, ESR1, GLI1, GLI2, LEF1, PEG3, PHGDH, REST, ROBO1, SIL1, SLC44A2, SLC6A3, SLITRK6, TCF7L2, YES1, ZNF423 | 22 |
| **Nervous System Development and Function,Neurological Disease,Organ Morphology,Organismal Development,Organismal Injury and Abnormalities** | Abnormal morphology of telencephalon | 0.0286 | B2M, CD81, CLCN3, COL18A1, GLI1, GLI2, PHGDH, ROBO1, ZNF423 | 9 |
| **Cancer,Organismal Injury and Abnormalities** | Frequency of tumor | 0.0312 | FAT1, GLI1, GLI2, RET | 4 |
| **Developmental Disorder,Neurological Disease,Organismal Injury and Abnormalities** | Congenital malformation of brain | 0.032 | IFT88, NOTCH4, PHGDH, SLC6A3, ZNF423 | 5 |
| **Neurological Disease,Organismal Injury and Abnormalities** | Brain lesion | 0.0327 | ANO2, C2orf40, CCDC80, COL11A1, COL18A1, DOCK11, ESR1, FAT1, GFM2, GLI1, GLI2, GPRIN2, GSN, LIMK2, NOTCH4, PTPRZ1, RET, RYR1, SCN7A, SCN9A, SLC6A3, TTLL3, XIAP, ZNF423 | 24 |
| **Nervous System Development and Function,Neurological Disease,Organ Morphology,Organismal Development,Organismal Injury and Abnormalities** | Abnormal morphology of brain | 0.0329 | AXL, B2M, C3, CD81, CLCN3, COL18A1, CP, ESR1, GLI1, GLI2, LEF1, PEG3, PHGDH, REST, ROBO1, SIL1, SLC6A3, TCF7L2, YES1, ZNF423 | 20 |
| **Embryonic Development,Organismal Development** | Growth of embryo | 0.033 | CNR2, DISC1, FOXO1, GLI1, GLI2, HUWE1, IFT88, ITGB1, ORC3 | 9 |
| **Cellular Compromise,Neurological Disease,Organismal Injury and Abnormalities** | Neurodegeneration of cerebral cortex cells | 0.0331 | CLCN3, NFIL3, XIAP | 3 |

| | | | | |
|---|---|---|---|---|
| **Cancer,Neurological Disease,Organismal Injury and Abnormalities** | Brain tumor | 0.035 | ANO2, C2orf40, CCDC80, COL11A1, COL18A1, DOCK11, FAT1, GFM2, GLI1, GLI2, GPRIN2, GSN, LIMK2, NOTCH4, PTPRZ1, RET, RYR1, SCN7A, SCN9A, SLC6A3, TTLL3, ZNF423 | 22 |
| **Nervous System Development and Function,Organ Morphology,Organismal Development** | Size of mesencephalon | 0.0372 | FTO, IRX6 | 2 |
| **Cell Morphology** | Morphology of cilia | 0.0372 | FTO, IFT88 | 2 |
| **Endocrine System Development and Function,Nervous System Development and Function,Organ Morphology,Tissue Morphology** | Quantity of somatotrophs | 0.0372 | PGR, SLC6A3 | 2 |
| **Neurological Disease,Organismal Injury and Abnormalities,Psychological Disorders** | Brain damage | 0.0384 | CNR2, CYP19A1, DTNA, GSN, NFIL3, SNTA1, XIAP | 7 |
| **Embryonic Development,Organismal Development,Tissue Development** | Growth of embryonic tissue | 0.0385 | CNR2, DISC1, GLI1, GLI2, HUWE1, IFT88, ITGB1, ORC3 | 8 |
| **Cardiovascular System Development and Function,Organismal Development** | Vasculogenesis | 0.0387 | GLI1, GSN, NOTCH4 | 3 |
| **Cell Morphology** | Morphology of cellular protrusions | 0.0397 | C3, CLCN6, CNR2, DISC1, ESR1, FTO, IFT88, PTPRZ1, RET, ROBO1, SERPINA3, ZNF423 | 12 |
| **Cancer,Neurological Disease,Organismal Injury and Abnormalities** | Development of primitive neuroectodermal tumor | 0.0416 | FAT1, GLI1, GLI2 | 3 |
| **Nervous System Development and Function,Neurological Disease,Organ Morphology,Organismal Development,Organismal Injury and Abnormalities** | Abnormal morphology of cerebral ventricles | 0.0423 | B2M, CLCN3, COL18A1, CP, PHGDH, ZNF423 | 6 |
| **Cancer,Neurological Disease,Organismal Injury and Abnormalities** | Development of medulloblastoma | 0.0424 | GLI1, GLI2 | 2 |
| **Cell-To-Cell Signaling and Interaction,Nervous System Development and Function** | Firing of central nervous system cells | 0.0424 | CNR2, SCN9A | 2 |
| **Cancer,Endocrine System Disorders,Organismal Injury and Abnormalities,Reproductive System Disease** | Hyperplasia of pituitary gland | 0.0424 | CYP19A1, TCF7L2 | 2 |
| **Cellular Movement,Nervous System Development and Function** | Migration of oligodendrocytes | 0.0424 | ITGB1, TNC | 2 |
| **Nervous System Development and Function,Neurological Disease,Organ Morphology,Organismal Development,Organismal Injury and Abnormalities** | Abnormal morphology of hippocampal commissure | 0.0424 | ROBO1, ZNF423 | 2 |
| **Cancer,Organismal Injury and Abnormalities** | Incidence of tumor | 0.0435 | FAT1, GLI1, GLI2, RET | 4 |

| | | | | |
|---|---|---|---|---|
| **Cell Morphology,Nervous System Development and Function,Neurological Disease,Tissue Morphology** | Abnormal morphology of parasympathetic neurons | 0.0438 | CPEB2 | 1 |
| **Cellular Development,Embryonic Development,Nervous System Development and Function,Organismal Development,Tissue Development** | Differentiation of cerebellar granule neuron precursor cells | 0.0438 | HUWE1 | 1 |
| **Cellular Movement,Nervous System Development and Function** | Guidance of corticocortical axons | 0.0438 | ROBO1 | 1 |
| **Cell Signaling,Small Molecule Biochemistry** | Generation of nitrite | 0.0438 | ITGB1 | 1 |
| **Auditory Disease,Cell Death and Survival,Cellular Compromise,Neurological Disease,Organismal Injury and Abnormalities,Tissue Morphology** | Neurodegeneration of spiral ganglion cells | 0.0438 | RET | 1 |
| **Cell-To-Cell Signaling and Interaction,Hematological System Development and Function,Immune Cell Trafficking,Inflammatory Response** | Adhesion of bone marrow-derived dendritic cells | 0.0438 | ITGB1 | 1 |
| **Nervous System Development and Function,Neurological Disease,Organ Morphology,Organismal Development,Organismal Injury and Abnormalities** | Abnormal morphology of entorhinal area | 0.0438 | CLCN3 | 1 |
| **Cell-To-Cell Signaling and Interaction,Nervous System Development and Function** | Activation of parasympathetic nerve | 0.0438 | REST | 1 |
| **Cell-To-Cell Signaling and Interaction,Drug Metabolism,Molecular Transport,Small Molecule Biochemistry** | Clearance of dopamine | 0.0438 | SLC6A3 | 1 |
| **Nervous System Development and Function,Organ Morphology,Organismal Development** | Size of hippocampal commissure | 0.0438 | ROBO1 | 1 |
| **Cellular Assembly and Organization,Cellular Development,Cellular Growth and Proliferation,Nervous System Development and Function,Tissue Development** | Growth of perikaryon | 0.0438 | RET | 1 |
| **Cellular Development,Cellular Growth and Proliferation,Nervous System Development and Function** | Expansion of satellite cells | 0.0438 | ITGB1 | 1 |
| **Hereditary Disorder,Neurological Disease,Organismal Injury and Abnormalities,Psychological Disorders** | Frontotemporal dementia with parkinsonism | 0.0438 | SERPINA3 | 1 |
| **Cell Cycle,Embryonic Development,Nervous System Development and Function** | Cell division of neuroblasts | 0.0438 | LEF1 | 1 |
| **Cell Morphology,Nervous System Development and Function** | Projection of olfactory receptor neurons | 0.0438 | ROBO1 | 1 |
| **Cell-To-Cell Signaling and Interaction,Nervous System Development and Function** | Firing of dorsal root ganglion cells | 0.0438 | SCN9A | 1 |
| **Cell Cycle** | Neurite outgrowth stage of dorsal root ganglion cells | 0.0438 | ITGB1 | 1 |
| **Cellular Assembly and Organization,Cellular Development,Cellular Growth and Proliferation,Nervous System Development and Function,Tissue Development** | Expansion of growth cone | 0.0438 | RET | 1 |
| **Infectious Diseases** | Pneumococcal infection | 0.0438 | C3 | 1 |
| **Neurological Disease,Organismal Injury and Abnormalities,Psychological Disorders** | Corticobasal degeneration | 0.0438 | SERPINA3 | 1 |
| **Organismal Functions** | Recovery of mice | 0.0438 | ITGB1 | 1 |
| **Cellular Movement,Nervous System Development and Function** | Guidance of corticothalamic axons | 0.0438 | ROBO1 | 1 |
| **Cellular Assembly and Organization,Nervous System Development and Function,Tissue Morphology** | Quantity of Merkel nerve endings | 0.0438 | NTRK1 | 1 |
| **Cell Morphology,Nervous System Development and Function,Tissue Morphology** | Size of cholinergic neurons | 0.0438 | NTRK1 | 1 |
| **Nervous System Development and Function** | Pathfinding of olfactory receptor neurons | 0.0438 | ROBO1 | 1 |
| **Nervous System Development and Function,Tissue Morphology** | Quantity of sympathetic preganglionic neurons | 0.0438 | NTRK1 | 1 |

| | | | | |
|---|---|---|---|---|
| **Cellular Compromise,Cellular Function and Maintenance,Organismal Injury and Abnormalities** | Endoplasmic reticulum stress response of Purkinje cells | 0.0438 | SIL1 | 1 |
| **Cell Death and Survival,Cellular Compromise,Neurological Disease,Organismal Injury and Abnormalities,Psychological Disorders,Tissue Morphology** | Damage of medium spiny neurons | 0.0438 | CNR2 | 1 |
| **Cellular Development,Cellular Growth and Proliferation,Embryonic Development,Nervous System Development and Function,Organ Development,Organismal Development,Tissue Development** | Proliferation of cortical precursor cells | 0.0438 | CNR2 | 1 |
| **Cell-To-Cell Signaling and Interaction,Nervous System Development and Function** | Action potential of AgRP neurons | 0.0438 | FOXO1 | 1 |
| **Cellular Movement,Nervous System Development and Function** | Guidance of corticofugal fibers | 0.0438 | ROBO1 | 1 |
| **Cell Death and Survival,Cellular Development,Cellular Function and Maintenance,Embryonic Development,Organismal Development,Tissue Development** | Self-renewal of neurosphere cells | 0.0438 | GLI1 | 1 |
| **Nervous System Development and Function** | Stabilization of postsynaptic density | 0.0438 | DTNA | 1 |
| **Nervous System Development and Function** | Quantity of enteric ganglion | 0.0438 | RET | 1 |
| **Cellular Function and Maintenance,Nervous System Development and Function,Tissue Development** | Function of cholinergic neurons | 0.0438 | NTRK1 | 1 |
| **Cell Morphology** | Length of cilia | 0.0438 | FTO | 1 |
| **Cardiovascular Disease,Developmental Disorder,Neurological Disease,Organismal Injury and Abnormalities** | Intracranial arteriovenous malformation | 0.0438 | NOTCH4 | 1 |
| **Cellular Movement,Nervous System Development and Function** | Guidance of corticospinal axon | 0.0438 | ROBO1 | 1 |
| **Cell Death and Survival,Cell Morphology,Nervous System Development and Function,Neurological Disease,Tissue Morphology** | Loss of otic neurons | 0.0438 | RET | 1 |
| **Nervous System Development and Function,Tissue Morphology** | Quantity of serotonergic neurons | 0.0438 | GLI2 | 1 |
| **Cell Death and Survival,Cell Morphology,Nervous System Development and Function,Neurological Disease,Tissue Morphology** | Loss of neurons | 0.0455 | CLCN3, CNR2, COL25A1, NTRK1, PTPRC, RET, SHC2, SIL1 | 8 |
| **Cell Morphology,Nervous System Development and Function,Neurological Disease,Tissue Morphology** | Loss of neurites | 0.0479 | C3, ESR1, SERPINA3 | 3 |

### Cluster 3

| Categories | Diseases or Functions Annotation | p-Value | Molecules | # Molecules |
|---|---|---|---|---|
| **Cellular Assembly and Organization,Cellular Function and Maintenance** | Microtubule dynamics | 2.92E-08 | ABI1, ADCY1, ADCY6, ADGRB3, AGRN, APBA1, APC, ARHGEF28, ATCAY, BCL11B, BSN, CAMSAP1, CDKL3, CELSR2, COL25A1, CRMP1, CYFIP1, DBN1, DCC, DVL1, EIF2AK4, | 58 |

| | | | | |
|---|---|---|---|---|
| | | | EIF4G3, ELK1, ERC2, FMN1, GABRB3, GAP43, GNAO1, GPRIN1, HERC1, HRAS, HTR1A, KIF3C, KNDC1, LAMB1, LRRK2, MEF2A, MRTFA, MYO16, MYO5B, NIN, NRN1, PALLD, PIP5K1C, PTGS2, PTK2B, PTPRZ1, ROBO1, ROCK2, RYR2, SEMA4F, SEMA5B, SHANK2, SIPA1L1, SORBS2, TNR, TRPC6, ULK4 | |
| **Cell Morphology,Cellular Assembly and Organization,Cellular Development,Cellular Function and Maintenance,Cellular Growth and Proliferation,Nervous System Development and Function,Organismal Development,Tissue Development** | Neuritogenesis | 5.71E-08 | ABI1, ADCY1, ADGRB3, AGRN, APBA1, APC, ARHGEF28, ATCAY, BCL11B, BSN, CAMSAP1, CDKL3, CELSR2, COL25A1, CRMP1, CYFIP1, DBN1, DCC, DVL1, EIF2AK4, EIF4G3, ELK1, ERC2, FMN1, GABRB3, GAP43, GNAO1, GPRIN1, HERC1, HRAS, KNDC1, LAMB1, LRRK2, MEF2A, MRTFA, MYO16, MYO5B, NIN, NRN1, PALLD, PIP5K1C, | 52 |

| | | | | |
|---|---|---|---|---|
| | | | PTGS2, PTK2B, PTPRZ1, ROBO1, ROCK2, RYR2, SHANK2, SIPA1L1, SORBS2, TNR, TRPC6 | |
| **Cellular Development,Cellular Growth and Proliferation,Nervous System Development and Function,Tissue Development** | Development of neurons | 3.95E-07 | ABI1, ADCY1, ADGRB3, AGRN, APBA1, APC, ARHGEF28, ATCAY, BCL11B, BSN, CAMSAP1, CDKL3, CELSR2, COL25A1, CRMP1, CYFIP1, DBN1, DCC, DVL1, EIF2AK4, EIF4G3, ELK1, ERC2, FMN1, GABRB3, GAP43, GNAO1, GPRIN1, HERC1, HRAS, KNDC1, LAMB1, LRRK2, LRRTM4, MEF2A, MRTFA, MYO16, MYO5B, NCAN, NIN, NRN1, PALLD, PCDH8, PIP5K1C, PLXND1, PTGS2, PTK2B, PTPRZ1, ROBO1, ROCK2, RYR2, SEZ6L, SHANK2, SIPA1L1, SORBS2, TNR, TRPC6 | 57 |
| **Cellular Function and Maintenance,Molecular Transport** | Exocytosis | 0.00000165 | APBA1, APBA2, ERC2, PIP5K1C, PPFIA3, RIMS2, RPH3A, SNAP25, | 10 |

| | | | | |
|---|---|---|---|---|
| | | | UNC13A, UNC13B | |
| **Cellular Assembly and Organization** | Transport of vesicles | 0.00000291 | AP2M1, APBA1, APBA2, APC, DENND1A, KIAA1107, PCLO, PIP5K1C, PPFIA3, RPH3A, SNAP25, UNC13A, UNC13B | 13 |
| **Cellular Assembly and Organization,Cellular Function and Maintenance,Cellular Movement,Molecular Transport,Nervous System Development and Function** | Exocytosis of synaptic vesicles | 0.00000365 | APBA1, APBA2, PIP5K1C, PPFIA3, RPH3A, SNAP25, UNC13A, UNC13B | 8 |
| **Cellular Assembly and Organization,Cellular Movement,Nervous System Development and Function** | Transport of synaptic vesicles | 0.00000535 | AP2M1, APBA1, APBA2, DENND1A, KIAA1107, PCLO, PIP5K1C, PPFIA3, RPH3A, SNAP25, UNC13A, UNC13B | 12 |
| **Cellular Assembly and Organization,Nervous System Development and Function** | Docking of synaptic vesicles | 0.0000396 | PPFIA3, RIMS2, SNAP25, UNC13A, UNC13B | 5 |
| **Cell Morphology,Cellular Assembly and Organization,Cellular Development,Cellular Function and Maintenance,Cellular Growth and Proliferation,Embryonic Development,Nervous System Development and Function,Organismal Development,Tissue Development** | Dendritic growth/branching | 0.0000634 | ABI1, AGRN, APBA1, BSN, CRMP1, DBN1, EIF4G3, ELK1, GABRB3, GAP43, GNAO1, HRAS, KNDC1, MYO5B, NRN1, PTGS2, ROBO1, ROCK2, RYR2, SHANK2, SIPA1L1, SORBS2, TNR | 23 |
| **Cell Morphology** | Shape change of neurites | 0.0000815 | ABI1, AGRN, APBA1, BCL11B, BSN, COL25A1, CRMP1, DBN1, DCC, EIF4G3, ELK1, GABRB3, GAP43, | 27 |

| | | | | |
|---|---|---|---|---|
| | | | GNAO1, HRAS, KNDC1, MYO5B, NR3C2, NRN1, PTGS2, ROBO1, ROCK2, RYR2, SHANK2, SIPA1L1, SORBS2, TNR | |
| **Cell Morphology,Nervous System Development and Function** | Innervation of optic nerve | 0.000263 | DCC, GAP43, VAX1 | 3 |
| **Cancer,Organismal Injury and Abnormalities** | Non-hematological solid tumor | 0.000285 | ABCF1, ADAMTS8, AGAP3, ANKRD30B, APC, ASXL3, ATF7IP2, ATR, COL11A1, CUL9, DHRS4, DNAH8, DPP10, ERC2, HERC1, HRAS, ITGA6, KMT2D, MYO5B, NPR1, P2RX5, PCLO, PIGQ, PIK3R1, PLB1, PRDM2, PROKR2, PTGS2, PTPRT, PTPRZ1, RIMS2, ROS1, RYR2, SEMA4F, SMARCA4, THEMIS, TSHZ2, UNC13B | 38 |
| **Cell Morphology,Cellular Assembly and Organization,Cellular Development,Cellular Function and Maintenance,Cellular Growth and Proliferation,Nervous System Development and Function,Organismal Development,Tissue Development** | Branching of neurons | 0.000446 | ABI1, AGRN, APBA1, BSN, COL25A1, CRMP1, DBN1, EIF4G3, ELK1, ERC2, GABRB3, GAP43, GNAO1, HRAS, KNDC1, MYO5B, NRN1, PTGS2, ROBO1, ROCK2, | 25 |

| | | | | |
|---|---|---|---|---|
| | | | RYR2, SHANK2, SIPA1L1, SORBS2, TNR | |
| **Nervous System Development and Function,Neurological Disease** | Abnormal morphology of nervous system | 0.000468 | ABR, ADCY1, AGRN, ARNT2, CACNA1C, CELSR2, COL25A1, CRMP1, DBN1, DCC, DVL1, E2F5, ERC2, FAT3, FES, GABRB3, GAP43, GMFB, GRIA2, HIVEP2, ITGA6, LRRK2, LRRTM4, MEGF8, MYO16, NOS2, NRN1, PRKN, PROKR2, PTPN13, PTPRZ1, RIMS2, ROBO1, ROCK2, SEMA5B, SHANK2, SHC2, SIL1, SLC4A3, SNAP25, SNTA1, SORBS2, TCF7L2, TENM2, TNR, ULK4, UNC13A, UNC13B, VAX1, VPS13A | 50 |
| **Cancer,Organismal Injury and Abnormalities** | Head and neck tumor | 0.000525 | ABCF1, ADAMTS8, AGAP3, ANKRD30B, APC, ASXL3, ATF7IP2, ATR, COL11A1, CUL9, DHRS4, DNAH8, DPP10, ERC2, HERC1, HRAS, KMT2D, MYO5B, NPR1, P2RX5, PCLO, PIK3R1, PLB1, PRDM2, PROKR2, | 36 |

| | | | | |
|---|---|---|---|---|
| | | | PTGS2, PTPRT, PTPRZ1, RIMS2, ROS1, RYR2, SEMA4F, SMARCA4, THEMIS, TSHZ2, UNC13B | |
| **Cellular Assembly and Organization** | Priming of vesicles | 0.000532 | ERC2, RIMS2, SNAP25, UNC13A, UNC13B | 5 |
| **Cancer,Neurological Disease,Organismal Injury and Abnormalities** | Brain astrocytoma | 0.000635 | ABCF1, ADAMTS8, AGAP3, ANKRD30B, APC, ASXL3, ATF7IP2, ATR, COL11A1, DNAH8, DPP10, ERC2, HERC1, HRAS, MYO5B, P2RX5, PCLO, PIK3R1, PLB1, PRDM2, PROKR2, PTGS2, PTPRZ1, RIMS2, ROS1, RYR2, SEMA4F, THEMIS, TSHZ2 | 29 |
| **Cell Morphology,Cellular Assembly and Organization,Cellular Development,Cellular Function and Maintenance,Cellular Growth and Proliferation,Embryonic Development,Nervous System Development and Function,Organismal Development,Tissue Development** | Branching of neurites | 0.000745 | ABI1, AGRN, APBA1, BSN, COL25A1, CRMP1, DBN1, EIF4G3, ELK1, GABRB3, GAP43, GNAO1, HRAS, KNDC1, MYO5B, NRN1, PTGS2, ROBO1, ROCK2, RYR2, SHANK2, SIPA1L1, SORBS2, TNR | 24 |
| **Cancer,Organismal Injury and Abnormalities** | Non-melanoma solid tumor | 0.000814 | ABCF1, ADAMTS8, AGAP3, ANKRD30B, APC, ASXL3, | 35 |

| | | | | |
|---|---|---|---|---|
| | | | ATF7IP2, ATR, COL11A1, DNAH8, DPP10, ERC2, HERC1, HRAS, ITGA6, KMT2D, MYO5B, NPR1, P2RX5, PCLO, PIK3R1, PLB1, PRDM2, PROKR2, PTGS2, PTPRT, PTPRZ1, RIMS2, ROS1, RYR2, SEMA4F, SMARCA4, THEMIS, TSHZ2, UNC13B | |
| **Cell Morphology,Cellular Assembly and Organization,Cellular Development,Cellular Function and Maintenance,Cellular Growth and Proliferation,Nervous System Development and Function,Organismal Development,Tissue Development** | Formation of dendrites | 0.000831 | ABI1, ADGRB3, CDKL3, CELSR2, DVL1, FMN1, KNDC1, MEF2A, MYO5B, PTPRZ1, ROBO1, ROCK2, SIPA1L1, TRPC6 | 14 |
| **Cancer,Neurological Disease,Organismal Injury and Abnormalities** | Brain tumor | 0.000836 | ABCF1, ADAMTS8, AGAP3, ANKRD30B, APC, ASXL3, ATF7IP2, ATR, COL11A1, DHRS4, DNAH8, DPP10, ERC2, HERC1, HRAS, KMT2D, MYO5B, NPR1, P2RX5, PCLO, PIK3R1, PLB1, PRDM2, PROKR2, PTGS2, PTPRT, PTPRZ1, RIMS2, ROS1, RYR2, SEMA4F, SMARCA4, | 35 |

| | | | | |
|---|---|---|---|---|
| | | | THEMIS, TSHZ2, UNC13B | |
| **Cellular Assembly and Organization** | Translocation of Golgi vesicles | 0.000915 | BSN, PCLO | 2 |
| **Cellular Assembly and Organization,Cellular Function and Maintenance** | Formation of Golgi vesicles | 0.000915 | BSN, PCLO | 2 |
| **Cancer,Neurological Disease,Organismal Injury and Abnormalities** | Nervous system neoplasm | 0.000972 | ABCF1, ADAMTS8, AGAP3, ANKRD30B, APC, ASXL3, ATF7IP2, ATR, COL11A1, DHRS4, DNAH8, DPP10, ERC2, HERC1, HRAS, KMT2D, MYO5B, NPR1, P2RX5, PCLO, PIGQ, PIK3R1, PLB1, PRDM2, PROKR2, PTGS2, PTPRT, PTPRZ1, RIMS2, ROS1, RYR2, SEMA4F, SMARCA4, THEMIS, TSHZ2, UNC13B | 36 |
| **Nervous System Development and Function** | Morphology of nervous system | 0.0013 | ABR, ADCY1, AGRN, AKAP13, APBA1, ARNT2, CACNA1C, CELSR2, COL25A1, CRMP1, DBN1, DCC, DVL1, E2F5, ERC2, FAT3, FES, GABRB3, GAP43, GMFB, GRIA2, HIVEP2, ITGA6, KIF5C, LRRK2, LRRTM4, MEGF8, MYO16, NOS2, NRN1, PLAGL1, PRDM16, PRKN, PROKR2, PTPN13, | 56 |

| | | | | |
|---|---|---|---|---|
| | | | PTPRZ1, RIMS2, ROBO1, ROCK2, SEMA5B, SHANK2, SHC2, SIL1, SIPA1L1, SLC4A3, SNAP25, SNTA1, SORBS2, TCF7L2, TENM2, TNR, ULK4, UNC13A, UNC13B, VAX1, VPS13A | |
| **Cancer,Neurological Disease,Organismal Injury and Abnormalities** | Brain glioma | 0.00136 | ABCF1, ADAMTS8, AGAP3, ANKRD30B, APC, ASXL3, ATF7IP2, ATR, COL11A1, DNAH8, DPP10, ERC2, HERC1, HRAS, KMT2D, MYO5B, NPR1, P2RX5, PCLO, PIK3R1, PLB1, PRDM2, PROKR2, PTGS2, PTPRT, PTPRZ1, RIMS2, ROS1, RYR2, SEMA4F, SMARCA4, THEMIS, TSHZ2 | 33 |
| **Cell-To-Cell Signaling and Interaction,Cellular Assembly and Organization** | Stabilization of synapse | 0.00138 | NCAN, NRN1, TNR | 3 |
| **Cellular Assembly and Organization,Nervous System Development and Function,Tissue Morphology** | Abnormal quantity of synaptic vesicles | 0.00189 | ERC2, PCLO, TNR, UNC13A, UNC13B | 5 |
| **Cell-To-Cell Signaling and Interaction,Cellular Assembly and Organization** | Cell-cell contact | 0.00248 | AGRN, BSN, DCC, ERC2, GAP43, LRRTM4, NCAN, NEURL1, NRN1, PCDH8, PLXND1, ROCK2, SEZ6L, SHANK2, SNTA1, SORBS2, TNR, UNC13A, UNC13B | 19 |

| | | | | |
|---|---|---|---|---|
| **Hereditary Disorder,Neurological Disease,Organismal Injury and Abnormalities,Psychological Disorders,Skeletal and Muscular Disorders** | Familial Parkinson disease | 0.00269 | GAP43, PRKN | 2 |
| **Hereditary Disorder,Neurological Disease,Organismal Injury and Abnormalities,Psychological Disorders,Skeletal and Muscular Disorders** | Huntington disease grade 4 Huntington disease | 0.00269 | RPH3A, SNAP25 | 2 |
| **Nervous System Development and Function,Organismal Development** | Secretion of cerebrospinal fluid | 0.00269 | CELSR2, E2F5 | 2 |
| **Cell Morphology,Nervous System Development and Function,Neurological Disease,Tissue Morphology** | Abnormal morphology of bipolar cells | 0.00273 | ERC2, FAT3, SEMA5B, SLC4A3 | 4 |
| **Cell-To-Cell Signaling and Interaction,Cellular Assembly and Organization,Cellular Function and Maintenance,Molecular Transport** | Priming of synaptic vesicles | 0.00273 | ERC2, RIMS2, UNC13A, UNC13B | 4 |
| **Neurological Disease,Organismal Injury and Abnormalities** | Hydrocephalus | 0.00273 | CELSR2, DVL1, E2F5, ULK4 | 4 |
| **Nervous System Development and Function** | Pathfinding of neurons | 0.00282 | DCC, GAP43, ROBO1 | 3 |
| **Cell-To-Cell Signaling and Interaction,Nervous System Development and Function** | Activation of synapse | 0.00282 | APBA1, APBA2, GRIA2 | 3 |
| **Nervous System Development and Function,Neurological Disease** | Abnormal morphology of ependyma | 0.00282 | E2F5, ULK4, VAX1 | 3 |
| **Embryonic Development,Nervous System Development and Function,Organ Development,Organismal Development,Tissue Development,Visual System Development and Function** | Formation of eye | 0.00282 | ADCY1, FAT3, SLC4A3 | 3 |
| **Cell Morphology,Cell-To-Cell Signaling and Interaction,Cellular Assembly and Organization,Nervous System Development and Function,Tissue Morphology** | Morphology of synapse | 0.00306 | AGRN, ERC2, GAP43, NRN1, ROCK2, SNTA1, TNR, UNC13A, UNC13B | 9 |
| **Tissue Morphology** | Quantity of cells | 0.00341 | ACVR1C, AGRN, BSN, CACNA1C, COL25A1, DCC, DVL1, ERC2, GBA, GNAO1, HIVEP2, HRAS, KIF5C, LRRK2, MGAT5B, NBN, NEURL1, NOS2, NR3C2, NRN1, PCLO, PRDM16, PRKN, PTGS2, PTPRZ1, RIMS2, SHC2, TCF7L2, TNR, TRPM2, UNC13A, UNC13B, VAX1 | 33 |
| **Organismal Development,Organismal Injury and Abnormalities** | Abnormal morphology of head | 0.00401 | ABR, ADCY1, | 32 |

| | | | | |
|---|---|---|---|---|
| | | | ADGRV1, ARNT2, CELSR2, CRMP1, DBN1, DCC, DVL1, E2F5, FAT3, FES, GABRB3, GAP43, GMFB, GRIA2, HIVEP2, ITGA6, LRRTM4, NOS2, PRKN, PROKR2, ROBO1, SHANK2, SIL1, SLC4A3, SNAP25, TCF7L2, TENM2, TNR, ULK4, VAX1 | |
| **Cell-To-Cell Signaling and Interaction,Cellular Assembly and Organization,Cellular Development,Cellular Function and Maintenance,Cellular Growth and Proliferation,Nervous System Development and Function,Tissue Development** | Developmental process of synapse | 0.00441 | AGRN, BSN, DCC, LRRTM4, NCAN, NRN1, PCDH8, PLXND1, SEZ6L, SHANK2, SORBS2, TNR | 12 |
| **Nervous System Development and Function,Tissue Morphology** | Density of neurons | 0.00483 | ACVR1C, CRMP1, ELK1, GABRB3, GAP43, GNAO1, HRAS, LRRTM4, MEF2A, PTGS2, RYR2 | 11 |
| **Nervous System Development and Function,Neurological Disease** | Abnormal morphology of central nervous system | 0.00495 | ABR, ADCY1, ARNT2, CELSR2, CRMP1, DBN1, DCC, E2F5, FES, GABRB3, GAP43, GMFB, GRIA2, HIVEP2, ITGA6, LRRTM4, NOS2, PRKN, PROKR2, PTPRZ1, RIMS2, ROBO1, SHANK2, SIL1, SLC4A3, SNAP25, TCF7L2, TENM2, | 31 |

| | | | | |
|---|---|---|---|---|
| | | | TNR, ULK4, VAX1 | |
| **Nervous System Development and Function,Neurological Disease,Ophthalmic Disease,Organ Morphology,Organismal Development,Organismal Injury and Abnormalities** | Abnormal morphology of visual cortex | 0.00527 | HIVEP2, TENM2 | 2 |
| **Behavior,Nervous System Development and Function** | Working memory | 0.00527 | KCNH3, NR3C2 | 2 |
| **Cell Morphology,Cell-To-Cell Signaling and Interaction,Cellular Assembly and Organization,Nervous System Development and Function,Neurological Disease,Tissue Morphology** | Abnormal morphology of synapse | 0.00548 | AGRN, ERC2, NRN1, ROCK2, SNTA1, TNR, UNC13A, UNC13B | 8 |
| **Cellular Assembly and Organization** | Quantity of vesicles | 0.00671 | BSN, ERC2, PCLO, TNR, UNC13A, UNC13B | 6 |
| **Organismal Development** | Morphology of head | 0.00683 | ABR, ADCY1, ADGRV1, AKAP13, ARNT2, CELSR2, CRMP1, DBN1, DCC, DVL1, E2F5, FAT3, FES, GABRB3, GAP43, GMFB, GRIA2, HIVEP2, ITGA6, KIF5C, LRRTM4, MYO16, NOS2, PRDM16, PRKN, PROKR2, ROBO1, SHANK2, SIL1, SLC4A3, SNAP25, TCF7L2, TENM2, TNR, ULK4, VAX1 | 36 |
| **Cell Morphology** | Projection | 0.0073 | BCL11B, DCC, NR3C2, ROBO1 | 4 |
| **Cellular Assembly and Organization,Cellular Function and Maintenance** | Stabilization of microtubules | 0.00782 | APC, HTR1A, KIF3C | 3 |
| **Nervous System Development and Function,Neurological Disease,Organ Morphology,Organismal Development,Organismal Injury and Abnormalities** | Third ventricle dilation | 0.00782 | CELSR2, E2F5, ULK4 | 3 |
| **Cell Morphology,Nervous System Development and Function,Neurological Disease,Tissue Morphology** | Abnormal morphology of neurons | 0.00819 | AGRN, CACNA1C, COL25A1, CRMP1, DBN1, DVL1, ERC2, FAT3, GAP43, HIVEP2, | 29 |

| | | | | |
|---|---|---|---|---|
| | | | LRRK2, MYO16, NOS2, NRN1, PRKN, PTPN13, ROBO1, ROCK2, SEMA5B, SHANK2, SHC2, SIL1, SLC4A3, SNTA1, SORBS2, TNR, UNC13A, UNC13B, VPS13A | |
| **Cellular Development,Nervous System Development and Function,Tissue Development** | Differentiation of nerve ending | 0.00861 | AGRN, MEF2A | 2 |
| **Hereditary Disorder,Neurological Disease,Organismal Injury and Abnormalities,Psychological Disorders,Skeletal and Muscular Disorders** | Huntington disease grade 3 Huntington disease | 0.00861 | RPH3A, SNAP25 | 2 |
| **Cell Morphology,Cellular Development,Embryonic Development,Nervous System Development and Function,Tissue Development** | Patterning of dendrites | 0.00861 | CELSR2, CRMP1 | 2 |
| **Nervous System Development and Function** | Morphology of central nervous system | 0.00933 | ABR, ADCY1, AKAP13, ARNT2, CELSR2, CRMP1, DBN1, DCC, E2F5, FES, GABRB3, GAP43, GMFB, GRIA2, HIVEP2, ITGA6, KIF5C, LRRTM4, MYO16, NOS2, PRDM16, PRKN, PROKR2, PTPRZ1, RIMS2, ROBO1, SHANK2, SIL1, SLC4A3, SNAP25, TCF7L2, TENM2, TNR, ULK4, VAX1 | 35 |
| **Nervous System Development and Function,Neurological Disease,Organ Morphology,Organismal Development,Organismal Injury and Abnormalities** | Abnormal morphology of brain | 0.0101 | ABR, ADCY1, ARNT2, CELSR2, CRMP1, DBN1, DCC, E2F5, FES, GABRB3, GAP43, GMFB, GRIA2, HIVEP2, ITGA6, LRRTM4, NOS2, | 28 |

| | | | | |
|---|---|---|---|---|
| | | | PRKN, PROKR2, ROBO1, SHANK2, SIL1, SNAP25, TCF7L2, TENM2, TNR, ULK4, VAX1 | |
| **Cell Morphology,Cell-To-Cell Signaling and Interaction,Nervous System Development and Function** | Plasticity of synapse | 0.0106 | DBN1, GRIA2, HRAS, KCNH3, NEURL1, PCDH8, PIP5K1C, PPFIA3, PTK2B, SIPA1L1 | 10 |
| **Cellular Development,Cellular Growth and Proliferation,Nervous System Development and Function,Tissue Development** | Growth of axons | 0.0108 | CRMP1, CYFIP1, DCC, GAP43, GNAO1, KIF3C, LRRK2, MEGF8, NRN1, PROKR2, ROBO1, ROCK2, SNAP25 | 13 |
| **Nervous System Development and Function,Tissue Morphology** | Quantity of neurons | 0.0109 | ACVR1C, AGRN, BSN, CACNA1C, COL25A1, DCC, DVL1, ERC2, GNAO1, HRAS, KIF5C, NBN, NEURL1, NRN1, PCLO, PRKN, PTPRZ1, RIMS2, SHC2, TNR, TRPM2, UNC13A, UNC13B, VAX1 | 24 |
| **Cellular Development,Cellular Growth and Proliferation,Nervous System Development and Function,Tissue Development** | Proliferation of neuronal cells | 0.0117 | ADCY6, AGRN, APBA2, CRMP1, CYFIP1, DCC, GAP43, GNAO1, HRAS, ITGA6, ITGA9, KIF3C, KNDC1, LRRK2, MEGF8, MYO10, NCAN, NRN1, PALLD, PLAGL1, PROKR2, PTK2B, | 28 |

| | | | | |
|---|---|---|---|---|
| | | | PTPRZ1, ROBO1, ROCK2, SEMA5B, SNAP25, VAX1 | |
| **Tissue Development** | Organization of nervous tissue | 0.0122 | AGRN, ARHGEF28, DCC, DVL1, LRRTM4, PTPRO, SHANK2, TNR | 8 |
| **Endocrine System Development and Function,Nervous System Development and Function,Organ Morphology,Tissue Morphology** | Quantity of lutenizing hormone-releasing hormone neurons | 0.0127 | DCC, VAX1 | 2 |
| **Nervous System Development and Function** | Depotentiation | 0.0127 | ADCY1, GRIA2 | 2 |
| **Cell Morphology,Nervous System Development and Function** | Innervation | 0.0128 | DCC, GABRB3, GAP43, GRIA2, RIMS2, SEZ6L, TENM2, VAX1 | 8 |
| **Amino Acid Metabolism,Molecular Transport,Small Molecule Biochemistry** | Uptake of amino acids | 0.0137 | GAP43, SLC32A1, TNR | 3 |
| **Cellular Movement,Nervous System Development and Function** | Guidance of axons | 0.0144 | DCC, DVL1, GAP43, HRAS, KIF5C, LMTK2, PIK3R1, PTPRO, RANBP9, ROBO1, SEMA4F, UNC5D, VAX1 | 13 |
| **Nervous System Development and Function,Organ Morphology,Organismal Development** | Morphology of brain | 0.0146 | ABR, ADCY1, AKAP13, ARNT2, CELSR2, CRMP1, DBN1, DCC, E2F5, FES, GABRB3, GAP43, GMFB, GRIA2, HIVEP2, ITGA6, KIF5C, LRRTM4, MYO16, NOS2, PRDM16, PRKN, PROKR2, ROBO1, SHANK2, SIL1, SNAP25, TCF7L2, TENM2, TNR, ULK4, VAX1 | 32 |
| **Cellular Compromise,Neurological Disease,Organismal Injury and Abnormalities** | Neurodegeneration of cerebral cortex cells | 0.0152 | GRIA2, LRRK2, PRKN, PTGS2 | 4 |

| | | | | |
|---|---|---|---|---|
| **Behavior** | Learning | 0.0162 | ADCY1, ADGRB3, GABRB3, KCNH3, KMT2D, NEURL1, NR3C2 | 7 |
| **Cellular Development,Cellular Growth and Proliferation,Nervous System Development and Function,Tissue Development** | Outgrowth of neurons | 0.0169 | ADCY6, APBA2, DCC, GAP43, GNAO1, HRAS, ITGA6, ITGA9, KIF3C, LRRK2, MEGF8, MYO10, NRN1, PALLD, PROKR2, PTK2B, PTPRZ1, ROBO1, ROCK2, SEMA5B | 20 |
| **Cell Signaling,Nucleic Acid Metabolism,Small Molecule Biochemistry** | Binding of guanosine 5'-O-(3-thiotriphosphate) | 0.0174 | GNAO1, HTR1A | 2 |
| **Cellular Assembly and Organization** | Clustering of synaptic vesicles | 0.0174 | BSN, PCLO | 2 |
| **Cellular Movement,Nervous System Development and Function** | Pathfinding of axons | 0.0174 | DCC, GAP43 | 2 |
| **Cell-To-Cell Signaling and Interaction,Drug Metabolism,Small Molecule Biochemistry** | Metabolism of dopamine | 0.0174 | PRKN, SORCS2 | 2 |
| **Cell Morphology,Cellular Assembly and Organization,Nervous System Development and Function** | Projection of axons | 0.0186 | BCL11B, DCC, NR3C2 | 3 |
| **Nervous System Development and Function,Neurological Disease,Organ Morphology,Organismal Development,Organismal Injury and Abnormalities** | Abnormal morphology of third cerebral ventricle | 0.0187 | ARNT2, CELSR2, E2F5, ULK4 | 4 |
| **Cellular Development,Cellular Growth and Proliferation,Nervous System Development and Function,Tissue Development** | Growth of neurites | 0.0187 | ADCY6, APBA2, CRMP1, CYFIP1, DCC, GAP43, GNAO1, HRAS, ITGA9, KIF3C, KNDC1, LRRK2, MEGF8, MYO10, NCAN, NRN1, PALLD, PROKR2, PTK2B, PTPRZ1, ROBO1, ROCK2, SEMA5B, SNAP25 | 24 |
| **Nervous System Development and Function** | Sensory system development | 0.0203 | ADCY1, DVL1, FAT3, KIF5C, SHC2, SLC4A3 | 6 |
| **Cell Morphology,Cellular Assembly and Organization,Cellular Development,Cellular** | Axonogenesis | 0.021 | ADCY1, AGRN, APC, | 15 |

| | | | | |
|---|---|---|---|---|
| **Function and Maintenance,Cellular Growth and Proliferation,Nervous System Development and Function,Organismal Development,Tissue Development** | | | ARHGEF28, BCL11B, COL25A1, CYFIP1, DCC, DVL1, GAP43, NIN, NRN1, PIP5K1C, PTPRZ1, ROBO1 | |
| **Cellular Development,Cellular Growth and Proliferation,Nervous System Development and Function,Tissue Development** | Outgrowth of axons | 0.0214 | DCC, GAP43, KIF3C, LRRK2, MEGF8, NRN1, PROKR2, ROBO1 | 8 |
| **Cell Morphology,Nervous System Development and Function,Tissue Morphology** | Morphology of neurons | 0.0221 | AGRN, APBA1, CACNA1C, COL25A1, CRMP1, DBN1, DVL1, ERC2, FAT3, GAP43, HIVEP2, LRRK2, MYO16, NOS2, NRN1, PLAGL1, PRKN, PTPN13, RIMS2, ROBO1, ROCK2, SEMA5B, SHANK2, SHC2, SIL1, SIPA1L1, SLC4A3, SNTA1, SORBS2, TNR, UNC13A, UNC13B, VPS13A | 33 |
| **Cell-To-Cell Signaling and Interaction,Nervous System Development and Function** | Action potential of motor endplates | 0.0227 | RIMS2, SNAP25 | 2 |
| **Cell Morphology,Embryonic Development,Nervous System Development and Function,Neurological Disease,Ophthalmic Disease,Organ Development,Organ Morphology,Organismal Development,Organismal Injury and Abnormalities,Tissue Development,Tissue Morphology,Visual System Development and Function** | Abnormal morphology of rod bipolar cells | 0.0227 | FAT3, SLC4A3 | 2 |
| **Cancer,Organismal Injury and Abnormalities** | Frequency of tumor | 0.0233 | CUL9, HRAS, PIGQ, SMARCA4, UNC13B | 5 |
| **Cancer,Organismal Injury and Abnormalities** | Malignant solid tumor | 0.0238 | APC, DHRS4, HRAS, ITGA6, PIGQ, ROS1, SMARCA4 | 7 |
| **Cellular Assembly and Organization,Cellular Function and Maintenance,Cellular** | Endocytosis of synaptic vesicles | 0.0249 | AP2M1, DENND1A, | 4 |

| | | | | |
|---|---|---|---|---|
| **Movement,Nervous System Development and Function** | | | KIAA1107, PIP5K1C | |
| **Embryonic Development,Nervous System Development and Function,Organ Development,Organismal Development,Tissue Development** | Formation of olfactory bulb | 0.025 | GNAO1, LRRK2, PROKR2, RTP1, VAX1 | 5 |
| **Cell Morphology,Cellular Assembly and Organization,Cellular Development,Cellular Function and Maintenance,Cellular Growth and Proliferation,Embryonic Development,Nervous System Development and Function,Organismal Development,Tissue Development,Tissue Morphology** | Density of dendritic spines | 0.0251 | CRMP1, ELK1, GABRB3, GNAO1, HRAS, PTGS2, RYR2 | 7 |
| **Neurological Disease,Organismal Injury and Abnormalities** | Abnormality of cerebral cortex | 0.0261 | ADCY1, CRMP1, DBN1, E2F5, FES, GAP43, GRIA2, HIVEP2, ITGA6, LRRK2, LRRTM4, PRKN, PTGS2, SNAP25, TNR, TP53BP2 | 16 |
| **Cellular Assembly and Organization,Cellular Compromise** | Retraction of cellular protrusions | 0.0268 | DVL1, GRIA2, HRAS, LRRK2, ROCK2 | 5 |
| **Cellular Compromise,Neurological Disease,Organismal Injury and Abnormalities** | Neurodegeneration of hippocampal cells | 0.0277 | GRIA2, PRKN, PTGS2 | 3 |
| **Cellular Development,Cellular Growth and Proliferation,Nervous System Development and Function,Tissue Development** | Outgrowth of neurites | 0.0278 | ADCY6, APBA2, DCC, GAP43, GNAO1, HRAS, ITGA9, KIF3C, LRRK2, MEGF8, MYO10, NRN1, PALLD, PROKR2, PTK2B, PTPRZ1, ROBO1, ROCK2, SEMA5B | 19 |
| **Cell Morphology,Cellular Assembly and Organization,Cellular Function and Maintenance,Nervous System Development and Function** | Extension of axons | 0.0281 | CDKL3, CYFIP1, DBN1, DVL1, MYO5B, SEMA4F, SEMA5B, TNR | 8 |
| **Cell-To-Cell Signaling and Interaction,Nervous System Development and Function** | Long-term potentiation of mossy fibers | 0.0286 | ADCY1, GRIA2 | 2 |
| **Amino Acid Metabolism,Molecular Transport,Small Molecule Biochemistry** | Uptake of L-glutamic acid | 0.0286 | GAP43, TNR | 2 |
| **Lipid Metabolism,Molecular Transport,Small Molecule Biochemistry** | Concentration of arachidonic acid | 0.0286 | ARNTL, PTGS2 | 2 |
| **Free Radical Scavenging,Lipid Metabolism,Molecular Transport,Small Molecule Biochemistry** | Accumulation of lipid peroxide | 0.0303 | KL | 1 |
| **Cellular Development,Nervous System Development and Function,Tissue Development** | Differentiation of sympathetic preganglionic neurons | 0.0303 | AGRN | 1 |

| | | | | |
|---|---|---|---|---|
| **Cancer,Organismal Injury and Abnormalities,Reproductive System Disease** | Prostatic carcinoma | 0.0303 | ITGA6 | 1 |
| **Cellular Assembly and Organization** | Aggregation of early endosomes | 0.0303 | KIF16B | 1 |
| **Cell-To-Cell Signaling and Interaction,Cellular Assembly and Organization,Nervous System Development and Function,Tissue Morphology** | Quantity of ribbon synapse | 0.0303 | BSN | 1 |
| **Nervous System Development and Function,Tissue Morphology** | Density of serotonergic neurons | 0.0303 | GAP43 | 1 |
| **Hereditary Disorder,Neurological Disease,Organismal Injury and Abnormalities,Psychological Disorders,Skeletal and Muscular Disorders** | Autosomal recessive juvenile Parkinson disease | 0.0303 | PRKN | 1 |
| **Cell-To-Cell Signaling and Interaction,Nervous System Development and Function** | Clustering of hippocampal neurons | 0.0303 | SHANK2 | 1 |
| **Cell-To-Cell Signaling and Interaction,Nervous System Development and Function** | Activation of medium spiny neurons | 0.0303 | LRRK2 | 1 |
| **Carbohydrate Metabolism,Lipid Metabolism,Small Molecule Biochemistry** | Incorporation of phosphatidylcholine | 0.0303 | GBA | 1 |
| **Cell Morphology,Cellular Assembly and Organization,Cellular Development,Cellular Function and Maintenance,Cellular Growth and Proliferation,Nervous System Development and Function,Organismal Development,Tissue Development** | Sprouting of rod bipolar cells | 0.0303 | ERC2 | 1 |
| **Cellular Development,Nervous System Development and Function,Tissue Development** | Differentiation of pyramidal neurons | 0.0303 | UNC5D | 1 |
| **Hereditary Disorder,Neurological Disease,Organismal Injury and Abnormalities,Psychological Disorders,Skeletal and Muscular Disorders** | Huntington disease grade 1 Huntington disease | 0.0303 | SNAP25 | 1 |
| **Cell-To-Cell Signaling and Interaction,Nervous System Development and Function** | Conduction of neuroglia | 0.0303 | TNR | 1 |
| **Embryonic Development,Nervous System Development and Function,Organ Development,Organismal Development,Tissue Development** | Development of septal area | 0.0303 | VAX1 | 1 |
| **Developmental Disorder,Endocrine System Disorders,Nutritional Disease,Organismal Injury and Abnormalities,Reproductive System Disease** | Delayed puberty | 0.0303 | VAX1 | 1 |
| **Cellular Assembly and Organization,Cellular Compromise,Nervous System Development and Function** | Retraction of glial cell projections | 0.0303 | GRIA2 | 1 |
| **Behavior** | Perseverance behavior | 0.0303 | NR3C2 | 1 |
| **Behavior** | Impulsivity | 0.0303 | ADCY1 | 1 |
| **Cell Cycle,Nervous System Development and Function** | Arrest in cell cycle progression of oligodendrocyte precursor cells | 0.0303 | HRAS | 1 |
| **Embryonic Development,Nervous System Development and Function,Organ Development,Organismal Development,Tissue Development** | Development of layer IV | 0.0303 | GAP43 | 1 |
| **Behavior** | Risk preference | 0.0303 | GABRB3 | 1 |
| **Metabolic Disease,Neurological Disease,Organismal Injury and Abnormalities,Psychological Disorders** | Stage V Alzheimer disease | 0.0303 | PTGS2 | 1 |
| **Cell-To-Cell Signaling and Interaction,Nervous System Development and Function** | Excitation of postsynaptic region | 0.0303 | PCDH8 | 1 |
| **Cellular Assembly and Organization,Cellular Development,Nervous System Development and Function,Tissue Development** | Differentiation of presynaptic terminals | 0.0303 | AGRN | 1 |
| **Cell-To-Cell Signaling and Interaction,Drug Metabolism,Small Molecule Biochemistry** | Catabolism of dopamine | 0.0303 | PRKN | 1 |
| **Nervous System Development and Function,Neurological Disease,Organ Morphology,Organismal Development,Organismal Injury and Abnormalities** | Lack of optic tract | 0.0303 | GAP43 | 1 |
| **Tissue Development** | Organization of axon initial segments | 0.0303 | DCC | 1 |
| **Cellular Development,Nervous System Development and Function,Tissue Development** | Differentiation of postsynaptic neurons | 0.0303 | AGRN | 1 |

| | | | | |
|---|---|---|---|---|
| **Nervous System Development and Function** | Discharge of cochlear nerve | 0.0303 | BSN | 1 |
| **Lipid Metabolism,Molecular Transport,Small Molecule Biochemistry** | Concentration of eicosapentenoic acid | 0.0303 | ARNTL | 1 |
| **Embryonic Development,Organismal Development,Tissue Morphology** | Morphology of visceral endoderm | 0.0303 | AMOT | 1 |
| **Metabolic Disease,Neurological Disease,Organismal Injury and Abnormalities,Psychological Disorders** | Stage I Alzheimer disease | 0.0303 | PTGS2 | 1 |
| **Cell-To-Cell Signaling and Interaction,Cellular Assembly and Organization,Nervous System Development and Function** | Modulation of synapse | 0.0303 | PCDH8 | 1 |
| **Cell-To-Cell Signaling and Interaction,Cellular Assembly and Organization,Nervous System Development and Function** | Binding of presynaptic membrane | 0.0303 | AP2M1 | 1 |
| **Developmental Disorder,Nervous System Development and Function,Neurological Disease,Organ Morphology,Organismal Development,Organismal Injury and Abnormalities** | Complete agenesis of corpus callosum | 0.0303 | DCC | 1 |
| **Cell-To-Cell Signaling and Interaction,Hematological System Development and Function,Immune Cell Trafficking,Inflammatory Response,Nervous System Development and Function** | Binding of microglia | 0.0303 | TNR | 1 |
| **Carbohydrate Metabolism,Cellular Function and Maintenance,Small Molecule Biochemistry** | Homeostasis of D-glucose | 0.0303 | FOXO1 | 1 |
| **Cell Signaling,Nucleic Acid Metabolism,Small Molecule Biochemistry** | Inhibition of cyclic AMP | 0.0303 | HTR1A | 1 |
| **Nervous System Development and Function** | Function of autonomic nerve | 0.0303 | SNAP25 | 1 |
| **Cell-To-Cell Signaling and Interaction,Nervous System Development and Function** | Inactivation of synapse | 0.0303 | BSN | 1 |
| **Cellular Development,Nervous System Development and Function,Organ Development** | Differentiation of preoptic area | 0.0303 | VAX1 | 1 |
| **Cellular Development,Nervous System Development and Function,Organ Development** | Differentiation of medial ganglionic eminences | 0.0303 | VAX1 | 1 |
| **Cellular Development,Cellular Growth and Proliferation,Embryonic Development,Nervous System Development and Function,Organ Development,Organismal Development,Tissue Development,Visual System Development and Function** | Formation of retinal cells | 0.0303 | ADCY1 | 1 |
| **Molecular Transport,Protein Trafficking** | Translocation of protein fragment | 0.0303 | AGAP3 | 1 |
| **Nervous System Development and Function,Organ Morphology,Organismal Development** | Size of optic chiasm | 0.0303 | GAP43 | 1 |
| **Cellular Development** | Delay in initiation of transition of radial glial cells | 0.0303 | PLAGL1 | 1 |
| **Cellular Development** | Delay in initiation of transition of neuronal progenitor cells | 0.0303 | PLAGL1 | 1 |
| **Nervous System Development and Function** | Curvature of nerves | 0.0303 | DCC | 1 |
| **Cellular Function and Maintenance,Skeletal and Muscular System Development and Function** | Depolarization of smooth muscle | 0.0303 | TRPC6 | 1 |
| **Behavior** | Social exploration | 0.0303 | APBA2 | 1 |
| **Hereditary Disorder,Neurological Disease,Organismal Injury and Abnormalities,Psychological Disorders,Skeletal and Muscular Disorders** | Parkinson-dementia syndrome | 0.0303 | GAP43 | 1 |
| **Lipid Metabolism,Small Molecule Biochemistry** | Incorporation of choline | 0.0303 | GBA | 1 |
| **Neurological Disease** | Mechanical hypersensitivity of neurons | 0.0303 | CACNA2D1 | 1 |
| **Cellular Development,Cellular Growth and Proliferation,Embryonic Development,Nervous System Development and Function,Organismal Development,Tissue Development** | Outgrowth of neuroblasts | 0.0303 | ITGA6 | 1 |
| **Lipid Metabolism,Molecular Transport,Small Molecule Biochemistry** | Accumulation of glucosylceramide | 0.0303 | GBA | 1 |

| | | | | |
|---|---|---|---|---|
| **Cellular Development,Cellular Growth and Proliferation,Nervous System Development and Function,Tissue Development** | Development of corticospinal neurons | 0.0303 | BCL11B | 1 |
| **Metabolic Disease,Neurological Disease,Organismal Injury and Abnormalities,Psychological Disorders** | Preclinical stage Alzheimer disease | 0.0303 | NOS2 | 1 |
| **Hematological Disease** | Hyperemia | 0.0303 | PTGS2 | 1 |
| **Endocrine System Development and Function,Endocrine System Disorders,Nervous System Development and Function,Neurological Disease,Organ Morphology,Organismal Development,Organismal Injury and Abnormalities,Reproductive System Disease** | Abnormal morphology of enlarged adenohypophysis | 0.0303 | TCF7L2 | 1 |
| **Cellular Assembly and Organization** | Delay in initiation of priming of vesicles | 0.0303 | SNAP25 | 1 |
| **Infectious Diseases** | Replication of Murine cytomegalovirus | 0.0303 | NOS2 | 1 |
| **Cell Morphology,Cellular Assembly and Organization,Nervous System Development and Function** | Projection of mossy fibers | 0.0303 | NR3C2 | 1 |
| **Lipid Metabolism,Small Molecule Biochemistry** | Incorporation of sphingomyelin | 0.0303 | GBA | 1 |
| **Amino Acid Metabolism,Lipid Metabolism,Small Molecule Biochemistry** | Translocation of GABA | 0.0303 | SLC32A1 | 1 |
| **Endocrine System Development and Function,Nervous System Development and Function,Organ Morphology,Tissue Morphology** | Quantity of oxytocin neurons | 0.0303 | DCC | 1 |
| **Small Molecule Biochemistry** | Generation of hydrogen sulfide | 0.0303 | HMOX2 | 1 |
| **Cell Morphology,Cellular Movement,Nervous System Development and Function** | Innervation of serotonergic neurons | 0.0303 | GAP43 | 1 |
| **Cell Morphology,Cellular Assembly and Organization** | Elongation of astrocyte projections | 0.0303 | GAP43 | 1 |
| **Cardiovascular Disease,Hematological Disease,Metabolic Disease** | Hypertriglyceridemia | 0.0303 | FOXO1 | 1 |
| **Nervous System Development and Function,Neurological Disease,Ophthalmic Disease** | Abnormal morphology of optic nerve | 0.0311 | DCC, SLC4A3, VAX1 | 3 |
| **Cell Morphology,Cellular Assembly and Organization,Cellular Function and Maintenance,Nervous System Development and Function** | Extension of neurites | 0.0318 | ADCY6, CDKL3, CYFIP1, DBN1, DVL1, KNDC1, MYO5B, SEMA4F, SEMA5B, TNR | 10 |
| **Cancer,Organismal Injury and Abnormalities** | Nonhematologic malignant neoplasm | 0.0324 | APC, DHRS4, ITGA6, PIGQ, ROS1, SMARCA4 | 6 |
| **Cellular Assembly and Organization** | Organization of organelle | 0.0333 | ADGRV1, AGRN, ARHGEF28, DVL1, LRRTM4, PTPRO, SHANK2, TNR | 8 |
| **Embryonic Development,Organismal Development** | Development of head | 0.0339 | ABCB6, ABR, ADCY1, ARNT2, CDK5RAP1, CELSR2, EIF2AK4, FAT3, GAP43, GNAO1, GRHL2, ITGA6, | 29 |

| | | | | |
|---|---|---|---|---|
| | | | LRRK2, MRTFA, MYO16, NBN, NIN, NME7, NOS2, PROKR2, ROBO1, ROGDI, RTP1, SLC4A3, SMARCA4, TACC2, TCF7L2, ULK4, VAX1 | |
| **Nervous System Development and Function,Neurological Disease,Organ Morphology,Organismal Development,Organismal Injury and Abnormalities** | Abnormal morphology of telencephalon | 0.0341 | CELSR2, DCC, E2F5, FES, HIVEP2, NOS2, PROKR2, ROBO1, TENM2, ULK4, VAX1 | 11 |
| **Cancer,Organismal Injury and Abnormalities** | Incidence of tumor | 0.0348 | CUL9, HRAS, PIGQ, SMARCA4, UNC13B | 5 |
| **Cellular Assembly and Organization,Nervous System Development and Function** | Recycling of synaptic vesicles | 0.0348 | DVL1, GAP43, PIP5K1C | 3 |
| **Cellular Assembly and Organization,Cellular Function and Maintenance,Cellular Movement,Nervous System Development and Function** | Axonal transport of mitochondria | 0.035 | PRKN, RANBP9 | 2 |
| **Cell Morphology,Cellular Movement,Nervous System Development and Function** | Innervation of Purkinje cells | 0.035 | GRIA2, SEZ6L | 2 |
| **Cell-To-Cell Signaling and Interaction,Cellular Assembly and Organization,Cellular Development,Cellular Function and Maintenance,Cellular Growth and Proliferation,Nervous System Development and Function,Tissue Development** | Formation of excitatory synapses | 0.035 | LRRTM4, SORBS2 | 2 |
| **Cell Morphology** | Shape change of axons | 0.0367 | AGRN, BCL11B, COL25A1, DCC, GAP43, NR3C2, NRN1 | 7 |
| **Nervous System Development and Function,Organ Morphology,Organismal Development** | Morphology of forebrain | 0.0369 | ADCY1, AKAP13, ARNT2, CELSR2, DCC, E2F5, FES, GAP43, HIVEP2, NOS2, PROKR2, ROBO1, TENM2, ULK4, VAX1 | 15 |
| **Neurological Disease,Organismal Injury and Abnormalities** | Neurodegeneration of cerebral cortex | 0.037 | GRIA2, LRRK2, PRKN, PTGS2, SNAP25 | 5 |
| **Cellular Function and Maintenance** | Endocytosis | 0.037 | AP2M1, DENND1A, KIAA1107, PIP5K1C, SNAP25 | 5 |

| | | | | |
|---|---|---|---|---|
| **Cell-To-Cell Signaling and Interaction,Cellular Assembly and Organization,Cellular Function and Maintenance,Tissue Development** | Organization of synapse | 0.0378 | AGRN, DVL1, LRRTM4, PTPRO, SHANK2, TNR | 6 |
| **Nervous System Development and Function,Neurological Disease,Organ Morphology,Organismal Development,Organismal Injury and Abnormalities** | Abnormal morphology of cerebral cortex | 0.0401 | ADCY1, CRMP1, DBN1, E2F5, FES, GAP43, GRIA2, HIVEP2, ITGA6, LRRTM4, SNAP25, TNR | 12 |
| **Cell Death and Survival,Cellular Compromise,Neurological Disease,Organismal Injury and Abnormalities,Tissue Morphology** | Neurodegeneration of pyramidal neurons | 0.042 | GRIA2, PTGS2 | 2 |
| **Neurological Disease,Organismal Injury and Abnormalities,Psychological Disorders,Skeletal and Muscular Disorders** | Parkinson disease with dementia | 0.042 | GAP43, NOS2 | 2 |
| **Cancer,Organismal Injury and Abnormalities** | Development of benign tumor | 0.042 | HRAS, UNC13B | 2 |
| **Cell-To-Cell Signaling and Interaction,Nervous System Development and Function** | Long-term potentiation | 0.0437 | ADCY1, CHST10, CRMP1, DPP6, EIF2AK4, GBA, GRIA2, KCNH3, NCAN, PCDH8, PTK2B, ROCK2, SHANK2, TNR | 14 |
| **Cell-To-Cell Signaling and Interaction,Nervous System Development and Function** | Synaptic transmission of cells | 0.0439 | BSN, ERC2, GRIA2, HTR1A, NR3C2, SNAP25 | 6 |
| **Nervous System Development and Function,Neurological Disease,Organ Morphology,Organismal Development,Organismal Injury and Abnormalities** | Abnormal morphology of forebrain | 0.0444 | ADCY1, ARNT2, CELSR2, DCC, E2F5, FES, HIVEP2, NOS2, PROKR2, ROBO1, TENM2, ULK4, VAX1 | 13 |
| **Developmental Disorder,Neurological Disease** | Hypoplasia of cranial nerve | 0.0494 | DCC, GABRB3 | 2 |
| **Inflammatory Disease,Inflammatory Response,Neurological Disease,Organismal Injury and Abnormalities** | Inflammation of cerebral cortex | 0.0494 | LRRK2, TP53BP2 | 2 |
| **Auditory and Vestibular System Development and Function,Cell Morphology,Cellular Assembly and Organization** | Orientation of stereocilia bundles | 0.0494 | ADGRV1, DVL1 | 2 |

### Cluster 5

| Categories | Diseases or Functions Annotation | p-Value | Molecules | # Molecules |
|---|---|---|---|---|
| **Cellular Function and Maintenance,Molecular Transport** | Exocytosis | 3.29E-07 | APBA2, CADPS2, ERC2, PIP5K1C, PPFIA3, RIMS2, | 11 |

| | | | | |
|---|---|---|---|---|
| | | | RPH3A, SNAP25, UNC13A, UNC13B, UNC13C | |
| **Cellular Assembly and Organization** | Priming of vesicles | 0.00000538 | CADPS2, ERC2, RIMS2, SNAP25, UNC13A, UNC13B, UNC13C | 7 |
| **Cellular Assembly and Organization,Cellular Function and Maintenance,Cellular Movement,Molecular Transport,Nervous System Development and Function** | Exocytosis of synaptic vesicles | 0.0000058 | APBA2, PIP5K1C, PPFIA3, RPH3A, SNAP25, UNC13A, UNC13B, UNC13C | 8 |
| **Cell-To-Cell Signaling and Interaction,Cellular Assembly and Organization,Cellular Function and Maintenance,Molecular Transport** | Priming of synaptic vesicles | 0.0000286 | CADPS2, ERC2, RIMS2, UNC13A, UNC13B, UNC13C | 6 |
| **Cellular Assembly and Organization,Nervous System Development and Function** | Docking of synaptic vesicles | 0.0000535 | PPFIA3, RIMS2, SNAP25, UNC13A, UNC13B | 5 |
| **Cellular Assembly and Organization,Cellular Movement,Nervous System Development and Function** | Transport of synaptic vesicles | 0.0000543 | APBA2, KIAA1107, PCLO, PIP5K1C, PPFIA3, RPH3A, SCRIB, SNAP25, UNC13A, UNC13B, UNC13C | 11 |
| **Cancer,Neurological Disease,Organismal Injury and Abnormalities** | Brain astrocytoma | 0.000178 | ABCF1, ADAMTS13, ADAMTS8, AGAP3, ANKRD30B, ASPM, ASXL3, ATF7IP2, CREBBP, DPP10, ERC2, GATAD2B, HERC1, MED12, MYO5B, P2RX5, PCLO, PIK3R1, PRDM2, PROKR2, PTGS2, RAB11FIP5, RALY, RIMS2, ROS1, RYR2, SEMA4F, SPTBN4, SYNE1, THEMIS, TSC2, UBQLN3 | 32 |

| | | | | |
|---|---|---|---|---|
| **Cancer,Organismal Injury and Abnormalities** | Non-melanoma solid tumor | 0.000337 | ABCF1, ADAMTS13, ADAMTS8, AGAP3, ANKRD30B, ASPM, ASXL3, ATF7IP2, CREBBP, DPP10, ERC2, GATAD2B, HERC1, MED12, MIIP, MYO15A, MYO5B, P2RX5, PCLO, PIK3R1, PRDM1, PRDM2, PROKR2, PTGS2, PTPRT, RAB11FIP5, RALY, RIMS2, ROS1, RYR2, SEMA4F, SETD2, SPTBN4, SYNE1, THEMIS, TSC2, UBQLN3, UNC13B | 38 |
| **Nervous System Development and Function,Neurological Disease** | Abnormal morphology of nervous system | 0.000357 | ABR, ADCY1, ADGRA2, AGRN, AHI1, ARNT2, ASPM, CADPS2, CELSR2, COL13A1, CREBBP, DVL1, EPHB3, ERC2, FAT3, FES, GAP43, GRIA2, HIVEP2, IDUA, KIF1B, LRRK2, LRRTM4, MAN2C1, MCPH1, MEGF8, MTHFR, NCOR1, NOS2, NRN1, PER2, PRKN, PROKR2, RIMS2, ROCK2, SCN1B, SEMA5B, SHANK2, | 53 |

| | | | | |
|---|---|---|---|---|
| | | | SLC2A8, SMPD1, SNAP25, SORBS2, SORL1, SOX4, SPTBN4, SYNE1, SYNE2, TOP2B, UNC13A, UNC13B, VAX1, VPS13A, ZIC5 | |
| **Cancer,Neurological Disease,Organismal Injury and Abnormalities** | Glioma | 0.000395 | ABCF1, ADAMTS13, ADAMTS8, AGAP3, ANKRD30B, ASPM, ASXL3, ATF7IP2, CREBBP, DPP10, ERC2, GATAD2B, HERC1, MED12, MIIP, MYO15A, MYO5B, P2RX5, PCLO, PIK3R1, PRDM1, PRDM2, PROKR2, PTGS2, PTPRT, RAB11FIP5, RALY, RIMS2, ROS1, RYR2, SEMA4F, SETD2, SPTBN4, SYNE1, THEMIS, TSC2, UBQLN3 | 37 |
| **Cancer,Organismal Injury and Abnormalities** | Head and neck tumor | 0.000439 | ABCF1, ADAMTS13, ADAMTS8, AGAP3, ANKRD30B, ASPM, ASXL3, ATF7IP2, CREBBP, CUL9, DPP10, ERC2, GATAD2B, HERC1, MED12, MYO15A, MYO5B, P2RX5, PCLO, PIK3R1, PRDM1, PRDM2, PROKR2, | 38 |

| | | | | |
|---|---|---|---|---|
| | | | PTGS2, PTPRT, RAB11FIP5, RALY, RIMS2, ROS1, RYR2, SEMA4F, SETD2, SPTBN4, SYNE1, THEMIS, TSC2, UBQLN3, UNC13B | |
| **Cancer,Organismal Injury and Abnormalities** | Non-hematological solid tumor | 0.000497 | ABCF1, ADAMTS13, ADAMTS8, AGAP3, ANKRD30B, ASPM, ASXL3, ATF7IP2, CREBBP, CUL9, DPP10, ERC2, GATAD2B, HERC1, MED12, MIIP, MYO15A, MYO5B, P2RX5, PCLO, PIK3R1, PRDM1, PRDM2, PROKR2, PTGS2, PTPRT, RAB11FIP5, RALY, RIMS2, ROS1, RYR2, SEMA4F, SETD2, SPTBN4, SYNE1, THEMIS, TSC2, UBQLN3, UNC13B | 39 |
| **Cancer,Neurological Disease,Organismal Injury and Abnormalities** | Brain glioma | 0.000538 | ABCF1, ADAMTS13, ADAMTS8, AGAP3, ANKRD30B, ASPM, ASXL3, ATF7IP2, CREBBP, DPP10, ERC2, GATAD2B, HERC1, MED12, MYO15A, MYO5B, P2RX5, PCLO, PIK3R1, PRDM1, PRDM2, | 36 |

| | | | | |
|---|---|---|---|---|
| | | | PROKR2, PTGS2, PTPRT, RAB11FIP5, RALY, RIMS2, ROS1, RYR2, SEMA4F, SETD2, SPTBN4, SYNE1, THEMIS, TSC2, UBQLN3 | |
| **Cancer,Neurological Disease,Organismal Injury and Abnormalities** | Central nervous system solid tumor | 0.000635 | ABCF1, ADAMTS13, ADAMTS8, AGAP3, ANKRD30B, ASPM, ASXL3, ATF7IP2, CREBBP, DPP10, ERC2, GATAD2B, HERC1, MED12, MIIP, MYO15A, MYO5B, P2RX5, PCLO, PIK3R1, PRDM1, PRDM2, PROKR2, PTGS2, PTPRT, RAB11FIP5, RALY, RIMS2, ROS1, RYR2, SEMA4F, SETD2, SPTBN4, SYNE1, THEMIS, TSC2, UBQLN3, UNC13B | 38 |
| **Cancer,Neurological Disease,Organismal Injury and Abnormalities** | Brain tumor | 0.000686 | ABCF1, ADAMTS13, ADAMTS8, AGAP3, ANKRD30B, ASPM, ASXL3, ATF7IP2, CREBBP, DPP10, ERC2, GATAD2B, HERC1, MED12, MYO15A, MYO5B, P2RX5, PCLO, PIK3R1, PRDM1, PRDM2, PROKR2, | 37 |

| | | | | |
|---|---|---|---|---|
| | | | PTGS2, PTPRT, RAB11FIP5, RALY, RIMS2, ROS1, RYR2, SEMA4F, SETD2, SPTBN4, SYNE1, THEMIS, TSC2, UBQLN3, UNC13B | |
| **Nervous System Development and Function** | Morphology of nervous system | 0.000756 | ABR, ADCY1, ADGRA2, AGRN, AHI1, ARNT2, ASPM, CADPS2, CELSR2, COL13A1, CREBBP, DVL1, EPHB3, ERC2, FAT3, FES, GAP43, GRIA2, HIVEP2, IDUA, KIF1B, KIF5C, KNG1, LRRK2, LRRTM4, MAN2C1, MCPH1, MEGF8, MTHFR, NCOR1, NOS2, NRN1, PCM1, PER2, PLAGL1, PRKN, PROKR2, RGS11, RIMS2, ROCK2, SCN1B, SEMA5B, SHANK2, SIPA1L1, SLC2A8, SMPD1, SNAP25, SORBS2, SORL1, SOX4, SPTBN4, SYNE1, SYNE2, TIAM1, TOP2B, UNC13A, UNC13B, VAX1, VPS13A, ZIC5 | 60 |

| | | | | |
|---|---|---|---|---|
| **Organismal Development,Organismal Injury and Abnormalities** | Abnormal morphology of head | 0.0009 | ABR, ADCY1, ADGRA2, ADGRV1, AHI1, ARNT2, ASPM, CADPS2, CELSR2, CREBBP, DVL1, EPHB3, FAT3, FES, GAP43, GRIA2, HIVEP2, IDUA, KIF1B, LRRTM4, MAN2C1, MCPH1, MTHFR, NCOR1, NOS2, PRKN, PROKR2, SHANK2, SLC2A8, SMPD1, SNAP25, SOX4, SPTBN4, TOP2B, VAX1, ZIC5 | 36 |
| **Cellular Assembly and Organization,Cellular Function and Maintenance** | Formation of Golgi vesicles | 0.00104 | BSN, PCLO | 2 |
| **Cellular Assembly and Organization** | Translocation of Golgi vesicles | 0.00104 | BSN, PCLO | 2 |
| **Cell Morphology,Cell-To-Cell Signaling and Interaction,Cellular Assembly and Organization,Nervous System Development and Function,Tissue Morphology** | Morphology of synapse | 0.00131 | AGRN, COL13A1, ERC2, GAP43, NRN1, ROCK2, SYNE1, SYNE2, UNC13A, UNC13B | 10 |
| **Cell-To-Cell Signaling and Interaction,Nervous System Development and Function** | Action potential of motor endplates | 0.00165 | COL13A1, RIMS2, SNAP25 | 3 |
| **Behavior,Nervous System Development and Function** | Short-term memory | 0.00165 | CREBBP, KCNH3, NR3C2 | 3 |
| **Nervous System Development and Function,Neurological Disease** | Abnormal morphology of central nervous system | 0.00207 | ABR, ADCY1, ADGRA2, AHI1, ARNT2, ASPM, CADPS2, CELSR2, CREBBP, EPHB3, FES, GAP43, GRIA2, HIVEP2, IDUA, KIF1B, LRRTM4, MAN2C1, MCPH1, MTHFR, NCOR1, | 34 |

| | | | | |
|---|---|---|---|---|
| | | | NOS2, PRKN, PROKR2, RIMS2, SCN1B, SHANK2, SLC2A8, SMPD1, SNAP25, SPTBN4, TOP2B, VAX1, ZIC5 | |
| **Nervous System Development and Function,Neurological Disease,Organ Morphology,Organismal Development,Organismal Injury and Abnormalities** | Abnormal morphology of brain | 0.00212 | ABR, ADCY1, ADGRA2, AHI1, ARNT2, ASPM, CADPS2, CELSR2, CREBBP, EPHB3, FES, GAP43, GRIA2, HIVEP2, IDUA, KIF1B, LRRTM4, MAN2C1, MCPH1, MTHFR, NCOR1, NOS2, PRKN, PROKR2, SHANK2, SLC2A8, SMPD1, SNAP25, SPTBN4, TOP2B, VAX1, ZIC5 | 32 |
| **Cellular Assembly and Organization,Cellular Function and Maintenance** | Microtubule dynamics | 0.00215 | ABI1, ADCY1, ADCY6, ADGRB3, AGRN, AHI1, ANAPC2, ARHGEF28, ATCAY, BCL11B, BSN, CAMSAP1, CELSR2, DAB2IP, DVL1, ELK1, EPHB3, ERC2, FMN1, GAP43, GPRIN1, HERC1, LAMB1, LRRK2, MEF2A, MRTFA, MYO5B, NIN, NRN1, PIP5K1C, PTGS2, PTK2B, ROCK2, | 45 |

| | | | | |
|---|---|---|---|---|
| | | | RYR2, SCN1B, SEMA4F, SEMA5B, SHANK2, SIPA1L1, SORBS2, SPTBN4, SYNE1, TIAM1, TOP2B, TSC2 | |
| **Cell Morphology,Cell-To-Cell Signaling and Interaction,Cellular Assembly and Organization,Nervous System Development and Function,Neurological Disease,Tissue Morphology** | Abnormal morphology of synapse | 0.00224 | AGRN, COL13A1, ERC2, NRN1, ROCK2, SYNE1, SYNE2, UNC13A, UNC13B | 9 |
| **Neurological Disease,Organismal Injury and Abnormalities** | Brain lesion | 0.00225 | ABCF1, ADAMTS13, ADAMTS8, AGAP3, ANKRD30B, ASPM, ASXL3, ATF7IP2, CREBBP, DPP10, ERC2, GATAD2B, HERC1, KNG1, MED12, MYO15A, MYO5B, P2RX5, PCLO, PIK3R1, PRDM1, PRDM2, PROKR2, PTGS2, PTPRT, RAB11FIP5, RALY, RIMS2, ROS1, RYR2, SEMA4F, SETD2, SPTBN4, SYNE1, THEMIS, TSC2, UBQLN3, UNC13B | 38 |
| **Lipid Metabolism,Molecular Transport,Small Molecule Biochemistry** | Concentration of arachidonic acid | 0.00242 | ARNTL, KNG1, PTGS2 | 3 |
| **Cell Morphology,Cellular Assembly and Organization,Cellular Development,Cellular Function and Maintenance,Cellular Growth and Proliferation,Nervous System Development and Function,Organismal Development,Tissue Development** | Neuritogenesis | 0.00244 | ABI1, ADCY1, ADGRB3, AGRN, ANAPC2, ARHGEF28, ATCAY, BCL11B, BSN, CAMSAP1, CELSR2, DAB2IP, DVL1, | 40 |

| | | | | |
|---|---|---|---|---|
| | | | ELK1, EPHB3, ERC2, FMN1, GAP43, GPRIN1, HERC1, LAMB1, LRRK2, MEF2A, MRTFA, MYO5B, NIN, NRN1, PIP5K1C, PTGS2, PTK2B, ROCK2, RYR2, SHANK2, SIPA1L1, SORBS2, SPTBN4, SYNE1, TIAM1, TOP2B, TSC2 | |
| **Cellular Assembly and Organization,Nervous System Development and Function,Tissue Morphology** | Abnormal quantity of synaptic vesicles | 0.00249 | ERC2, KIF1B, PCLO, UNC13A, UNC13B | 5 |
| **Nervous System Development and Function,Tissue Morphology** | Density of neurons | 0.00269 | ACVR1C, DAB2IP, ELK1, EPHB3, GAP43, LRRTM4, MEF2A, PTGS2, RYR2, SCN1B, TIAM1, TSC2 | 12 |
| **Hereditary Disorder,Neurological Disease,Organismal Injury and Abnormalities,Psychological Disorders,Skeletal and Muscular Disorders** | Huntington disease grade 4 Huntington disease | 0.00304 | RPH3A, SNAP25 | 2 |
| **Hereditary Disorder,Neurological Disease,Organismal Injury and Abnormalities,Psychological Disorders,Skeletal and Muscular Disorders** | Familial Parkinson disease | 0.00304 | GAP43, PRKN | 2 |
| **Neurological Disease,Organismal Injury and Abnormalities** | Hydrocephalus | 0.00342 | CELSR2, DVL1, DVL3, NCOR1 | 4 |
| **Organismal Development** | Morphology of head | 0.00373 | ABR, ADCY1, ADGRA2, ADGRV1, AHI1, ARNT2, ASPM, CADPS2, CELSR2, CREBBP, DVL1, EPHB3, FAT3, FES, GAP43, GRIA2, HIVEP2, IDUA, KIF1B, KIF5C, KNG1, | 39 |

| | | | | |
|---|---|---|---|---|
| | | | LRRTM4, MAN2C1, MCPH1, MTHFR, NCOR1, NOS2, PCM1, PRKN, PROKR2, SHANK2, SLC2A8, SMPD1, SNAP25, SOX4, SPTBN4, TOP2B, VAX1, ZIC5 | |
| **Neurological Disease,Psychological Disorders** | Pervasive developmental disorder | 0.00412 | MATK, RYR2, SLC6A7, TSC2 | 4 |
| **Cellular Development,Cellular Growth and Proliferation,Nervous System Development and Function,Tissue Development** | Development of neurons | 0.00555 | ABI1, ADCY1, ADGRB3, AGRN, ANAPC2, ARHGEF28, ATCAY, BCL11B, BSN, CADPS2, CAMSAP1, CELSR2, DAB2IP, DVL1, ELK1, ELP1, EPHB3, ERC2, FMN1, GAP43, GPRIN1, HERC1, LAMB1, LRRK2, LRRTM4, MEF2A, MRTFA, MYO5B, NIN, NRN1, PIP5K1C, PLXND1, PRDM1, PTGS2, PTK2B, ROCK2, RYR2, SHANK2, SIPA1L1, SORBS2, SPTBN4, SYNE1, TIAM1, TOP2B, TSC2 | 45 |
| **Auditory and Vestibular System Development and Function,Cell Morphology,Cellular Assembly and Organization** | Orientation of stereocilia bundles | 0.00589 | ADGRV1, DVL1, DVL3 | 3 |
| **Behavior,Nervous System Development and Function** | Working memory | 0.00596 | KCNH3, NR3C2 | 2 |
| **Cell Morphology,Nervous System Development and Function,Tissue Morphology** | Morphology of bipolar cells | 0.00678 | ERC2, FAT3, RGS11, SEMA5B | 4 |

| | | | | |
|---|---|---|---|---|
| **Nervous System Development and Function,Organ Morphology,Organismal Development** | Morphology of brain | 0.00739 | ABR, ADCY1, ADGRA2, AHI1, ARNT2, ASPM, CADPS2, CELSR2, CREBBP, EPHB3, FES, GAP43, GRIA2, HIVEP2, IDUA, KIF1B, KIF5C, KNG1, LRRTM4, MAN2C1, MCPH1, MTHFR, NCOR1, NOS2, PCM1, PRKN, PROKR2, SHANK2, SLC2A8, SMPD1, SNAP25, SPTBN4, TOP2B, VAX1, ZIC5 | 35 |
| **Nervous System Development and Function,Neurological Disease,Organ Morphology,Organismal Development,Organismal Injury and Abnormalities** | Abnormal morphology of barrel cortex | 0.00748 | ADCY1, ASPM, GAP43 | 3 |
| **Nervous System Development and Function** | Morphology of central nervous system | 0.00855 | ABR, ADCY1, ADGRA2, AHI1, ARNT2, ASPM, CADPS2, CELSR2, CREBBP, EPHB3, FES, GAP43, GRIA2, HIVEP2, IDUA, KIF1B, KIF5C, KNG1, LRRTM4, MAN2C1, MCPH1, MTHFR, NCOR1, NOS2, PCM1, PRKN, PROKR2, RIMS2, SCN1B, SHANK2, SLC2A8, SMPD1, SNAP25, SPTBN4, TOP2B, VAX1, ZIC5 | 37 |

| | | | | |
|---|---|---|---|---|
| **Cellular Assembly and Organization** | Quantity of vesicles | 0.00901 | BSN, ERC2, KIF1B, PCLO, UNC13A, UNC13B | 6 |
| **Cellular Assembly and Organization,Cellular Compromise** | Retraction of cellular protrusions | 0.00901 | ARHGEF11, DVL1, GRIA2, LRRK2, PITPNM1, ROCK2 | 6 |
| **Cell Morphology,Cellular Function and Maintenance** | Autophagy of cortical neurons | 0.00972 | PRKN, TSC2 | 2 |
| **Cell Morphology,Nervous System Development and Function** | Innervation of optic nerve | 0.00972 | GAP43, VAX1 | 2 |
| **Cellular Development,Nervous System Development and Function,Tissue Development** | Differentiation of nerve ending | 0.00972 | AGRN, MEF2A | 2 |
| **Hereditary Disorder,Neurological Disease,Organismal Injury and Abnormalities,Psychological Disorders,Skeletal and Muscular Disorders** | Huntington disease grade 3 Huntington disease | 0.00972 | RPH3A, SNAP25 | 2 |
| **Cell-To-Cell Signaling and Interaction,Cellular Assembly and Organization** | Cell-cell contact | 0.0104 | AGRN, BSN, COL13A1, EPHB3, ERC2, GAP43, GJD2, LRRTM4, NEURL1, NRN1, PLXND1, ROCK2, SHANK2, SORBS2, SYNE1, SYNE2, UNC13A, UNC13B | 18 |
| **Lipid Metabolism,Molecular Transport,Small Molecule Biochemistry** | Concentration of phospholipid | 0.0136 | PIP5K1C, PTGS2, SMPD1 | 3 |
| **Cell-To-Cell Signaling and Interaction,Nervous System Development and Function** | Long term synaptic depression of CA1 neuron | 0.0136 | GRIA2, PIP5K1C, RAB11FIP5 | 3 |
| **Nervous System Development and Function,Neurological Disease,Organ Morphology,Organismal Development,Organismal Injury and Abnormalities** | Abnormal morphology of cerebral cortex | 0.0139 | ADCY1, ASPM, FES, GAP43, GRIA2, HIVEP2, KIF1B, LRRTM4, MAN2C1, MCPH1, SLC2A8, SMPD1, SNAP25, ZIC5 | 14 |
| **Nervous System Development and Function** | Depotentiation | 0.0143 | ADCY1, GRIA2 | 2 |
| **Amino Acid Metabolism,Endocrine System Development and Function,Molecular Transport,Small Molecule Biochemistry** | Quantity of L-triiodothyronine | 0.0143 | NCOR1, SLCO1C1 | 2 |
| **Nervous System Development and Function,Neurological Disease,Organ Morphology,Organismal Development,Organismal Injury and Abnormalities** | Abnormal morphology of forebrain | 0.0172 | ADCY1, ADGRA2, ARNT2, ASPM, CELSR2, CREBBP, FES, HIVEP2, MAN2C1, NCOR1, NOS2, | 15 |

| | | | | |
|---|---|---|---|---|
| | | | PROKR2, TOP2B, VAX1, ZIC5 | |
| **Cellular Compromise,Neurological Disease,Organismal Injury and Abnormalities** | Neurodegeneration of cerebral cortex cells | 0.0188 | GRIA2, LRRK2, PRKN, PTGS2 | 4 |
| **Lipid Metabolism,Small Molecule Biochemistry** | Synthesis of eicosanoid | 0.0189 | KNG1, NOS2, PTGS2 | 3 |
| **Cellular Assembly and Organization** | Clustering of synaptic vesicles | 0.0196 | BSN, PCLO | 2 |
| **Behavior,Nervous System Development and Function** | Memory consolidation | 0.0196 | CREBBP, NR3C2 | 2 |
| **Cell-To-Cell Signaling and Interaction,Drug Metabolism,Small Molecule Biochemistry** | Metabolism of dopamine | 0.0196 | PRKN, SORCS2 | 2 |
| **Cellular Assembly and Organization,Cellular Compromise,Nervous System Development and Function** | Retraction of neurites | 0.0219 | ARHGEF11, DVL1, LRRK2, PITPNM1, ROCK2 | 5 |
| **Cell Morphology,Cellular Assembly and Organization,Cellular Development,Cellular Function and Maintenance,Cellular Growth and Proliferation,Embryonic Development,Nervous System Development and Function,Organismal Development,Tissue Development** | Dendritic growth/branching | 0.022 | ABI1, AGRN, BSN, DAB2IP, ELK1, EPHB3, GAP43, MYO5B, NRN1, PTGS2, ROCK2, RYR2, SHANK2, SIPA1L1, SORBS2, TIAM1, TSC2 | 17 |
| **Neurological Disease,Organismal Injury and Abnormalities** | Abnormality of cerebral cortex | 0.0227 | ADCY1, ASPM, FES, GAP43, GRIA2, HIVEP2, KIF1B, LRRK2, LRRTM4, MAN2C1, MCPH1, PRKN, PTGS2, SLC2A8, SMPD1, SNAP25, ZIC5 | 17 |
| **Cell Morphology,Cellular Compromise** | Collapse of growth cone | 0.0251 | ARHGEF11, EPHB3, FES, ROCK2, SEMA4F, SORCS2 | 6 |
| **Cell Morphology,Nervous System Development and Function,Neurological Disease,Tissue Morphology** | Abnormal morphology of bipolar cells | 0.0252 | ERC2, FAT3, SEMA5B | 3 |
| **Cell Morphology,Cell-To-Cell Signaling and Interaction,Cellular Assembly and Organization,Nervous System Development and Function,Neurological Disease,Tissue Morphology** | Abnormal morphology of neuromuscular synapse | 0.0253 | AGRN, COL13A1, SYNE1, SYNE2 | 4 |
| **Cell-To-Cell Signaling and Interaction,Nervous System Development and Function** | Long term synaptic depression of brain cells | 0.0253 | ADCY5, GRIA2, PIP5K1C, RAB11FIP5 | 4 |
| **Cell Morphology,Cellular Assembly and Organization,Cellular Development,Cellular** | Formation of dendrites | 0.0254 | ABI1, ADGRB3, | 11 |

| | | | | |
|---|---|---|---|---|
| **Function and Maintenance,Cellular Growth and Proliferation,Nervous System Development and Function,Organismal Development,Tissue Development** | | | ANAPC2, CELSR2, DVL1, FMN1, MEF2A, MYO5B, ROCK2, SIPA1L1, SYNE1 | |
| **Behavior,Nervous System Development and Function** | Memory | 0.0275 | ADCY1, CREBBP, KCNH3, NEURL1, NR3C2 | 5 |
| **Embryonic Development,Organismal Development** | Development of head | 0.0277 | ABR, ADCY1, ADGRA2, AHI1, ARNT2, ASPM, CDK5RAP1, CELSR2, ELP1, EPHB3, FAT3, GAP43, GRHL2, LRRK2, MCPH1, MED12, MRTFA, NCOR1, NIN, NOS2, PITPNM1, PROKR2, ROGDI, RTP1, SCRIB, SETD2, SOX4, TACC2, TOP2B, VAX1, ZIC5 | 31 |
| **Embryonic Development,Nervous System Development and Function,Organ Development,Organismal Development,Tissue Development** | Formation of forebrain | 0.0297 | ADGRA2, ARNT2, ELP1, EPHB3, GAP43, LRRK2, MRTFA, NCOR1, NIN, PROKR2, RTP1, SETD2, TOP2B, VAX1, ZIC5 | 15 |
| **Cell Morphology,Nervous System Development and Function,Neurological Disease,Tissue Morphology** | Abnormal morphology of neurons | 0.0299 | AGRN, CADPS2, COL13A1, DVL1, EPHB3, ERC2, FAT3, GAP43, HIVEP2, IDUA, LRRK2, NOS2, NRN1, PRKN, ROCK2, SCN1B, SEMA5B, SHANK2, | 28 |

| | | | | |
|---|---|---|---|---|
| | | | SMPD1, SORBS2, SORL1, SOX4, SPTBN4, SYNE1, SYNE2, UNC13A, UNC13B, VPS13A | |
| **Nervous System Development and Function,Organ Morphology,Organismal Development** | Morphology of forebrain | 0.0314 | ADCY1, ADGRA2, ARNT2, ASPM, CELSR2, CREBBP, FES, GAP43, HIVEP2, MAN2C1, NCOR1, NOS2, PROKR2, TOP2B, VAX1, ZIC5 | 16 |
| **Cell-To-Cell Signaling and Interaction,Nervous System Development and Function** | Synaptic depression of collateral synapses | 0.0321 | ADCY1, RAB11FIP5 | 2 |
| **Cell-To-Cell Signaling and Interaction,Nervous System Development and Function** | Long-term potentiation of mossy fibers | 0.0321 | ADCY1, GRIA2 | 2 |
| **Amino Acid Metabolism,Drug Metabolism,Endocrine System Development and Function,Molecular Transport,Small Molecule Biochemistry** | Concentration of levothyroxine | 0.0321 | NCOR1, SLCO1C1 | 2 |
| **Nervous System Development and Function,Neurological Disease,Organ Morphology,Organismal Development,Organismal Injury and Abnormalities** | Abnormal morphology of anterior horn of the spinal cord | 0.0321 | KIF1B, RIMS2 | 2 |
| **Nervous System Development and Function,Neurological Disease,Organ Morphology,Organismal Development,Organismal Injury and Abnormalities** | Abnormal morphology of optic tract | 0.0321 | EPHB3, GAP43 | 2 |
| **Carbohydrate Metabolism,Drug Metabolism,Molecular Transport,Small Molecule Biochemistry** | Quantity of heparan sulfate | 0.0322 | IDUA | 1 |
| **Cellular Development,Nervous System Development and Function,Tissue Development** | Differentiation of sympathetic preganglionic neurons | 0.0322 | AGRN | 1 |
| **Cellular Assembly and Organization** | Aggregation of early endosomes | 0.0322 | KIF16B | 1 |
| **Cell Morphology,Cellular Assembly and Organization,Cellular Function and Maintenance** | Formation of axon-like process | 0.0322 | TIAM1 | 1 |
| **Cell Morphology,Cell-To-Cell Signaling and Interaction** | Length of stereocilia | 0.0322 | MYO15A | 1 |
| **Cell-To-Cell Signaling and Interaction,Cellular Assembly and Organization,Nervous System Development and Function,Tissue Morphology** | Quantity of ribbon synapse | 0.0322 | BSN | 1 |
| **Nervous System Development and Function,Tissue Morphology** | Density of serotonergic neurons | 0.0322 | GAP43 | 1 |
| **Behavior,Developmental Disorder,Neurological Disease,Psychological Disorders** | Autistic-like traits | 0.0322 | TSC2 | 1 |
| **Nervous System Development and Function** | Function of autonomic nerve | 0.0322 | SNAP25 | 1 |
| **Hereditary Disorder,Neurological Disease,Organismal Injury and Abnormalities,Psychological Disorders,Skeletal and Muscular Disorders** | Autosomal recessive juvenile Parkinson disease | 0.0322 | PRKN | 1 |
| **Cellular Compromise** | Acidification of cytosol | 0.0322 | CA4 | 1 |
| **Cell-To-Cell Signaling and Interaction,Nervous System Development and Function** | Inactivation of synapse | 0.0322 | BSN | 1 |
| **Cellular Development,Nervous System Development and Function,Organ Development** | Differentiation of preoptic area | 0.0322 | VAX1 | 1 |

| | | | | |
|---|---|---|---|---|
| **Cellular Development,Nervous System Development and Function,Organ Development** | Differentiation of medial ganglionic eminences | 0.0322 | VAX1 | 1 |
| **Cellular Development,Cellular Growth and Proliferation,Embryonic Development,Nervous System Development and Function,Organ Development,Organismal Development,Tissue Development,Visual System Development and Function** | Formation of retinal cells | 0.0322 | ADCY1 | 1 |
| **Nervous System Development and Function,Organ Morphology,Tissue Morphology** | Quantity of rod bipolar cells | 0.0322 | PRDM1 | 1 |
| **Lipid Metabolism,Small Molecule Biochemistry** | Storage of sphingomyelin | 0.0322 | SMPD1 | 1 |
| **Cell Morphology,Cellular Assembly and Organization,Cellular Development,Cellular Function and Maintenance,Cellular Growth and Proliferation,Embryonic Development,Nervous System Development and Function,Organismal Development,Tissue Development** | Delay in initiation of outgrowth of dendrites | 0.0322 | DAB2IP | 1 |
| **Cell-To-Cell Signaling and Interaction,Nervous System Development and Function** | Clustering of hippocampal neurons | 0.0322 | SHANK2 | 1 |
| **Molecular Transport,Protein Trafficking** | Translocation of protein fragment | 0.0322 | AGAP3 | 1 |
| **Cell-To-Cell Signaling and Interaction,Nervous System Development and Function** | Activation of medium spiny neurons | 0.0322 | LRRK2 | 1 |
| **Hematological System Development and Function,Lymphoid Tissue Structure and Development,Tissue Morphology** | Quantity of CD8+ T lymphocyte | 0.0322 | SMPD1 | 1 |
| **Nervous System Development and Function,Tissue Morphology** | Density of parallel fiber | 0.0322 | DAB2IP | 1 |
| **Cell Death and Survival,Cellular Development,Nervous System Development and Function,Tissue Development,Tissue Morphology** | Regeneration of bipolar cells | 0.0322 | PRDM1 | 1 |
| **Nervous System Development and Function,Organ Morphology,Organismal Development** | Size of optic chiasm | 0.0322 | GAP43 | 1 |
| **Cell Morphology,Cellular Assembly and Organization,Cellular Development,Cellular Function and Maintenance,Cellular Growth and Proliferation,Nervous System Development and Function,Organismal Development,Tissue Development** | Sprouting of rod bipolar cells | 0.0322 | ERC2 | 1 |
| **Cellular Development,Nervous System Development and Function,Tissue Development** | Differentiation of pyramidal neurons | 0.0322 | UNC5D | 1 |
| **Inflammatory Response** | Inflammatory response of cerebral cortex | 0.0322 | KNG1 | 1 |
| **Cellular Development** | Delay in initiation of transition of radial glial cells | 0.0322 | PLAGL1 | 1 |
| **Cellular Development** | Delay in initiation of transition of neuronal progenitor cells | 0.0322 | PLAGL1 | 1 |
| **Hereditary Disorder,Neurological Disease,Organismal Injury and Abnormalities,Psychological Disorders,Skeletal and Muscular Disorders** | Huntington disease grade 1 Huntington disease | 0.0322 | SNAP25 | 1 |
| **Embryonic Development,Nervous System Development and Function,Organ Development,Organismal Development,Tissue Development** | Development of septal area | 0.0322 | VAX1 | 1 |
| **Behavior** | Social exploration | 0.0322 | APBA2 | 1 |
| **Hereditary Disorder,Neurological Disease,Organismal Injury and Abnormalities,Psychological Disorders,Skeletal and Muscular Disorders** | Parkinson-dementia syndrome | 0.0322 | GAP43 | 1 |
| **Cardiovascular System Development and Function,Organismal Development** | Penetration by blood vessel | 0.0322 | ADGRA2 | 1 |
| **Developmental Disorder,Endocrine System Disorders,Nutritional Disease,Organismal Injury and Abnormalities,Reproductive System Disease** | Delayed puberty | 0.0322 | VAX1 | 1 |

| | | | | |
|---|---|---|---|---|
| **Cell-To-Cell Signaling and Interaction,Nervous System Development and Function** | Sensitization of trigeminovascular neuron | 0.0322 | KNG1 | 1 |
| **Cellular Assembly and Organization,Cellular Compromise,Nervous System Development and Function** | Retraction of glial cell projections | 0.0322 | GRIA2 | 1 |
| **Behavior** | Perseverance behavior | 0.0322 | NR3C2 | 1 |
| **Nervous System Development and Function** | Formation of neuronal layer | 0.0322 | SOX4 | 1 |
| **Cellular Development,Cellular Growth and Proliferation,Nervous System Development and Function,Tissue Development** | Development of corticospinal neurons | 0.0322 | BCL11B | 1 |
| **Behavior** | Impulsivity | 0.0322 | ADCY1 | 1 |
| **Metabolic Disease,Neurological Disease,Organismal Injury and Abnormalities,Psychological Disorders** | Preclinical stage Alzheimer disease | 0.0322 | NOS2 | 1 |
| **Hematological Disease** | Hyperemia | 0.0322 | PTGS2 | 1 |
| **Nervous System Development and Function,Tissue Morphology** | Density of climbing fiber | 0.0322 | DAB2IP | 1 |
| **Embryonic Development,Nervous System Development and Function,Organ Development,Organismal Development,Tissue Development** | Development of layer IV | 0.0322 | GAP43 | 1 |
| **Cellular Assembly and Organization** | Delay in initiation of priming of vesicles | 0.0322 | SNAP25 | 1 |
| **Cell-To-Cell Signaling and Interaction,Drug Metabolism,Small Molecule Biochemistry** | Conversion of norepinephrine | 0.0322 | DBH | 1 |
| **Infectious Diseases** | Replication of Murine cytomegalovirus | 0.0322 | NOS2 | 1 |
| **Metabolic Disease,Neurological Disease,Organismal Injury and Abnormalities,Psychological Disorders** | Stage V Alzheimer disease | 0.0322 | PTGS2 | 1 |
| **Cell Morphology,Cellular Assembly and Organization,Nervous System Development and Function** | Projection of mossy fibers | 0.0322 | NR3C2 | 1 |
| **Cell-To-Cell Signaling and Interaction,Nervous System Development and Function** | Activation of mitral cells | 0.0322 | GJD2 | 1 |
| **Cell Morphology,Cellular Assembly and Organization** | Orientation of plasma membrane projections | 0.0322 | SCN1B | 1 |
| **Cellular Assembly and Organization,Cellular Development,Nervous System Development and Function,Tissue Development** | Differentiation of presynaptic terminals | 0.0322 | AGRN | 1 |
| **Cell-To-Cell Signaling and Interaction,Drug Metabolism,Small Molecule Biochemistry** | Catabolism of dopamine | 0.0322 | PRKN | 1 |
| **Nervous System Development and Function,Neurological Disease,Organ Morphology,Organismal Development,Organismal Injury and Abnormalities** | Lack of optic tract | 0.0322 | GAP43 | 1 |
| **Cell Morphology,Cellular Assembly and Organization,Nervous System Development and Function,Tissue Morphology** | Diameter of nodes of Ranvier | 0.0322 | SPTBN4 | 1 |
| **Cellular Development,Nervous System Development and Function,Tissue Development** | Differentiation of postsynaptic neurons | 0.0322 | AGRN | 1 |
| **Nervous System Development and Function** | Discharge of cochlear nerve | 0.0322 | BSN | 1 |
| **Cardiovascular System Development and Function,Nervous System Development and Function,Organ Morphology,Organismal Development,Tissue Morphology** | Diameter of basilar artery | 0.0322 | KNG1 | 1 |
| **Cell Morphology,Cellular Movement,Nervous System Development and Function** | Innervation of serotonergic neurons | 0.0322 | GAP43 | 1 |
| **Cell-To-Cell Signaling and Interaction,Nervous System Development and Function** | Firing of trigeminovascular neuron | 0.0322 | KNG1 | 1 |
| **Cell Morphology,Cellular Assembly and Organization** | Elongation of astrocyte projections | 0.0322 | GAP43 | 1 |
| **Nervous System Development and Function,Neurological Disease,Organ Morphology,Organismal Development,Organismal Injury and Abnormalities** | Abnormal morphology of cerebellar hemisphere | 0.0322 | AHI1 | 1 |

| | | | | |
|---|---|---|---|---|
| **Lipid Metabolism,Molecular Transport,Small Molecule Biochemistry** | Concentration of eicosapentenoic acid | 0.0322 | ARNTL | 1 |
| **Infectious Diseases** | Infection by measles virus | 0.0322 | SMPD1 | 1 |
| **Cell-To-Cell Signaling and Interaction,Cellular Function and Maintenance,Nervous System Development and Function** | Paired-pulse facilitation of parallel fiber-Purkinje cell synapses | 0.0322 | UNC13C | 1 |
| **Cancer,Neurological Disease,Organismal Injury and Abnormalities,Tumor Morphology** | Progression of glioma | 0.0322 | MIIP | 1 |
| **Metabolic Disease,Neurological Disease,Organismal Injury and Abnormalities,Psychological Disorders** | Stage I Alzheimer disease | 0.0322 | PTGS2 | 1 |
| **Cellular Compromise,Neurological Disease,Organismal Injury and Abnormalities** | Neurodegeneration of hippocampal cells | 0.0325 | GRIA2, PRKN, PTGS2 | 3 |
| **Cell-To-Cell Signaling and Interaction,Drug Metabolism,Molecular Transport,Small Molecule Biochemistry** | Concentration of norepinephrine | 0.0325 | DBH, GAP43, PRKN | 3 |
| **Nervous System Development and Function,Neurological Disease,Ophthalmic Disease** | Abnormal morphology of optic nerve | 0.0365 | SCN1B, SPTBN4, VAX1 | 3 |
| **Behavior** | Cognition | 0.0371 | ADCY1, ADGRB3, CREBBP, KCNH3, NEURL1, NR3C2, SMPD1 | 7 |
| **Cell-To-Cell Signaling and Interaction,Nervous System Development and Function** | Activation of synapse | 0.0393 | APBA2, GRIA2 | 2 |
| **Nervous System Development and Function,Tissue Morphology** | Density of axons | 0.0393 | ACVR1C, DAB2IP | 2 |
| **Embryonic Development,Nervous System Development and Function,Organ Development,Organismal Development,Tissue Development,Visual System Development and Function** | Formation of eye | 0.0393 | ADCY1, FAT3 | 2 |
| **Cell-To-Cell Signaling and Interaction,Cellular Assembly and Organization,Cellular Development,Cellular Function and Maintenance,Cellular Growth and Proliferation,Nervous System Development and Function,Tissue Development** | Formation of excitatory synapses | 0.0393 | LRRTM4, SORBS2 | 2 |
| **Cellular Compromise,Neurological Disease,Organismal Injury and Abnormalities** | Neurodegeneration of brain cells | 0.0398 | GRIA2, IDUA, LRRK2, PRKN, PTGS2, SMPD1 | 6 |
| **Cellular Assembly and Organization,Nervous System Development and Function** | Recycling of synaptic vesicles | 0.0408 | DVL1, GAP43, PIP5K1C | 3 |
| **Embryonic Development,Nervous System Development and Function,Organ Development,Organismal Development,Tissue Development** | Formation of brain | 0.041 | ABR, ADGRA2, AHI1, ARNT2, ASPM, CDK5RAP1, CELSR2, ELP1, EPHB3, GAP43, GRHL2, LRRK2, MCPH1, MED12, MRTFA, NCOR1, NIN, NOS2, PITPNM1, PROKR2, ROGDI, RTP1, | 29 |

| | | | | |
|---|---|---|---|---|
| | | | SCRIB, SETD2, SOX4, TACC2, TOP2B, VAX1, ZIC5 | |
| **Embryonic Development,Nervous System Development and Function,Organ Development,Organismal Development,Tissue Development** | Formation of telencephalon | 0.0411 | ADGRA2, ELP1, EPHB3, GAP43, LRRK2, NIN, PROKR2, RTP1, TOP2B, VAX1 | 10 |
| **Nervous System Development and Function,Tissue Morphology** | Morphology of nervous tissue | 0.0414 | AGRN, CADPS2, COL13A1, DVL1, EPHB3, ERC2, FAT3, GAP43, HIVEP2, IDUA, LRRK2, MAN2C1, NOS2, NRN1, PLAGL1, PRKN, RGS11, RIMS2, ROCK2, SCN1B, SEMA5B, SHANK2, SIPA1L1, SMPD1, SORBS2, SORL1, SOX4, SPTBN4, SYNE1, SYNE2, TIAM1, UNC13A, UNC13B, VPS13A | 34 |
| **Neurological Disease,Organismal Injury and Abnormalities** | Neurodegeneration of cerebral cortex | 0.0464 | GRIA2, LRRK2, PRKN, PTGS2, SNAP25 | 5 |
| **Cell Death and Survival,Cellular Compromise,Neurological Disease,Organismal Injury and Abnormalities,Tissue Morphology** | Neurodegeneration of pyramidal neurons | 0.0471 | GRIA2, PTGS2 | 2 |
| **Neurological Disease,Organismal Injury and Abnormalities,Psychological Disorders,Skeletal and Muscular Disorders** | Parkinson disease with dementia | 0.0471 | GAP43, NOS2 | 2 |
| **Cellular Movement,Nervous System Development and Function** | Guidance of axons | 0.0473 | DVL1, EPHB3, GAP43, KIF5C, LMTK2, PIK3R1, SCN1B, SEMA4F, SPTB, SPTBN4, UNC5D, VAX1 | 12 |
| **Cell Morphology,Nervous System Development and Function,Tissue Morphology** | Morphology of neurons | 0.0474 | AGRN, CADPS2, | 33 |

| | | | | |
|---|---|---|---|---|
| | | | COL13A1, DVL1, EPHB3, ERC2, FAT3, GAP43, HIVEP2, IDUA, LRRK2, NOS2, NRN1, PLAGL1, PRKN, RGS11, RIMS2, ROCK2, SCN1B, SEMA5B, SHANK2, SIPA1L1, SMPD1, SORBS2, SORL1, SOX4, SPTBN4, SYNE1, SYNE2, TIAM1, UNC13A, UNC13B, VPS13A | |
| **Nervous System Development and Function,Neurological Disease,Organ Morphology,Organismal Development,Organismal Injury and Abnormalities** | Abnormal morphology of telencephalon | 0.0497 | ADGRA2, ASPM, CELSR2, FES, HIVEP2, MAN2C1, NOS2, PROKR2, TOP2B, VAX1, ZIC5 | 11 |

### Cluster 11

| Categories | Diseases or Functions Annotation | p-Value | Molecules | # Molecules |
|---|---|---|---|---|
| **Cellular Function and Maintenance,Molecular Transport** | Exocytosis | 5.35E-08 | APBA2, CADPS2, ERC2, PIP5K1C, PPFIA3, RIMS2, RPH3A, SNAP25, UNC13A, UNC13B | 10 |
| **Cellular Assembly and Organization,Cellular Function and Maintenance,Cellular Movement,Molecular Transport,Nervous System Development and Function** | Exocytosis of synaptic vesicles | 0.00000345 | APBA2, PIP5K1C, PPFIA3, RPH3A, SNAP25, UNC13A, UNC13B | 7 |
| **Cellular Assembly and Organization** | Priming of vesicles | 0.00000588 | CADPS2, ERC2, RIMS2, SNAP25, UNC13A, UNC13B | 6 |
| **Cellular Assembly and Organization,Nervous System Development and Function** | Docking of synaptic vesicles | 0.00000657 | PPFIA3, RIMS2, SNAP25, UNC13A, UNC13B | 5 |

| | | | | |
|---|---|---|---|---|
| **Cell-To-Cell Signaling and Interaction,Cellular Assembly and Organization,Cellular Function and Maintenance,Molecular Transport** | Priming of synaptic vesicles | 0.0000459 | CADPS2, ERC2, RIMS2, UNC13A, UNC13B | 5 |
| **Cellular Assembly and Organization,Cellular Movement,Nervous System Development and Function** | Transport of synaptic vesicles | 0.0000464 | AP2M1, APBA2, KIAA1107, PIP5K1C, PPFIA3, RPH3A, SNAP25, UNC13A, UNC13B | 9 |
| **Nervous System Development and Function,Neurological Disease** | Abnormal morphology of nervous system | 0.000165 | ABR, ADCY1, AGRN, ARNT2, ASPM, CACNA1C, CADPS2, CELSR2, COL13A1, DVL1, ECE1, EPHB3, ERC2, FAT3, FES, GAP43, GRIA2, HIVEP2, LRRK2, LRRTM4, MEGF8, NCOR1, NOS2, NRN1, PER2, PRKN, RIMS2, ROCK2, SCN1B, SEMA5B, SNAP25, SORL1, SPTBN4, TCF7L2, TENM2, UNC13A, UNC13B, VAX1, VPS13A | 39 |
| **Cancer,Organismal Injury and Abnormalities** | Head and neck tumor | 0.000329 | ABCF1, ADAMTS8, ANKRD30B, ASPM, ASXL3, ATF7IP2, CAMTA1, CUL9, DPP10, ERC2, HERC1, MYO15A, MYO5B, P2RX5, PIK3R1, PRDM1, PRDM2, PTGS2, PTPRT, RAB11FIP5, RALY, RIMS2, ROS1, | 28 |

| | | | | |
|---|---|---|---|---|
| | | | RYR2, SEMA4F, SPTBN4, THEMIS, UNC13B | |
| **Cancer,Neurological Disease,Organismal Injury and Abnormalities** | Astrocytoma | 0.000436 | ABCF1, ADAMTS8, ANKRD30B, ASPM, ASXL3, ATF7IP2, CAMTA1, DPP10, ERC2, HERC1, MYO5B, P2RX5, PIK3R1, PRDM2, PTGS2, RAB11FIP5, RALY, RIMS2, ROS1, RYR2, SEMA4F, SPTBN4, THEMIS | 23 |
| **Cell-To-Cell Signaling and Interaction,Nervous System Development and Function** | Action potential of motor endplates | 0.000467 | COL13A1, RIMS2, SNAP25 | 3 |
| **Nervous System Development and Function** | Morphology of nervous system | 0.000535 | ABR, ADCY1, AGRN, ARNT2, ASPM, CACNA1C, CADPS2, CELSR2, COL13A1, DVL1, ECE1, EPHB3, ERC2, FAT3, FES, GAP43, GRIA2, HIVEP2, KIF5C, KNG1, LRRK2, LRRTM4, MEGF8, NCOR1, NOS2, NRN1, PER2, PLAGL1, PRKN, RIMS2, ROCK2, SCN1B, SEMA5B, SIPA1L1, SNAP25, SORL1, SPTBN4, TCF7L2, TENM2, UNC13A, UNC13B, VAX1, VPS13A | 43 |
| **Cancer,Organismal Injury and Abnormalities** | Non-melanoma solid tumor | 0.000599 | ABCF1, ADAMTS8, | 27 |

| | | | | |
|---|---|---|---|---|
| | | | ANKRD30B, ASPM, ASXL3, ATF7IP2, CAMTA1, DPP10, ERC2, HERC1, MYO15A, MYO5B, P2RX5, PIK3R1, PRDM1, PRDM2, PTGS2, PTPRT, RAB11FIP5, RALY, RIMS2, ROS1, RYR2, SEMA4F, SPTBN4, THEMIS, UNC13B | |
| **Cancer,Neurological Disease,Organismal Injury and Abnormalities** | Brain tumor | 0.000613 | ABCF1, ADAMTS8, ANKRD30B, ASPM, ASXL3, ATF7IP2, CAMTA1, DPP10, ERC2, HERC1, MYO15A, MYO5B, P2RX5, PIK3R1, PRDM1, PRDM2, PTGS2, PTPRT, RAB11FIP5, RALY, RIMS2, ROS1, RYR2, SEMA4F, SPTBN4, THEMIS, UNC13B | 27 |
| **Cancer,Organismal Injury and Abnormalities** | Non-hematological solid tumor | 0.000633 | ABCF1, ADAMTS8, ANKRD30B, ASPM, ASXL3, ATF7IP2, CAMTA1, CUL9, DPP10, ERC2, HERC1, MYO15A, MYO5B, P2RX5, PIK3R1, PRDM1, PRDM2, PTGS2, PTPRT, RAB11FIP5, RALY, RIMS2, ROS1, | 28 |

| | | | | |
|---|---|---|---|---|
| | | | RYR2, SEMA4F, SPTBN4, THEMIS, UNC13B | |
| **Cancer,Neurological Disease,Organismal Injury and Abnormalities** | Brain glioma | 0.000648 | ABCF1, ADAMTS8, ANKRD30B, ASPM, ASXL3, ATF7IP2, CAMTA1, DPP10, ERC2, HERC1, MYO15A, MYO5B, P2RX5, PIK3R1, PRDM1, PRDM2, PTGS2, PTPRT, RAB11FIP5, RALY, RIMS2, ROS1, RYR2, SEMA4F, SPTBN4, THEMIS | 26 |
| **Lipid Metabolism,Molecular Transport,Small Molecule Biochemistry** | Concentration of arachidonic acid | 0.00069 | ARNTL, KNG1, PTGS2 | 3 |
| **Cell Morphology,Cell-To-Cell Signaling and Interaction,Nervous System Development and Function** | Plasticity of synapse | 0.000747 | CAMK2G, GRIA2, KCNH3, NEURL1, PCDH8, PIP5K1C, PPFIA3, PTK2B, RASGRF1, SIPA1L1 | 10 |
| **Cancer,Neurological Disease,Organismal Injury and Abnormalities** | Brain astrocytoma | 0.000827 | ABCF1, ADAMTS8, ANKRD30B, ASPM, ASXL3, ATF7IP2, DPP10, ERC2, HERC1, MYO5B, P2RX5, PIK3R1, PRDM2, PTGS2, RAB11FIP5, RALY, RIMS2, ROS1, RYR2, SEMA4F, SPTBN4, THEMIS | 22 |
| **Cellular Assembly and Organization,Cellular Compromise** | Retraction of cellular protrusions | 0.00106 | ARHGEF11, DVL1, GRIA2, LRRK2, PITPNM1, ROCK2 | 6 |
| **Cell Morphology,Cell-To-Cell Signaling and Interaction,Cellular Assembly and** | Morphology of synapse | 0.00108 | AGRN, COL13A1, ERC2, | 8 |

| | | | | |
|---|---|---|---|---|
| **Organization,Nervous System Development and Function,Tissue Morphology** | | | GAP43, NRN1, ROCK2, UNC13A, UNC13B | |
| **Hereditary Disorder,Neurological Disease,Organismal Injury and Abnormalities,Psychological Disorders,Skeletal and Muscular Disorders** | Familial Parkinson disease | 0.00129 | GAP43, PRKN | 2 |
| **Hereditary Disorder,Neurological Disease,Organismal Injury and Abnormalities,Psychological Disorders,Skeletal and Muscular Disorders** | Huntington disease grade 4 Huntington disease | 0.00129 | RPH3A, SNAP25 | 2 |
| **Neurological Disease,Organismal Injury and Abnormalities** | Brain lesion | 0.00131 | ABCF1, ADAMTS8, ANKRD30B, ASPM, ASXL3, ATF7IP2, CAMTA1, DPP10, ERC2, HERC1, KNG1, MYO15A, MYO5B, P2RX5, PIK3R1, PRDM1, PRDM2, PTGS2, PTPRT, RAB11FIP5, RALY, RIMS2, ROS1, RYR2, SEMA4F, SPTBN4, THEMIS, UNC13B | 28 |
| **Cell-To-Cell Signaling and Interaction,Cellular Assembly and Organization** | Cell-cell contact | 0.00196 | AGRN, BSN, COL13A1, EPHB3, ERC2, GAP43, GJD2, LRRTM4, NEURL1, NRN1, PCDH8, PLXND1, ROCK2, UNC13A, UNC13B | 15 |
| **Cellular Movement,Nervous System Development and Function** | Guidance of axons | 0.00196 | DVL1, EPHB3, GAP43, KIF5C, LMTK2, PIK3R1, RANBP9, SCN1B, SEMA4F, SPTBN4, UNC5D, VAX1 | 12 |
| **Nervous System Development and Function,Neurological Disease,Organ Morphology,Organismal Development,Organismal Injury and Abnormalities** | Abnormal morphology of barrel cortex | 0.00221 | ADCY1, ASPM, GAP43 | 3 |
| **Nervous System Development and Function,Neurological Disease,Ophthalmic** | Abnormal morphology of visual cortex | 0.00254 | HIVEP2, TENM2 | 2 |

| | | | | |
|---|---|---|---|---|
| **Disease,Organ Morphology,Organismal Development,Organismal Injury and Abnormalities** | | | | |
| **Behavior,Nervous System Development and Function** | Working memory | 0.00254 | KCNH3, NR3C2 | 2 |
| **Cell Morphology,Cell-To-Cell Signaling and Interaction,Cellular Assembly and Organization,Nervous System Development and Function,Neurological Disease,Tissue Morphology** | Abnormal morphology of synapse | 0.00257 | AGRN, COL13A1, ERC2, NRN1, ROCK2, UNC13A, UNC13B | 7 |
| **Tissue Morphology** | Quantity of cells | 0.00309 | ACVR1C, AGRN, BSN, CACNA1C, CADPS2, DVL1, EPHB3, ERC2, HIVEP2, KIF5C, LRRK2, MGAT5B, NEURL1, NOS2, NR3C2, NRN1, PRDM1, PRKN, PTGS2, RIMS2, TCF7L2, TRPM2, UNC13A, UNC13B, VAX1 | 25 |
| **Cell Morphology,Nervous System Development and Function,Neurological Disease,Tissue Morphology** | Abnormal morphology of neurons | 0.00346 | AGRN, CACNA1C, CADPS2, COL13A1, DVL1, ECE1, EPHB3, ERC2, FAT3, GAP43, HIVEP2, LRRK2, NOS2, NRN1, PRKN, ROCK2, SCN1B, SEMA5B, SORL1, SPTBN4, UNC13A, UNC13B, VPS13A | 23 |
| **Cellular Assembly and Organization,Cellular Compromise,Nervous System Development and Function** | Retraction of neurites | 0.00377 | ARHGEF11, DVL1, LRRK2, PITPNM1, ROCK2 | 5 |
| **Cell-To-Cell Signaling and Interaction,Nervous System Development and Function** | Long term synaptic depression of CA1 neuron | 0.00413 | GRIA2, PIP5K1C, RAB11FIP5 | 3 |
| **Cellular Development,Nervous System Development and Function,Tissue Development** | Differentiation of nerve ending | 0.00417 | AGRN, MEF2A | 2 |
| **Hereditary Disorder,Neurological Disease,Organismal Injury and Abnormalities,Psychological Disorders,Skeletal and Muscular Disorders** | Huntington disease grade 3 Huntington disease | 0.00417 | RPH3A, SNAP25 | 2 |

| | | | | |
|---|---|---|---|---|
| **Cell Morphology,Nervous System Development and Function** | Innervation of optic nerve | 0.00417 | GAP43, VAX1 | 2 |
| **Cellular Compromise,Neurological Disease,Organismal Injury and Abnormalities** | Neurodegeneration of cerebral cortex cells | 0.00423 | GRIA2, LRRK2, PRKN, PTGS2 | 4 |
| **Cellular Development,Cellular Growth and Proliferation,Nervous System Development and Function,Tissue Development** | Development of neurons | 0.00472 | ADCY1, ADGRB3, AGRN, ANAPC2, ATCAY, BSN, CADPS2, CELSR2, DVL1, ECE1, EPHB3, ERC2, FMN1, GAP43, HERC1, LRRK2, LRRTM4, MEF2A, MRTFA, MYO5B, NRN1, PCDH8, PIP5K1C, PLXND1, PRDM1, PTGS2, PTK2B, RASGRF1, ROCK2, RYR2, SIPA1L1, SPTBN4 | 32 |
| **Lipid Metabolism,Small Molecule Biochemistry** | Synthesis of eicosanoid | 0.00583 | KNG1, NOS2, PTGS2 | 3 |
| **Cell-To-Cell Signaling and Interaction,Nervous System Development and Function** | Long term synaptic depression of brain cells | 0.00586 | ADCY5, GRIA2, PIP5K1C, RAB11FIP5 | 4 |
| **Nervous System Development and Function** | Depotentiation | 0.00617 | ADCY1, GRIA2 | 2 |
| **Cell Morphology,Nervous System Development and Function,Neurological Disease,Tissue Morphology** | Abnormal morphology of bipolar cells | 0.0079 | ERC2, FAT3, SEMA5B | 3 |
| **Neurological Disease,Organismal Injury and Abnormalities** | Hydrocephalus | 0.0079 | CELSR2, DVL1, NCOR1 | 3 |
| **Cell Morphology,Nervous System Development and Function,Tissue Morphology** | Morphology of neurons | 0.00859 | AGRN, CACNA1C, CADPS2, COL13A1, DVL1, ECE1, EPHB3, ERC2, FAT3, GAP43, HIVEP2, LRRK2, NOS2, NRN1, PLAGL1, PRKN, RIMS2, ROCK2, SCN1B, SEMA5B, SIPA1L1, SORL1, | 26 |

| | | | | |
|---|---|---|---|---|
| | | | SPTBN4, UNC13A, UNC13B, VPS13A | |
| **Cell-To-Cell Signaling and Interaction,Nervous System Development and Function** | Synaptic transmission of cells | 0.00868 | BSN, ERC2, GRIA2, NR3C2, RAB11FIP5, SNAP25 | 6 |
| **Neurological Disease,Organismal Injury and Abnormalities** | Neurodegeneration of cerebral cortex | 0.0088 | GRIA2, LRRK2, PRKN, PTGS2, SNAP25 | 5 |
| **Neurological Disease,Psychological Disorders** | Pervasive developmental disorder | 0.00908 | MATK, RYR2, SLC6A7 | 3 |
| **Cellular Compromise,Neurological Disease,Organismal Injury and Abnormalities** | Neurodegeneration of hippocampal cells | 0.0104 | GRIA2, PRKN, PTGS2 | 3 |
| **Cell-To-Cell Signaling and Interaction,Drug Metabolism,Molecular Transport,Small Molecule Biochemistry** | Concentration of norepinephrine | 0.0104 | DBH, GAP43, PRKN | 3 |
| **Cell Morphology,Cellular Assembly and Organization,Cellular Development,Cellular Function and Maintenance,Cellular Growth and Proliferation,Nervous System Development and Function,Organismal Development,Tissue Development** | Formation of dendrites | 0.0109 | ADGRB3, ANAPC2, CELSR2, DVL1, FMN1, MEF2A, MYO5B, ROCK2, SIPA1L1 | 9 |
| **Nervous System Development and Function,Tissue Morphology** | Density of neurons | 0.0116 | ACVR1C, EPHB3, GAP43, LRRTM4, MEF2A, PTGS2, RYR2, SCN1B | 8 |
| **Nervous System Development and Function,Neurological Disease,Ophthalmic Disease** | Abnormal morphology of optic nerve | 0.0117 | SCN1B, SPTBN4, VAX1 | 3 |
| **Nervous System Development and Function,Tissue Morphology** | Quantity of neurons | 0.0118 | ACVR1C, AGRN, BSN, CACNA1C, CADPS2, DVL1, EPHB3, ERC2, KIF5C, NEURL1, NRN1, PRDM1, PRKN, RIMS2, TRPM2, UNC13A, UNC13B, VAX1 | 18 |
| **Nervous System Development and Function,Neurological Disease** | Abnormal morphology of central nervous system | 0.0121 | ABR, ADCY1, ARNT2, ASPM, CADPS2, CELSR2, EPHB3, FES, GAP43, GRIA2, HIVEP2, LRRTM4, NCOR1, NOS2, | 22 |

| | | | | |
|---|---|---|---|---|
| | | | PRKN, RIMS2, SCN1B, SNAP25, SPTBN4, TCF7L2, TENM2, VAX1 | |
| **Cell Morphology,Cellular Assembly and Organization,Cellular Development,Cellular Function and Maintenance,Cellular Growth and Proliferation,Nervous System Development and Function,Organismal Development,Tissue Development** | Neuritogenesis | 0.0125 | ADCY1, ADGRB3, AGRN, ANAPC2, ATCAY, BSN, CELSR2, DVL1, EPHB3, ERC2, FMN1, GAP43, HERC1, LRRK2, MEF2A, MRTFA, MYO5B, NRN1, PIP5K1C, PTGS2, PTK2B, RASGRF1, ROCK2, RYR2, SIPA1L1, SPTBN4 | 26 |
| **Cellular Assembly and Organization,Cellular Function and Maintenance** | Microtubule dynamics | 0.0132 | ADCY1, ADGRB3, AGRN, ANAPC2, ATCAY, BSN, CELSR2, DVL1, EPHB3, ERC2, FMN1, GAP43, HERC1, LRRK2, MEF2A, MRTFA, MYO5B, NRN1, PIP5K1C, PTGS2, PTK2B, RASGRF1, ROCK2, RYR2, SCN1B, SEMA4F, SEMA5B, SIPA1L1, SPTBN4 | 29 |
| **Cellular Assembly and Organization,Nervous System Development and Function** | Recycling of synaptic vesicles | 0.0132 | DVL1, GAP43, PIP5K1C | 3 |
| **Cell-To-Cell Signaling and Interaction,Nervous System Development and Function** | Synaptic depression of collateral synapses | 0.0142 | ADCY1, RAB11FIP5 | 2 |
| **Cell-To-Cell Signaling and Interaction,Nervous System Development and Function** | Long-term potentiation of mossy fibers | 0.0142 | ADCY1, GRIA2 | 2 |
| **Nervous System Development and Function,Neurological Disease,Organ Morphology,Organismal Development,Organismal Injury and Abnormalities** | Abnormal morphology of optic tract | 0.0142 | EPHB3, GAP43 | 2 |

| | | | | |
|---|---|---|---|---|
| **Cell-To-Cell Signaling and Interaction,Nervous System Development and Function** | Long term synaptic depression of hippocampus | 0.0153 | GRIA2, PIP5K1C, PTK2B, RAB11FIP5 | 4 |
| **Cell Morphology,Cellular Compromise** | Collapse of growth cone | 0.0154 | ARHGEF11, EPHB3, FES, ROCK2, SEMA4F | 5 |
| **Organismal Development,Organismal Injury and Abnormalities** | Abnormal morphology of head | 0.016 | ABR, ADCY1, ARNT2, ASPM, CADPS2, CELSR2, DVL1, EPHB3, FAT3, FES, GAP43, GRIA2, HIVEP2, LRRTM4, NCOR1, NOS2, PRKN, SNAP25, SPTBN4, TCF7L2, TENM2, VAX1 | 22 |
| **Nervous System Development and Function** | Activation of brain | 0.0164 | ADCY1, GJD2, LRRK2, SCN1B | 4 |
| **Cell-To-Cell Signaling and Interaction,Nervous System Development and Function** | Activation of synapse | 0.0175 | APBA2, GRIA2 | 2 |
| **Cellular Assembly and Organization,Cellular Function and Maintenance,Cellular Movement,Nervous System Development and Function** | Axonal transport of mitochondria | 0.0175 | PRKN, RANBP9 | 2 |
| **Embryonic Development,Nervous System Development and Function,Organ Development,Organismal Development,Tissue Development,Visual System Development and Function** | Formation of eye | 0.0175 | ADCY1, FAT3 | 2 |
| **Nervous System Development and Function,Neurological Disease,Organ Morphology,Organismal Development,Organismal Injury and Abnormalities** | Abnormal morphology of brain | 0.0193 | ABR, ADCY1, ARNT2, ASPM, CADPS2, CELSR2, EPHB3, FES, GAP43, GRIA2, HIVEP2, LRRTM4, NCOR1, NOS2, PRKN, SNAP25, SPTBN4, TCF7L2, TENM2, VAX1 | 20 |
| **Cell Morphology,Nervous System Development and Function** | Innervation | 0.0209 | EPHB3, GAP43, GRIA2, RIMS2, TENM2, VAX1 | 6 |
| **Cellular Development,Nervous System Development and Function,Tissue Development** | Differentiation of sympathetic preganglionic neurons | 0.0209 | AGRN | 1 |

| | | | | |
|---|---|---|---|---|
| **Cellular Assembly and Organization** | Aggregation of early endosomes | 0.0209 | KIF16B | 1 |
| **Cell-To-Cell Signaling and Interaction,Cellular Assembly and Organization,Nervous System Development and Function,Tissue Morphology** | Quantity of ribbon synapse | 0.0209 | BSN | 1 |
| **Nervous System Development and Function,Tissue Morphology** | Density of serotonergic neurons | 0.0209 | GAP43 | 1 |
| **Hereditary Disorder,Neurological Disease,Organismal Injury and Abnormalities,Psychological Disorders,Skeletal and Muscular Disorders** | Autosomal recessive juvenile Parkinson disease | 0.0209 | PRKN | 1 |
| **Nervous System Development and Function,Organ Morphology,Tissue Morphology** | Quantity of rod bipolar cells | 0.0209 | PRDM1 | 1 |
| **Cell-To-Cell Signaling and Interaction,Nervous System Development and Function** | Activation of medium spiny neurons | 0.0209 | LRRK2 | 1 |
| **Cellular Development,Cellular Growth and Proliferation,Digestive System Development and Function,Nervous System Development and Function,Tissue Development** | Development of enteric neurons | 0.0209 | ECE1 | 1 |
| **Cell Death and Survival,Cellular Development,Nervous System Development and Function,Tissue Development,Tissue Morphology** | Regeneration of bipolar cells | 0.0209 | PRDM1 | 1 |
| **Cell Morphology,Cellular Assembly and Organization,Cellular Development,Cellular Function and Maintenance,Cellular Growth and Proliferation,Nervous System Development and Function,Organismal Development,Tissue Development** | Sprouting of rod bipolar cells | 0.0209 | ERC2 | 1 |
| **Cellular Development,Nervous System Development and Function,Tissue Development** | Differentiation of pyramidal neurons | 0.0209 | UNC5D | 1 |
| **Inflammatory Response** | Inflammatory response of cerebral cortex | 0.0209 | KNG1 | 1 |
| **Hereditary Disorder,Neurological Disease,Organismal Injury and Abnormalities,Psychological Disorders,Skeletal and Muscular Disorders** | Huntington disease grade 1 Huntington disease | 0.0209 | SNAP25 | 1 |
| **Embryonic Development,Nervous System Development and Function,Organ Development,Organismal Development,Tissue Development** | Development of septal area | 0.0209 | VAX1 | 1 |
| **Developmental Disorder,Endocrine System Disorders,Nutritional Disease,Organismal Injury and Abnormalities,Reproductive System Disease** | Delayed puberty | 0.0209 | VAX1 | 1 |
| **Cell-To-Cell Signaling and Interaction,Nervous System Development and Function** | Sensitization of trigeminovascular neuron | 0.0209 | KNG1 | 1 |
| **Cellular Assembly and Organization,Cellular Compromise,Nervous System Development and Function** | Retraction of glial cell projections | 0.0209 | GRIA2 | 1 |
| **Behavior** | Perseverance behavior | 0.0209 | NR3C2 | 1 |
| **Behavior** | Impulsivity | 0.0209 | ADCY1 | 1 |
| **Embryonic Development,Nervous System Development and Function,Organ Development,Organismal Development,Tissue Development** | Development of layer IV | 0.0209 | GAP43 | 1 |
| **Metabolic Disease,Neurological Disease,Organismal Injury and Abnormalities,Psychological Disorders** | Stage V Alzheimer disease | 0.0209 | PTGS2 | 1 |
| **Cell Morphology,Cellular Assembly and Organization** | Orientation of plasma membrane projections | 0.0209 | SCN1B | 1 |
| **Cell-To-Cell Signaling and Interaction,Nervous System Development and Function** | Excitation of postsynaptic region | 0.0209 | PCDH8 | 1 |
| **Cellular Assembly and Organization,Cellular Development,Nervous System Development and Function,Tissue Development** | Differentiation of presynaptic terminals | 0.0209 | AGRN | 1 |
| **Cell-To-Cell Signaling and Interaction,Drug Metabolism,Small Molecule Biochemistry** | Catabolism of dopamine | 0.0209 | PRKN | 1 |
| **Nervous System Development and Function,Neurological Disease,Organ Morphology,Organismal** | Lack of optic tract | 0.0209 | GAP43 | 1 |

| | | | | |
|---|---|---|---|---|
| **Development,Organismal Injury and Abnormalities** | | | | |
| **Cell Morphology,Cellular Assembly and Organization,Nervous System Development and Function,Tissue Morphology** | Diameter of nodes of Ranvier | 0.0209 | SPTBN4 | 1 |
| **Cellular Development,Nervous System Development and Function,Tissue Development** | Differentiation of postsynaptic neurons | 0.0209 | AGRN | 1 |
| **Nervous System Development and Function** | Discharge of cochlear nerve | 0.0209 | BSN | 1 |
| **Cell-To-Cell Signaling and Interaction,Nervous System Development and Function** | Firing of trigeminovascular neuron | 0.0209 | KNG1 | 1 |
| **Lipid Metabolism,Molecular Transport,Small Molecule Biochemistry** | Concentration of eicosapentenoic acid | 0.0209 | ARNTL | 1 |
| **Metabolic Disease,Neurological Disease,Organismal Injury and Abnormalities,Psychological Disorders** | Stage I Alzheimer disease | 0.0209 | PTGS2 | 1 |
| **Cell-To-Cell Signaling and Interaction,Cellular Assembly and Organization,Nervous System Development and Function** | Modulation of synapse | 0.0209 | PCDH8 | 1 |
| **Cancer,Neurological Disease,Organismal Injury and Abnormalities** | Growth of glioma | 0.0209 | CAMTA1 | 1 |
| **Cell-To-Cell Signaling and Interaction,Cellular Assembly and Organization,Nervous System Development and Function** | Binding of presynaptic membrane | 0.0209 | AP2M1 | 1 |
| **Cell Morphology,Cell-To-Cell Signaling and Interaction** | Length of stereocilia | 0.0209 | MYO15A | 1 |
| **Nervous System Development and Function** | Function of autonomic nerve | 0.0209 | SNAP25 | 1 |
| **Cell-To-Cell Signaling and Interaction,Nervous System Development and Function** | Inactivation of synapse | 0.0209 | BSN | 1 |
| **Cellular Development,Nervous System Development and Function,Organ Development** | Differentiation of preoptic area | 0.0209 | VAX1 | 1 |
| **Cellular Development,Nervous System Development and Function,Organ Development** | Differentiation of medial ganglionic eminences | 0.0209 | VAX1 | 1 |
| **Cellular Development,Cellular Growth and Proliferation,Embryonic Development,Nervous System Development and Function,Organ Development,Organismal Development,Tissue Development,Visual System Development and Function** | Formation of retinal cells | 0.0209 | ADCY1 | 1 |
| **Nervous System Development and Function,Organ Morphology,Organismal Development** | Size of optic chiasm | 0.0209 | GAP43 | 1 |
| **Cellular Development** | Delay in initiation of transition of radial glial cells | 0.0209 | PLAGL1 | 1 |
| **Cellular Development** | Delay in initiation of transition of neuronal progenitor cells | 0.0209 | PLAGL1 | 1 |
| **Behavior** | Social exploration | 0.0209 | APBA2 | 1 |
| **Hereditary Disorder,Neurological Disease,Organismal Injury and Abnormalities,Psychological Disorders,Skeletal and Muscular Disorders** | Parkinson-dementia syndrome | 0.0209 | GAP43 | 1 |
| **Cell-To-Cell Signaling and Interaction,Nervous System Development and Function** | Hyperactivation of neurons | 0.0209 | SLC24A4 | 1 |
| **Metabolic Disease,Neurological Disease,Organismal Injury and Abnormalities,Psychological Disorders** | Preclinical stage Alzheimer disease | 0.0209 | NOS2 | 1 |
| **Hematological Disease** | Hyperemia | 0.0209 | PTGS2 | 1 |
| **Endocrine System Development and Function,Endocrine System Disorders,Nervous System Development and Function,Neurological Disease,Organ Morphology,Organismal Development,Organismal Injury and Abnormalities,Reproductive System Disease** | Abnormal morphology of enlarged adenohypophysis | 0.0209 | TCF7L2 | 1 |
| **Cellular Assembly and Organization** | Delay in initiation of priming of vesicles | 0.0209 | SNAP25 | 1 |
| **Cell-To-Cell Signaling and Interaction,Drug Metabolism,Small Molecule Biochemistry** | Conversion of norepinephrine | 0.0209 | DBH | 1 |
| **Infectious Diseases** | Replication of Murine cytomegalovirus | 0.0209 | NOS2 | 1 |

| | | | | |
|---|---|---|---|---|
| **Cell-To-Cell Signaling and Interaction,Nervous System Development and Function** | Activation of mitral cells | 0.0209 | GJD2 | 1 |
| **Cell Morphology,Cellular Assembly and Organization,Nervous System Development and Function** | Projection of mossy fibers | 0.0209 | NR3C2 | 1 |
| **Cardiovascular System Development and Function,Nervous System Development and Function,Organ Morphology,Organismal Development,Tissue Morphology** | Diameter of basilar artery | 0.0209 | KNG1 | 1 |
| **Cell Morphology,Cellular Movement,Nervous System Development and Function** | Innervation of serotonergic neurons | 0.0209 | GAP43 | 1 |
| **Cell Morphology,Cellular Assembly and Organization** | Elongation of astrocyte projections | 0.0209 | GAP43 | 1 |
| **Cell Death and Survival,Cellular Compromise,Neurological Disease,Organismal Injury and Abnormalities,Tissue Morphology** | Neurodegeneration of pyramidal neurons | 0.0211 | GRIA2, PTGS2 | 2 |
| **Neurological Disease,Organismal Injury and Abnormalities,Psychological Disorders,Skeletal and Muscular Disorders** | Parkinson disease with dementia | 0.0211 | GAP43, NOS2 | 2 |
| **Cell-To-Cell Signaling and Interaction,Nervous System Development and Function** | Neurotransmission | 0.024 | BSN, ERC2, GRIA2, KNG1, NR3C2, PRKN, RAB11FIP5, RIMS2, SNAP25 | 9 |
| **Cell Death and Survival,Cellular Compromise,Neurological Disease,Tissue Morphology** | Neurodegeneration of dopaminergic neurons | 0.0242 | LRRK2, PRKN, PTGS2 | 3 |
| **Cellular Assembly and Organization,Nervous System Development and Function,Tissue Morphology** | Abnormal quantity of synaptic vesicles | 0.0242 | ERC2, UNC13A, UNC13B | 3 |
| **Behavior,Nervous System Development and Function** | Memory | 0.0245 | ADCY1, KCNH3, NEURL1, NR3C2 | 4 |
| **Nervous System Development and Function,Neurological Disease,Organ Morphology,Organismal Development,Organismal Injury and Abnormalities** | Abnormal morphology of thalamus | 0.0263 | ADCY1, NCOR1, TENM2 | 3 |
| **Cell-To-Cell Signaling and Interaction,Molecular Transport,Small Molecule Biochemistry** | Concentration of 5-hydroxytryptamine | 0.0286 | APBA2, DBH, GAP43 | 3 |
| **Neurological Disease,Organismal Injury and Abnormalities** | Seizures | 0.0286 | BSN, GAP43, GRIA2 | 3 |
| **Drug Metabolism,Lipid Metabolism,Small Molecule Biochemistry** | Synthesis of prostaglandin E2 | 0.0291 | NOS2, PTGS2 | 2 |
| **Carbohydrate Metabolism,Lipid Metabolism,Molecular Transport,Small Molecule Biochemistry** | Concentration of phosphatidic acid | 0.0291 | PIP5K1C, PTGS2 | 2 |
| **Cancer,Neurological Disease,Organismal Injury and Abnormalities** | Glioblastoma | 0.0291 | CAMTA1, ROS1 | 2 |
| **Cellular Assembly and Organization** | Quantity of vesicles | 0.0293 | BSN, ERC2, UNC13A, UNC13B | 4 |
| **Nervous System Development and Function** | Morphology of central nervous system | 0.0298 | ABR, ADCY1, ARNT2, ASPM, CADPS2, CELSR2, EPHB3, FES, GAP43, GRIA2, HIVEP2, KIF5C, KNG1, LRRTM4, NCOR1, NOS2, | 24 |

| | | | | |
|---|---|---|---|---|
| | | | PRKN, RIMS2, SCN1B, SNAP25, SPTBN4, TCF7L2, TENM2, VAX1 | |
| **Cell-To-Cell Signaling and Interaction,Nervous System Development and Function** | Synaptic transmission of cerebral cortex cells | 0.031 | ERC2, GRIA2, RAB11FIP5, SNAP25 | 4 |
| **Cell-To-Cell Signaling and Interaction,Cellular Assembly and Organization,Nervous System Development and Function,Tissue Morphology** | Quantity of synapse | 0.0328 | AGRN, BSN, EPHB3, NEURL1 | 4 |
| **Nervous System Development and Function,Neurological Disease,Organ Morphology,Organismal Development,Organismal Injury and Abnormalities** | Abnormal morphology of third cerebral ventricle | 0.0335 | ARNT2, CELSR2, NCOR1 | 3 |
| **Nervous System Development and Function,Neurological Disease,Organ Morphology,Organismal Development,Organismal Injury and Abnormalities** | Third ventricle dilation | 0.0335 | CELSR2, NCOR1 | 2 |
| **Cell Signaling,Molecular Transport,Vitamin and Mineral Metabolism** | Quantity of Ca2+ | 0.034 | CACNA1C, CCL4, KNG1, NRN1, RANBP9 | 5 |
| **Organismal Development** | Morphology of head | 0.0344 | ABR, ADCY1, ARNT2, ASPM, CADPS2, CELSR2, DVL1, EPHB3, FAT3, FES, GAP43, GRIA2, HIVEP2, KIF5C, KNG1, LRRTM4, NCOR1, NOS2, PRKN, SNAP25, SPTBN4, TCF7L2, TENM2, VAX1 | 24 |
| **Neurological Disease,Psychological Disorders** | Schizophrenia | 0.035 | APBA2, GAP43, MAP10, NELL1, PCDH8, PIK3R1, SNAP25, TCF7L2, TRERF1 | 9 |
| **Behavior** | Learning | 0.0355 | ADCY1, ADGRB3, KCNH3, NEURL1, NR3C2 | 5 |
| **Cell-To-Cell Signaling and Interaction** | Synaptic depression | 0.0358 | ADCY1, ADCY5, GRIA2, PIP5K1C, PTK2B, RAB11FIP5, UNC13A | 7 |

| | | | | |
|---|---|---|---|---|
| **Cellular Assembly and Organization,Nervous System Development and Function** | Fasciculation of axons | 0.0361 | EPHB3, GAP43, MEGF8 | 3 |
| **Cell Morphology,Nervous System Development and Function,Neurological Disease,Tissue Morphology** | Abnormal morphology of neurites | 0.0374 | CADPS2, EPHB3, GAP43, LRRK2, NRN1, SCN1B, SEMA5B, SPTBN4 | 8 |
| **Cell-To-Cell Signaling and Interaction,Cellular Assembly and Organization,Nervous System Development and Function** | Fusion of synaptic vesicles | 0.0382 | BSN, SNAP25 | 2 |
| **Cell-To-Cell Signaling and Interaction** | Activation of central nervous system cells | 0.0384 | GJD2, MGAT5B, PTGS2, SCN1B | 4 |
| **Cellular Function and Maintenance** | Endocytosis | 0.0384 | AP2M1, KIAA1107, PIP5K1C, SNAP25 | 4 |
| **Nervous System Development and Function,Organ Morphology,Organismal Development** | Morphology of brain | 0.0395 | ABR, ADCY1, ARNT2, ASPM, CADPS2, CELSR2, EPHB3, FES, GAP43, GRIA2, HIVEP2, KIF5C, KNG1, LRRTM4, NCOR1, NOS2, PRKN, SNAP25, SPTBN4, TCF7L2, TENM2, VAX1 | 22 |
| **Nervous System Development and Function,Neurological Disease,Organ Morphology,Organismal Development,Organismal Injury and Abnormalities** | Abnormal morphology of forebrain | 0.04 | ADCY1, ARNT2, ASPM, CELSR2, FES, HIVEP2, NCOR1, NOS2, TENM2, VAX1 | 10 |
| **Nervous System Development and Function** | Dedepression of CA1 neuron | 0.0414 | GRIA2 | 1 |
| **Cellular Development,Nervous System Development and Function,Tissue Development** | Differentiation of neuron-like cells | 0.0414 | PIP5K1C | 1 |
| **Cardiovascular System Development and Function,Organismal Development,Tissue Morphology** | Vasodilation of capillary vessel | 0.0414 | UNC13B | 1 |
| **Nervous System Development and Function** | Depotentiation of CA1 neuron | 0.0414 | GRIA2 | 1 |
| **Cell Morphology,Cellular Assembly and Organization,Cellular Development,Cellular Function and Maintenance,Cellular Growth and Proliferation,Nervous System Development and Function,Organismal Development,Tissue Development** | Branching of axon branches | 0.0414 | EPHB3 | 1 |
| **Cellular Assembly and Organization,Cellular Movement,Nervous System Development and Function** | Migration of axons | 0.0414 | EPHB3 | 1 |
| **Cell Death and Survival,Cell Morphology,Nervous System Development and** | Loss of basal ganglia neuron | 0.0414 | VPS13A | 1 |

| | | | | |
|---|---|---|---|---|
| **Function,Neurological Disease,Tissue Morphology** | | | | |
| **Inflammatory Disease,Inflammatory Response,Neurological Disease,Organismal Injury and Abnormalities** | Inflammation of substantia nigra | 0.0414 | LRRK2 | 1 |
| **Cellular Growth and Proliferation,Nervous System Development and Function** | Inhibition of dorsal root ganglion cells | 0.0414 | ROCK2 | 1 |
| **Cell Morphology,Cellular Assembly and Organization,Cellular Development,Cellular Function and Maintenance,Cellular Growth and Proliferation,Embryonic Development,Nervous System Development and Function,Organismal Development,Tissue Development** | Branching of sympathetic neurites | 0.0414 | GAP43 | 1 |
| **Cell Cycle,Embryonic Development,Nervous System Development and Function** | Cell division of neuroblasts | 0.0414 | ASPM | 1 |
| **Developmental Disorder,Endocrine System Disorders,Hereditary Disorder,Metabolic Disease,Neurological Disease,Organismal Injury and Abnormalities,Psychological Disorders** | Childhood adrenoleukodystrophy | 0.0414 | PTGS2 | 1 |
| **Cell-To-Cell Signaling and Interaction,Nervous System Development and Function** | Action potential of superior cervical ganglion neurons | 0.0414 | KNG1 | 1 |
| **Lipid Metabolism,Small Molecule Biochemistry** | Liberation of stearic acid | 0.0414 | KNG1 | 1 |
| **Cellular Assembly and Organization** | Quantity of dense-core vesicles | 0.0414 | BSN | 1 |
| **Cellular Assembly and Organization** | Activation of mitochondria | 0.0414 | RANBP9 | 1 |
| **Cell-To-Cell Signaling and Interaction,Nervous System Development and Function** | Coupling of neurons | 0.0414 | GJD2 | 1 |
| **Cellular Assembly and Organization** | Translocation of Golgi vesicles | 0.0414 | BSN | 1 |
| **Cell-To-Cell Signaling and Interaction,Nervous System Development and Function** | Long term synaptic depression of corticostriatal neurons | 0.0414 | ADCY5 | 1 |
| **Cancer,Neurological Disease,Organismal Injury and Abnormalities,Tumor Morphology** | Production of glioblastoma | 0.0414 | ROS1 | 1 |
| **Cell-To-Cell Signaling and Interaction,Nervous System Development and Function** | Short-term depression of synapse | 0.0414 | UNC13A | 1 |
| **Cell Morphology,Cellular Assembly and Organization,Nervous System Development and Function** | Extension of axon branches | 0.0414 | EPHB3 | 1 |
| **Cellular Assembly and Organization,Cellular Function and Maintenance,Cellular Movement,Nervous System Development and Function** | Arrest in axonal transport of mitochondria | 0.0414 | PRKN | 1 |
| **Cell-To-Cell Signaling and Interaction,Nervous System Development and Function** | Neurotransmission of thalamocortical axons | 0.0414 | RIMS2 | 1 |
| **Cardiovascular Disease,Hematological Disease,Organismal Injury and Abnormalities** | Thrombus | 0.0414 | KNG1 | 1 |
| **Cardiovascular System Development and Function** | Hypoperfusion of brain | 0.0414 | NOS2 | 1 |
| **Nervous System Development and Function,Tissue Morphology** | Quantity of catecholaminergic neurons | 0.0414 | PRKN | 1 |
| **Nervous System Development and Function,Neurological Disease** | Abnormal morphology of maxillary nerve | 0.0414 | MEGF8 | 1 |
| **Cell-To-Cell Signaling and Interaction** | Synaptic strength of GABAergic synapse | 0.0414 | AGRN | 1 |
| **Reproductive System Development and Function** | Estrous cyclicity | 0.0414 | GJD2 | 1 |
| **Carbohydrate Metabolism,Lipid Metabolism,Molecular Transport,Small Molecule Biochemistry** | Concentration of phosphatidylserine | 0.0414 | PTGS2 | 1 |
| **Cellular Assembly and Organization,Cellular Function and Maintenance** | Formation of Golgi vesicles | 0.0414 | BSN | 1 |
| **Lipid Metabolism,Small Molecule Biochemistry** | Liberation of arachidonic acid | 0.0414 | KNG1 | 1 |
| **Endocrine System Development and Function,Endocrine System Disorders,Nervous System Development and Function,Neurological Disease,Organ Morphology,Organismal Development,Organismal Injury and Abnormalities,Reproductive System Disease** | Abnormal morphology of neurohypophysis | 0.0414 | ARNT2 | 1 |

| | | | | |
|---|---|---|---|---|
| **Cell-To-Cell Signaling and Interaction,Nervous System Development and Function** | Long term synaptic depression of striatopallidal neurons | 0.0414 | ADCY5 | 1 |
| **Metabolic Disease,Neurological Disease,Organismal Injury and Abnormalities,Psychological Disorders** | Stage II Alzheimer disease | 0.0414 | PTGS2 | 1 |
| **Neurological Disease,Organismal Injury and Abnormalities,Psychological Disorders** | Damage of brain tissue | 0.0414 | NOS2 | 1 |
| **Cellular Assembly and Organization,Cellular Movement** | Segregation of retinogeniculate projection | 0.0414 | GJD2 | 1 |
| **Cell-To-Cell Signaling and Interaction,Cellular Assembly and Organization,Cellular Development,Cellular Function and Maintenance,Cellular Growth and Proliferation,Nervous System Development and Function,Tissue Development** | Formation of neuromuscular synapse | 0.0414 | AGRN | 1 |
| **Cardiovascular Disease,Organismal Injury and Abnormalities** | Failure of heart | 0.0414 | PTGS2 | 1 |
| **Nervous System Development and Function,Organ Morphology,Tissue Morphology** | Quantity of spiny stellate neurons | 0.0414 | RIMS2 | 1 |
| **Endocrine System Development and Function,Endocrine System Disorders,Nervous System Development and Function,Neurological Disease,Organ Development,Organ Morphology,Organismal Development,Organismal Injury and Abnormalities** | Abnormal morphology of supraoptic nucleus | 0.0414 | ARNT2 | 1 |
| **Cell-To-Cell Signaling and Interaction,Inflammatory Response** | Cytotoxic reaction of macrophages | 0.0414 | NOS2 | 1 |
| **Cell-To-Cell Signaling and Interaction** | Dissociation of endothelial cells | 0.0414 | UNC13B | 1 |
| **Cellular Assembly and Organization,Cellular Function and Maintenance,Cellular Movement,Nervous System Development and Function** | Endocytosis of synaptic vesicles | 0.0416 | AP2M1, KIAA1107, PIP5K1C | 3 |
| **Cell-To-Cell Signaling and Interaction,Nervous System Development and Function** | Synaptic transmission of hippocampal cells | 0.0416 | ERC2, GRIA2, RAB11FIP5 | 3 |
| **Cell-To-Cell Signaling and Interaction,Cellular Growth and Proliferation,Nervous System Development and Function** | Excitation of neurons | 0.0425 | KCNH3, KNG1, NRN1, PRKN | 4 |
| **Cell-To-Cell Signaling and Interaction** | Activation of cells | 0.0427 | GJD2, LRRK2, MGAT5B, NOS2, PTGS2, SCN1B, SLC24A4, TRPM2 | 8 |
| **Behavior** | Social behavior | 0.043 | DVL1, GRIA2 | 2 |
| **Cellular Assembly and Organization,Nervous System Development and Function** | Remodeling of neurites | 0.043 | ADGRB3, DVL1 | 2 |
| **Cancer,Neurological Disease,Organismal Injury and Abnormalities** | Astrocytoma in brainstem | 0.043 | RYR2, THEMIS | 2 |
| **Cell Morphology,Cellular Movement,Nervous System Development and Function** | Innervation of neurons | 0.0435 | EPHB3, GAP43, GRIA2, RIMS2, TENM2 | 5 |
| **Cell-To-Cell Signaling and Interaction,Cellular Assembly and Organization,Nervous System Development and Function,Tissue Morphology** | Density of synapse | 0.0445 | EPHB3, LRRTM4, MEF2A | 3 |
| **Nervous System Development and Function,Neurological Disease,Organ Morphology,Organismal Development,Organismal Injury and Abnormalities** | Abnormal morphology of diencephalon | 0.0452 | ADCY1, ARNT2, CELSR2, NCOR1, TENM2 | 5 |
| **Nervous System Development and Function,Organ Morphology,Organismal Development** | Morphology of forebrain | 0.0477 | ADCY1, ARNT2, ASPM, CELSR2, | 11 |

| | | | | |
|---|---|---|---|---|
| | | | FES, GAP43, HIVEP2, NCOR1, NOS2, TENM2, VAX1 | |
| **Hematological System Development and Function,Organismal Functions** | Flow of blood | 0.0481 | KNG1, PTGS2 | 2 |
| **Cell Death and Survival,Cellular Compromise,Neurological Disease,Organismal Injury and Abnormalities,Tissue Morphology** | Neurodegeneration of hippocampal neurons | 0.0481 | GRIA2, PRKN | 2 |

**Cluster 12**

| Categories | Diseases or Functions Annotation | p-Value | Molecules | # Molecules |
|---|---|---|---|---|
| **Cellular Assembly and Organization,Cellular Function and Maintenance** | Microtubule dynamics | 0.000021 | ABI1, ADCY1, ADGRB3, AGRN, ARHGEF28, ATCAY, BCL11B, BSN, CELSR2, CRMP1, DBN1, DVL1, ERC2, FMN1, GABRB3, GAP43, GPRIN1, HTR1A, LAMB1, MEF2A, MRTFA, MYO16, MYO5B, NRN1, PTGS2, SEMA4F, SEMA5B, SHANK2, SIPA1L1, SORBS2 | 30 |
| **Cellular Development,Cellular Growth and Proliferation,Nervous System Development and Function,Tissue Development** | Development of neurons | 0.0000213 | ABI1, ADCY1, ADGRB3, AGRN, ARHGEF28, ATCAY, BCL11B, BSN, CELSR2, CRMP1, DBN1, DVL1, ERC2, FMN1, GABRB3, GAP43, GPRIN1, LAMB1, LRRTM4, MEF2A, MRTFA, MYO16, MYO5B, NCAN, NRN1, PCDH8, PLXND1, | 31 |

| | | | | |
|---|---|---|---|---|
| | | | PTGS2, SHANK2, SIPA1L1, SORBS2 | |
| **Cell Morphology,Cellular Assembly and Organization,Cellular Development,Cellular Function and Maintenance,Cellular Growth and Proliferation,Nervous System Development and Function,Organismal Development,Tissue Development** | Neuritogenesis | 0.00003 | ABI1, ADCY1, ADGRB3, AGRN, ARHGEF28, ATCAY, BCL11B, BSN, CELSR2, CRMP1, DBN1, DVL1, ERC2, FMN1, GABRB3, GAP43, GPRIN1, LAMB1, MEF2A, MRTFA, MYO16, MYO5B, NRN1, PTGS2, SHANK2, SIPA1L1, SORBS2 | 27 |
| **Cell-To-Cell Signaling and Interaction,Cellular Assembly and Organization** | Cell-cell contact | 0.0000482 | AGRN, BSN, CAMK2D, ERC2, GAP43, LRRTM4, NCAN, NEURL1, NRN1, PCDH8, PLXND1, SHANK2, SNTA1, SORBS2, UNC13A | 15 |
| **Cancer,Neurological Disease,Organismal Injury and Abnormalities** | Brain astrocytoma | 0.000444 | ADAMTS8, AGAP3, ANKRD30B, ASXL3, ATF7IP2, COL11A1, DPP10, ERC2, ICE1, MYO5B, P2RX5, PIK3R1, PLB1, PRDM2, PTGS2, RIMS2, SEMA4F, THEMIS | 18 |
| **Nervous System Development and Function,Neurological Disease** | Abnormal morphology of nervous system | 0.000475 | ABR, ADCY1, AGRN, ARNT2, CACNA1C, CELSR2, CRMP1, DBN1, DVL1, E2F5, ERC2, | 29 |

| | | | | |
|---|---|---|---|---|
| | | | FAT3, GABRB3, GAP43, GMFB, GRIA2, LRRTM4, MYO16, NOS2, NRN1, PRKN, RIMS2, SEMA5B, SHANK2, SHC2, SNTA1, SORBS2, UNC13A, VPS13A | |
| **Hereditary Disorder,Neurological Disease,Organismal Injury and Abnormalities,Psychological Disorders,Skeletal and Muscular Disorders** | Familial Parkinson disease | 0.000645 | GAP43, PRKN | 2 |
| **Nervous System Development and Function,Organismal Development** | Secretion of cerebrospinal fluid | 0.000645 | CELSR2, E2F5 | 2 |
| **Cell Morphology,Cellular Assembly and Organization,Cellular Development,Cellular Function and Maintenance,Cellular Growth and Proliferation,Embryonic Development,Nervous System Development and Function,Organismal Development,Tissue Development** | Dendritic growth/branching | 0.000657 | ABI1, AGRN, BSN, CRMP1, DBN1, GABRB3, GAP43, MYO5B, NRN1, PTGS2, SHANK2, SIPA1L1, SORBS2 | 13 |
| **Cell-To-Cell Signaling and Interaction,Cellular Assembly and Organization,Cellular Development,Cellular Function and Maintenance,Cellular Growth and Proliferation,Nervous System Development and Function,Tissue Development** | Developmental process of synapse | 0.000764 | AGRN, BSN, LRRTM4, NCAN, NRN1, PCDH8, PLXND1, SHANK2, SORBS2 | 9 |
| **Nervous System Development and Function** | Morphology of nervous system | 0.00111 | ABR, ADCY1, AGRN, ARNT2, CACNA1C, CELSR2, CRMP1, DBN1, DVL1, E2F5, ERC2, FAT3, GABRB3, GAP43, GMFB, GRIA2, KIF5C, LRRTM4, MYO16, NOS2, NRN1, PRDM16, PRKN, RIMS2, SEMA5B, SHANK2, SHC2, SIPA1L1, SNTA1, | 32 |

| | | | | |
|---|---|---|---|---|
| | | | SORBS2, UNC13A, VPS13A | |
| **Cell Morphology,Nervous System Development and Function,Neurological Disease,Tissue Morphology** | Abnormal morphology of neurons | 0.00133 | AGRN, CACNA1C, CRMP1, DBN1, DVL1, ERC2, FAT3, GAP43, MYO16, NOS2, NRN1, PRKN, SEMA5B, SHANK2, SHC2, SNTA1, SORBS2, UNC13A, VPS13A | 19 |
| **Cell Morphology,Cellular Development,Embryonic Development,Nervous System Development and Function,Tissue Development** | Patterning of dendrites | 0.00211 | CELSR2, CRMP1 | 2 |
| **Cellular Development,Nervous System Development and Function,Tissue Development** | Differentiation of nerve ending | 0.00211 | AGRN, MEF2A | 2 |
| **Cell Morphology,Cellular Assembly and Organization,Cellular Development,Cellular Function and Maintenance,Cellular Growth and Proliferation,Nervous System Development and Function,Organismal Development,Tissue Development** | Branching of neurons | 0.00229 | ABI1, AGRN, BSN, CRMP1, DBN1, ERC2, GABRB3, GAP43, MYO5B, NRN1, PTGS2, SHANK2, SIPA1L1, SORBS2 | 14 |
| **Cellular Assembly and Organization** | Organization of organelle | 0.00231 | ADGRV1, AGRN, ARHGEF28, DVL1, LRRTM4, PTPRO, SHANK2 | 7 |
| **Cell Morphology** | Shape change of neurites | 0.0024 | ABI1, AGRN, BCL11B, BSN, CRMP1, DBN1, GABRB3, GAP43, MYO5B, NRN1, PTGS2, SHANK2, SIPA1L1, SORBS2 | 14 |
| **Cell Morphology,Nervous System Development and Function,Neurological Disease,Tissue Morphology** | Abnormal morphology of bipolar cells | 0.003 | ERC2, FAT3, SEMA5B | 3 |
| **Cell-To-Cell Signaling and Interaction,Cellular Assembly and Organization,Cellular Function and Maintenance,Molecular Transport** | Priming of synaptic vesicles | 0.003 | ERC2, RIMS2, UNC13A | 3 |
| **Neurological Disease,Organismal Injury and Abnormalities** | Hydrocephalus | 0.003 | CELSR2, DVL1, E2F5 | 3 |
| **Nervous System Development and Function** | Depotentiation | 0.00313 | ADCY1, GRIA2 | 2 |
| **Cancer,Organismal Injury and Abnormalities** | Non-hematological solid tumor | 0.00319 | ADAMTS8, AGAP3, | 20 |

| | | | | |
|---|---|---|---|---|
| | | | ANKRD30B, ASXL3, ATF7IP2, COL11A1, CUL9, DPP10, ERC2, ICE1, MYO5B, P2RX5, PIGQ, PIK3R1, PLB1, PRDM2, PTGS2, RIMS2, SEMA4F, THEMIS | |
| **Cell Morphology,Cell-To-Cell Signaling and Interaction,Cellular Assembly and Organization,Nervous System Development and Function,Tissue Morphology** | Morphology of synapse | 0.00345 | AGRN, ERC2, GAP43, NRN1, SNTA1, UNC13A | 6 |
| **Cell Morphology,Nervous System Development and Function,Tissue Morphology** | Morphology of neurons | 0.00394 | AGRN, CACNA1C, CRMP1, DBN1, DVL1, ERC2, FAT3, GAP43, MYO16, NOS2, NRN1, PRKN, RIMS2, SEMA5B, SHANK2, SHC2, SIPA1L1, SNTA1, SORBS2, UNC13A, VPS13A | 21 |
| **Nervous System Development and Function** | Sensory system development | 0.00395 | ADCY1, DVL1, FAT3, KIF5C, SHC2 | 5 |
| **Cellular Compromise,Neurological Disease,Organismal Injury and Abnormalities** | Neurodegeneration of hippocampal cells | 0.00397 | GRIA2, PRKN, PTGS2 | 3 |
| **Tissue Development** | Organization of nervous tissue | 0.004 | AGRN, ARHGEF28, DVL1, LRRTM4, PTPRO, SHANK2 | 6 |
| **Cell Morphology,Cellular Assembly and Organization,Cellular Development,Cellular Function and Maintenance,Cellular Growth and Proliferation,Nervous System Development and Function,Organismal Development,Tissue Development** | Formation of dendrites | 0.0043 | ABI1, ADGRB3, CELSR2, DVL1, FMN1, MEF2A, MYO5B, SIPA1L1 | 8 |
| **Cell-To-Cell Signaling and Interaction,Drug Metabolism,Small Molecule Biochemistry** | Metabolism of dopamine | 0.00434 | PRKN, SORCS2 | 2 |
| **Cancer,Organismal Injury and Abnormalities** | Head and neck tumor | 0.0045 | ADAMTS8, AGAP3, ANKRD30B, ASXL3, ATF7IP2, COL11A1, | 19 |

| | | | | |
|---|---|---|---|---|
| | | | CUL9, DPP10, ERC2, ICE1, MYO5B, P2RX5, PIK3R1, PLB1, PRDM2, PTGS2, RIMS2, SEMA4F, THEMIS | |
| **Organismal Development** | Morphology of head | 0.00515 | ABR, ADCY1, ADGRV1, ARNT2, CELSR2, CRMP1, DBN1, DVL1, E2F5, FAT3, GABRB3, GAP43, GMFB, GRIA2, KIF5C, LRRTM4, MYO16, NOS2, PRDM16, PRKN, SHANK2 | 21 |
| **Cell-To-Cell Signaling and Interaction,Cellular Assembly and Organization** | Stabilization of synapse | 0.00573 | NCAN, NRN1 | 2 |
| **Cellular Function and Maintenance,Molecular Transport** | Exocytosis | 0.00582 | ERC2, RIMS2, RPH3A, UNC13A | 4 |
| **Cancer,Neurological Disease,Organismal Injury and Abnormalities** | Nervous system neoplasm | 0.00643 | ADAMTS8, AGAP3, ANKRD30B, ASXL3, ATF7IP2, COL11A1, DPP10, ERC2, ICE1, MYO5B, P2RX5, PIGQ, PIK3R1, PLB1, PRDM2, PTGS2, RIMS2, SEMA4F, THEMIS | 19 |
| **Cellular Assembly and Organization,Cellular Movement,Nervous System Development and Function** | Transport of synaptic vesicles | 0.0069 | AP2M1, DENND1A, KIAA1107, RPH3A, UNC13A | 5 |
| **Organismal Development,Organismal Injury and Abnormalities** | Abnormal morphology of head | 0.00695 | ABR, ADCY1, ADGRV1, ARNT2, CELSR2, CRMP1, DBN1, DVL1, E2F5, FAT3, GABRB3, | 18 |

| | | | | |
|---|---|---|---|---|
| | | | GAP43, GMFB, GRIA2, LRRTM4, NOS2, PRKN, SHANK2 | |
| **Cell-To-Cell Signaling and Interaction,Cellular Assembly and Organization,Cellular Function and Maintenance,Tissue Development** | Organization of synapse | 0.00727 | AGRN, DVL1, LRRTM4, PTPRO, SHANK2 | 5 |
| **Cell-To-Cell Signaling and Interaction,Nervous System Development and Function** | Long-term potentiation of mossy fibers | 0.0073 | ADCY1, GRIA2 | 2 |
| **Embryonic Development,Nervous System Development and Function,Organ Development,Organismal Development,Tissue Development,Visual System Development and Function** | Formation of eye | 0.00904 | ADCY1, FAT3 | 2 |
| **Cell-To-Cell Signaling and Interaction,Cellular Assembly and Organization,Cellular Development,Cellular Function and Maintenance,Cellular Growth and Proliferation,Nervous System Development and Function,Tissue Development** | Formation of excitatory synapses | 0.00904 | LRRTM4, SORBS2 | 2 |
| **Tissue Morphology** | Quantity of cells | 0.00925 | AGRN, BSN, CACNA1C, CAMK2D, DVL1, ERC2, GBA, KIF5C, MGAT5B, NEURL1, NOS2, NRN1, PRDM16, PRKN, PTGS2, RIMS2, SHC2, UNC13A | 18 |
| **Behavior** | Learning | 0.00928 | ADCY1, ADGRB3, GABRB3, KCNH3, NEURL1 | 5 |
| **Cell-To-Cell Signaling and Interaction,Molecular Transport,Small Molecule Biochemistry** | Release of neurotransmitter | 0.0099 | DVL1, GABRB3, HTR1A, PRKN | 4 |
| **Cell Morphology,Cell-To-Cell Signaling and Interaction,Cellular Assembly and Organization,Nervous System Development and Function,Neurological Disease,Tissue Morphology** | Abnormal morphology of synapse | 0.0102 | AGRN, ERC2, NRN1, SNTA1, UNC13A | 5 |
| **Cell-To-Cell Signaling and Interaction,Cellular Assembly and Organization,Nervous System Development and Function,Tissue Morphology** | Quantity of synapse | 0.0105 | AGRN, BSN, CAMK2D, NEURL1 | 4 |
| **Cell Death and Survival,Cellular Compromise,Neurological Disease,Organismal Injury and Abnormalities,Tissue Morphology** | Neurodegeneration of pyramidal neurons | 0.0109 | GRIA2, PTGS2 | 2 |
| **Neurological Disease,Organismal Injury and Abnormalities,Psychological Disorders,Skeletal and Muscular Disorders** | Parkinson disease with dementia | 0.0109 | GAP43, NOS2 | 2 |
| **Neurological Disease,Organismal Injury and Abnormalities** | Seizures | 0.0115 | BSN, GAP43, GRIA2 | 3 |
| **Auditory and Vestibular System Development and Function,Cell Morphology,Cellular Assembly and Organization** | Orientation of stereocilia bundles | 0.013 | ADGRV1, DVL1 | 2 |

| | | | | |
|---|---|---|---|---|
| **Nervous System Development and Function,Neurological Disease,Organ Morphology,Organismal Development,Organismal Injury and Abnormalities** | Abnormal morphology of third cerebral ventricle | 0.0136 | ARNT2, CELSR2, E2F5 | 3 |
| **Cell-To-Cell Signaling and Interaction,Cellular Assembly and Organization,Nervous System Development and Function** | Modulation of synapse | 0.0148 | PCDH8 | 1 |
| **Cellular Development,Nervous System Development and Function,Tissue Development** | Differentiation of sympathetic preganglionic neurons | 0.0148 | AGRN | 1 |
| **Cell-To-Cell Signaling and Interaction,Cellular Assembly and Organization,Nervous System Development and Function** | Binding of presynaptic membrane | 0.0148 | AP2M1 | 1 |
| **Cell-To-Cell Signaling and Interaction,Cellular Assembly and Organization,Nervous System Development and Function,Tissue Morphology** | Quantity of ribbon synapse | 0.0148 | BSN | 1 |
| **Cell Signaling,Nucleic Acid Metabolism,Small Molecule Biochemistry** | Inhibition of cyclic AMP | 0.0148 | HTR1A | 1 |
| **Nervous System Development and Function,Tissue Morphology** | Density of serotonergic neurons | 0.0148 | GAP43 | 1 |
| **Hereditary Disorder,Neurological Disease,Organismal Injury and Abnormalities,Psychological Disorders,Skeletal and Muscular Disorders** | Autosomal recessive juvenile Parkinson disease | 0.0148 | PRKN | 1 |
| **Cell-To-Cell Signaling and Interaction,Nervous System Development and Function** | Inactivation of synapse | 0.0148 | BSN | 1 |
| **Cellular Development,Cellular Growth and Proliferation,Embryonic Development,Nervous System Development and Function,Organ Development,Organismal Development,Tissue Development,Visual System Development and Function** | Formation of retinal cells | 0.0148 | ADCY1 | 1 |
| **Cell-To-Cell Signaling and Interaction,Nervous System Development and Function** | Clustering of hippocampal neurons | 0.0148 | SHANK2 | 1 |
| **Molecular Transport,Protein Trafficking** | Translocation of protein fragment | 0.0148 | AGAP3 | 1 |
| **Carbohydrate Metabolism,Lipid Metabolism,Small Molecule Biochemistry** | Incorporation of phosphatidylcholine | 0.0148 | GBA | 1 |
| **Nervous System Development and Function,Organ Morphology,Organismal Development** | Size of optic chiasm | 0.0148 | GAP43 | 1 |
| **Nervous System Development and Function,Organ Morphology,Tissue Morphology** | Quantity of place cells | 0.0148 | CAMK2D | 1 |
| **Cell Morphology,Cellular Assembly and Organization,Cellular Development,Cellular Function and Maintenance,Cellular Growth and Proliferation,Nervous System Development and Function,Organismal Development,Tissue Development** | Sprouting of rod bipolar cells | 0.0148 | ERC2 | 1 |
| **Cellular Development,Nervous System Development and Function,Tissue Development** | Differentiation of pyramidal neurons | 0.0148 | UNC5D | 1 |
| **Hereditary Disorder,Neurological Disease,Organismal Injury and Abnormalities,Psychological Disorders,Skeletal and Muscular Disorders** | Parkinson-dementia syndrome | 0.0148 | GAP43 | 1 |
| **Lipid Metabolism,Small Molecule Biochemistry** | Incorporation of choline | 0.0148 | GBA | 1 |
| **Neurological Disease** | Mechanical hypersensitivity of neurons | 0.0148 | CACNA2D1 | 1 |
| **Cellular Assembly and Organization,Cellular Compromise,Nervous System Development and Function** | Retraction of glial cell projections | 0.0148 | GRIA2 | 1 |
| **Lipid Metabolism,Molecular Transport,Small Molecule Biochemistry** | Accumulation of glucosylceramide | 0.0148 | GBA | 1 |
| **Cellular Development,Cellular Growth and Proliferation,Nervous System Development and Function,Tissue Development** | Development of corticospinal neurons | 0.0148 | BCL11B | 1 |
| **Behavior** | Impulsivity | 0.0148 | ADCY1 | 1 |
| **Cell-To-Cell Signaling and Interaction,Nervous System Development and Function** | Hyperactivation of neurons | 0.0148 | SLC24A4 | 1 |

| | | | | |
|---|---|---|---|---|
| **Metabolic Disease,Neurological Disease,Organismal Injury and Abnormalities,Psychological Disorders** | Preclinical stage Alzheimer disease | 0.0148 | NOS2 | 1 |
| **Hematological Disease** | Hyperemia | 0.0148 | PTGS2 | 1 |
| **Embryonic Development,Nervous System Development and Function,Organ Development,Organismal Development,Tissue Development** | Development of layer IV | 0.0148 | GAP43 | 1 |
| **Infectious Diseases** | Replication of Murine cytomegalovirus | 0.0148 | NOS2 | 1 |
| **Behavior** | Risk preference | 0.0148 | GABRB3 | 1 |
| **Metabolic Disease,Neurological Disease,Organismal Injury and Abnormalities,Psychological Disorders** | Stage V Alzheimer disease | 0.0148 | PTGS2 | 1 |
| **Lipid Metabolism,Small Molecule Biochemistry** | Incorporation of sphingomyelin | 0.0148 | GBA | 1 |
| **Cell-To-Cell Signaling and Interaction,Nervous System Development and Function** | Excitation of postsynaptic region | 0.0148 | PCDH8 | 1 |
| **Cellular Assembly and Organization,Cellular Development,Nervous System Development and Function,Tissue Development** | Differentiation of presynaptic terminals | 0.0148 | AGRN | 1 |
| **Cell-To-Cell Signaling and Interaction,Drug Metabolism,Small Molecule Biochemistry** | Catabolism of dopamine | 0.0148 | PRKN | 1 |
| **Nervous System Development and Function,Neurological Disease,Organ Morphology,Organismal Development,Organismal Injury and Abnormalities** | Lack of optic tract | 0.0148 | GAP43 | 1 |
| **Cellular Development,Nervous System Development and Function,Tissue Development** | Differentiation of postsynaptic neurons | 0.0148 | AGRN | 1 |
| **Nervous System Development and Function** | Discharge of cochlear nerve | 0.0148 | BSN | 1 |
| **Cell Morphology,Cellular Movement,Nervous System Development and Function** | Innervation of serotonergic neurons | 0.0148 | GAP43 | 1 |
| **Cell Morphology,Cellular Assembly and Organization** | Elongation of astrocyte projections | 0.0148 | GAP43 | 1 |
| **Metabolic Disease,Neurological Disease,Organismal Injury and Abnormalities,Psychological Disorders** | Stage I Alzheimer disease | 0.0148 | PTGS2 | 1 |
| **Nervous System Development and Function,Neurological Disease,Organ Morphology,Organismal Development,Organismal Injury and Abnormalities** | Abnormal morphology of barrel cortex | 0.0152 | ADCY1, GAP43 | 2 |
| **Drug Metabolism,Lipid Metabolism,Small Molecule Biochemistry** | Synthesis of prostaglandin E2 | 0.0152 | NOS2, PTGS2 | 2 |
| **Neurological Disease,Organismal Injury and Abnormalities** | Abnormality of cerebral cortex | 0.0165 | ADCY1, CRMP1, DBN1, E2F5, GAP43, GRIA2, LRRTM4, PRKN, PTGS2, TP53BP2 | 10 |
| **Nervous System Development and Function,Organ Morphology,Organismal Development** | Morphology of brain | 0.0167 | ABR, ADCY1, ARNT2, CELSR2, CRMP1, DBN1, E2F5, GABRB3, GAP43, GMFB, GRIA2, KIF5C, LRRTM4, MYO16, NOS2, PRDM16, PRKN, SHANK2 | 18 |

| | | | | |
|---|---|---|---|---|
| **Cellular Assembly and Organization,Cellular Function and Maintenance,Cellular Movement,Nervous System Development and Function** | Endocytosis of synaptic vesicles | 0.0171 | AP2M1, DENND1A, KIAA1107 | 3 |
| **Cellular Assembly and Organization,Nervous System Development and Function** | Docking of synaptic vesicles | 0.0176 | RIMS2, UNC13A | 2 |
| **Nervous System Development and Function,Neurological Disease,Organ Morphology,Organismal Development,Organismal Injury and Abnormalities** | Third ventricle dilation | 0.0176 | CELSR2, E2F5 | 2 |
| **Nervous System Development and Function** | Morphology of central nervous system | 0.0179 | ABR, ADCY1, ARNT2, CELSR2, CRMP1, DBN1, E2F5, GABRB3, GAP43, GMFB, GRIA2, KIF5C, LRRTM4, MYO16, NOS2, PRDM16, PRKN, RIMS2, SHANK2 | 19 |
| **Cell Morphology,Cell-To-Cell Signaling and Interaction,Nervous System Development and Function** | Plasticity of synapse | 0.0182 | DBN1, GRIA2, KCNH3, NEURL1, PCDH8, SIPA1L1 | 6 |
| **Nervous System Development and Function,Tissue Morphology** | Density of neurons | 0.0212 | CRMP1, GABRB3, GAP43, LRRTM4, MEF2A, PTGS2 | 6 |
| **Cell-To-Cell Signaling and Interaction,Nervous System Development and Function** | Long-term potentiation | 0.0224 | ADCY1, CHST10, CRMP1, GBA, GRIA2, KCNH3, NCAN, PCDH8, SHANK2 | 9 |
| **Behavior** | Social behavior | 0.0227 | DVL1, GRIA2 | 2 |
| **Cellular Assembly and Organization,Nervous System Development and Function** | Remodeling of neurites | 0.0227 | ADGRB3, DVL1 | 2 |
| **Nervous System Development and Function,Neurological Disease** | Abnormal morphology of central nervous system | 0.0234 | ABR, ADCY1, ARNT2, CELSR2, CRMP1, DBN1, E2F5, GABRB3, GAP43, GMFB, GRIA2, LRRTM4, NOS2, PRKN, RIMS2, SHANK2 | 16 |
| **Nervous System Development and Function,Neurological Disease,Organ Morphology,Organismal** | Abnormal morphology of brain | 0.025 | ABR, ADCY1, ARNT2, CELSR2, | 15 |

| | | | | |
|---|---|---|---|---|
| **Development,Organismal Injury and Abnormalities** | | | CRMP1, DBN1, E2F5, GABRB3, GAP43, GMFB, GRIA2, LRRTM4, NOS2, PRKN, SHANK2 | |
| **Cell Death and Survival,Nervous System Development and Function** | Cell viability of cortical neurons | 0.0254 | ADCY1, NRN1, PRKN | 3 |
| **Nervous System Development and Function,Tissue Morphology** | Quantity of neurons | 0.0255 | AGRN, BSN, CACNA1C, CAMK2D, DVL1, ERC2, KIF5C, NEURL1, NRN1, PRKN, RIMS2, SHC2, UNC13A | 13 |
| **Cell Death and Survival,Cellular Compromise,Neurological Disease,Organismal Injury and Abnormalities,Tissue Morphology** | Neurodegeneration of hippocampal neurons | 0.0255 | GRIA2, PRKN | 2 |
| **Nervous System Development and Function** | Dedepression of CA1 neuron | 0.0293 | GRIA2 | 1 |
| **Cell-To-Cell Signaling and Interaction,Nervous System Development and Function** | Short-term depression of synapse | 0.0293 | UNC13A | 1 |
| **Cellular Assembly and Organization,Cellular Function and Maintenance,Cellular Movement,Nervous System Development and Function** | Arrest in axonal transport of mitochondria | 0.0293 | PRKN | 1 |
| **Cell-To-Cell Signaling and Interaction,Nervous System Development and Function** | Action potential of spinal neuron | 0.0293 | HTR1A | 1 |
| **Nervous System Development and Function** | Depotentiation of CA1 neuron | 0.0293 | GRIA2 | 1 |
| **Cell-To-Cell Signaling and Interaction,Nervous System Development and Function** | Neurotransmission of thalamocortical axons | 0.0293 | RIMS2 | 1 |
| **Cell Morphology,Cell-To-Cell Signaling and Interaction,Cellular Assembly and Organization** | Structural integrity of synapse | 0.0293 | SNTA1 | 1 |
| **Cardiovascular System Development and Function** | Hypoperfusion of brain | 0.0293 | NOS2 | 1 |
| **Cell Death and Survival,Cell Morphology,Nervous System Development and Function,Neurological Disease,Tissue Morphology** | Loss of basal ganglia neuron | 0.0293 | VPS13A | 1 |
| **Nervous System Development and Function,Tissue Morphology** | Quantity of catecholaminergic neurons | 0.0293 | PRKN | 1 |
| **Nervous System Development and Function,Neurological Disease,Organ Morphology,Organismal Development,Organismal Injury and Abnormalities** | Abnormal morphology of lobule II | 0.0293 | GABRB3 | 1 |
| **Cellular Function and Maintenance** | Respiration of neurons | 0.0293 | SLC25A23 | 1 |
| **Behavior** | Pavlovian learning | 0.0293 | GABRB3 | 1 |
| **Cell-To-Cell Signaling and Interaction** | Synaptic strength of GABAergic synapse | 0.0293 | AGRN | 1 |
| **Carbohydrate Metabolism,Lipid Metabolism,Molecular Transport,Small Molecule Biochemistry** | Concentration of phosphatidylserine | 0.0293 | PTGS2 | 1 |
| **Cellular Assembly and Organization,Cellular Function and Maintenance** | Formation of Golgi vesicles | 0.0293 | BSN | 1 |
| **Neurological Disease,Organismal Injury and Abnormalities** | Hyperinnervation of frontal cortex | 0.0293 | SORCS2 | 1 |
| **Endocrine System Development and Function,Endocrine System Disorders,Nervous System Development and Function,Neurological** | Abnormal morphology of neurohypophysis | 0.0293 | ARNT2 | 1 |

| | | | | |
|---|---|---|---|---|
| **Disease,Organ Morphology,Organismal Development,Organismal Injury and Abnormalities,Reproductive System Disease** | | | | |
| **Neurological Disease,Organismal Injury and Abnormalities,Psychological Disorders** | Damage of brain tissue | 0.0293 | NOS2 | 1 |
| **Metabolic Disease,Neurological Disease,Organismal Injury and Abnormalities,Psychological Disorders** | Stage II Alzheimer disease | 0.0293 | PTGS2 | 1 |
| **Cell Morphology,Cellular Assembly and Organization,Cellular Development,Cellular Function and Maintenance,Cellular Growth and Proliferation,Embryonic Development,Nervous System Development and Function,Organismal Development,Tissue Development** | Branching of sympathetic neurites | 0.0293 | GAP43 | 1 |
| **Nervous System Development and Function,Neurological Disease,Organ Morphology,Organismal Development,Organismal Injury and Abnormalities** | Abnormal morphology of lobule III | 0.0293 | GABRB3 | 1 |
| **Cell-To-Cell Signaling and Interaction,Cellular Assembly and Organization,Cellular Development,Cellular Function and Maintenance,Cellular Growth and Proliferation,Nervous System Development and Function,Tissue Development** | Formation of neuromuscular synapse | 0.0293 | AGRN | 1 |
| **Cardiovascular Disease,Organismal Injury and Abnormalities** | Failure of heart | 0.0293 | PTGS2 | 1 |
| **Developmental Disorder,Endocrine System Disorders,Hereditary Disorder,Metabolic Disease,Neurological Disease,Organismal Injury and Abnormalities,Psychological Disorders** | Childhood adrenoleukodystrophy | 0.0293 | PTGS2 | 1 |
| **Nervous System Development and Function,Organ Morphology,Tissue Morphology** | Quantity of spiny stellate neurons | 0.0293 | RIMS2 | 1 |
| **Cellular Assembly and Organization** | Quantity of dense-core vesicles | 0.0293 | BSN | 1 |
| **Endocrine System Development and Function,Endocrine System Disorders,Nervous System Development and Function,Neurological Disease,Organ Development,Organ Morphology,Organismal Development,Organismal Injury and Abnormalities** | Abnormal morphology of supraoptic nucleus | 0.0293 | ARNT2 | 1 |
| **Cellular Assembly and Organization** | Translocation of Golgi vesicles | 0.0293 | BSN | 1 |
| **Cell-To-Cell Signaling and Interaction,Inflammatory Response** | Cytotoxic reaction of macrophages | 0.0293 | NOS2 | 1 |
| **Cellular Compromise** | Depletion of neural stem cells | 0.0293 | PRDM16 | 1 |
| **Cell Morphology,Cellular Assembly and Organization,Cellular Function and Maintenance,Nervous System Development and Function** | Extension of axons | 0.0316 | DBN1, DVL1, MYO5B, SEMA4F, SEMA5B | 5 |
| **Cell Morphology,Cellular Assembly and Organization,Cellular Development,Cellular Function and Maintenance,Cellular Growth and Proliferation,Embryonic Development,Nervous System Development and Function,Organismal Development,Tissue Development,Tissue Morphology** | Length of dendritic spines | 0.0346 | DBN1, SIPA1L1 | 2 |
| **Cell-To-Cell Signaling and Interaction,Nervous System Development and Function** | Synaptic transmission of cells | 0.037 | BSN, ERC2, GRIA2, HTR1A | 4 |
| **Behavior,Nervous System Development and Function** | Spatial memory | 0.0379 | ADCY1, KCNH3 | 2 |
| **Cellular Function and Maintenance,Nervous System Development and Function,Tissue Development** | Function of sensory neurons | 0.0413 | ADGRV1, SLC24A4 | 2 |
| **Cell-To-Cell Signaling and Interaction,Drug Metabolism,Molecular Transport,Small Molecule Biochemistry** | Concentration of norepinephrine | 0.0413 | GAP43, PRKN | 2 |
| **Cellular Movement,Nervous System Development and Function** | Guidance of axons | 0.0413 | DVL1, GAP43, KIF5C, | 7 |

| | | | | |
|---|---|---|---|---|
| | | | PIK3R1, PTPRO, SEMA4F, UNC5D | |
| **Cellular Function and Maintenance,Nervous System Development and Function,Tissue Development** | Function of neurons | 0.0414 | ADGRV1, LRRTM4, SLC24A4 | 3 |
| **Nervous System Development and Function,Tissue Morphology** | Quantity of sensory neurons | 0.0434 | DVL1, KIF5C, SHC2 | 3 |
| **Behavior,Nervous System Development and Function** | Memory | 0.0434 | ADCY1, KCNH3, NEURL1 | 3 |
| **Cell Signaling,Molecular Transport,Vitamin and Mineral Metabolism** | Uptake of Ca2+ | 0.0437 | GRIA2 | 1 |
| **Cellular Assembly and Organization** | Fusion of mitochondria | 0.0437 | PRKN | 1 |
| **Nervous System Development and Function,Neurological Disease,Organ Morphology,Organismal Development,Organismal Injury and Abnormalities** | Lack of barrel cortex | 0.0437 | ADCY1 | 1 |
| **Cancer,Endocrine System Disorders,Organismal Injury and Abnormalities,Reproductive System Disease** | Development of pituitary gland tumor | 0.0437 | CUL9 | 1 |
| **Endocrine System Development and Function,Endocrine System Disorders,Nervous System Development and Function,Neurological Disease,Organ Development,Organ Morphology,Organismal Development,Organismal Injury and Abnormalities** | Abnormal morphology of paraventricular nucleus | 0.0437 | ARNT2 | 1 |
| **Nervous System Development and Function** | Neuroprotection of CA1 neuron | 0.0437 | PTGS2 | 1 |
| **Nervous System Development and Function** | Activation of superior colliculus | 0.0437 | ADCY1 | 1 |
| **Behavior** | Motor learning | 0.0437 | ADGRB3 | 1 |
| **Developmental Disorder,Neurological Disease** | Hypoplasia of vestibular ganglion | 0.0437 | GABRB3 | 1 |
| **Cell Morphology,Cellular Assembly and Organization** | Elongation of mitochondria | 0.0437 | PRKN | 1 |
| **Cell Death and Survival,Nervous System Development and Function** | Survival of hypoglossal motor neurons | 0.0437 | AGRN | 1 |
| **Cellular Assembly and Organization,Nervous System Development and Function** | Turnover of synaptic vesicles | 0.0437 | AGRN | 1 |
| **Cell-To-Cell Signaling and Interaction,Molecular Transport,Small Molecule Biochemistry** | Release of 5-hydroxytryptamine | 0.0437 | HTR1A | 1 |
| **Embryonic Development,Nervous System Development and Function,Organ Development,Organismal Development,Tissue Development** | Formation of optic tract | 0.0437 | GAP43 | 1 |
| **Cell-To-Cell Signaling and Interaction,Cellular Assembly and Organization,Cellular Development,Cellular Function and Maintenance,Cellular Growth and Proliferation,Nervous System Development and Function,Tissue Development** | Formation of ribbon synapse | 0.0437 | BSN | 1 |
| **Amino Acid Metabolism,Small Molecule Biochemistry** | Removal of L-glutamic acid | 0.0437 | GRIA2 | 1 |
| **Cell Death and Survival,Nervous System Development and Function** | Survival of facial branchiomotor neurons | 0.0437 | AGRN | 1 |
| **Nervous System Development and Function** | Olfactory response | 0.0437 | SLC24A4 | 1 |
| **Hereditary Disorder,Neurological Disease,Organismal Injury and Abnormalities,Psychological Disorders,Skeletal and Muscular Disorders** | Huntington disease grade 4 Huntington disease | 0.0437 | RPH3A | 1 |
| **Cell Death and Survival,Nervous System Development and Function** | Survival of trochlear motor neurons | 0.0437 | AGRN | 1 |
| **Cell Death and Survival,Nervous System Development and Function** | Survival of trigeminal motor neurons | 0.0437 | AGRN | 1 |
| **Cell Morphology,Cellular Assembly and Organization,Nervous System Development and Function** | Elongation of dendrites | 0.0437 | AGRN | 1 |

| | | | | |
|---|---|---|---|---|
| **Cell-To-Cell Signaling and Interaction,Cellular Growth and Proliferation,Nervous System Development and Function** | Excitation of striatal neurons | 0.0437 | PRKN | 1 |
| **Cell Death and Survival,Cellular Compromise,Neurological Disease,Organismal Injury and Abnormalities,Tissue Morphology** | Neurodegeneration of CA1 neuron | 0.0437 | PTGS2 | 1 |
| **Cell-To-Cell Signaling and Interaction,Cellular Growth and Proliferation** | Excitation of cerebral cortex cells | 0.0448 | KCNH3, NRN1 | 2 |
| **Auditory Disease,Cell Death and Survival,Cellular Compromise,Neurological Disease,Organismal Injury and Abnormalities,Tissue Morphology** | Neurodegeneration of inner hair cells | 0.0448 | ADGRV1, GABRB3 | 2 |
| **Nervous System Development and Function** | Neuroprotection of cerebral cortex cells | 0.0448 | LRP1B, PTGS2 | 2 |
| **Cell Morphology,Cellular Movement,Nervous System Development and Function** | Innervation of neurons | 0.0473 | GABRB3, GAP43, GRIA2, RIMS2 | 4 |
| **Cell-To-Cell Signaling and Interaction** | Quantity of neurotransmitter | 0.0473 | GAP43, HTR1A, PRKN, SORCS2 | 4 |
| **Cellular Assembly and Organization,Nervous System Development and Function** | Recycling of synaptic vesicles | 0.0484 | DVL1, GAP43 | 2 |
| **Nervous System Development and Function,Neurological Disease,Organ Morphology,Organismal Development,Organismal Injury and Abnormalities** | Abnormal morphology of cerebral cortex | 0.0486 | ADCY1, CRMP1, DBN1, E2F5, GAP43, GRIA2, LRRTM4 | 7 |
| **Cell-To-Cell Signaling and Interaction,Molecular Transport,Small Molecule Biochemistry** | Quantity of monoamines | 0.0488 | GAP43, HTR1A, PRKN, SORCS2 | 4 |
| **Nervous System Development and Function,Neurological Disease,Organ Morphology,Organismal Development,Organismal Injury and Abnormalities** | Abnormal morphology of diencephalon | 0.0488 | ADCY1, ARNT2, CELSR2, E2F5 | 4 |
| **Cellular Assembly and Organization** | Quantity of vesicles | 0.0497 | BSN, ERC2, UNC13A | 3 |

### Supplementary Table 10: Module 10
### Cluster 10

| Categories | Diseases or Functions Annotation | p-Value | Molecules | # Molecules |
|---|---|---|---|---|
| **Cellular Function and Maintenance,Molecular Transport** | Exocytosis | 4.44E-07 | APBA2, CADPS2, ERC2, PIP5K1C, PPFIA3, RIMS2, RPH3A, SNAP25, UNC13A, UNC13B, UNC13C | 11 |
| **Cellular Assembly and Organization** | Priming of vesicles | 0.00000656 | CADPS2, ERC2, RIMS2, SNAP25, UNC13A, UNC13B, UNC13C | 7 |
| **Cellular Assembly and Organization,Cellular Function and Maintenance,Cellular Movement,Molecular Transport,Nervous System Development and Function** | Exocytosis of synaptic vesicles | 0.00000723 | APBA2, PIP5K1C, PPFIA3, RPH3A, SNAP25, UNC13A, UNC13B, UNC13C | 8 |

| | | | | |
|---|---|---|---|---|
| **Cellular Assembly and Organization,Cellular Function and Maintenance** | Microtubule dynamics | 0.0000308 | ABI1, ADCY1, ADCY6, ADGRB3, AGRN, AHI1, ANAPC2, ARHGEF28, ATCAY, BCL11B, BSN, CAMSAP1, CDKL3, CELSR2, CRMP1, DAB2IP, DBN1, DVL1, ELK1, EPHB3, ERC2, FMN1, GABRB3, GAP43, GPRIN1, HERC1, HTR1A, LAMB1, LRRK2, MEF2A, MRTFA, MYO16, MYO5B, NIN, NRN1, PIP5K1C, PTGS2, PTK2B, RBFOX2, ROCK2, RYR2, SCN1B, SEMA4F, SEMA5B, SHANK2, SIPA1L1, SMN1/SMN2, SORBS2, SPTBN4, SYNE1, TIAM1, TOP2B, TSC2 | 53 |
| **Cell-To-Cell Signaling and Interaction,Cellular Assembly and Organization,Cellular Function and Maintenance,Molecular Transport** | Priming of synaptic vesicles | 0.0000338 | CADPS2, ERC2, RIMS2, UNC13A, UNC13B, UNC13C | 6 |
| **Nervous System Development and Function,Neurological Disease** | Abnormal morphology of nervous system | 0.0000354 | ABR, ADCY1, ADGRA2, AGRN, AHI1, AKT3, ARNT2, ASPM, BRCA2, CACNA1C, CADPS2, CELSR2, COL13A1, CREBBP, CRMP1, DBN1, DVL1, EPHB3, ERC2, FAT3, FES, GABRB3, | 58 |

| | | | | |
|---|---|---|---|---|
| | | | GAP43, GRIA2, HIVEP2, IDUA, KIF1B, LRRK2, LRRTM4, MAN2C1, MCPH1, MEGF8, MTHFR, MYO16, NCOR1, NOS2, NRN1, PER2, PRKN, PROKR2, RIMS2, ROCK2, SCN1B, SEMA5B, SHANK2, SMN1/SMN2, SNAP25, SORBS2, SORL1, SPTBN4, SYNE1, TCF7L2, TENM2, TOP2B, UNC13A, UNC13B, VAX1, VPS13A | |
| **Cell Morphology,Cellular Assembly and Organization,Cellular Development,Cellular Function and Maintenance,Cellular Growth and Proliferation,Nervous System Development and Function,Organismal Development,Tissue Development** | Neuritogenesis | 0.0000517 | ABI1, ADCY1, ADGRB3, AGRN, ANAPC2, ARHGEF28, ATCAY, BCL11B, BSN, CAMSAP1, CDKL3, CELSR2, CRMP1, DAB2IP, DBN1, DVL1, ELK1, EPHB3, ERC2, FMN1, GABRB3, GAP43, GPRIN1, HERC1, LAMB1, LRRK2, MEF2A, MRTFA, MYO16, MYO5B, NIN, NRN1, PIP5K1C, PTGS2, PTK2B, RBFOX2, ROCK2, RYR2, SHANK2, SIPA1L1, SMN1/SMN2, SORBS2, | 47 |

| | | | | |
|---|---|---|---|---|
| | | | SPTBN4, SYNE1, TIAM1, TOP2B, TSC2 | |
| **Cellular Assembly and Organization,Nervous System Development and Function** | Docking of synaptic vesicles | 0.0000617 | PPFIA3, RIMS2, SNAP25, UNC13A, UNC13B | 5 |
| **Cellular Assembly and Organization,Cellular Movement,Nervous System Development and Function** | Transport of synaptic vesicles | 0.0000711 | APBA2, KIAA1107, PCLO, PIP5K1C, PPFIA3, RPH3A, SCRIB, SNAP25, UNC13A, UNC13B, UNC13C | 11 |
| **Nervous System Development and Function** | Morphology of nervous system | 0.00011 | ABR, ADCY1, ADGRA2, AGRN, AHI1, AKT3, ARNT2, ASPM, BRCA2, CACNA1C, CADPS2, CELSR2, COL13A1, CREBBP, CRMP1, DBN1, DVL1, EPHB3, ERC2, FAT3, FES, GABRB3, GAP43, GRIA2, HIVEP2, IDUA, KIF1B, KIF5C, KNG1, LRRK2, LRRTM4, MAN2C1, MCPH1, MEGF8, MTHFR, MYO16, NCOR1, NOS2, NRN1, PER2, PLAGL1, PRDM16, PRKN, PROKR2, RGS11, RIMS2, ROCK2, SCN1B, SEMA5B, SHANK2, SIPA1L1, SMN1/SMN2, SNAP25, SORBS2, SORL1, SPTBN4, SYNE1, TCF7L2, | 65 |

| | | | | |
|---|---|---|---|---|
| | | | TENM2, TIAM1, TOP2B, UNC13A, UNC13B, VAX1, VPS13A | |
| **Cellular Development,Cellular Growth and Proliferation,Nervous System Development and Function,Tissue Development** | Development of neurons | 0.000117 | ABI1, ADCY1, ADGRB3, AGRN, ANAPC2, ARHGEF28, ATCAY, BCL11B, BSN, CADPS2, CAMSAP1, CDKL3, CELSR2, CRMP1, DAB2IP, DBN1, DVL1, ELK1, ELP1, EPHB3, ERC2, FMN1, GABRB3, GAP43, GPRIN1, HERC1, LAMB1, LRRK2, LRRTM4, MEF2A, MRTFA, MYO16, MYO5B, NIN, NRN1, PCDH8, PIP5K1C, PLXND1, PRDM1, PTGS2, PTK2B, RBFOX2, ROCK2, RYR2, SHANK2, SIPA1L1, SMN1/SMN2, SORBS2, SPTBN4, SYNE1, TIAM1, TOP2B, TSC2 | 53 |
| **Cancer,Neurological Disease,Organismal Injury and Abnormalities** | Brain astrocytoma | 0.000305 | ABCF1, ADAMTS13, ADAMTS8, AGAP3, ANKRD30B, ASPM, ASXL3, ATF7IP2, CREBBP, DNAH8, DPP10, ERC2, GATAD2B, HERC1, ICE1, MYO5B, P2RX5, PCLO, | 32 |

| | | | | |
|---|---|---|---|---|
| | | | PIK3R1, PRDM2, PROKR2, PTGS2, RAB11FIP5, RALY, RIMS2, ROS1, RYR2, SEMA4F, SPTBN4, SYNE1, THEMIS, TSC2 | |
| **Nervous System Development and Function,Tissue Morphology** | Density of neurons | 0.000356 | ACVR1C, CRMP1, DAB2IP, ELK1, EPHB3, GABRB3, GAP43, LRRTM4, MEF2A, PTGS2, RYR2, SCN1B, TIAM1, TSC2 | 14 |
| **Organismal Development,Organismal Injury and Abnormalities** | Abnormal morphology of head | 0.000411 | ABR, ADCY1, ADGRA2, ADGRV1, AHI1, AKT3, ARNT2, ASPM, CADPS2, CELSR2, CREBBP, CRMP1, DBN1, DVL1, EPHB3, FAT3, FES, GABRB3, GAP43, GRIA2, HIVEP2, IDUA, KIF1B, LRRTM4, MAN2C1, MCPH1, MTHFR, NCOR1, NOS2, PRKN, PROKR2, SHANK2, SNAP25, SPTBN4, TCF7L2, TENM2, TOP2B, VAX1 | 38 |
| **Nervous System Development and Function,Neurological Disease,Organ Morphology,Organismal Development,Organismal Injury and Abnormalities** | Abnormal morphology of brain | 0.000464 | ABR, ADCY1, ADGRA2, AHI1, AKT3, ARNT2, ASPM, CADPS2, CELSR2, CREBBP, CRMP1, DBN1, EPHB3, FES, GABRB3, GAP43, | 35 |

| | | | | |
|---|---|---|---|---|
| | | | GRIA2, HIVEP2, IDUA, KIF1B, LRRTM4, MAN2C1, MCPH1, MTHFR, NCOR1, NOS2, PRKN, PROKR2, SHANK2, SNAP25, SPTBN4, TCF7L2, TENM2, TOP2B, VAX1 | |
| **Nervous System Development and Function,Neurological Disease** | Abnormal morphology of central nervous system | 0.000489 | ABR, ADCY1, ADGRA2, AHI1, AKT3, ARNT2, ASPM, CADPS2, CELSR2, CREBBP, CRMP1, DBN1, EPHB3, FES, GABRB3, GAP43, GRIA2, HIVEP2, IDUA, KIF1B, LRRTM4, MAN2C1, MCPH1, MTHFR, NCOR1, NOS2, PRKN, PROKR2, RIMS2, SCN1B, SHANK2, SNAP25, SPTBN4, TCF7L2, TENM2, TOP2B, VAX1 | 37 |
| **Cancer,Organismal Injury and Abnormalities** | Head and neck tumor | 0.000779 | ABCF1, ADAMTS13, ADAMTS8, AGAP3, ANKRD30B, ASPM, ASXL3, ATF7IP2, CREBBP, CUL9, DNAH8, DPP10, ERC2, GATAD2B, HERC1, ICE1, MYO15A, MYO5B, P2RX5, PCLO, PIK3R1, PRDM1, PRDM2, | 38 |

| | | | | |
|---|---|---|---|---|
| | | | PROKR2, PTGS2, PTPRT, RAB11FIP5, RALY, RIMS2, ROS1, RYR2, SEMA4F, SETD2, SPTBN4, SYNE1, THEMIS, TSC2, UNC13B | |
| **Nervous System Development and Function** | Morphology of central nervous system | 0.000873 | ABR, ADCY1, ADGRA2, AHI1, AKT3, ARNT2, ASPM, CADPS2, CELSR2, CREBBP, CRMP1, DBN1, EPHB3, FES, GABRB3, GAP43, GRIA2, HIVEP2, IDUA, KIF1B, KIF5C, KNG1, LRRTM4, MAN2C1, MCPH1, MTHFR, MYO16, NCOR1, NOS2, PRDM16, PRKN, PROKR2, RIMS2, SCN1B, SHANK2, SMN1/SMN2, SNAP25, SPTBN4, TCF7L2, TENM2, TOP2B, VAX1 | 42 |
| **Cancer,Organismal Injury and Abnormalities** | Non-hematological solid tumor | 0.000887 | ABCF1, ADAMTS13, ADAMTS8, AGAP3, ANKRD30B, ASPM, ASXL3, ATF7IP2, CREBBP, CUL9, DNAH8, DPP10, ERC2, GATAD2B, HERC1, ICE1, MYO15A, MYO5B, P2RX5, PCLO, PIGQ, PIK3R1, | 39 |

| | | | | |
|---|---|---|---|---|
| | | | PRDM1, PRDM2, PROKR2, PTGS2, PTPRT, RAB11FIP5, RALY, RIMS2, ROS1, RYR2, SEMA4F, SETD2, SPTBN4, SYNE1, THEMIS, TSC2, UNC13B | |
| **Cancer,Neurological Disease,Organismal Injury and Abnormalities** | Brain glioma | 0.00093 | ABCF1, ADAMTS13, ADAMTS8, AGAP3, ANKRD30B, ASPM, ASXL3, ATF7IP2, CREBBP, DNAH8, DPP10, ERC2, GATAD2B, HERC1, ICE1, MYO15A, MYO5B, P2RX5, PCLO, PIK3R1, PRDM1, PRDM2, PROKR2, PTGS2, PTPRT, RAB11FIP5, RALY, RIMS2, ROS1, RYR2, SEMA4F, SETD2, SPTBN4, SYNE1, THEMIS, TSC2 | 36 |
| **Cellular Assembly and Organization,Cellular Function and Maintenance** | Formation of Golgi vesicles | 0.0011 | BSN, PCLO | 2 |
| **Cellular Assembly and Organization** | Translocation of Golgi vesicles | 0.0011 | BSN, PCLO | 2 |
| **Organismal Development** | Morphology of head | 0.00115 | ABR, ADCY1, ADGRA2, ADGRV1, AHI1, AKT3, ARNT2, ASPM, CADPS2, CELSR2, CREBBP, CRMP1, DBN1, DVL1, EPHB3, FAT3, FES, GABRB3, GAP43, GRIA2, HIVEP2, | 42 |

| | | | | |
|---|---|---|---|---|
| | | | IDUA, KIF1B, KIF5C, KNG1, LRRTM4, MAN2C1, MCPH1, MTHFR, MYO16, NCOR1, NOS2, PRDM16, PRKN, PROKR2, SHANK2, SNAP25, SPTBN4, TCF7L2, TENM2, TOP2B, VAX1 | |
| **Cancer,Organismal Injury and Abnormalities** | Non-melanoma solid tumor | 0.00116 | ABCF1, ADAMTS13, ADAMTS8, AGAP3, ANKRD30B, ASPM, ASXL3, ATF7IP2, CREBBP, DNAH8, DPP10, ERC2, GATAD2B, HERC1, ICE1, MYO15A, MYO5B, P2RX5, PCLO, PIK3R1, PRDM1, PRDM2, PROKR2, PTGS2, PTPRT, RAB11FIP5, RALY, RIMS2, ROS1, RYR2, SEMA4F, SETD2, SPTBN4, SYNE1, THEMIS, TSC2, UNC13B | 37 |
| **Cancer,Neurological Disease,Organismal Injury and Abnormalities** | Brain tumor | 0.00119 | ABCF1, ADAMTS13, ADAMTS8, AGAP3, ANKRD30B, ASPM, ASXL3, ATF7IP2, CREBBP, DNAH8, DPP10, ERC2, GATAD2B, HERC1, ICE1, MYO15A, MYO5B, P2RX5, | 37 |

| | | | | |
|---|---|---|---|---|
| | | | PCLO, PIK3R1, PRDM1, PRDM2, PROKR2, PTGS2, PTPRT, RAB11FIP5, RALY, RIMS2, ROS1, RYR2, SEMA4F, SETD2, SPTBN4, SYNE1, THEMIS, TSC2, UNC13B | |
| **Nervous System Development and Function,Organ Morphology,Organismal Development** | Morphology of brain | 0.00122 | ABR, ADCY1, ADGRA2, AHI1, AKT3, ARNT2, ASPM, CADPS2, CELSR2, CREBBP, CRMP1, DBN1, EPHB3, FES, GABRB3, GAP43, GRIA2, HIVEP2, IDUA, KIF1B, KIF5C, KNG1, LRRTM4, MAN2C1, MCPH1, MTHFR, MYO16, NCOR1, NOS2, PRDM16, PRKN, PROKR2, SHANK2, SNAP25, SPTBN4, TCF7L2, TENM2, TOP2B, VAX1 | 39 |
| **Cancer,Neurological Disease,Organismal Injury and Abnormalities** | Nervous system neoplasm | 0.00145 | ABCF1, ADAMTS13, ADAMTS8, AGAP3, ANKRD30B, ASPM, ASXL3, ATF7IP2, CREBBP, DNAH8, DPP10, ERC2, GATAD2B, HERC1, ICE1, MYO15A, MYO5B, P2RX5, PCLO, PIGQ, PIK3R1, | 38 |

| | | | | |
|---|---|---|---|---|
| | | | PRDM1, PRDM2, PROKR2, PTGS2, PTPRT, RAB11FIP5, RALY, RIMS2, ROS1, RYR2, SEMA4F, SETD2, SPTBN4, SYNE1, THEMIS, TSC2, UNC13B | |
| **Cell Morphology,Cell-To-Cell Signaling and Interaction,Cellular Assembly and Organization,Nervous System Development and Function,Tissue Morphology** | Morphology of synapse | 0.00164 | AGRN, COL13A1, ERC2, GAP43, NRN1, ROCK2, SMN1/SMN2, SYNE1, UNC13A, UNC13B | 10 |
| **Cell-To-Cell Signaling and Interaction,Nervous System Development and Function** | Action potential of motor endplates | 0.0018 | COL13A1, RIMS2, SNAP25 | 3 |
| **Behavior,Nervous System Development and Function** | Short-term memory | 0.0018 | CREBBP, KCNH3, NR3C2 | 3 |
| **Lipid Metabolism,Molecular Transport,Small Molecule Biochemistry** | Concentration of arachidonic acid | 0.00263 | ARNTL, KNG1, PTGS2 | 3 |
| **Cell Morphology,Cell-To-Cell Signaling and Interaction,Cellular Assembly and Organization,Nervous System Development and Function,Neurological Disease,Tissue Morphology** | Abnormal morphology of synapse | 0.00274 | AGRN, COL13A1, ERC2, NRN1, ROCK2, SMN1/SMN2, SYNE1, UNC13A, UNC13B | 9 |
| **Cellular Assembly and Organization,Nervous System Development and Function,Tissue Morphology** | Abnormal quantity of synaptic vesicles | 0.00283 | ERC2, KIF1B, PCLO, UNC13A, UNC13B | 5 |
| **Hereditary Disorder,Neurological Disease,Organismal Injury and Abnormalities,Psychological Disorders,Skeletal and Muscular Disorders** | Huntington disease grade 4 Huntington disease | 0.00323 | RPH3A, SNAP25 | 2 |
| **Hereditary Disorder,Neurological Disease,Organismal Injury and Abnormalities,Psychological Disorders,Skeletal and Muscular Disorders** | Familial Parkinson disease | 0.00323 | GAP43, PRKN | 2 |
| **Cell Morphology,Cellular Assembly and Organization,Cellular Development,Cellular Function and Maintenance,Cellular Growth and Proliferation,Embryonic Development,Nervous System Development and Function,Organismal Development,Tissue Development** | Dendritic growth/branching | 0.00334 | ABI1, AGRN, BSN, CRMP1, DAB2IP, DBN1, ELK1, EPHB3, GABRB3, GAP43, MYO5B, NRN1, PTGS2, ROCK2, RYR2, SHANK2, SIPA1L1, SORBS2, TIAM1, TSC2 | 20 |

| | | | | |
|---|---|---|---|---|
| **Neurological Disease,Organismal Injury and Abnormalities** | Brain lesion | 0.00376 | ABCF1, ADAMTS13, ADAMTS8, AGAP3, ANKRD30B, ASPM, ASXL3, ATF7IP2, CREBBP, DNAH8, DPP10, ERC2, GATAD2B, HERC1, ICE1, KNG1, MYO15A, MYO5B, P2RX5, PCLO, PIK3R1, PRDM1, PRDM2, PROKR2, PTGS2, PTPRT, RAB11FIP5, RALY, RIMS2, ROS1, RYR2, SEMA4F, SETD2, SPTBN4, SYNE1, THEMIS, TSC2, UNC13B | 38 |
| **Neurological Disease,Organismal Injury and Abnormalities** | Hydrocephalus | 0.00381 | CELSR2, DVL1, DVL3, NCOR1 | 4 |
| **Neurological Disease,Psychological Disorders** | Pervasive developmental disorder | 0.00458 | MATK, RYR2, SLC6A7, TSC2 | 4 |
| **Cell Morphology,Cellular Assembly and Organization,Cellular Development,Cellular Function and Maintenance,Cellular Growth and Proliferation,Nervous System Development and Function,Organismal Development,Tissue Development** | Formation of dendrites | 0.00536 | ABI1, ADGRB3, ANAPC2, CDKL3, CELSR2, DVL1, FMN1, MEF2A, MYO5B, RBFOX2, ROCK2, SIPA1L1, SYNE1 | 13 |
| **Behavior,Nervous System Development and Function** | Working memory | 0.00632 | KCNH3, NR3C2 | 2 |
| **Nervous System Development and Function,Neurological Disease,Ophthalmic Disease,Organ Morphology,Organismal Development,Organismal Injury and Abnormalities** | Abnormal morphology of visual cortex | 0.00632 | HIVEP2, TENM2 | 2 |
| **Auditory and Vestibular System Development and Function,Cell Morphology,Cellular Assembly and Organization** | Orientation of stereocilia bundles | 0.0064 | ADGRV1, DVL1, DVL3 | 3 |
| **Cell-To-Cell Signaling and Interaction,Cellular Assembly and Organization** | Cell-cell contact | 0.00671 | AGRN, BSN, COL13A1, EPHB3, ERC2, GAP43, GJD2, LRRTM4, NEURL1, NRN1, | 19 |

| | | | | |
|---|---|---|---|---|
| | | | PCDH8, PLXND1, ROCK2, SHANK2, SMN1/SMN2, SORBS2, SYNE1, UNC13A, UNC13B | |
| **Cell Morphology,Nervous System Development and Function,Tissue Morphology** | Morphology of bipolar cells | 0.00753 | ERC2, FAT3, RGS11, SEMA5B | 4 |
| **Nervous System Development and Function,Neurological Disease,Organ Morphology,Organismal Development,Organismal Injury and Abnormalities** | Abnormal morphology of barrel cortex | 0.00812 | ADCY1, ASPM, GAP43 | 3 |
| **Cell Morphology,Nervous System Development and Function,Neurological Disease,Tissue Morphology** | Abnormal morphology of neurons | 0.00916 | AGRN, BRCA2, CACNA1C, CADPS2, COL13A1, CRMP1, DBN1, DVL1, EPHB3, ERC2, FAT3, GAP43, HIVEP2, IDUA, LRRK2, MYO16, NOS2, NRN1, PRKN, ROCK2, SCN1B, SEMA5B, SHANK2, SMN1/SMN2, SORBS2, SORL1, SPTBN4, SYNE1, UNC13A, UNC13B, VPS13A | 31 |
| **Cell Morphology,Cellular Development,Embryonic Development,Nervous System Development and Function,Tissue Development** | Patterning of dendrites | 0.0103 | CELSR2, CRMP1 | 2 |
| **Cell Morphology,Cellular Function and Maintenance** | Autophagy of cortical neurons | 0.0103 | PRKN, TSC2 | 2 |
| **Cell Morphology,Nervous System Development and Function** | Innervation of optic nerve | 0.0103 | GAP43, VAX1 | 2 |
| **Cellular Development,Nervous System Development and Function,Tissue Development** | Differentiation of nerve ending | 0.0103 | AGRN, MEF2A | 2 |
| **Hereditary Disorder,Neurological Disease,Organismal Injury and Abnormalities,Psychological Disorders,Skeletal and Muscular Disorders** | Huntington disease grade 3 Huntington disease | 0.0103 | RPH3A, SNAP25 | 2 |
| **Cellular Assembly and Organization** | Quantity of vesicles | 0.0104 | BSN, ERC2, KIF1B, PCLO, UNC13A, UNC13B | 6 |
| **Cellular Assembly and Organization,Cellular Compromise** | Retraction of cellular protrusions | 0.0104 | ARHGEF11, DVL1, GRIA2, LRRK2, PITPNM1, ROCK2 | 6 |
| **Cell Morphology** | Shape change of neurites | 0.014 | ABI1, AGRN, BCL11B, BSN, CRMP1, | 22 |

| | | | | |
|---|---|---|---|---|
| | | | DAB2IP, DBN1, ELK1, EPHB3, GABRB3, GAP43, MYO5B, NR3C2, NRN1, PTGS2, ROCK2, RYR2, SHANK2, SIPA1L1, SORBS2, TIAM1, TSC2 | |
| **Cell-To-Cell Signaling and Interaction,Nervous System Development and Function** | Long term synaptic depression of CA1 neuron | 0.0148 | GRIA2, PIP5K1C, RAB11FIP5 | 3 |
| **Nervous System Development and Function** | Depotentiation | 0.0151 | ADCY1, GRIA2 | 2 |
| **Amino Acid Metabolism,Endocrine System Development and Function,Molecular Transport,Small Molecule Biochemistry** | Quantity of L-triiodothyronine | 0.0151 | NCOR1, SLCO1C1 | 2 |
| **Embryonic Development,Organismal Development** | Development of head | 0.0155 | ABR, ADCY1, ADGRA2, AHI1, AKT3, ARNT2, ASPM, BRCA2, CDK5RAP1, CELSR2, ELP1, EPHB3, FAT3, GAP43, GRHL2, LRRK2, MCPH1, MRTFA, MYO16, NCOR1, NIN, NME7, NOS2, PITPNM1, PROKR2, RBFOX2, RTP1, SCRIB, SETD2, TACC2, TCF7L2, TOP2B, VAX1 | 33 |
| **Nervous System Development and Function,Tissue Morphology** | Morphology of nervous tissue | 0.0164 | AGRN, BRCA2, CACNA1C, CADPS2, COL13A1, CRMP1, DBN1, DVL1, EPHB3, ERC2, FAT3, GAP43, HIVEP2, IDUA, LRRK2, MAN2C1, MYO16, NOS2, NRN1, PLAGL1, PRKN, RGS11, RIMS2, | 37 |

| | | | | |
|---|---|---|---|---|
| | | | ROCK2, SCN1B, SEMA5B, SHANK2, SIPA1L1, SMN1/SMN2, SORBS2, SORL1, SPTBN4, SYNE1, TIAM1, UNC13A, UNC13B, VPS13A | |
| **Cell Morphology,Nervous System Development and Function,Tissue Morphology** | Morphology of neurons | 0.0188 | AGRN, BRCA2, CACNA1C, CADPS2, COL13A1, CRMP1, DBN1, DVL1, EPHB3, ERC2, FAT3, GAP43, HIVEP2, IDUA, LRRK2, MYO16, NOS2, NRN1, PLAGL1, PRKN, RGS11, RIMS2, ROCK2, SCN1B, SEMA5B, SHANK2, SIPA1L1, SMN1/SMN2, SORBS2, SORL1, SPTBN4, SYNE1, TIAM1, UNC13A, UNC13B, VPS13A | 36 |
| **Cell Morphology,Cell-To-Cell Signaling and Interaction,Nervous System Development and Function** | Plasticity of synapse | 0.0191 | CAMK2G, DBN1, GRIA2, KCNH3, NEURL1, PCDH8, PIP5K1C, PPFIA3, PTK2B, SIPA1L1 | 10 |
| **Lipid Metabolism,Small Molecule Biochemistry** | Synthesis of eicosanoid | 0.0205 | KNG1, NOS2, PTGS2 | 3 |
| **Cellular Compromise,Neurological Disease,Organismal Injury and Abnormalities** | Neurodegeneration of cerebral cortex cells | 0.0207 | GRIA2, LRRK2, PRKN, PTGS2 | 4 |
| **Cell-To-Cell Signaling and Interaction,Molecular Transport,Small Molecule Biochemistry** | Concentration of 5-hydroxytryptamine | 0.0207 | APBA2, DBH, GAP43, HTR1A | 4 |
| **Cell Signaling,Nucleic Acid Metabolism,Small Molecule Biochemistry** | Binding of guanosine 5'-O-(3-thiotriphosphate) | 0.0207 | HTR1A, KNG1 | 2 |
| **Cellular Assembly and Organization** | Clustering of synaptic vesicles | 0.0207 | BSN, PCLO | 2 |

| | | | | |
|---|---|---|---|---|
| **Cell Morphology,Cellular Assembly and Organization** | Size of mitochondria | 0.0207 | AKT3, PRKN | 2 |
| **Behavior,Nervous System Development and Function** | Memory consolidation | 0.0207 | CREBBP, NR3C2 | 2 |
| **Cell-To-Cell Signaling and Interaction,Drug Metabolism,Small Molecule Biochemistry** | Metabolism of dopamine | 0.0207 | PRKN, SORCS2 | 2 |
| **Nervous System Development and Function,Organ Morphology,Organismal Development** | Morphology of forebrain | 0.0208 | ADCY1, ADGRA2, AKT3, ARNT2, ASPM, CELSR2, CREBBP, FES, GAP43, HIVEP2, MAN2C1, NCOR1, NOS2, PROKR2, TENM2, TOP2B, VAX1 | 17 |
| **Cell Morphology,Nervous System Development and Function** | Innervation | 0.0212 | EPHB3, GABRB3, GAP43, GRIA2, RIMS2, TENM2, TOP2B, VAX1 | 8 |
| **Nervous System Development and Function,Neurological Disease,Organ Morphology,Organismal Development,Organismal Injury and Abnormalities** | Abnormal morphology of forebrain | 0.0219 | ADCY1, ADGRA2, ARNT2, ASPM, CELSR2, CREBBP, FES, HIVEP2, MAN2C1, NCOR1, NOS2, PROKR2, TENM2, TOP2B, VAX1 | 15 |
| **Cell-To-Cell Signaling and Interaction,Nervous System Development and Function** | Synaptic transmission of cells | 0.0227 | BSN, ERC2, GRIA2, HTR1A, NR3C2, RAB11FIP5, SNAP25 | 7 |
| **Cell Morphology,Cellular Assembly and Organization,Cellular Development,Cellular Function and Maintenance,Cellular Growth and Proliferation,Nervous System Development and Function,Organismal Development,Tissue Development** | Branching of neurons | 0.0243 | ABI1, AGRN, BSN, CRMP1, DAB2IP, DBN1, ELK1, EPHB3, ERC2, GABRB3, GAP43, MYO5B, NRN1, PTGS2, ROCK2, RYR2, SHANK2, SIPA1L1, SORBS2, TIAM1, TSC2 | 21 |
| **Cell Morphology,Cellular Assembly and Organization,Cellular Function and Maintenance,Nervous System Development and Function** | Extension of neurites | 0.0245 | ADCY6, AHI1, ANAPC2, CDKL3, DBN1, DVL1, MYO5B, | 11 |

| | | | | |
|---|---|---|---|---|
| | | | SCN1B, SEMA4F, SEMA5B, TIAM1 | |
| **Cellular Assembly and Organization,Cellular Compromise,Nervous System Development and Function** | Retraction of neurites | 0.0246 | ARHGEF11, DVL1, LRRK2, PITPNM1, ROCK2 | 5 |
| **Cell-To-Cell Signaling and Interaction,Cellular Function and Maintenance,Nervous System Development and Function** | Paired-pulse facilitation of synapse | 0.0246 | APBA2, CADPS2, HTR1A, ROCK2, UNC13C | 5 |
| **Behavior** | Learning | 0.0255 | ADCY1, ADGRB3, CREBBP, GABRB3, KCNH3, NEURL1, NR3C2 | 7 |
| **Cell Morphology,Nervous System Development and Function,Neurological Disease,Tissue Morphology** | Abnormal morphology of bipolar cells | 0.0273 | ERC2, FAT3, SEMA5B | 3 |
| **Cell-To-Cell Signaling and Interaction,Nervous System Development and Function** | Long term synaptic depression of brain cells | 0.0279 | ADCY5, GRIA2, PIP5K1C, RAB11FIP5 | 4 |
| **Cellular Movement,Nervous System Development and Function** | Guidance of axons | 0.0283 | DVL1, EPHB3, GAP43, KIF5C, LMTK2, PIK3R1, PTPRO, RANBP9, SCN1B, SEMA4F, SPTBN4, UNC5D, VAX1 | 13 |
| **Cell Morphology,Cellular Compromise** | Collapse of growth cone | 0.0285 | ARHGEF11, EPHB3, FES, ROCK2, SEMA4F, SORCS2 | 6 |
| **Behavior,Nervous System Development and Function** | Memory | 0.0307 | ADCY1, CREBBP, KCNH3, NEURL1, NR3C2 | 5 |
| **Carbohydrate Metabolism,Drug Metabolism,Molecular Transport,Small Molecule Biochemistry** | Quantity of heparan sulfate | 0.0332 | IDUA | 1 |
| **Cell-To-Cell Signaling and Interaction,Cellular Assembly and Organization,Nervous System Development and Function** | Modulation of synapse | 0.0332 | PCDH8 | 1 |
| **Cellular Development,Nervous System Development and Function,Tissue Development** | Differentiation of sympathetic preganglionic neurons | 0.0332 | AGRN | 1 |
| **Cellular Assembly and Organization** | Aggregation of early endosomes | 0.0332 | KIF16B | 1 |
| **Cell Morphology,Cellular Assembly and Organization,Cellular Function and Maintenance** | Formation of axon-like process | 0.0332 | TIAM1 | 1 |
| **Cell Morphology,Cell-To-Cell Signaling and Interaction** | Length of stereocilia | 0.0332 | MYO15A | 1 |
| **Cell-To-Cell Signaling and Interaction,Cellular Assembly and Organization,Nervous System Development and Function,Tissue Morphology** | Quantity of ribbon synapse | 0.0332 | BSN | 1 |
| **Cell Signaling,Nucleic Acid Metabolism,Small Molecule Biochemistry** | Inhibition of cyclic AMP | 0.0332 | HTR1A | 1 |
| **Nervous System Development and Function,Tissue Morphology** | Density of serotonergic neurons | 0.0332 | GAP43 | 1 |

| | | | | |
|---|---|---|---|---|
| **Behavior,Developmental Disorder,Neurological Disease,Psychological Disorders** | Autistic-like traits | 0.0332 | TSC2 | 1 |
| **Nervous System Development and Function** | Function of autonomic nerve | 0.0332 | SNAP25 | 1 |
| **Hereditary Disorder,Neurological Disease,Organismal Injury and Abnormalities,Psychological Disorders,Skeletal and Muscular Disorders** | Autosomal recessive juvenile Parkinson disease | 0.0332 | PRKN | 1 |
| **Cellular Compromise** | Acidification of cytosol | 0.0332 | CA4 | 1 |
| **Cell-To-Cell Signaling and Interaction,Nervous System Development and Function** | Inactivation of synapse | 0.0332 | BSN | 1 |
| **Cellular Development,Nervous System Development and Function,Organ Development** | Differentiation of preoptic area | 0.0332 | VAX1 | 1 |
| **Cellular Development,Nervous System Development and Function,Organ Development** | Differentiation of medial ganglionic eminences | 0.0332 | VAX1 | 1 |
| **Cellular Development,Cellular Growth and Proliferation,Embryonic Development,Nervous System Development and Function,Organ Development,Organismal Development,Tissue Development,Visual System Development and Function** | Formation of retinal cells | 0.0332 | ADCY1 | 1 |
| **Nervous System Development and Function,Organ Morphology,Tissue Morphology** | Quantity of rod bipolar cells | 0.0332 | PRDM1 | 1 |
| **Cell Morphology,Cellular Assembly and Organization,Cellular Development,Cellular Function and Maintenance,Cellular Growth and Proliferation,Embryonic Development,Nervous System Development and Function,Organismal Development,Tissue Development** | Delay in initiation of outgrowth of dendrites | 0.0332 | DAB2IP | 1 |
| **Cell-To-Cell Signaling and Interaction,Nervous System Development and Function** | Clustering of hippocampal neurons | 0.0332 | SHANK2 | 1 |
| **Molecular Transport,Protein Trafficking** | Translocation of protein fragment | 0.0332 | AGAP3 | 1 |
| **Cell-To-Cell Signaling and Interaction,Nervous System Development and Function** | Activation of medium spiny neurons | 0.0332 | LRRK2 | 1 |
| **Nervous System Development and Function,Tissue Morphology** | Density of parallel fiber | 0.0332 | DAB2IP | 1 |
| **Cell Death and Survival,Cellular Development,Nervous System Development and Function,Tissue Development,Tissue Morphology** | Regeneration of bipolar cells | 0.0332 | PRDM1 | 1 |
| **Nervous System Development and Function,Organ Morphology,Organismal Development** | Size of optic chiasm | 0.0332 | GAP43 | 1 |
| **Cell Morphology,Cellular Assembly and Organization,Cellular Development,Cellular Function and Maintenance,Cellular Growth and Proliferation,Nervous System Development and Function,Organismal Development,Tissue Development** | Sprouting of rod bipolar cells | 0.0332 | ERC2 | 1 |
| **Cellular Development,Nervous System Development and Function,Tissue Development** | Differentiation of pyramidal neurons | 0.0332 | UNC5D | 1 |
| **Inflammatory Response** | Inflammatory response of cerebral cortex | 0.0332 | KNG1 | 1 |
| **Cellular Development** | Delay in initiation of transition of radial glial cells | 0.0332 | PLAGL1 | 1 |
| **Cellular Development** | Delay in initiation of transition of neuronal progenitor cells | 0.0332 | PLAGL1 | 1 |
| **Hereditary Disorder,Neurological Disease,Organismal Injury and Abnormalities,Psychological Disorders,Skeletal and Muscular Disorders** | Huntington disease grade 1 Huntington disease | 0.0332 | SNAP25 | 1 |
| **Embryonic Development,Nervous System Development and Function,Organ Development,Organismal Development,Tissue Development** | Development of septal area | 0.0332 | VAX1 | 1 |
| **Behavior** | Social exploration | 0.0332 | APBA2 | 1 |
| **Hereditary Disorder,Neurological Disease,Organismal Injury and** | Parkinson-dementia syndrome | 0.0332 | GAP43 | 1 |

| | | | | |
|---|---|---|---|---|
| **Abnormalities,Psychological Disorders,Skeletal and Muscular Disorders** | | | | |
| **Cardiovascular System Development and Function,Organismal Development** | Penetration by blood vessel | 0.0332 | ADGRA2 | 1 |
| **Developmental Disorder,Endocrine System Disorders,Nutritional Disease,Organismal Injury and Abnormalities,Reproductive System Disease** | Delayed puberty | 0.0332 | VAX1 | 1 |
| **Cell-To-Cell Signaling and Interaction,Nervous System Development and Function** | Sensitization of trigeminovascular neuron | 0.0332 | KNG1 | 1 |
| **Cell Morphology,Nervous System Development and Function,Organ Morphology,Organismal Development,Tissue Morphology** | Size of anterior horn cells | 0.0332 | SMN1/SMN2 | 1 |
| **Neurological Disease,Organismal Injury and Abnormalities** | Pathogenesis of motor neurons | 0.0332 | SMN1/SMN2 | 1 |
| **Cellular Assembly and Organization,Cellular Compromise,Nervous System Development and Function** | Retraction of glial cell projections | 0.0332 | GRIA2 | 1 |
| **Cell Death and Survival** | Delay in cell death of motor neurons | 0.0332 | SMN1/SMN2 | 1 |
| **Behavior** | Perseverance behavior | 0.0332 | NR3C2 | 1 |
| **Cell Death and Survival,Cell Morphology,Nervous System Development and Function,Neurological Disease,Tissue Morphology** | Loss of alpha-motor neurons | 0.0332 | SMN1/SMN2 | 1 |
| **Cellular Development,Cellular Growth and Proliferation,Nervous System Development and Function,Tissue Development** | Development of corticospinal neurons | 0.0332 | BCL11B | 1 |
| **Behavior** | Impulsivity | 0.0332 | ADCY1 | 1 |
| **Cell-To-Cell Signaling and Interaction,Nervous System Development and Function** | Hyperactivation of neurons | 0.0332 | SLC24A4 | 1 |
| **Metabolic Disease,Neurological Disease,Organismal Injury and Abnormalities,Psychological Disorders** | Preclinical stage Alzheimer disease | 0.0332 | NOS2 | 1 |
| **Hematological Disease** | Hyperemia | 0.0332 | PTGS2 | 1 |
| **Nervous System Development and Function,Tissue Morphology** | Density of climbing fiber | 0.0332 | DAB2IP | 1 |
| **Embryonic Development,Nervous System Development and Function,Organ Development,Organismal Development,Tissue Development** | Development of layer IV | 0.0332 | GAP43 | 1 |
| **Endocrine System Development and Function,Endocrine System Disorders,Nervous System Development and Function,Neurological Disease,Organ Morphology,Organismal Development,Organismal Injury and Abnormalities,Reproductive System Disease** | Abnormal morphology of enlarged adenohypophysis | 0.0332 | TCF7L2 | 1 |
| **Cellular Assembly and Organization** | Delay in initiation of priming of vesicles | 0.0332 | SNAP25 | 1 |
| **Cell-To-Cell Signaling and Interaction,Drug Metabolism,Small Molecule Biochemistry** | Conversion of norepinephrine | 0.0332 | DBH | 1 |
| **Infectious Diseases** | Replication of Murine cytomegalovirus | 0.0332 | NOS2 | 1 |
| **Behavior** | Risk preference | 0.0332 | GABRB3 | 1 |
| **Metabolic Disease,Neurological Disease,Organismal Injury and Abnormalities,Psychological Disorders** | Stage V Alzheimer disease | 0.0332 | PTGS2 | 1 |
| **Cell-To-Cell Signaling and Interaction,Nervous System Development and Function** | Activation of mitral cells | 0.0332 | GJD2 | 1 |
| **Cell Morphology,Cellular Assembly and Organization,Nervous System Development and Function** | Projection of mossy fibers | 0.0332 | NR3C2 | 1 |
| **Cell Morphology,Cellular Assembly and Organization** | Orientation of plasma membrane projections | 0.0332 | SCN1B | 1 |
| **Cell-To-Cell Signaling and Interaction,Nervous System Development and Function** | Excitation of postsynaptic region | 0.0332 | PCDH8 | 1 |
| **Cellular Assembly and Organization,Cellular Development,Nervous System Development and Function,Tissue Development** | Differentiation of presynaptic terminals | 0.0332 | AGRN | 1 |
| **Cell-To-Cell Signaling and Interaction,Drug Metabolism,Small Molecule Biochemistry** | Catabolism of dopamine | 0.0332 | PRKN | 1 |

| | | | | |
|---|---|---|---|---|
| **Nervous System Development and Function,Neurological Disease,Organ Morphology,Organismal Development,Organismal Injury and Abnormalities** | Lack of optic tract | 0.0332 | GAP43 | 1 |
| **Cell Morphology,Cellular Assembly and Organization,Nervous System Development and Function,Tissue Morphology** | Diameter of nodes of Ranvier | 0.0332 | SPTBN4 | 1 |
| **Cellular Development,Nervous System Development and Function,Tissue Development** | Differentiation of postsynaptic neurons | 0.0332 | AGRN | 1 |
| **Cell-To-Cell Signaling and Interaction** | Restoration of synapse | 0.0332 | SMN1/SMN2 | 1 |
| **Nervous System Development and Function** | Discharge of cochlear nerve | 0.0332 | BSN | 1 |
| **Cardiovascular System Development and Function,Nervous System Development and Function,Organ Morphology,Organismal Development,Tissue Morphology** | Diameter of basilar artery | 0.0332 | KNG1 | 1 |
| **Cell Morphology,Cellular Movement,Nervous System Development and Function** | Innervation of serotonergic neurons | 0.0332 | GAP43 | 1 |
| **Cell-To-Cell Signaling and Interaction,Nervous System Development and Function** | Firing of trigeminovascular neuron | 0.0332 | KNG1 | 1 |
| **Cell Morphology,Cellular Assembly and Organization** | Elongation of astrocyte projections | 0.0332 | GAP43 | 1 |
| **Nervous System Development and Function,Neurological Disease,Organ Morphology,Organismal Development,Organismal Injury and Abnormalities** | Abnormal morphology of cerebellar hemisphere | 0.0332 | AHI1 | 1 |
| **Lipid Metabolism,Molecular Transport,Small Molecule Biochemistry** | Concentration of eicosapentenoic acid | 0.0332 | ARNTL | 1 |
| **Cell-To-Cell Signaling and Interaction,Cellular Function and Maintenance,Nervous System Development and Function** | Paired-pulse facilitation of parallel fiber-Purkinje cell synapses | 0.0332 | UNC13C | 1 |
| **Metabolic Disease,Neurological Disease,Organismal Injury and Abnormalities,Psychological Disorders** | Stage I Alzheimer disease | 0.0332 | PTGS2 | 1 |
| **Cell Morphology,Cellular Movement,Nervous System Development and Function** | Innervation of neurons | 0.0332 | EPHB3, GABRB3, GAP43, GRIA2, RIMS2, TENM2, TOP2B | 7 |
| **Cell-To-Cell Signaling and Interaction,Nervous System Development and Function** | Synaptic depression of collateral synapses | 0.034 | ADCY1, RAB11FIP5 | 2 |
| **Cell-To-Cell Signaling and Interaction,Nervous System Development and Function** | Long-term potentiation of mossy fibers | 0.034 | ADCY1, GRIA2 | 2 |
| **Amino Acid Metabolism,Drug Metabolism,Endocrine System Development and Function,Molecular Transport,Small Molecule Biochemistry** | Concentration of levothyroxine | 0.034 | NCOR1, SLCO1C1 | 2 |
| **Cellular Assembly and Organization** | Quantity of mitochondria | 0.034 | AKT3, PRKN | 2 |
| **Nervous System Development and Function,Neurological Disease,Organ Morphology,Organismal Development,Organismal Injury and Abnormalities** | Abnormal morphology of anterior horn of the spinal cord | 0.034 | KIF1B, RIMS2 | 2 |
| **Nervous System Development and Function,Neurological Disease,Organ Morphology,Organismal Development,Organismal Injury and Abnormalities** | Abnormal morphology of optic tract | 0.034 | EPHB3, GAP43 | 2 |
| **Cellular Compromise,Neurological Disease,Organismal Injury and Abnormalities** | Neurodegeneration of hippocampal cells | 0.0351 | GRIA2, PRKN, PTGS2 | 3 |
| **Cell-To-Cell Signaling and Interaction,Drug Metabolism,Molecular Transport,Small Molecule Biochemistry** | Concentration of norepinephrine | 0.0351 | DBH, GAP43, PRKN | 3 |

| | | | | |
|---|---|---|---|---|
| **Nervous System Development and Function,Neurological Disease,Organ Morphology,Organismal Development,Organismal Injury and Abnormalities** | Abnormal morphology of cerebral cortex | 0.0365 | ADCY1, ASPM, CRMP1, DBN1, FES, GAP43, GRIA2, HIVEP2, KIF1B, LRRTM4, MAN2C1, MCPH1, SNAP25 | 13 |
| **Embryonic Development,Nervous System Development and Function,Organ Development,Organismal Development,Tissue Development** | Formation of brain | 0.0365 | ABR, ADGRA2, AHI1, AKT3, ARNT2, ASPM, BRCA2, CDK5RAP1, CELSR2, ELP1, EPHB3, GAP43, GRHL2, LRRK2, MCPH1, MRTFA, MYO16, NCOR1, NIN, NME7, NOS2, PITPNM1, PROKR2, RBFOX2, RTP1, SCRIB, SETD2, TACC2, TOP2B, VAX1 | 30 |
| **Cell Morphology,Cellular Assembly and Organization,Cellular Development,Cellular Function and Maintenance,Cellular Growth and Proliferation,Embryonic Development,Nervous System Development and Function,Organismal Development,Tissue Development,Tissue Morphology** | Density of dendritic spines | 0.0386 | CRMP1, ELK1, GABRB3, PTGS2, RYR2, TIAM1, TSC2 | 7 |
| **Nervous System Development and Function,Neurological Disease,Ophthalmic Disease** | Abnormal morphology of optic nerve | 0.0394 | SCN1B, SPTBN4, VAX1 | 3 |
| **Cell-To-Cell Signaling and Interaction,Nervous System Development and Function** | Activation of synapse | 0.0415 | APBA2, GRIA2 | 2 |
| **Nervous System Development and Function,Tissue Morphology** | Density of axons | 0.0415 | ACVR1C, DAB2IP | 2 |
| **Cellular Assembly and Organization,Cellular Function and Maintenance,Cellular Movement,Nervous System Development and Function** | Axonal transport of mitochondria | 0.0415 | PRKN, RANBP9 | 2 |
| **Embryonic Development,Nervous System Development and Function,Organ Development,Organismal Development,Tissue Development,Visual System Development and Function** | Formation of eye | 0.0415 | ADCY1, FAT3 | 2 |
| **Cell-To-Cell Signaling and Interaction,Cellular Assembly and Organization,Cellular Development,Cellular Function and Maintenance,Cellular Growth and Proliferation,Nervous System Development and Function,Tissue Development** | Formation of excitatory synapses | 0.0415 | LRRTM4, SORBS2 | 2 |
| **Cell Morphology,Cellular Assembly and Organization,Cellular Development,Cellular Function and Maintenance,Cellular Growth and Proliferation,Nervous System Development and Function,Organismal Development,Tissue Development** | Axonogenesis | 0.0425 | ADCY1, AGRN, ARHGEF28, BCL11B, DVL1, EPHB3, | 15 |

| | | | | |
|---|---|---|---|---|
| | | | GAP43, NIN, NRN1, PIP5K1C, SMN1/SMN2, SPTBN4, TIAM1, TOP2B, TSC2 | |
| **Cellular Assembly and Organization,Nervous System Development and Function** | Recycling of synaptic vesicles | 0.0439 | DVL1, GAP43, PIP5K1C | 3 |
| **Embryonic Development,Nervous System Development and Function,Organ Development,Organismal Development,Tissue Development** | Formation of telencephalon | 0.0486 | ADGRA2, ELP1, EPHB3, GAP43, LRRK2, NIN, PROKR2, RTP1, TOP2B, VAX1 | 10 |
| **Cell Death and Survival,Cellular Compromise,Neurological Disease,Organismal Injury and Abnormalities,Tissue Morphology** | Neurodegeneration of pyramidal neurons | 0.0497 | GRIA2, PTGS2 | 2 |
| **Neurological Disease,Organismal Injury and Abnormalities,Psychological Disorders,Skeletal and Muscular Disorders** | Parkinson disease with dementia | 0.0497 | GAP43, NOS2 | 2 |
